# SRC-mediated and TKS5-enabled podosome formation is an inherent property of IPF fibroblasts, promoting ECM invasion and pulmonary fibrosis

**DOI:** 10.1101/2023.01.25.522705

**Authors:** Ilianna Barbayianni, Paraskevi Kanellopoulou, Dionysios Fanidis, Eleftheria-Dimitra Ntouskou, Dimitris Nastos, Apostolos Galaris, Vaggelis Harokopos, Pantelis Hatzis, Eliza Tsitoura, Robert Homer, Naftali Kaminski, Katerina M. Antoniou, Bruno Crestani, Argyrios Tzouvelekis, Vassilis Aidinis

**Author notes:** Equal contribution (Ilianna Barbayianni, Paraskevi Kanellopoulou).

## Abstract

The activation and accumulation of lung fibroblasts (LFs), resulting in aberrant deposition of collagens and other extracellular matrix (ECM) components, is a pathogenic hallmark of Idiopathic Pulmonary Fibrosis (IPF), a lethal and incurable disease. In this report, increased expression of TKS5, a scaffold protein essential for the formation of podosomes, was detected in the lung tissue of IPF patients and bleomycin (BLM)-treated mice, correlating with increased collagen type I alpha 1 chain (COL1A1) expression. The profibrotic milieu, TGFβ, as well as a stiff Col1a1-rich acellular fibrotic ECM, were found to induce *TKS5* expression and the formation of prominent podosome rosettes in LFs, culminating in increased ECM invasion. Podosomes were retained *ex vivo* in the absence of any stimulation, indicating that the formation of TKS5-enabled podosomes is an inherent property of IPF LFs. Remarkably, haploinsufficient *Tks5*^+/-^ mice were relatively resistant to BLM-induced pulmonary fibrosis. Disease protection was largely attributable to diminished podosome formation in LFs and decreased ECM invasion, thus indicating TKS5-enabled and podosome-mediated ECM invasion as a major pathogenic mechanism in pulmonary fibrosis. Expression profiling revealed an ECM-podosome cross talk, and pharmacologic connectivity map analysis suggested several inhibitors that could prevent podosome formation and thus pulmonary fibrosis. Among them, inhibition of src kinase was shown to potently attenuate podosome formation in LFs, ECM invasion, as well as pulmonary fibrosis in post BLM precision cut lung slices, suggesting that pharmacological targeting of TKS5-enabled podosome formation is a very promising therapeutic option in pulmonary fibrosis.

## Introduction

Tissue fibrosis is a pathogenic process that affects most organs and constitutes a complication of many chronic diseases including cancer; such fibroproliferative disorders account for >45% of all disease-related deaths worldwide^1^. Among them, Idiopathic pulmonary fibrosis (IPF) is a chronic, progressive, interstitial lung disease affecting mostly older adults. IPF patients exhibit progressive worsening of respiratory functions, which lead to dyspnea and eventually to respiratory failure. Histologically, IPF is characterized by lung parenchymal scarring, as evident by lung honeycombing, and a usual interstitial pneumonia (UIP) profile, distinguished by the presence of fibroblast foci^2^. Although the etiopathogenesis of IPF remains largely elusive, the prevailing hypothesis suggests that the mechanisms driving IPF involve age-related aberrant recapitulation of developmental programs and reflect abnormal, deregulated wound healing in response to persistent alveolar epithelial damage, resulting in the accumulation of lung fibroblasts^3^.

Lung fibroblasts (LFs) are the main effector cells in pulmonary fibrosis, secreting exuberant amounts of extracellular matrix (ECM) components, such as different types of collagens. LFs also secrete a variety of ECM remodeling enzymes, such as matrix metalloproteinases (MMPs), thus coordinating the overall ECM structural organization and consequently the mechanical properties of the lung^4^. LF activation upon fibrogenic cues, such as TGFβ or other growth factors (including PDGF and VEGF), is characterized by the expression of alpha smooth muscle actin (αSMA/ACTA2), and/or increased collagen expression, as exemplified by COL1A1^5,6^. ECM fibrotic remodeling and resulting mechanical cues are considered as crucial stimulating and perpetuating factors for LF activation^6,7^, while the chemoattraction of LFs to various signals and their resistance to apoptosis has been suggested to promote respectively their recruitment and accumulation^5,6^. Fibroblast accumulation in pulmonary fibrosis has also been suggested to be mediated by their ability to invade the underlying ECM, and increased ECM invasion of fibroblasts isolated from the lung tissue of IPF patients or animal models has been reported^8-11^. Activation of invasion, critical for embryonic development, is among the well-established hallmarks of cancer^12^, and one of the many shared hallmarks between cancer cells and activated LFs^13^. Invasion critically relies on the proteolysis of the underlying ECM via invadopodia in cancer cells and podosomes in other cell types^14,15^. Podosomes are comprised of a filamentous (F)-actin-rich core enriched in actin-regulating proteins, such as the Arp2/3 complex and cortactin (CTTN), and are surrounded by a ring of scaffold proteins, most notably SH3 and PX domains 2A (SH3PXD2A; commonly known as tyrosine kinase substrate with 5 SH3 domains, TKS5)^14-16^.

*Tks5* expression is necessary for neural crest cell migration during embryonic development in zebrafish^17^, and homozygous disruption of *Tks5* in mice resulted in neonatal death^18^. Beyond embryonic development, which heavily relies on migration and invasion, increased TKS5 expression has been reported in different types of cancers^14-16^, including lung adenocarcinoma, where it was suggested to mediate metastatic invasion^19^. Pulmonary fibrosis confers one of the highest risks for lung cancer development, while many similarities between activated LFs and cancer cells have been suggested, including ECM invasion^13^. Therefore, in this report we investigated a possible role of TKS5 and podosomes in the pathogenesis of pulmonary fibrosis.

## Results

### Increased *TKS5* expression in pulmonary fibrosis

Increased *TKS5* mRNA levels were detected *in silico* in the lung tissue of IPF patients as compared with control samples (Fig. 1A), in most publicly available IPF transcriptomic datasets (Table S1) at Fibromine^20^, including three of the largest ones (Figs. 1B and S1A, C). Importantly, *TKS5* mRNA expression in fibrotic lungs correlated with the expression of *COL1A1* (Figs. 1C and S1B, D), a well-established marker of fibrotic gene expression. Confirming the *in silico* results, increased *TKS5* mRNA levels were detected with Q-RT-PCR in lung tissue isolated from IPF patients (n=20), as compared with COPD patients (n=19) and healthy lung tissue (n=9) (Table S2; Fig. 1D). Moreover, positive TKS5 immunostaining was detected in the lungs of IPF/UIP patients (n=3), as opposed to control samples (n=3), mainly localized in the alveolar epithelium and fibrotic areas (Figs. 1E, S2). Similar conclusions were derived from the analysis of a publicly available single cell RNA sequencing (scRNAseq) dataset of lung tissue from transplant recipients with pulmonary fibrosis (n=4) and healthy lung tissue from transplant donors (n=8)^21^. *TKS5* mRNA expression was mostly detected in subsets of epithelial cells, basal cells and especially fibroblasts (Fig. S1E-F), where TKS5-expressing LFs were found to belong to a *COL1A1*-expressing subpopulation (Fig. S1G, 1F).

**Figure 1.**
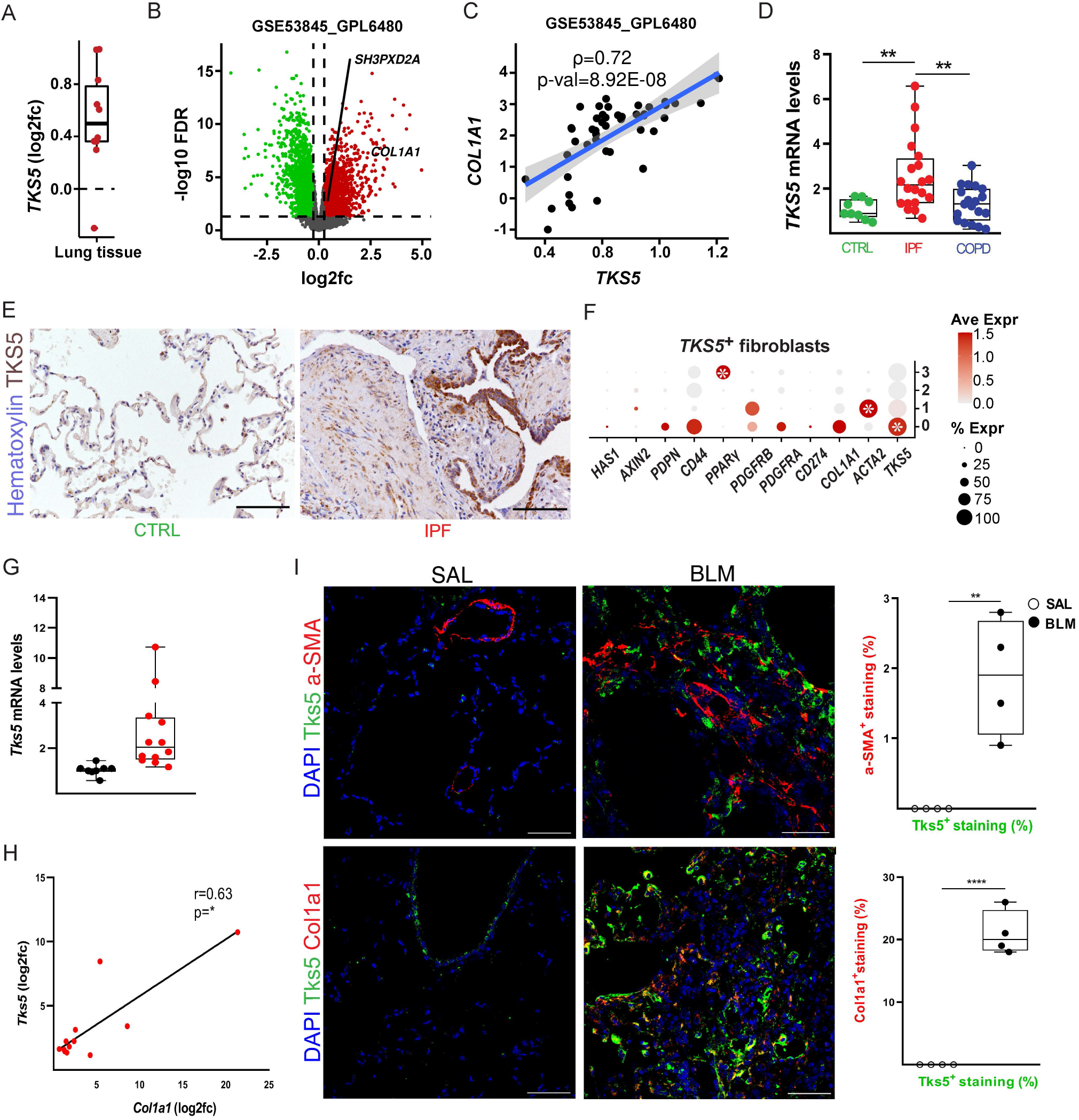
Increased *TKS5* expression in pulmonary fibrosis. **A**. *TKS5* mRNA expression in lung tissue from IPF patients as compared (log2FC) to controls (CTRL) in different publicly available datasets (Table S1) at Fibromine. **B**. Volcano plot from a representative large dataset (FC>1.2, FDR<0.05). **C**. Spearman correlation plot of *TKS5* and *COL1A1* expression in the dataset of B (rho>0.6, p<0.05). **D**. Increased *TKS5* mRNA levels in the lung tissue of IPF (UIP) patients were detected with Q-RT-PCR (r2=0.98, E=97%), as compared with lung tissue from COPD patients and control (CTRL) lung tissue isolated from lung cancer patients (Table S2). Values were normalized to the expression values of the housekeeping gene *B2M* and presented as fold change to CTRL values. Statistical significance was assessed with one-way ANOVA, followed by Holm-Sidak’s multiple comparison test; **denotes p<0.01. **E**. Increased TKS5 immunostaining in fibrotic lungs. Representative images from immunohistochemistry for TKS5 in IPF and CTRL lung tissue (n=3; Fig. S2); scale bars=50 μm. **F**. *TKS5* is expressed mainly by a *COL1A1*-expressing cluster/LF subpopulation. (^*^FC>1.2, Bonferroni corrected p<0.05) in a publicly available scRNAseq dataset (Reyfman, Walter et al. 2019). **G-H**. *Tks5* and *Col1a1* mRNA expression was interrogated with Q-RT-PCR (r^2^=0.89/0.93; E=103%/96%); cumulative result from 3 different experiments. Values were normalized over the expression of *B2m* and presented as fold change (log2) over control. Statistical significance in G was assessed with Mann-Whitney test; ^***^denotes p<0.001. Statistical significance in H was assessed with Spearman r=0.63; ^*^denotes p<0.05 **I**. Double immunostaining against Tks5 and aSMA (Acta2) or Col1a1; representative images are shown, followed by their respective quantification with Image J; statistical significance was assessed with unpaired t-test; ^*/**/***^ denote p<0.05/0.01/0.001 respectively; scale bars=50 μm; representative experiment out of 3 independent ones.

Increased *Tks5* mRNA levels, correlating with *Col1a1* mRNA levels, were also detected in the lung tissue of mice post bleomycin (BLM) administration (Fig. 1G-H), a widely used animal model of pulmonary fibrosis ^22-24^; immunostaining localized Tks5 in the alveolar epithelium and fibrotic areas (Fig. 1I), as in human patients. Moreover, double immunostaining for aSMA or Col1a1, prominent activation markers of fibroblasts in both mice and humans, indicated that Tks5 localized mainly to a Col1a1 expressing fibroblast subset (Fig. 1I).

Therefore, pulmonary fibrosis in both humans and mice is associated with increased *TKS5* expression, consistently correlated with the expression of COL1A1, especially in LFs.

### TGFβ-induced podosome rosettes is an inherent property of fibrotic LFs

TGFβ, among the main pro-fibrotic factors driving disease development *in vivo*, was found to stimulate *TKS5* mRNA expression in different primary normal human lung fibroblast (NHLF) clones (Fig. 2A), correlating with *COL1A1* mRNA expression (Fig. 2B); identical results were obtained from an independently derived NHLF cell line at a different lab/setting (Fig. S3 A, B), as well as from the human fibroblastic cell line MRC5 (Fig. S3 C, D). In agreement with the essential role of TKS5 on podosome formation^25^, TGFβ potently stimulated the formation of podosomes in NHLFs *in vitro*, organized in distinctive rosettes (Figs. 2C-F and S4). Moreover, TGFβ-induced podosomes in LFs were enriched in MMP9 (Fig. 2G), likely contributing to the increased degradation of a fluorescein-conjugated gelatin substrate (Fig. 2H), a nominal podosome property.

**Figure 2.**
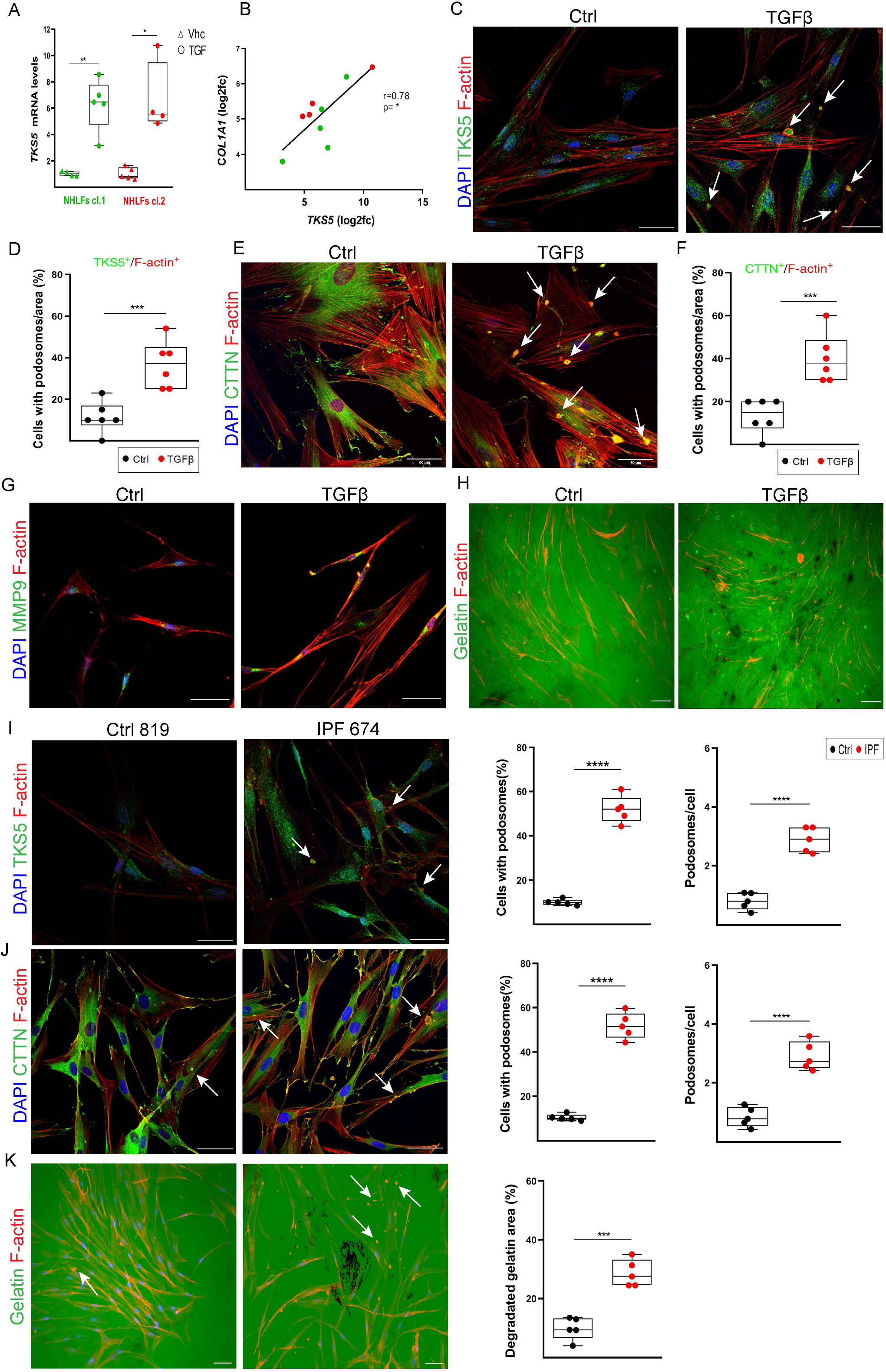
TGFβ-induced podosome rosettes is an inherent property of fibrotic LFs. Serum starved, sub-confluent (70-80%), primary NHLFs were stimulated with recombinant human TGFβ (10 ng/ml) for 24h. Scale bars=50 μm; representative experiment out of 4 independent ones. **A-B**. *TKS5* and *COL1A1* mRNA expression was interrogated with Q-RT-PCR (r^2^=0,94/0,92; E=98,3%/93% respectively). Values were normalized to the expression values of the housekeeping gene *B2M* and presented as fold change over control; statistical significance was assessed with unpaired t-test (cl.1) followed by Welch’s correction, or Mann-Whitney test (cl.2); ^*/**^denote p<0.05/0.01 respectively. **B**. Pearson correlation plot of *COL1A1* expression in the same samples (r=0.78 and R2=0.62); ^*^denote p<0.05. **C, E**. Representative composite images from double immunostaining for F-actin and TKS5 (C) or Cortactin (CTTN; E) counter stained with DAPI; arrows indicate representative podosomes; separate images and proof of colocalization of signals can be found at Figure S4. **D, F**. Quantification of the number of podosome-containing cells per optical field (x6); statistical significance was assessed with unpaired t-test, followed by Welch’s correction; ^***^denotes p<0.001. **G**. Representative composite images from double immunostaining for F-actin and MMP-9, counter stained with DAPI. **H**. Representative images of the TGFβ-induced degradation (black holes) of a fluorescein-conjugated gelatin substrate by LFs. Serum starved, sub-confluent (70-80%), primary IPF-HLFs and NHLFs (n=5 each) were immunostained for F-actin and **I**. TKS5 or **J**. cortactin (CTTN) and counter stained with DAPI. Representative images from representative clones are shown, followed by their respective quantification of the number of podosome-containing cells and the number of podosomes per cell per optical field (x6); scale bars=50 μm; additional clones and controls at figure S5; ^****^denotes p<0.0001 **K**. The same clones were cultured on a fluorescein-conjugated gelatin substrate and were stained for F-actin and counter stained with DAPI; representative images are shown, followed by the quantification of the percentage of the degraded gelatin for all IPF and control clones cumulatively (means +/-SE), as quantified with ImageJ; statistical significance was assessed with unpaired t-test, followed by Welch’s correction; ^****^denotes p<0.0001. Representative images from representative clones are shown; scale bars=50 μm; additional clones and controls are shown at figure S5.

To examine if the pro-fibrotic milieu in the lungs of IPF patients, which includes TGFβ, also stimulate podosome formation *in vivo*, HLFs from IPF patients (Table S3) were cultured in the absence of any stimulation and were stained for podosomes in comparison, under the same conditions (and 7-8 passages), with different NHLF lines derived from healthy tissue. Remarkably, IPF HLFs presented with prominent podosome rosettes (Fig. 2I-J, Fig. S5A-B, Video S1), identical in structure as those stimulated *in vitro* by TGFβ, that persist upon prolonged culture *ex vivo* (7-8 passages). As shown for TGFβ-stimulated NHLFs, IPF HLFs degraded more potently than NHLFs a fluorescein-conjugated gelatin substrate (Fig. 2K; Fig. S5C).

Phenocopying the human experiments, exposure to TGFβ of primary, normal mouse lung fibroblasts (NMLFs) stimulated *Tks5* mRNA expression (Fig. S6A), correlating with *Col1a1* expression (Fig. S6B), the formation of podosome rosettes (Fig. S6C, D) and the degradation of a fluorescein-conjugated gelatin substrate (Fig. S6E-F); similar results were obtained with 3T3 embryonic fibroblasts (Fig. S6G-K). Moreover, and as in the case of IPF LFs, mouse primary LFs isolated post BLM administration presented with prominent podosome rosettes in the absence of any stimulation (Fig. S6L-N).

Therefore, the pro-fibrotic milieu in the lungs of IPF patients and BLM-treated mice, as well as TGFβ, induce TKS5 expression and the formation of podosome rosettes, an inherent fibrotic LF property.

### *Tks5* haploinsufficiency in mice attenuates BLM-induced pulmonary fibrosis

To genetically dissect the likely role of Tks5 in pulmonary fibrosis and pathophysiology in mice, we created a series of constitutive and conditional knock out mice for *Tks5* (*Sh3pxd2a*), as described in detail in the online supplement and outlined in Figure S7A. The haploinsufficient *Sh3pxd2a*^tm1b(EUCOMM)WtsiFlmg/+^ and *Sh3pxd2a*^tm1d/(EUCOMM)WtsiFlmg/+^ (*Tks5*^+/-^) mice, presented with a 50% reduction of *Tks5* mRNA levels in the lung (Fig. S7G-H), while X-gal staining (in the reporter tm1b strain) localized transcriptional *Tks5* activation (throughout development, neonatal and adult life) mainly in endothelial and smooth muscle cells in healthy conditions (Fig. S7I). No obvious gross macroscopic abnormalities were observed, while heterozygous mice were healthy and fertile. Intercrossing of heterozygous mice *Tks5*^+/-^ mice yielded no homozygous knockout mice, indicating that *Tks5* has an essential role in mouse development, as previously reported for an obligatory knock out strain ^18^.

Bleomycin (BLM) was then administered to 8-10-week-old C57Bl6/J *Tks5*^+/-^ mice and WT littermates (Fig. 3A,B), as previously described^23^. No weight loss, an overall systemic health indicator, was observed in *Tks5*^+/-^ mice (Fig. 3C), as opposed to wt mice. Vascular leak and pulmonary oedema were significantly reduced in *Tks5*^+/-^ mice, as indicated by the total protein concentration in the bronchoalveolar lavage fluid (BALF), determined with the Bradford assay (Fig. 3D). Inflammatory cells in the BALF, as measured by hematocytometer, were found significantly reduced in *Tks5*^+/-^ mice (Fig. 3E); so were soluble collagen BALF levels, as determined by the Sirius red assay (Fig. 3F), in concordance with *Col1a1* mRNA expression in the lung tissue from the same mice, as determined with Q-RT-PCR (Fig. 3G-H). Histological analysis revealed decreased collagen deposition in *Tks5*^+/-^ mice post BLM, as quantified by Sirius red/Fast green staining (Fig. 3I), and fewer peribronchiolar and parenchymal fibrotic regions were detected (Fig. 3I), as reflected in the Ashcroft score (Fig. 3J); similar conclusions were drawn upon the histological evaluation of Precision Cut Lung Slices (PCLS) prepared from the same mice and cultured *ex vivo* (Fig. 3I). The relative protection from the BLM-induced tissue architecture distortion upon the genetic reduction of *Tks5* expression was also reflected in lung respiratory functions, as measured with FlexiVent (Fig. 3 K-M). Therefore, Tks5 expression, and likely the formation of podosomes, were shown to have a major role in BLM-induced pulmonary fibrosis, and therefore likely IPF.

**Figure 3.**
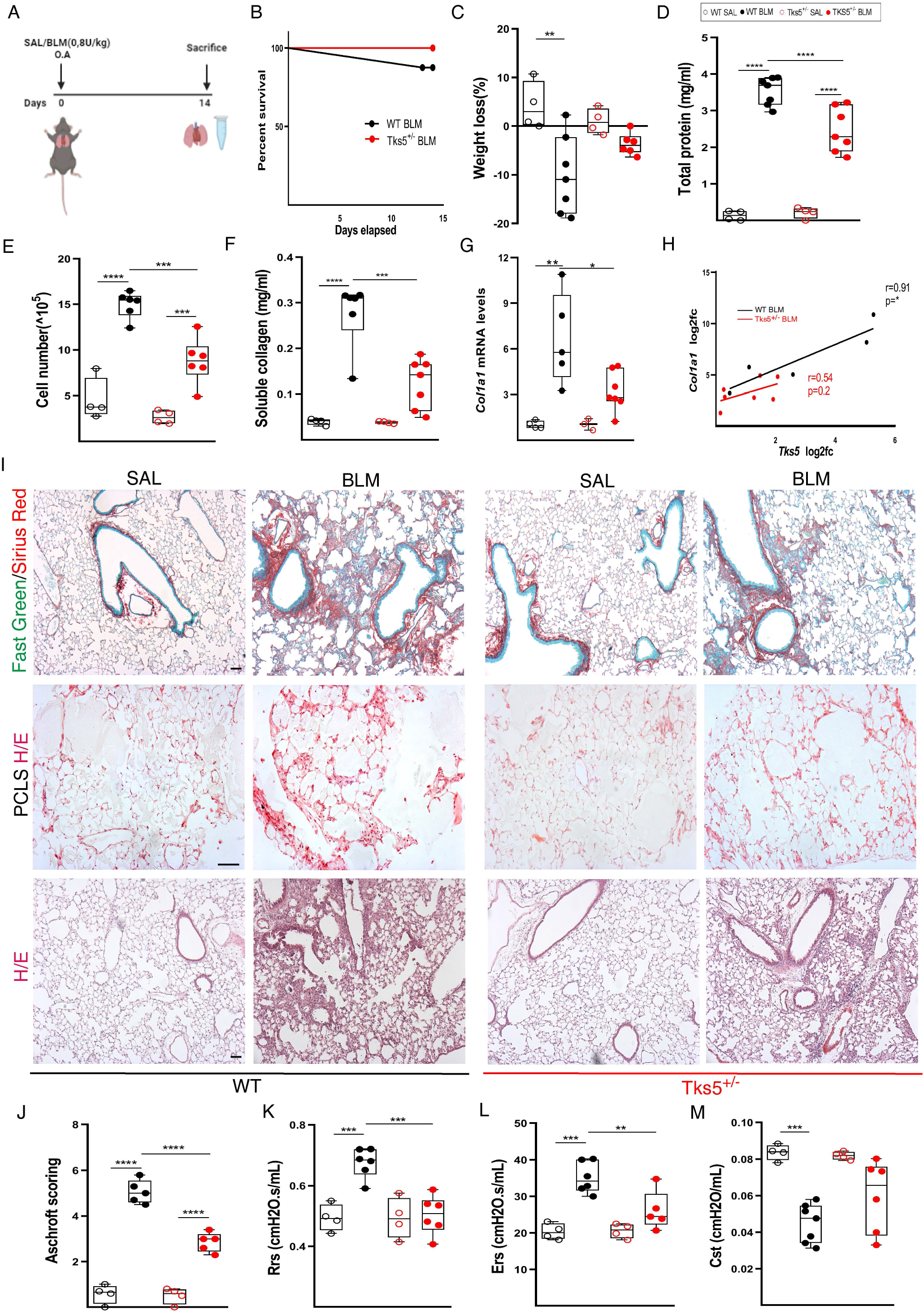
*Tks5* haploinsufficiency in mice attenuates BLM-induced pulmonary fibrosis. **A**. Schematic presentation of the BLM induced PF model. **B**. Kaplan Meyer survival curve post BLM administration. **C**. Weight change post BLM administration. **D**. Total protein concentration in BALFs, as determined with the Bradford assay. **E**. Inflammatory cell numbers in BALFs, as counted with a hematocytometer. **F. S**oluble collagen levels in the BALFs were detected with the direct red assay. **G-H**. *Tks5* and *Col1a1* mRNA expression was interrogated with Q-RT-PCR; Values were normalized over the expression of the housekeeping gene *B2m* and presented as fold change over control; Statistical significance was assessed with one-way anova, followed by Bonferroni post hoc correction, ^*/**^denote p<0.05/0.01; **H**. Pearson correlation plot of Col1a1 expression. Statistical significance was assessed with Pearson r= 0.91/0.54 and as indicated. **I**. Representative images from lung sections of murine lungs of the indicated genotypes, stained with Fast Green/Sirius Red (first row), from H&E-stained Precision cut lung slices (PCLS) (second row) and H&E-stained lung sections (third row). **J**. Quantification of fibrosis severity via Ashcroft scoring. **K-M** Respiratory functions were measured with FlexiVent. Statistical significance was assessed with one-way ANOVA, followed by Tukey’s correction; cumulative result from 2 different experiments; ^*/**/***^ denote p<0.05/0.01/0.001 respectively.

### *Tks5* haploinsufficiency in mouse LFs decreases ECM-regulated podosome formation and ECM invasion

To functionally dissect the relative protection of *Tks5*^+/-^ mice from BLM-induced pulmonary fibrosis, primary LFs were isolated from littermate wt and *Tks5*^+/-^ mice and were exposed to TGFβ, as before. *Tks5*^+/-^ LFs, expressing ∼50% of Tks5 (Fig. 4A), presented with decreased numbers of podosomes in response to TGFβ (Fig. 4B, C), reaffirming the seminal role of Tks5 in podosome formation^25^, as well as with decreased proliferation (24h; Fig. 4D). As podosomes are known to promote ECM invasion, we then examined the ability of LFs to invade acellular ECM (aECM) prepared from the lungs of mice (Fig. S8 A, B), in a transwell invasion chamber (6h; Fig. 4E). The reduction of podosomes was associated with a decreased TGFβ-induced invasion of *Tks5*^+/-^ LFs in aECM (Fig. 4F). Moreover, reaffirming in mice the inherent character of podosome formation in LFs, post BLM *Tks5*^+/-^ LFs presented with reduced numbers of podosomes in comparison with wt LFs isolated from littermate mice (Fig. 4G-H), resulting in defective aECM invasion (Fig. 4I). Therefore, the *in vivo* demonstrated pathogenic role of Tks5 in pulmonary fibrosis include the formation of podosomes in LFs and the promotion of their ECM invasion.

**Figure 4.**
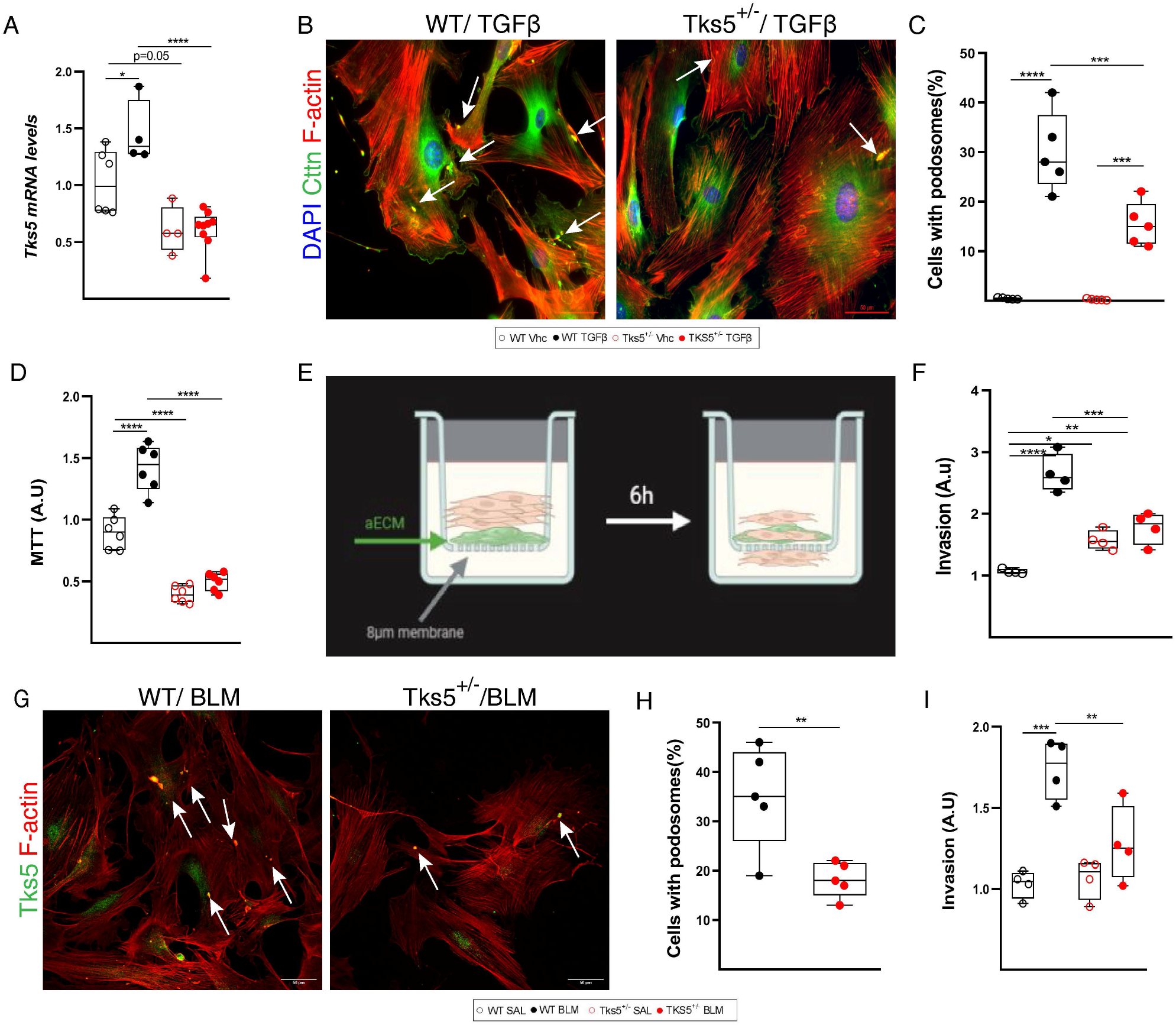
Tks5 haploinsufficiency in mouse LFs decreases podosome formation and ECM invasion. Serum starved primary NMLFs from WT and Tks5^+/-^ mice were stimulated with recombinant TGF-β1 (10 ng/ml for 24 h); ^*/**/***/****^ denote p<0.05/0.01/0.001/0.0001 respectively. **A**. *Tks5* mRNA expression was interrogated with Q-RT-PCR. Values were normalized over the expression of the housekeeping gene *B2m* and presented as fold change over control; statistical significance was assessed with one-way anova, followed by Tukey’s correction. **B**. Representative composite images from double immunostaining for F-actin and Tks5 counter stained with DAPI; arrows indicate representative podosomes. **C**. Quantification of the number of podosome-containing cells per optical field (x5). Statistical significance was assessed with one-way anova. **D**. TGFβ-induced NMLFs proliferation was assessed with the MTT assay. Statistical significance was assessed with one-way anova, followed by Tukey’s correction. **E**. Schematic presentation of LFs invasion into aECM, upon TGFB stimulation. After 6 h, cells that had invaded into the lower surface of the upper chamber were stained, lysed and absorbance values were measured **F**. Invasion capacity of LFs, upon TGF-β stimulation, as detected with the transwell invasion assay. Statistical significance was assessed with one-way anova, followed by Tukey’s correction; ^*/**/***^denote p<0.05/0.01/0.001 respectively. **G**. Representative composite images from double immunostaining for F-actin and Tks5 in NMLFs isolated from WT and *Tks5*^+/-^ mice, post BLM administration; arrows indicate representative podosomes **H**. Quantification of the number of podosome-containing cells per optical field (x5). Statistical significance was assessed as in C. **I**. Invasion capacity of LFs, post BLM, as detected with the transwell invasion assay. Statistical significance was assessed as in F.

To obtain additional mechanistic insights, wt and *Tks5*^+/-^ LFs were exposed to TGFβ, as above, and their global expression profile was interrogated with 3’ UTR RNA sequencing (Quant-Seq LEXOGEN). Differential expression analysis between TGFβ-induced *Tks5*^+/-^ and wt LFs, revealed 3648 differentially expressed genes (DEGs; FC>1.2, FDR corr. p<0.05) (Table S4; Fig. S9A); among them 418 DEGs have been previously associated with pulmonary fibrosis, as detected with text mining of abstract co-occurrence of identified DEGs with fibrosis keywords (Table S4).

*Stat1, Cebpa* and *Ar* transcription factors (TFs), where found downregulated in *Tks5*^+/-^ LFs along with several of their target genes (Table S4, Fig S9B). Gene set enrichment analysis (GSEA) performed on DEGs revealed that the most affected cellular components (CC), molecular functions (MF) and biological processes (BP) all relate to the ECM (Fig. 5A, Table S5). “Collagen containing ECM” (GO:0062023) was most prominent (Fig. 5B) due to the down regulation of several ECM related genes such as collagens and MMPs/TIMPS/Adamts (Fig. 5B, S9C), In this context and given the observed consistent correlation of *Tks5* and *Col1a1* expression, *Tks5*^+/-^ LFs post BLM, containing fewer podosomes and exhibiting defective aECM invasion (Fig. 4G-I), were found to produce significantly less Col1a1 (Fig. 5C). *Vice versa*, culture of primary NMLFs on Col1a1-rich aECM prepared from the lungs of mice post BLM (Fig. S8C), stimulated *Tks5* expression (Fig. 5D-E) and the formation of podosomes (Fig. 5F-G), and further stimulated *Col1a1* expression (Fig. 5H), indicating an ECM-podosome cross talk in the perpetuation of LF activation.

**Figure 5.**
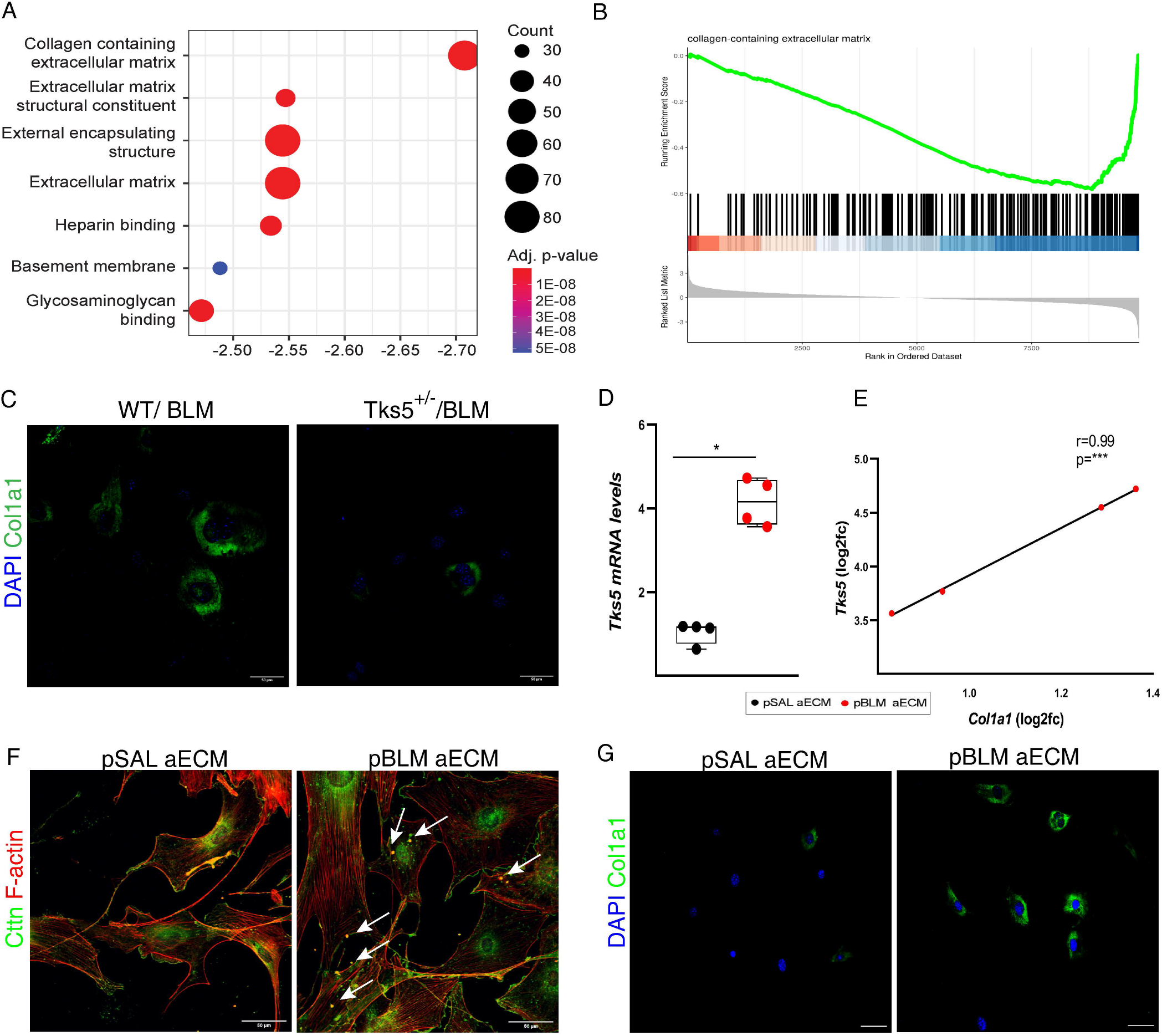
Tks5 haploinsufficiency in mouse LFs disrupts ECM homeostasis, that critically controls podosome formation and ECM invasion. **A**. ECM-related gene ontology components are enriched for genes down-regulated in TGFβ stimulated Tks5^+/-^ LFs compared to their WT TGFβ treated counterparts. **B**. Collagen containing extracellular matrix is the term most enriched in down-regulated genes according to gene-set enrichment analysis (GSEA). **C**. Serum starved WT and Tks5^+/-^ LFs were immunostained for Col1a1 and counter stained with DAPI. Representative images are shown. **D-E**. Serum starved WT primary LFs were cultured in pSAL and pBLM aECM. *Tks5* and *Col1a1* mRNA expression was interrogated with Q-RT-PCR; values were normalized over the expression of the housekeeping gene *B2m* and presented as fold change over control; statistical significance was assessed with unpaired t-test; */***denote p<0.05/0.001.**E**. Pearson correlation plot of *Col1a1* expression in the same samples (r=0.99). **F, G**. Representative composite images from double immunostaining for F-actin and Cttn (F) or Col1a1(G) counter stained with DAPI; arrows indicate representative podosomes; scale bars=50 μm.

### Src-inhibition potently reduces podosome formation and attenuate pulmonary fibrosis

To identify pharmaceutical compounds that can induce a similar transcriptional profile as that of the defective in ECM invasion *Tks5*^+/-^ LFs, the TGFβ-induced *Tks5*^+/-^ LFs profile was queried against the connectivity map (CMap) <u>LINCSL1000</u> database (Fig. 6A), a public resource that contains >10^6^ gene expression signatures of different cell types treated with a large variety of small molecule compounds^26^. Among the identified compounds with similar expression signatures, several have already been shown to have a positive effect in disease pathogenesis in animal models (Fig. 6A, Table S6). The identified possible therapeutic targets include the PDGF and VEGF receptors, which are pharmacologically targeted by the current IPF standard of care (SOC) compound nintedanib^27^. More importantly, the list also includes an inhibitor of Src, a TGFβ/PDGF-inducible, non-receptor tyrosine kinase essential for podosome formation^28^. To verify the *in silico* findings in our experimental settings, TGFβ-activated NHLFs were incubated with nontoxic, increasing concentrations of nintedanib and A-419259, a commercially available src inhibitor. Both nintedanib but especially A-419259 reduced both *TKS5* and *COL1A1* expression (Fig. 6B-E), as well as podosome formation (Fig. 6F-G). Moreover, the same src inhibitor reduced aECM LF invasion (Fig. 6H) and reversed fibrosis in fibrotic mouse PCLS (Fig. 6I). Therefore, the TKS5-mediated podosome formation is a druggable pathologic process, which can be potently targeted by src inhibition.

**Figure 6.**
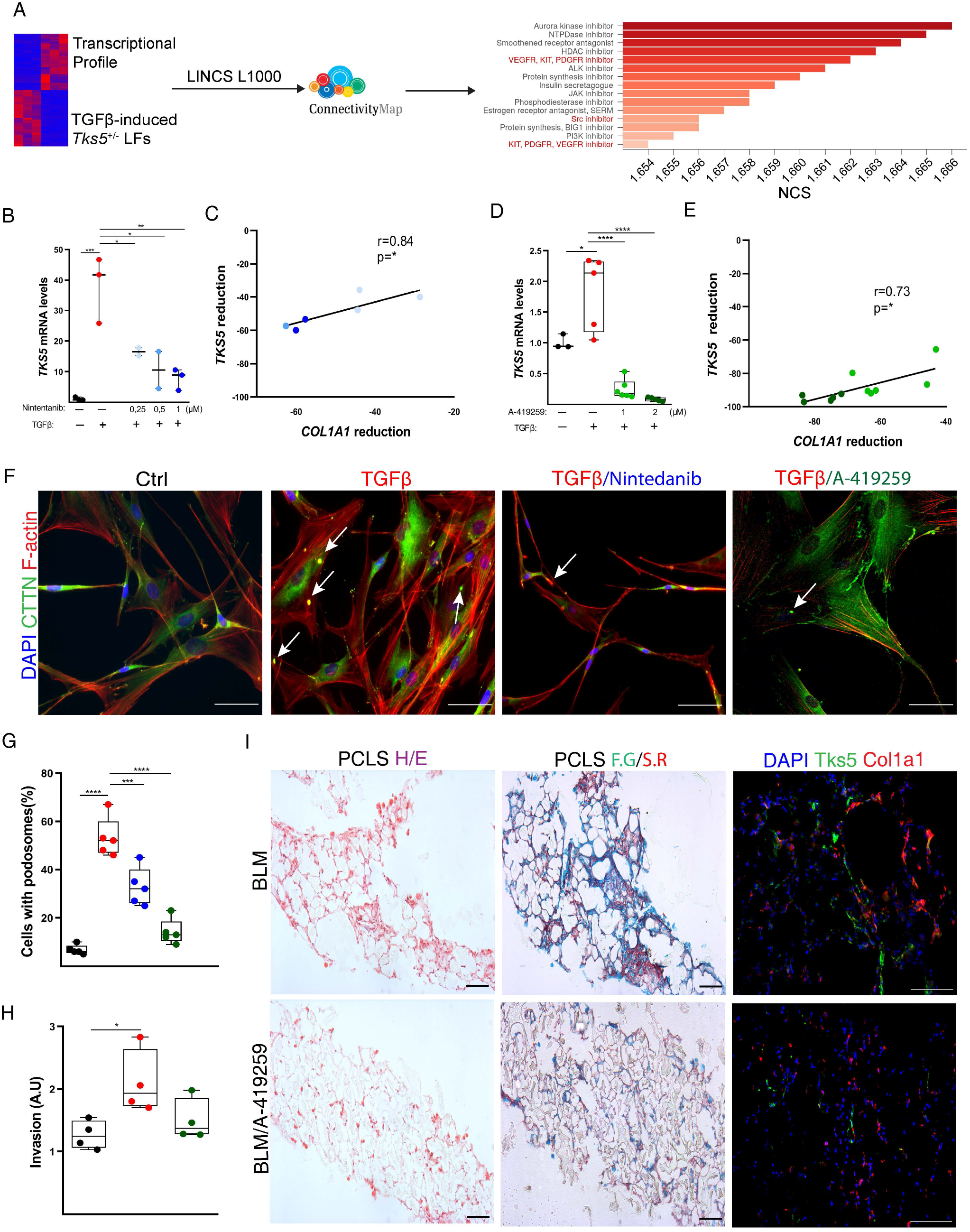
Src-inhibition potently reduces podosome formation, ECM invasion and attenuates pulmonary fibrosis. Serum starved, primary NHLFs were pretreated with A-419259 (SRC inhibitor) and Nintedanib, and then stimulated with recombinant human TGFβ (10 ng/ml) for 24h. Scale bars=50 μm; ^*/**/***/****^denote p<0.05/0.01/0.001/0.0001. **A**. Graphical representation of connectivity (CMap) analysis using LINCS1000 resource of the TGFβ-induced *TKS5*^*+/-*^ expression profile. **B-E**. *TKS5* and *COL1A1* mRNA expression was interrogated with Q-RT-PCR. Values were normalized to the expression values of the housekeeping gene *B2M* and presented as fold change over control; statistical significance was assessed with one-way ANOVA; ^*/**/***/****^denote p<0.05/0.01/0.001/0.0001. **C, E**. Pearson correlation plot of *COL1A1* reduction in the same samples (r=0.84, 0.73). **F**. Representative composite images from double immunostaining for F-actin and CTTN (E) counter stained with DAPI; arrows indicate representative podosomes. **G**. Quantification of the number of podosome-containing cells per optical field (x5). Statistical significance was assessed with one-way anova; ^***/****^denote p<0.001/0.0001. **H**. Invasion capacity of NHLFs, upon A-419259 pretreatment and TGF-β stimulation, as detected with the transwell invasion assay. Statistical significance was assessed with one-way anova; ^*^denote p<0.05 **I**. SRC-inhibition attenuates pulmonary fibrosis in mouse PCLS models, post BLM administration. Mouse precision cut lung slices (PCLS) were generated from both saline and bleomycin treated mice (d11). Treatment with A-419259, was administered in the first 24 hours after slicing for 3 consecutive days. Representative images from PCLS stained with H&E, with Fast green/Sirius red and from double immunostaining for Tks5 and Col1a1 are shown. Scale bars=50 μm; representative experiment out of 2 independent ones.

## Discussion

Increased *TKS5* expression was detected, for the first time in a non-malignant disease^15^, in the lung tissue of IPF patients and BLM-treated mice (Figs. 1 and S1-2). Increased *TKS5* expression has been previously reported, beyond cancer cell lines, in lung adenocarcinoma^19^, further extending the similarities of IPF and lung cancer^13^. *TKS5* mRNA expression in the lung tissue, of both humans and mice, correlated with the mRNA expression of *COL1A1*, a hallmark of deregulated expression in IPF, while *TKS5* expression in fibrotic lungs was predominantly localized in the alveolar epithelium and COL1A1-expressing LFs (Figs. 1 and S1-2).

TGFβ, the prototypic pro-fibrotic factor, was found to be a very potent inducer of *TKS5* expression and podosome formation in fibroblasts (NHLFs, NMLFs, MRC5, 3T3)(Figs. 2, S3-4), as previously reported only for THP-1 macrophages^29,30^ and primary aortic endothelial cells^31,32^. Other well established pro-fibrotic growth factors in the lung have been reported to modulate podosome formation in different cell types: PDGF in synovial fibroblasts^33^ and smooth muscle cells^34^, and VEGF in endothelial cells^35^, suggesting that they could exert similar stimulatory effects on LFs. Moreover, PGE_2_, which suppresses pulmonary fibrosis^36^, has been reported to promote the dissolution of podosomes in dendritic cells^37^, suggesting that the diminished PGE_2_ levels in IPF^36^ also favor the formation of podosomes. Remarkably, the formation of podosomes in LFs was shown to be an inherent property of IPF and post BLM LFs that can be maintained in culture in the absence of any stimulation (Figs. 2I-J, S6L-N). Therefore, podosome formation is an unappreciated central response of LFs to pro-fibrotic factors.

*Col1a1* mRNA levels were found to consistently correlate with *Tks5* mRNA levels in both humans and mouse lung tissue or LFs (Figs. 1C,H, S1B,D, 2B, S3B,D, S9B,H,M, 3F, 5E). *Tks5*^+/-^ LFs were shown to produce less Col1a1 post BLM (Fig. 5C), as also reflected in the reduced overall collagen deposition in the lungs of *Tks5*^+/-^ mice (Fig. 3G). Culture of LFs on Col1a1-rich aECM promoted *Tks5* expression and podosome formation (Figs. 5D-F), as well as further *Col1a1* expression, emphasizing a TGFβ-induced Col1a1-podosomes interdependency in the context of the suggested crosstalk of ECM with podosomes^38^. Accordingly, Col1 has been shown to stimulate Tks5-dependent growth^38^, while the degree of collagen fibrilization has also been reported to have a decisive effect on podosome formation^39^, likely though the Discoidin domain receptors (DDRs) that mediate collagen binding^40^. The expression of both DDR1 and 2 were found downregulated in TGFβ-induced *Tks5*^+/-^ LFs (Table S4), suggesting that DDR signaling, and consequent tyrosine kinase activation could mediate the observed TGFβ-Col1a-induced podosome formation.

Another potent podosome inducer is tropoelastin^38^, the soluble precursor of the cross-linked ECM protein elastin (Eln) whose expression was found accordingly downregulated in podosome deficient *Tks5*^+/-^ LFs (Table S4). Moreover, and beyond individual fibrosis-modulating factors, the stiff fibrotic post-BLM aECM was shown to stimulate *Tks5* expression and podosome formation in LFs and to perpetuate the increased expression of Col1a1 (Fig. 5). In agreement, increased substrate rigidity, modeled with gelatin or polyacrylamide, has been previously shown to promote invadopodia activity^41^. Therefore, the formation of podosomes in LFs upon mechanical cues from the stiff ECM of fibrotic lungs is a major component of the suggested crosstalk of ECM with fibroblasts^42,43^, especially considering the age-related increase of ECM stiffness in the lungs^7^, and the suggested role of mechanosensitive signaling in LF activation and pulmonary fibrosis^44^.

Ubiquitous genetic Tks5 haploinsufficiency was shown to attenuate BLM-induced pulmonary fibrosis with a plethora of readout assays (Fig. 3), and *Tks5*^+/-^ LFs were shown to form fewer podosomes, resulting in diminished aECM invasion (Fig. 4), thus establishing a major pathogenetic role for Tks5-enabled podosomes and LF ECM invasion in pulmonary fibrosis. ECM invasion by non-leukocytes, a hallmark of cancer^12^, is gaining increased attention in pulmonary fibrosis. IPF HLFs were shown to invade matrigel more efficiently than NHLFs or HLFs from other interstitial diseases^8,10,11^. Enhanced invasion correlated with increased actin stress fibers^10^, and was suggested to be mediated, in part, by fibronectin and integrin α4β1 signaling^11^, or hyalrounan (HA) and CD44 signaling ^8^. CD44 has been localized in invadopodia in breast cancer cells and has been shown to be required for invadopodia activity^45^, while TKS5^+^ LFs were found to preferentially express CD44 (Fig. 1F), suggesting that HA/CD44 participate in the regulation of podosome formation in LFs. TKS5^+^ LFs were also found to express preferentially CD274 (Fig. 1F), a proposed marker of invasive IPF LFs^46^, suggesting yet another potential signaling input for podosome formation. Moreover, BALFs from BLM-treated mice or IPF patients stimulated ECM invasion of LFs^47,48^, shown to be attenuated upon silencing LPAR1, EGFR and FGFR2 receptors^47^, or by interfering with Sdc4-CXCL10 interactions^48^, suggesting additional signals that could modulate LF invasion. HER2 has been also proposed to drive invasion in LFs^49^, suggesting again similarities with metastatic ADC; interestingly, several of the identified invasion-associated genes (Fstl3, Il11, Hbegf, Ccn2, Inhba, Podxl, Sema7a, Bcl2a1b, Bcl2a1d, Sh3rf1) were found down regulated in the *Tks5*^+/-^ invasion-defective LFs (Table S4; Fig. S9D), further supporting the functional results on the role of TKS5 and podosomes in LF ECM invasion.

Connectivity MAP (CMap) analysis has emerged as an invaluable tool to connect gene expression, drugs and disease states^26^. CMap analysis of scRNAseq of IPF bronchial brushings suggested that src inhibition can reverse the observed pro-fibrotic transcriptional changes in IPF bronchial airway basal cells^50^. Moreover, CMap analysis of IPF transcriptional profiles and the nintedanib and pirfenidone corresponding transcriptional signatures, indicated src inhibition as the strongest connection^51^. As shown here, CMap analysis of the TGFβ-induced *Tks5*^+/-^ LFs profile identified, among established and promising others, src inhibition as a possible treatment to limit LF invasion (Fig. 6) and therefore pulmonary fibrosis. In agreement, src inhibition was shown to reduce *Tks5* levels and podosome numbers, to decrease aECM invasion and to reverse fibrosis in PCLS (Fig. 6). Src inhibition has been previously shown to attenuate BLM-induced pulmonary fibrosis^51-53^, while the Src inhibitor Saracatanib^51^ has entered clinical trials (<u>NCT04598919</u>). Therefore, targeting podosome formation, an inherent pathogenic LF property as shown here, as well as LF interactions with the ECM including invasion, are very promising therapeutic targets in pulmonary fibrosis.

## Materials and Methods

### Datasets

All analyzed, re-normalized datasets (Table S1) were sourced from Fibromine^20^; DEGs: FC>1.2, FDR corr. p<0.05.

### Patients

All studies were performed in accordance with the Helsinki Declaration principles. Lung tissue samples (Table S2) were obtained through the University of Pittsburgh Health Sciences Tissue Bank and Yale University Pathology Tissue service, a subset of previously well characterized and published samples^54^. Lung fibroblasts were isolated from the lung tissue of IPF patients and from the adjacent healthy tissue of patients undergoing open lung surgery for cancer (Table S3) at the Department of Pulmonology, Bichat-Claude Bernard Hospital (#0811760-25032010), Paris/France.

### Mice

Mice were bred at the animal facilities of Biomedical Sciences Research Center ‘Alexander Fleming’, under SPF conditions, at 20–22°C,55 ±5% humidity, and a 12-h light-dark cycle; food and water were provided ad libitum. Pulmonary fibrosis was induced by a single oropharyngeal administration (OA) of 0.8U/kg bleomycin hydrogen chloride (BLM) to 8-10-week-old C57Bl6/J mice, as previously described^23,36^ and as analyzed in detail in the online supplement that includes all related readout assays. All experimentation was approved by the Institutional Animal Ethical Committee (IAEC) of Biomedical Sciences Research Center “Alexander Fleming”, as well as by the Veterinary Service of the governmental prefecture of Attica, Greece (# 8441/2017). The creation of a series of obligatory and conditional knock out mice for *Tks5* (*Sh3pxd2a*), as well as the corresponding genotyping strategy, is described in detail in the online supplement (and summarized in Fig. S9).

The preparation of **Precision cut lung slices (PCLS)** and **acellular ECM (aECM)**, the **i*n vitro/ex vivo* LF culture**, as well as **RT-PCR and RNAseq** were performed as described in detail in the online supplement.

### Bioinformatics and statistics

Differential expression analysis, single cell RNA-seq data re-analysis, Gene Set Enrichment Analysis, CMap/LINCS analysis are described in the online supplement. Statistical significance was assessed with the Prism (GraphPad) software, as detailed at each figure legend.

## Supporting information

All supplementary figures

All supplementary Tables

IPF podosomes

## Acknowledgements

We would like to thank Vassia Papadaki (Kafasla lab, BSRC Fleming) for support on confocal imaging, and Dimitris Kletsas (NCSR Demokritos) for NHLF clones and advice on culture optimal conditions.

