## Supplementary material for "SRC-mediated and TKS5-enabled podosome formation is an inherent property of IPF fibroblasts, promoting ECM invasion and pulmonary fibrosis": All supplementary figures

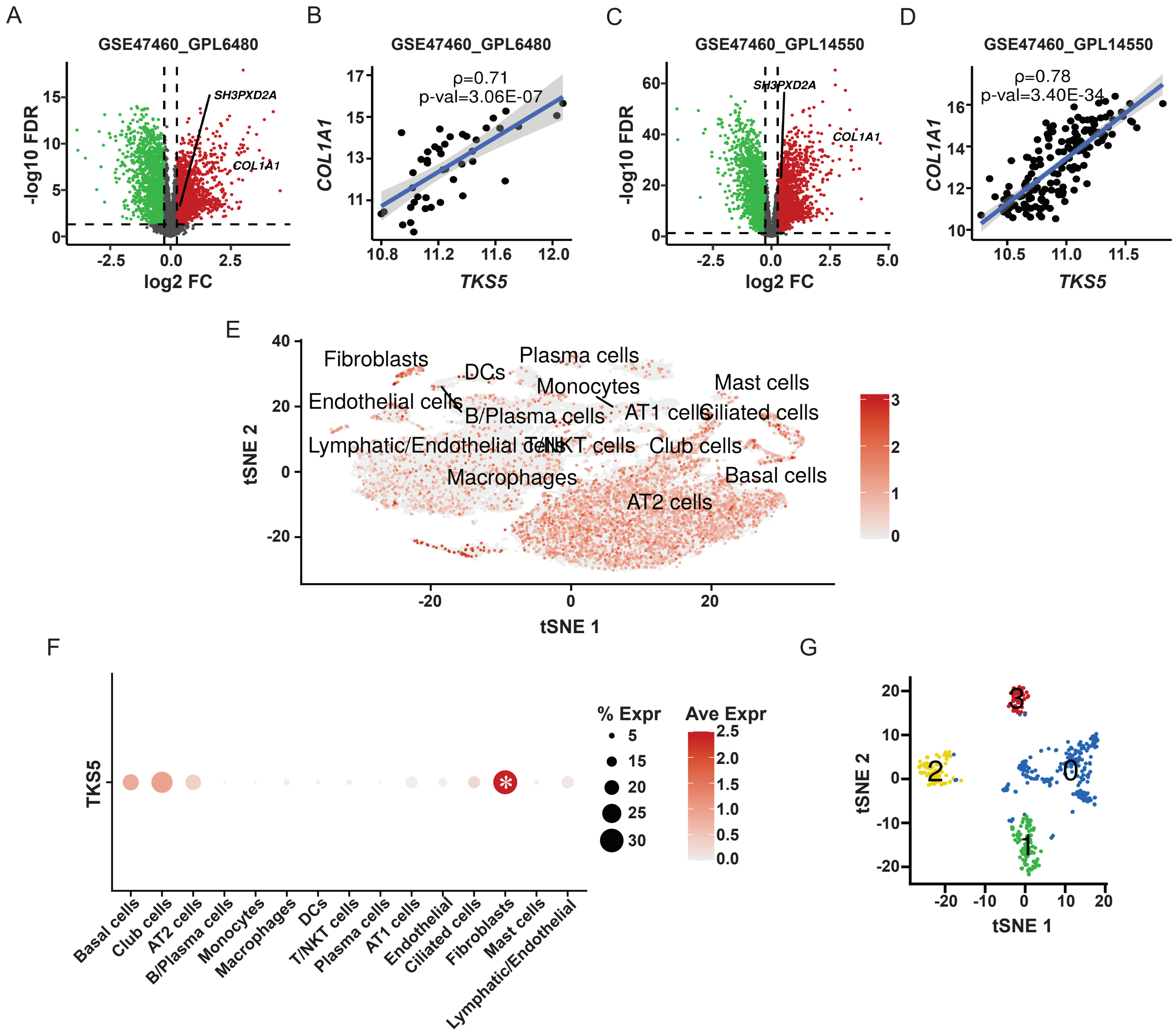


**Figure S1. Increased *TKS5* mRNA expression in the lungs of IPF patients. A, C.** Differential expression profiling of IPF lung tissue vs controls: two of the largest publicly available datasets (Table S1) are shown (FC>1.2, FDR<0.05). **B, D.** Spearman correlation plot of *TKS5* with *COL1A1* expression in the indicated datasets **(A-C)** (A; rho>0.6, p< 0.05). **E.** Visual representation of *TKS5* abundance in the detected lung tissue cell clusters from reanalysis of the scRNA-seq dataset of (Reyfman, Walter et al. 2019). **F.** In the same dataset, *TKS5* is expressed primarily by fibroblasts as compared to other cells; statistical significance was assessed with the Wilcoxon rank sum test (FC>1.2, Bonferroni adj-p<0.05). **G.** Fibroblast sub-clusters defined from the same data using a resolution of 0.1 on isolated re-processed fibroblasts.

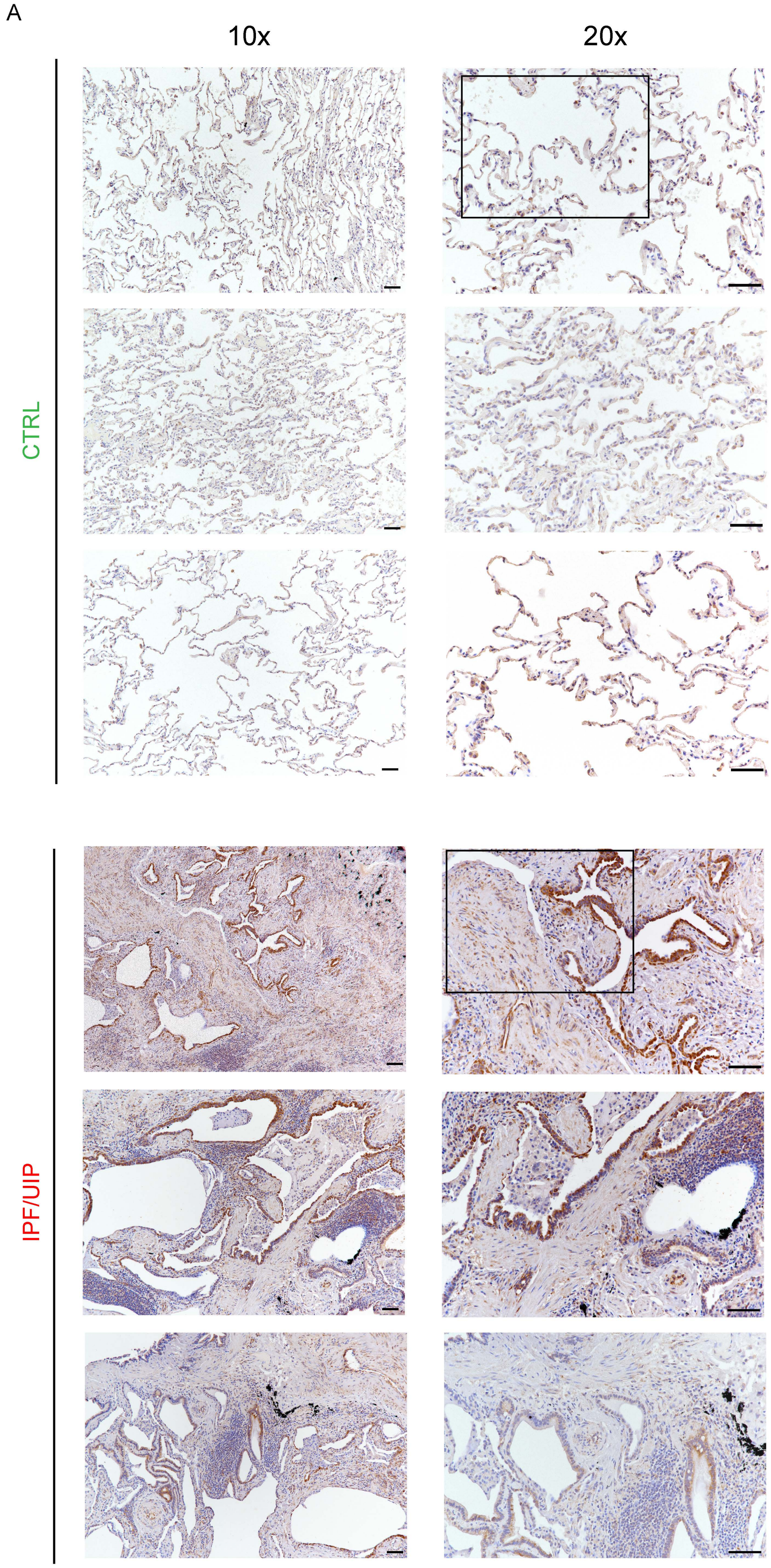


**Figure S2. A. Increased TKS5 immunostaining in the lungs of IPF patients**. Images from immunohistochemistry for TKS5 in fibrotic (IPF/UIP) and healthy (CTRL) lung tissue (n=3). Scale bars=100 μm; the indicated regions are shown in Figure 1E.

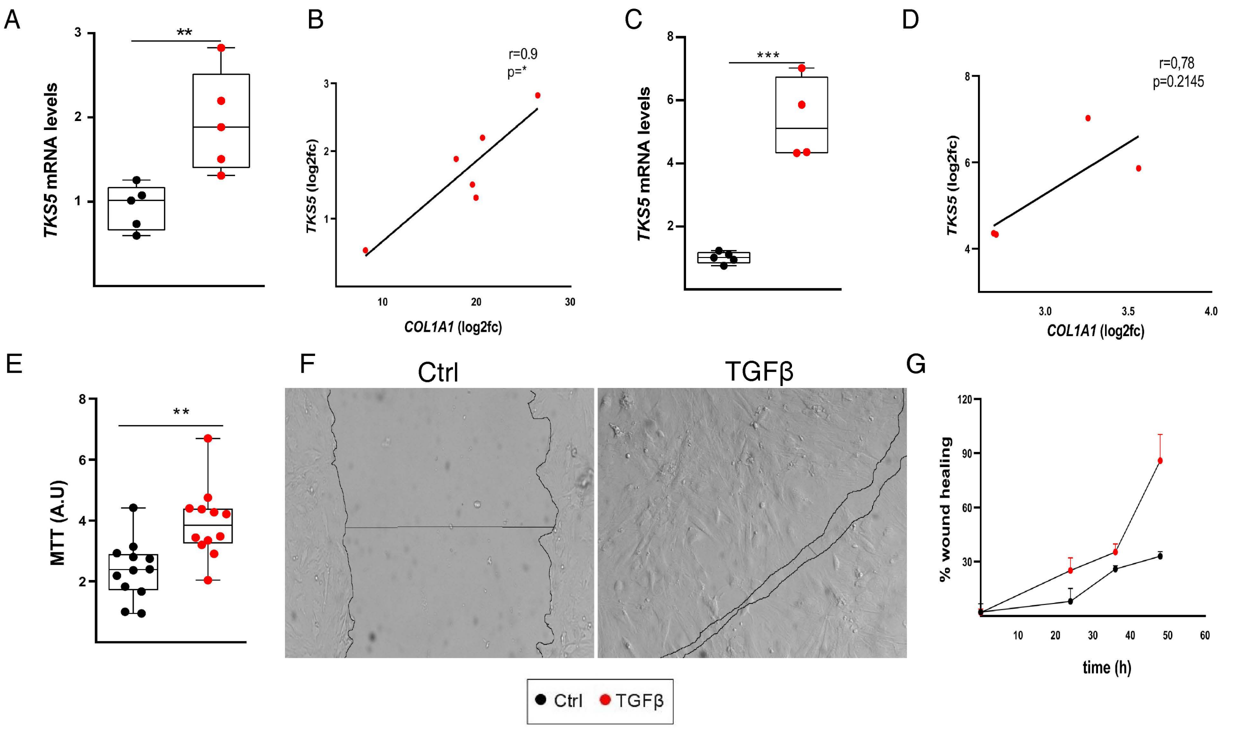


**Figure S3. TGF stimulates *TKS5* mRNA expression in lung fibroblasts (LFs), correlating with *COL1A1* expression.** Serum starved primary normal human lung fibroblasts (NHLFs; A-B) and MRC5 (human lung fibroblasts; C-D) were stimulated with recombinant human TGFβ (10 ng/ml for 24 h); representative experiment out of 2 independent ones. **A, C** *TKS5* and *COL1A1* mRNA expression was interrogated with Q-RT-PCR ($r^2$=0,93/0,95; E=102%/97%); values were normalized over the expression of the housekeeping gene *B2M* and presented as fold change over control; statistical significance was assessed with unpaired t-test; **B, D** Pearson correlation plot of *COL1A1* expression in the same samples (r=0.90,0.78 and R2=0.62); $^{*/**/***}$denote p<0.05/0.01/0.001 respectively. **E.** TGFβ-induced NHLF proliferation was assessed with the MTT assay; statistical significance was assessed with unpaired t-test; $^{**}$denotes p<0.01. **F-G.** TGFβ-induced wound healing of NHLFs, as evaluated with the scratch assay. Representative images 48h upon TGFβ stimulation are shown; **G.** Quantification of "wound closure" over time as quantified with Image J.

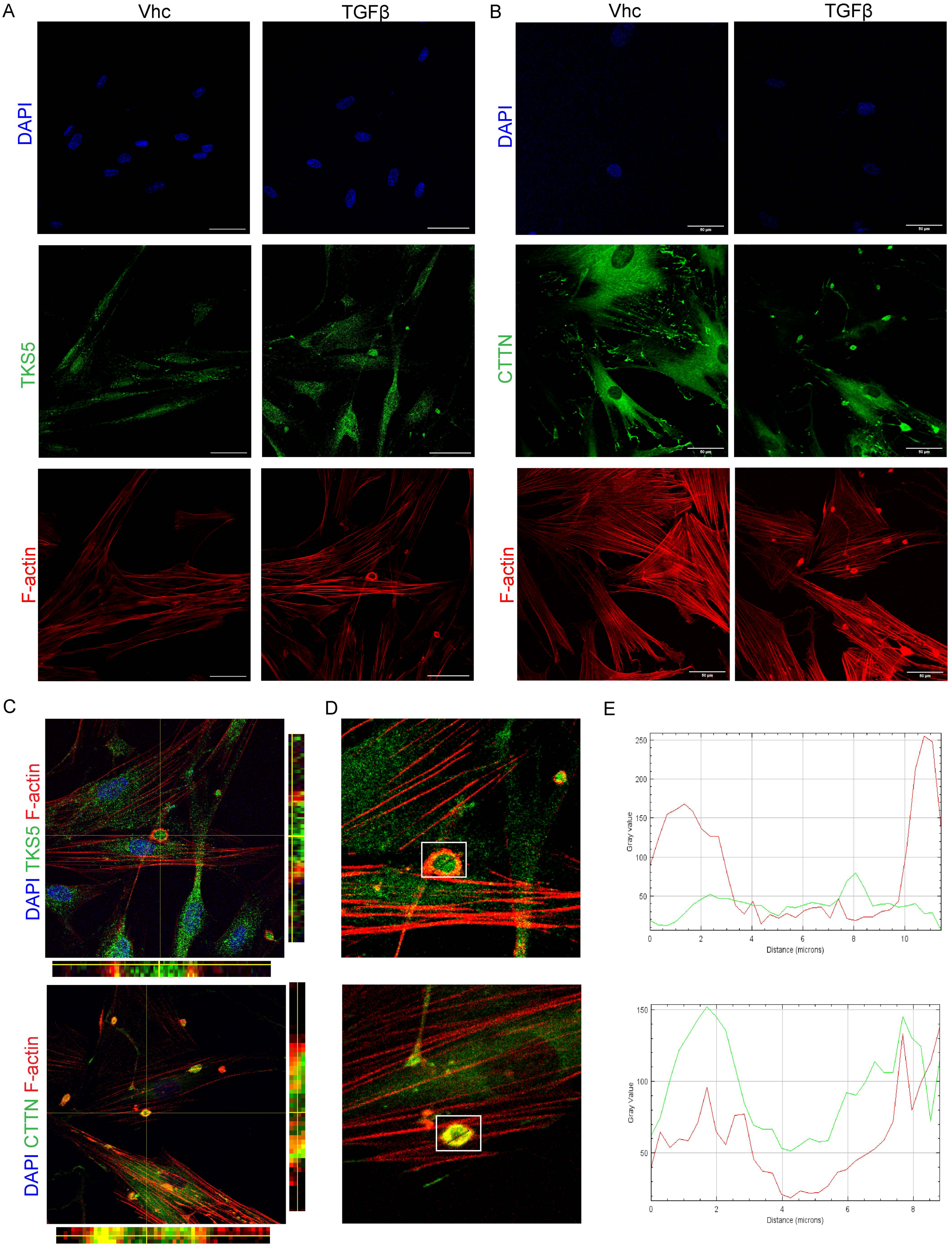


**Figure S4. TGF stimulates the formation of podosome rosettes in normal human lung fibroblasts (NHLFs).** Serum starved, sub-confluent (70-80%), primary NHLFs were stimulated with recombinant human TGFβ (10 ng/ml) for 24h. Scale bars=50 μm; representative experiment out of 3 independent ones. **A-B.** The individual images of the corresponding composite images in Figure 2 (C, E) are shown. **C.** Orthogonal projections of indicated podosomes through z-stacking. **D-E.** K-curves, showing the spatial intensity of fluorescent signals.

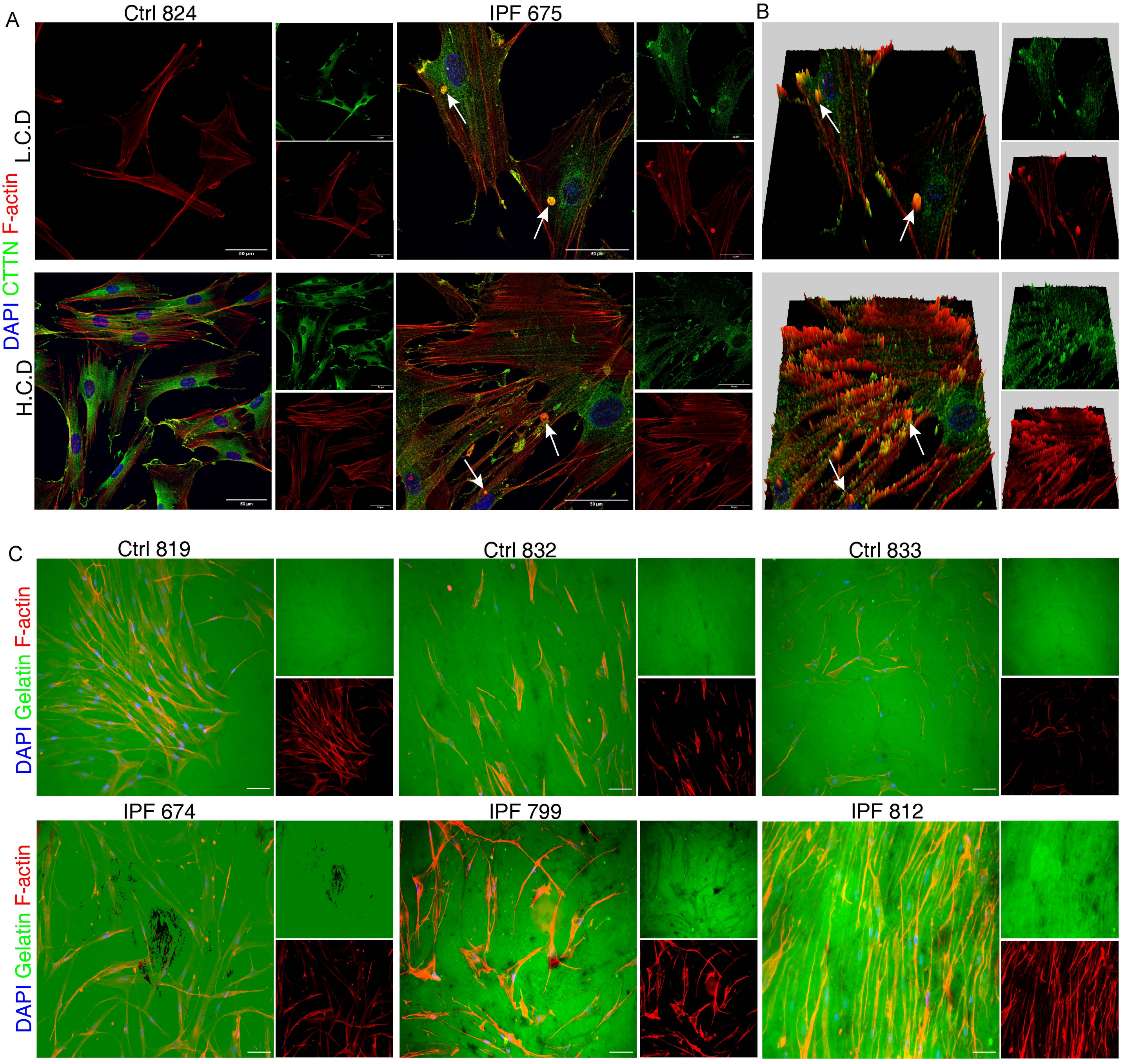


**Figure S5. The formation of ECM-degrading podosome rosettes is an imprinted property of IPF LFs. A.** Serum starved primary IPF-HLFs and NHLFs were cultured at low (LCD) or high cell density (HCD) and were immunostained for F-actin and cortactin (CTTN) and counter stained with DAPI. Representative images are shown. **B.** 3D surface plots of IPF-HLFs as captured with Image J, presenting the 3D podosome structure. **C.** The indicated clones were cultured on a fluorescein-conjugated gelatin substrate and were stained for F-actin and counter stained with DAPI; representative images are shown; scale bars=50 μm.

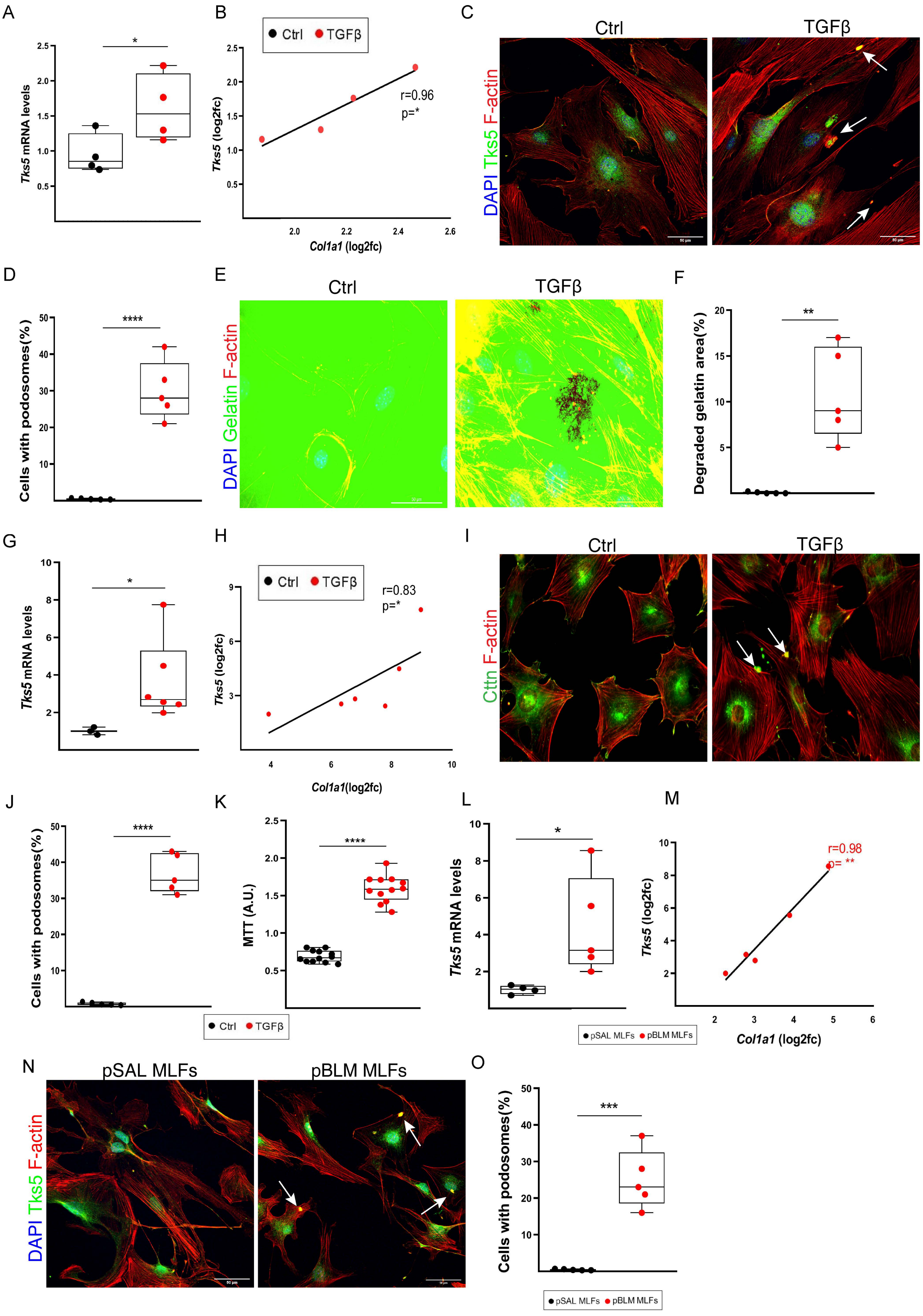


**Figure S6. TGFβ-induced, podosome rosettes is an inherent property of post BLM LFs.**

**Figure S6. TGFβ-induced, podosome rosettes is an inherent property of post BLM LFs.** Serum starved primary NMLFs or 3T3 cells were stimulated with recombinant TGF-β1 (10 ng/ml for 24 h); */**/***/**** denote p<0.05/0.01/0.001/0.0001 respectively. **A-B**, **G-H.** *Tks5* and *Col1a1* mRNA expression were interrogated with Q-RT-PCR in NMLFs and 3T3 cells respectively. Values were normalized over the expression of the housekeeping gene *B2m* and presented as fold change over control; statistical significance was assessed with unpaired t-test. **B, H**. Pearson correlation plot of *Col1a1* expression in the same samples (r=0.96,0.83). **C**, **I**. Representative composite images from double immunostaining for F-actin and Tks5 (C.) or Cttn (I.), counter stained with DAPI in NMLFs and 3T3 cells respectively; arrows indicate representative podosomes. **D, J**. Quantification of the number of podosome-containing cells per optical field (x6); statistical significance was assessed with unpaired t-test, followed by Welch's correction. **E.** Representative composite images of the TGFβ-induced degradation (black holes) of a fluorescein-conjugated gelatin substrate by NMLFs. **F.** Quantification of gelatin degradation, as quantified with ImageJ; statistical significance was assessed with unpaired t-test, followed by Welch's correction; **K.** TGFβ-induced 3T3 cells proliferation was assessed with the MTT assay. **L-M.** *Tks5* and *Col1a1* mRNA expression in NMLFs isolated post BLM administration, were detected with Q-RT-PCR, performed as in A,G; **M**. Pearson correlation plot of *Col1a1* expression in the same samples (r=0.98). **N**. Representative composite images from double immunostaining for F-actin and Tks5 counter stained with DAPI. **O.** Quantification of the number of podosome-containing cells per optical field (x5); statistical significance was assessed as in E. Scale bars 50μm; representative experiment out of 2 independent ones.

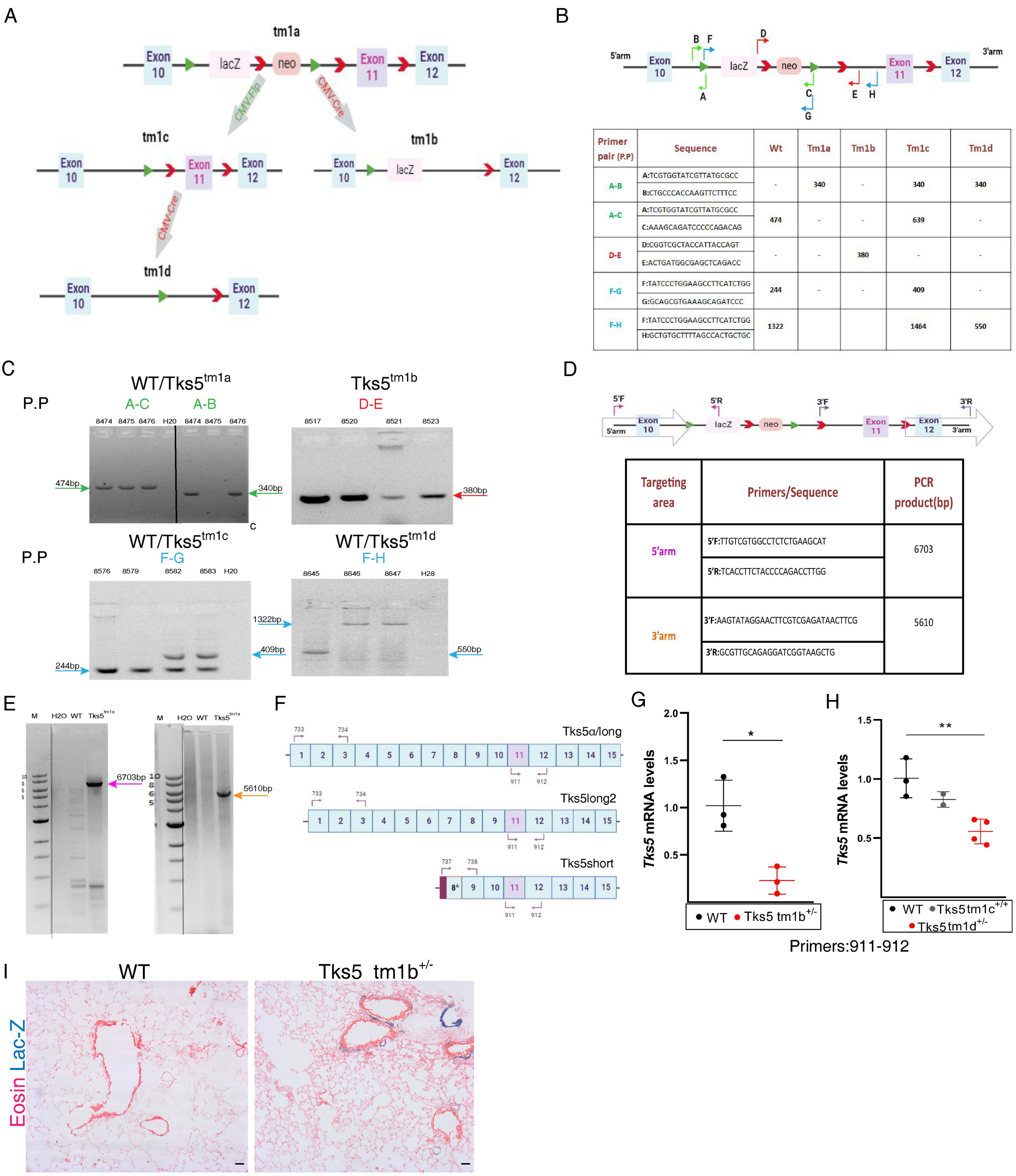


| Primer pair (P.P) | Sequence | Wt | Tm1a | Tm1b | Tm1c | Tm1d |
|---|---|---|---|---|---|---|
| A-B | A:TCGTGGTATCGTTATGCGCC<br>B:CTGCCCACCAAGTTCTTTCC | - | 340 | - | 340 | 340 |
| A-C | A:TCGTGGTATCGTTATGCGCC<br>C:AAAGCAGATCCCCCAGACAG | 474 | - | - | 639 | - |
| D-E | D:CGGTCGCTACCATTACCAGT<br>E:ACTGATGGCGAGCTCAGACC | - | - | 380 | - | - |
| F-G | F:TATCCCTGGAAGCCTTCATCTGG<br>G:GCAGCGTGAAAGCAGATCCC | 244 | - | - | 409 | - |
| F-H | F:TATCCCTGGAAGCCTTCATCTGG<br>H:GCTGTGCTTTTAGCCACTGCTGC | 1322 | | | 1464 | 550 |

| Targeting area | Primers/Sequence | PCR product(bp) |
|---|---|---|
| 5'arm | 5'F:TTGTCGTGGCCTCTCTGAAGCAT<br>5'R:TCACCTTCTACCCCAGACCTTGG | 6703 |
| 3'arm | 3'F:AAGTATAGGAACTTCGTCGAGATAACTTCG<br>3'R:GCGTTGCAGAGGATCGGTAAGCTG | 5610 |

**Figure S7. Generation of mouse conditional or obligatory knock out alleles for *Tks5*. A.** Schematic presentation of knock out alleles and genetic strategy. **B.** Genotyping strategy, as well as primer sequences and the expected product length of genotyping are indicated. **C.** Confirmation of recombination for all different alleles with genomic PCR. The primer pairs (P.P) used from the Table (b) are also indicated **D.** Genotyping strategy, as well as primers sequences and the expected product length of long-range PCR are indicated. **E.** Confirmation of successful targeting with long range genomic PCR. **F.** Schematic presentation of Tks5 mouse isoforms and the location of the real-time PCR primers, which are used to identify the deletion and the different isoforms. **G-H.** *Tks5* mRNA levels in lungs of tm1b and tm1d strains were detected with Q-RT-PCR. PCR performed with primers 911-912, detecting the deletion of critical exon 11 from all isoforms. Values were normalized over the expression of the housekeeping gene *B2m* and presented as fold change over control; statistical significance was assessed with unpaired t-test (G) or one-way ANOVA (H). **I.** Lac-Z staining in the lungs from Tks5$^{tm1b}$ mice and control littermates, indicating Tks5 lifelong transcriptional activation.

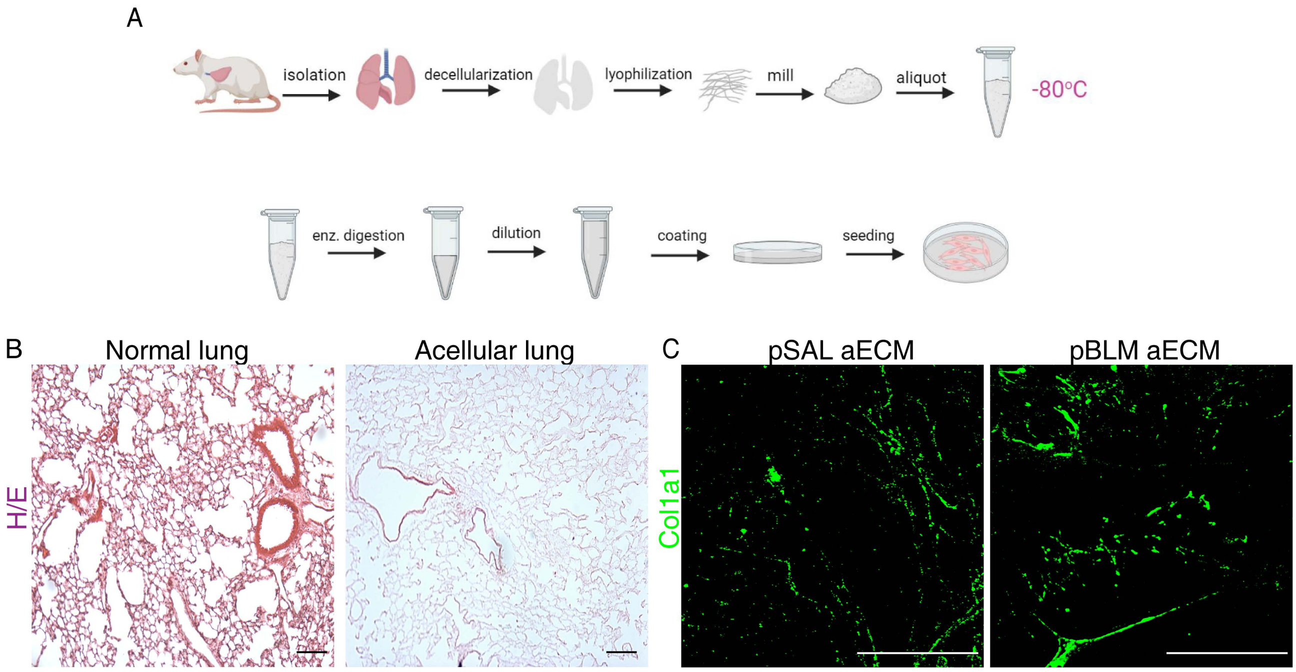


**Figure S8. Acellular extracellular matrix (aECM) as a cell substrate for aututologus *in vitro* culture of lung fibroblasts. A.** Schematic presentation of generation of aECM from mouse lungs. **B**. Confirmation of decellularization (absence of cells) with H&E staining. **C.** Representative images of immunofluorescent staining of aECM, generated from lungs upon BLM administration, for Col1a1. As expected, aECM generated from mouse lungs post BLM, is enriched in Col1a1 compared with the pSAL aECM. Representative experiment out of 2 independent ones; scale bars 50μm.

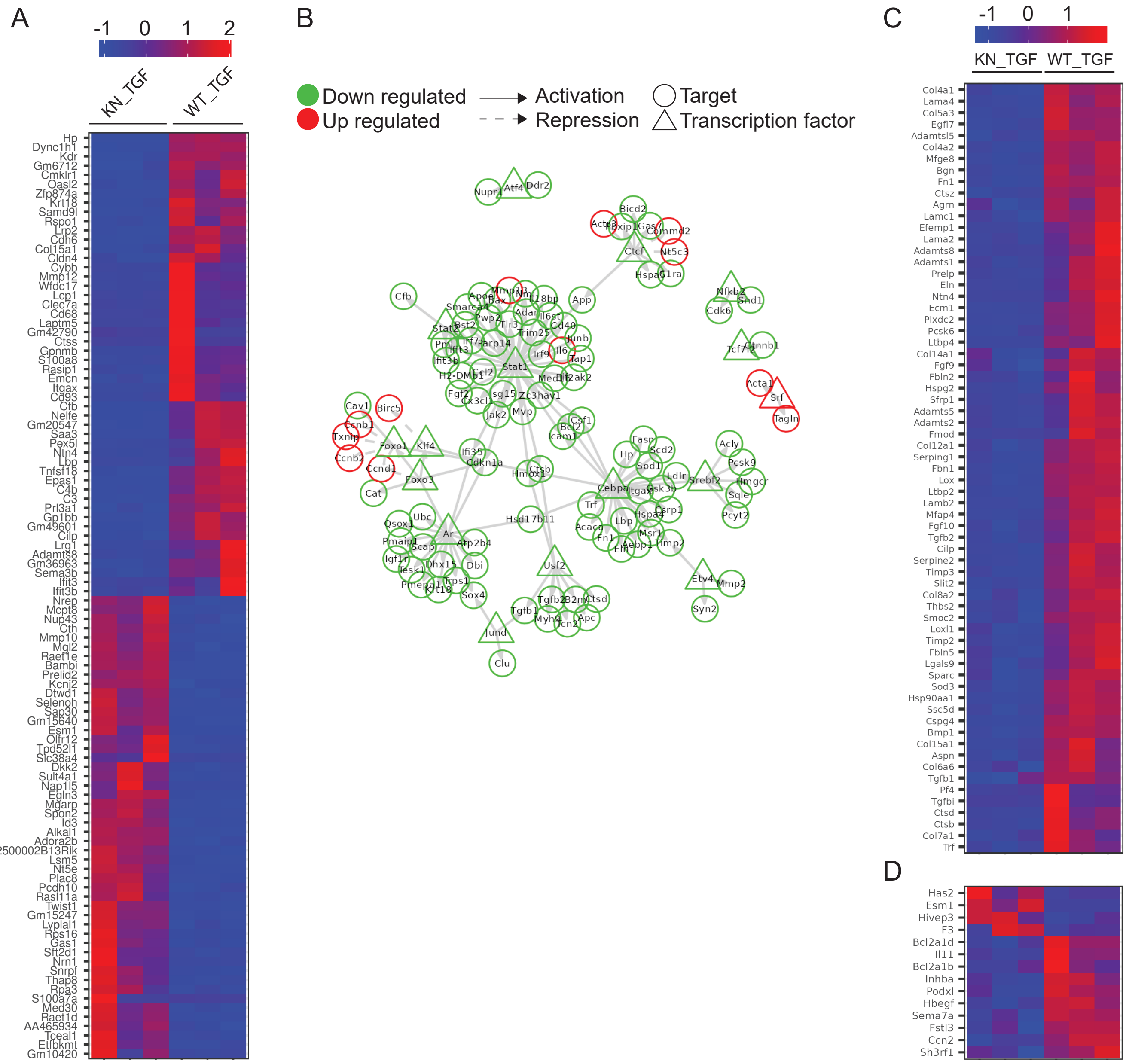


**Figure S9. Differential expression analysis of haploinsufficient *Tks5*$^{+/-}$ TGFβ-induced lung fibroblasts (LFs).** **A.** Heatmap of top 100 statistically significant (|FC| ≥ 1.2 Benjamini-Hochberg corrected PANDORA $p < 0.05$) differentially expressed genes (DEGs) in *Tks5*$^{+/-}$ LFs. **B.** Interaction network of differentially expressed transcription factors – targets in *Tks5*$^{+/-}$ LFs. **C.** Heatmap of the features in the core enrichment set of the *GO:0062023* gene ontology term as revealed during pathway analysis. **D.** Heatmap of the deregulated mouse homologous genes of the top 50 up-regulated features in invasive fibroblasts as given in the supplementary data of PMID: 30763282.
