## Supplementary material for "SRC-mediated and TKS5-enabled podosome formation is an inherent property of IPF fibroblasts, promoting ECM invasion and pulmonary fibrosis": Online supplement

*[1]Institute for Fundamental Biomedical Research, Biomedical Sciences Research Center Alexander Fleming, Athens, Greece. [2]Department of Respiratory Medicine, School of Medicine, University of Crete, Heraklion, Greece. [3]Department of Pathology, and [4]Department of Internal Medicine, Yale School of Medicine, New Haven CT, USA; [5]Department of Pulmonology, Bichat-Claude Bernard Hospital, Paris, France. [6]Department of Respiratory Medicine, School of Medicine, University of Patras, Patras, Greece.*

### Online supplement

**Creation of a series of obligatory and conditional knock out mice for Tks5**

To genetically dissect the likely role of Tks5 in pulmonary fibrosis and pathophysiology in mice, we then created a series of obligatory and conditional knock out mice for *Tks5* (*Sh3pxd2a*). The *Sh3pxd2a* locus has been already targeted by the European Conditional Mouse Mutagenesis Program (EUCOMM), aiming to knock out all mouse genes in a high throughput approach [1]. In this context, the exon 11 of the *Sh3pxd2a* gene was loxP-flanked, while a LacZ/neomycin reporter/selection cassette was placed upstream, including two FRT sites; this allele is referred to as "targeted mutation 1a" (tm1a; Fig. S7A)[1]. The targeted ES cells were then microinjected into C57Bl/6N blastocysts by the Welcome Trust Sanger Institute (WTSI), that were transferred in pseudopregnant females to yield the *Sh3pxd2a*$^{tm1a(EUCOMM)Wtsi/+}$ heterozygous mice (Fig. S7A). Frozen sperm of these mice was obtained from WTSI, via the INFAFRONTIER consortium [2,3] and the European Mutant Mouse Archive (EMMA), that was directly injected to mice in the transgenic facility of "BSRC Fleming" via IVF technology to yield the *Sh3pxd2a*$^{tm1a(EUCOMM)WtsiFlmg/+}$ heterozygous mice. Mice were genotyped following the corresponding strategy from EUCOMM, that queries three different genomic fragments (lacz, WT allele, tm1a allele) by performing three independent PCR reactions (Fig S7B, C). Moreover, the successful targeting was also verified with long range PCR for both the 5' and 3' arms flanking the floxed region with primers against inserted sequences (Fig. S7D, E).

To obtain the tm1b reporter allele (Fig. S7A) *Sh3pxd2a*$^{tm1a/Fleming/+}$ mice were mated with transgenic mice expressing the Cre recombinase under the control of the Cytomegalovirus (CMV) promoter in all mouse tissues and cells (*Tg-CMV-Cre*) [4]. Genetic recombination of the obtained *Sh3pxd2a*$^{tm1b\ (EUCOMM)WtsiFlmg/+}$ mice was verified with genomic PCR (Fig. S7B, C). Q-RT-PCR in lung tissues indicated a 50% reduction of *Sh3pxd2a* mRNA levels indicating proper gene targeting (Fig. S7G). X-gal staining, detecting LacZ expression from the promoter of Tks5 (Fig. S7A, Tm1b), localized transcriptional *Tks5* activation (throughout development, neonatal and adult life) mainly in arterial endothelium of the lung (Fig. S7I). No obvious gross macroscopic abnormalities were observed, while heterozygous mice were healthy and fertile. Intercrossing of heterozygous mice *Sh3pxd2a*$^{tm1b(EUCOMM)WtsiFlmg\ /+}$ (*Tks5*$^{+/-}$) mice yielded no homozygous knockout mice, indicating that *Sh3pxd2a* has an essential role in mouse development, as previously reported for an obligatory knock out strain [5].

A similar genetic strain, *Sh3pxd2a*$^{tm1b/(EUCOMM)Wtsi/+}$ was created by WTSI from the *Sh3pxd2a*$^{tm1a(EUCOMM)Wtsi/+}$ mice via a cell permeable HTN-Cre [6]. *Sh3pxd2a*$^{tm1b/(EUCOMM)Wtsi/+}$ heterozygous mice were systematically phenotyped from the INFAFRONTIER consortium [2,3] on our behalf, following a relative competitive call. Results indicated that Sh3*pxd2a*$^{tm1b/(EUCOMM)Wtsi/+}$ mice present with no major pathophysiological abnormalities, apart from an increase of serum alkaline phosphatase in females (measurements chart). Moreover, a viability primary screen phenotypic assay was performed on the novel mutant strain by WTSI (data chart), confirming the requirement of *Tks5* for embryonic development.

To generate the conditional tm1c allele (Fig. S7A), *Sh3pxd2a*$^{tm1a(EUCOMM)WtsiFlmg\,/+}$ mice were crossed with transgenic mice expressing the Flp recombinase under the control of the Cytomegalovirus (CMV) promoter in all mouse tissues and cells (*Tg-CMV-Flp*)[7]. Genetic recombination of the obtained *Sh3pxd2a*$^{tm1c(EUCOMM)WtsiFlmg/+}$ mice was verified with genomic PCR (Fig. S7C).

To generate the conditional tm1d allele (Fig. S7A), *Sh3pxd2a*$^{tm1c/(EUCOMM)WtsiFlmg/+}$ mice were crossed with transgenic mice expressing the Cre recombinase under the control of the Cytomegalovirus (CMV) promoter in all mouse tissues and cells (*Tg-CMV-Cre*)[4]. Genetic recombination of the obtained *Sh3pxd2a*$^{tm1d(EUCOMM)WtsiFlmg\,/+}$ mice was verified with genomic PCR (Fig. S7C). Q-RT-PCR in lung tissue indicated a 50% reduction of *Sh3pxd2a* mRNA levels indicating proper gene targeting (Fig. S7H).

#### Bleomycin (BLM)-induced pulmonary fibrosis

Pulmonary fibrosis was induced by a single oropharyngeal administration (OA) of 0.8U/kg bleomycin hydrogen chloride (BLM) (Nippon Kayaku Co., Ltd., Tokyo, Japan) at day 0 into anesthetized (i.p.; xylazine, ketamine, and atropine, 10, 100, and 0.05 mg/kg, respectively) 8-10-week-old mice; control groups received sodium chloride (SAL). Dose and route were selected upon prior extensive local testing to induce a solid fibrotic profile, while minimizing lethality. All randomly assigned experimental groups consisted of littermate mice. The health status of the mice was monitored at least once per day; no unexpected deaths were observed; all measures were taken to minimize animal suffering and distress. Disease development was assessed in comparison with WT littermates 14 days post-BLM, at the peak of the disease (which resolves at d21 post BLM in these settings).

Following weighing, **respiratory functions** were measured with **FlexiVent** (SCIREQ, Montreal, Canada), according to manufacturer instructions and as previously described [8,9]. Euthanasia was humanly performed in a $CO_2$ chamber with gradual filling followed by exsanguination.

**Bronchoalveolar Lavage fluid (BALF**) was obtained by lavaging the lungs with 1ml of 0.9% sterile sodium chloride three times. After the isolation, the samples were centrifuged at 1200g for 10 min at 4°C, the first BALF supernatant was stored at -80°C for protein and collagen measurements. To estimate **pulmonary inflammation**, BALF cell pellets were redissolved in 1ml saline, stained with 0.4% Trypan Blue solution and were counted with the use of a Neubauer hematocytometer. Total protein levels in BALFs, an indication of **pulmonary edema and vascular leak**, were assessed with the Bradford assay according to the manufacturer's instructions (Bio-Rad, Hercules, CA, USA). In a 96-well plate 5μl of every BALF sample is placed, followed by the addition of 245 μl of 1x Bradford reagent and incubation for 5 minutes in the dark. Absorbance values were then measured at 595nm, using a spectrometer, and were converted in concentration values (mg/ml) using a bovine serum albumin standard curve (BSA 0–2 mg/mL). Total **soluble collagen** in BALFs was quantified using the Sirius Red assay. 50μl of BALF samples, diluted in 350μl of 0.5M acetic acid, were incubated for 30min with 400μl of Direct at RT, in the dark. This was followed by centrifugation, at 12000g for 10 min and isolation of 200μl of the supernatants. Absorbance values were measured at 540nm, using a spectrometer, and were converted in concentration values (μg/ml) using a rat tail collagen I standard curve (0–500 μg/mL).

**Histology**

The right lung was fixed overnight in 10% neutral buffered formalin and embedded in paraffin. 5μm lung sections were cut using a Microtome and stained with Hematoxylin/eosin (**H&E**) with standard protocols. Fibrosis development was quantified by two independent reviewers, in a blinded manner, based on a modified **Ashcroft score** (0, normal lung; 1, isolated alveolar septa with gentle fibrotic changes; 2, fibrotic changes of alveolar septa with knot-like formation; 3, contiguous fibrotic walls of alveolar septa; 4, single fibrotic masses; 5, confluent fibrotic masses; 6, large contiguous fibrotic masses; 7, air bubbles; 8, fibrous obliteration). For **Fast green-Sirius red (F.G/S.R)** collagen staining, lung sections were deparaffinized in xylene and ethanol and incubated in Bouin's solution (75% picric acid/ 25% formaldehyde/ 1% acetic acid), for 1 hour at $56^0$C, followed by staining with Fast Green 0,04% in picric acid for 15 minutes and Sirius Red 0,1%/Fast Green 0,04% dissolved in picric acid for 40 minutes. Stained sections were washed in acetic acid, then dehydrated and mounted with DPX. PCLS, isolated and cut as described below stained with H&E. For **X-gal (Lac-Z)**, freshly isolated mouse lungs were inflated with 0,1 g/ml sucrose in 50% OCT/PBS, followed by the simultaneous embedding and freezing in OCT, using isopentane and dry ice. Sections of 6-10 μm were cut using a cryotome and fixed in 2% formaldehyde/ 0,2% glutaraldehyde for 15 minutes at 4°C. Next, they were washed twice in cold PBS/ 2 mM $MgCl_2$ for 10 minutes and stained overnight with X-gal staining solution (2 mg/mL X-gal in 0.1 M Sodium phosphate buffer pH=7.3, 0,01% Sodium deoxycholate, 5 mM $K_3Fe(CN)_6$, 5,7 mM $K_4Fe(CN)_6$, 2 mM $MgCl_2$, 0,02% NP-40) at 37°C in the dark. The sections were then rinsed twice with PBS/2 mM $MgCl_2$ and $dH_2O$ for 5 minutes at room temperature, counterstained with eosin following by dehydration and mounting with DPX. **Imaging** was performed using a Nikon Eclipse E800 microscope (Nikon Corp., Shinagawa-ku, Japan) attached to a Q Imaging EXI Aqua digital camera, using the Q-Capture Pro 7 software. For **immunohistochemistry** studies, lung sections were deparaffinized in xylene, rehydrated in a gradient of ethanol, and briefly washed with water. The slides were kept in tap water until ready to perform antigen retrieval with sodium citrate buffer with pH 6.0 by autoclave for 20 min. Then they were treated with blocking solution (10% normal goat serum/2% BSA) at room temperature for 1 h and incubated with primary antibodies overnight at 4°C. After washing, they were incubated with fluorophore-conjugated secondary antibodies diluted in the blocking solution. Following this, sections were washed 3 times with PBS-T and mounted with medium containing DAPI for nuclear visualization. **Imaging** was performed using a TCS SP8X White Light Laser confocal system (Leica).

***In vitro/ex vivo* lung fibroblast (LF) cell model**

Normal human lung fibroblasts (**NHLFs**) and **IPF-HLFs** were isolated from fresh tissue samples by plating several 2-3 mm pieces on 10 cm tissue culture plates in DMEM supplemented by penicillin/streptomycin solution, 10% FBS at 37° C and 5% $CO_2$ in a humidified atmosphere. 10-14 days following plating, proliferating fibroblasts surrounded the tissue pieces, which were then removed, and cells are detached with trypsin-EDTA solution and replated in F75 tissue culture flasks (P0) until confluent. Removed tissue pieces were replated in fresh 10 cm tissue culture plates for a second round of fibroblast outgrowth in the same conditions as above. Following 3 passages, homogeneous fibroblasts' colonies are observed; HLFs are used until passage 7-8. A similar procedure was independently used for an additional NHLF clone (Fig. S3A-B), as previously published[10].

Primary normal mouse lung fibroblasts (**NMLFs**) were isolated from 8-10-week-old C57Bl6/J mice and/or from BLM-challenged mice. Upon sacrifice, perfused lungs were excised in DMEM. Then lungs were minced and digested with 0.7 mg/ml collagenase type IV (C5138-Sigma), for 1h at 37°C. Digestion was followed by filtration and the suspension was centrifuged at 1200 rpm for 5 min. Finally, the pellet was resuspended in DMEM containing 10% FBS. All experiments were performed at passage 2-3.

NHLFs, IPF-HLFs, NMLFs and murine and human embryonic lung fibroblasts (**3T3** and **MRC5** cell lines respectively) were cultured in DMEM supplemented with 10% fetal bovine serum (FBS) and streptomycin/penicillin and amphotericin and incubated at 37°C and 5% $CO_2$. Cells were cultured to 60-80% confluency, were starved overnight with serum-free DMEM (+ 0.1% BSA) and were exposed to 10 ng/ml TGF-β1 for 24h in serum-free DMEM. In control samples the diluent of TGFβ (7.5% BSA in $H_2O$) was used. For pharmacologic studies, NHLFs were seeded at 6-well plates, were serum starved and pre-treated for 1h with the indicated increasing concentrations of various agents and theirs diluents in controls. After one hour of pre-treatment, cells were incubated with TGFβ as usual.

The **proliferation** of all LF cultures was quantified with the **MTT** assay in 96-well plates, where a common solution of DMEM and MTT was added into each well. After incubation for 4 h and having confirmed the formation of purple crystals, the media was removed, and acidified isopropanol was added into each well to dissolve the formazan crystalline product. Absorbance values were determined at 570 nm and background subtracted at 660 nm using an OPTImax Microplate Photometer (Molecular Devises).

The spreading and **migration** capabilities of LFs were assessed in a scratch wound assay. Cells were cultured into a 12-well plate as above and left to grow until confluency. Upon confluency, TGFβ was added, as described above. A wound was generated using a sterile pipette tip. The plate was placed in a tissue culture incubator at 37 °C, and photos were taken under a reverse microscope at specific intervals.

Migration and **invasion** were quantified also with Boyden chambers according to manufacturer's instructions, as shown schematically in Figure 4E. Briefly, LFs were added to the upper chamber which was pre-coated with aECM substrate (or not for migration) and allowed for 6 hours to invade/migrate through the transwell membrane to the lower side which was in touch with the starvation medium. Then, the cells which remained in the upper chamber were removed, while invasive or migratory cells, after washes and fixation, were stained with crystal violet. Additionally, stained cells were lysed with Lysis Buffer for 20 minutes and absorbance was measured at 550 nm using the TECAN Sunrise Microplate Photometer.

The **proteolytic capacity** of LFs was assessed with the Fluorescein gelatin degradation assay. 12-mm glass coverslips were acid washed with 20% nitric acid, incubated with 50 μg/ml poly-L-lysine, crosslinked with 0.5% glutaraldehyde and then coated for 20 min with 0.2% fluorescein-conjugated gelatin (Invitrogen, Gelatin From Pig Skin, Fluorescein Conjugate: G13187) in 2% sucrose-containing phosphate buffered saline (PBS). Then, the coverslips were treated with sodium borohydride (NaBH4), washed with PBS, and transferred to a new 24 well-plate. Fibroblasts were seeded in gelatin-coated coverslips and treated with TGF-β1 as previously described. After 24 hours, medium was replaced with full medium and cells were processed for immunofluorescence, 24-48 hours later. The quantification of degradation was performed with Image J.

For **IF stainings**, cells were seeded in coverslips (20.000 cells/well) and after starvation, incubated with TGF-β1 as usual. The cells were fixed with 4% PFA for 15 min and permeabilized with 0.1% Triton X for 10 min. This was followed by blocking with 2% BSA in PBS for 1h at RT. The cells were incubated overnight at 4°C with primary antibodies. The next day, cells were washed, and incubated with a secondary antibody and conjugated phalloidin in 1%BSA/PBS for 60min at RT. Finally, after washes, coverslips were mounted with a drop of mounting Fluoroshield medium (containing DAPI for nucleus labelling).

The decellularization and generation of **acellular ECM (aECM)** from mouse lungs was performed, based on similar protocols for other tissues[11,12]. Briefly, whole lungs were isolated and treated with increasing concentrations of SDS (0.01, 0.1, 1%) in a PBS solution, with 24 h incubation for each SDS concentration. For the final step, decellularized lung tissues were washed with PBS for at least 3 days, cut into small pieces and stored at -80ºC. Frozen tissue was lyophilized using a lyophilizer and then milled in liquid nitrogen. To produce an ECM substrate, the milled form of the matrix was solubilized through enzymatic digestion. Pepsin (Sigma-Aldrich, P6887) was dissolved in 0.1 M HCl to make a concentration of 1 mg/ml. Approximately 10 mg of the ECM powder were digested in 1 mL of pepsin solution, in order to solubilize the ECM components. After approximately 48 hours, the matrix was diluted using 0.1 M acetic acid to make a 5 mg/ml concentration of lung ECM solution, which was used as a coating substrate for cells.

**Precision cut lung slices (PCLS)**

C57-Bl/6, 8-10-week-old mice, were administered with SAL/BLM as described above. On day 11, mice were sacrificed and lungs, after perfusion, were inflated with 1ml of with 1.5% Low Melting Agarose (15517-014-Invitrogen) in saline. Lungs were then isolated and incubated at RPMI medium (Thermo Fisher Scientific) supplemented with 10% FBS and 1% penicillin-streptomycin for 30 min at $4C^0$ to allow agarose polymerization. The left lobe of the agarose filled lungs, was cut into 200μm slices (PCLS) with Vibratome. PCLS were then cultured in 700 μL of RPMI medium supplemented with 10% FBS and 1% penicillin-streptomycin in 24-well plates at standard conditions (37°C and 5% $CO_2$) overnight. Next, PCLS were incubated to 2μm of A-419259 (Src-family inhibitor) and $H_2O$ for 3 consecutive days, changing the treatment daily. In day 14, PCLS were fixed overnight with PFA at 4°C, until slices embedded in paraffin. Finally, 5μm sections of PCLS were cut using a Microtome and stained for H&E, F.G/S.R and antibodies accordingly.

**Antibodies- Reagents**

Antibodies used in this study included anti-Tks5 (SH3 domain) rabbit monoclonal antibody (Merck, 3174822, 1:100), anti-SH3PXD2A mouse monoclonal antibody (Origene, clone OTI1F5-TA811757S, 1:250), col1a1 rabbit monoclonal antibody (Thermo Scientific, RE2209815, 1:100), anti-A-actin (sma) mouse monoclonal antibody,(Origene, UM800129, 1:250), recombinant Anti-Cortactin antibody (Abcam,ab81208, 1:500), Alexa Fluor™ 633 Phalloidin (Invitrogen, A22284, 1:50), MMP-9 (D6O3H) XP Rabbit monoclonal Antibody (Cell signaling #13667, 1:100) Secondary antibodies included: Goat anti-Rabbit IgG (H+L) Cross-Adsorbed Secondary Antibody Alexa Fluor 488 (a11008), Goat anti-Rabbit IgG (H+L) Cross-Adsorbed Secondary Antibody, Alexa Fluor 555 (a21428), Goat anti-Mouse IgG (H+L) Highly Cross-Adsorbed Secondary Antibody Alexa Fluor 488 (a11029) Goat anti-Mouse IgG (H+L) Highly Cross-Adsorbed Secondary Antibody Alexa Fluor 555 (a221424). For pharmacological studies A-419259 inhibitor SML0446 (Sigma-Aldrich), Nintedanib SML2848 (Sigma-Aldrich) and Pirferidone P2116 (Sigma-Aldrich) were used.

**Real Time quantitative RNA RT-PCR (Q-RT-PCR)**

In human samples, RNA was extracted from 30 – 50 mg of frozen lung tissue in 700 μL of Qiazol (Lysis buffer, Qiagen, Valencia, CA) by tissue disruption and homogenization using an electric homogenizer (PolyTron homogenizer H3660-2A, Cardinal Health, Dublin, OH) at 19.000 rpm for 15 seconds, according to the manufacturer's instructions. RNA was purified using the miRNeasy Mini kit (217004, Qiagen, Valencia, CA) with the assistance of the Qiacube automated system (9001292, Qiagen, Valencia, CA). The purity of the RNA was verified using NanoDrop at 260 nm and the quality of the RNA was assessed using the Agilent 2100 Bioanalyzer (Agilent, Technologies, Santa Clara, CA). Real-time PCR was performed with Taqman primers as described in the table below. Values were normalized to the expression of B2M.

In mouse samples RNA was extracted from the left lung lobe using the Tri Reagent (TR-118) obtained from Invitrogen and treated with DNAse (RQ1 RNAse-free DNAse) prior to RT-PCR according to manufacturer's instructions. cDNA synthesis was performed using 2 µg of total RNA per sample in 20-µl reaction using M-MLV RT (Promega). Real-time PCR was performed on a BioRad CFX96 Touch™ Real-Time PCR Detection System (Bio-Rad Laboratories). Values were normalized to the expression of b-2 microglobulin (b2m). The annealing temperature for all primers was 58°C. Primers sequences for RT and genomic PCRs are depicted on the table below.

**Primers and sequences used for RT and genomic PCR.**

| | **Description** | **Sequence- Lot (for taqman)** |
|---|---|---|
| **Human** | m1 SH3PXD2A | Hs00206037 |
| | m1 B2M | Hs00984230 |
| | B2M F | AGATGAGTATGCCTGCCGTG |
| | B2M R | CTGCTTACATGTCTGGATCCCA |
| | TKS5 LONG F | CTCCCAAGAAGGACGTGACA |
| | TKS5 LONG R | CTCTTGGACACTTCCCCAGT |
| | COL1A1 F | CGAAGACATCCCACCAATCAC |
| | COL1A1 R | CATCGCACAACACCTTGCC |
| **Mouse** | B2m F | TTCTGGTGCTTGTCTCACTGA |
| | B2m R | CAGTATGTTCGGCTTCCCATTC |
| | Tks5 F | GGAGCCCCTCTAAACACTATGT |
| | Tks5 R | GGCCACCTTCAATAGGAAACTT |
| | Tks5 long F | TTATCAACGTGACCTGGTCTG |
| | Tks5 long R | TTCGGATCCTTCTGGCCAC |
| | Tks5short F | TGGCTCACCGCGTGCTTTCTG |
| | Tks5 short R | CCTTGCTCTTCAGATGTGCTCACAA |
| | Tks5 tm1b-d (ex11-12) F | AAGACGAGATCGGCTTCGAG |
| | Tks5 tm1b-d (ex11-12) R | TCCCTATGATCTCCACCGGA |
| | Col1a1 F | CTACTACCGGGCCGATGATG |
| | Col1a1 R | CGATCCAGTACTCTCCGCTC |
| **Mouse genomic primers** | (A) CAS_R1_Term | TCGTGGTATCGTTATGCGCC |
| | (B) Sh3pxd2a 270538 F | CTGCCCACCAAGTTCTTTCC |
| | (C) Sh3pxd2a 270538 R | AAAGCAGATCCCCCAGACAG |
| | (D) Tm1b prom F | CGGTCGCTACCATTACCAGT |
| | (E) Floxed LR | ACTGATGGCGAGCTCAGACC |
| | (F) Sh3pxd2a F | TATCCCTGGAAGCCTTCATCTGG |
| | (G) Sh3pxd2a R1 | GCAGCGTGAAAGCAGATCCC |
| | (H) Sh3pxd2a R2 | GCTGTGCTTTTAGCCACTGCTGC |
| | Long range 5' arm F | TTGTCGTGGCCTCTCTGAAGCAT |
| | Long range 5' arm R | TCACCTTCTACCCCAGACCTTGG |
| | Long range 3' arm F | AAGTATAGAACTTCGTCGAGATAACTTCG |
| | Long range 3' arm R | GCGTTGCAGAGGATCGGTAAGCTG |

**RNA sequencing**

Six total RNA samples were prepared, and their concentration was measured with nanodrop (ND1000 Spectrophotometer – PEQLAB). The samples measured to a concentration of 400-500ng/μl and therefore 1μl of RNA, from each sample was used to proceed with the library preparation. The RNA quality of each sample was measured in bioanalyzer (Agilent Technologies) using the Agilent RNA 6000 Nano Kit reagents and protocol. For the preparation of per sample libraries, the 3' mRNA-Seq Library Prep Kit Protocol for Ion Torrent (QuantSeq-LEXOGEN™ Vienna, Austria) was used according to manufacturer's instruction. Briefly, library generation was initiated by oligodT priming which contains the Ion Torrent compatible linker sequences. 5 to 500ng per 5μl of RNA from each sample was used to perform the first strand synthesis. After first strand synthesis any remaining RNA was removed and second strand synthesis was initiated by a random primer, containing Ion Torrent compatible linker sequences at its 5' end, and a sequence polymerase. In line barcodes were introduced at this point. Second strand synthesis was followed by a magnetic bead-based purification step and the resulted purified library was amplified for 14 cycles and re-purified. Quality and quantity of each library was assessed in a bioanalyzer using the DNA High Sensitivity Kit reagents and protocol (Agilent Technologies). The quantified libraries were pooled together at a final concentration of 7pM. The libraries pool was processed on the Ion Proton One Touch system where the libraries were templated and enriched using either the Ion PI™ Hi-Q™ OT2 200 Kit (ThermoFisher Scientific) and sequenced, with the Ion PI™ Hi-Q™ Sequencing 200 Kit on Ion Proton PI™ V2 chips (ThermoFisher Scientific) according to commercially available protocols. 3' RNA-sequencing was performed on an Ion Proton™ System [13], according to the manufacturer's instructions. Initial analysis took place in Ion Torrent server.

**Single cell RNA-seq data re-analysis**

Single cell RNA-seq data were downloaded from GSE122960 and processed with the R package Seurat (v.3.1.2 & 4.0.5) [14,15]. A similar to the original data analysis strategy was applied. Initially, each sample was processed on its own. After removing low-quality cells and genes, data were normalized using the LogNormalize method of Seurat and then top variable features were selected using the vst method. Data were scaled prior to principal component analysis (PCA) application and selection of the top principal components. The latter were used for a Shared Nearest Neighbor (SNN) graph-guided cell clustering and last, t-SNE dimensionality reduction was performed. Cell typing followed that of the original analysis as much as possible. Samples integration was performed with the standard Seurat v3 integration pipeline first for samples within each and then across phenotypes (donor and IPF). Integrated data were re-scaled, clustering and dimensionality reduction were repeated. Cell types inherited from single-sample analysis were validated and corrected whenever required according to the original publication of the dataset. Marker genes were identified using the Wilcoxon Rank Sum test applied on the "RNA" slot of the integrated samples object. Absolute fold change of at least 1.2 on natural scale and Bonferroni-corrected p-value less than 0.05 were used as differential expression thresholds. Fibroblast sub-clusters were identified with a resolution of 0.1 after fibroblast cells were isolated from the rest of the dataset, re-scaled and new variable features were found as above.

**Data processing**

Quant-Seq (Lexogen) FASTQ files obtained from the Ion Proton sequencing procedure were trimmed with Trim Galore (v.0.6.51) to remove low quality read ends using a Phred score of 20. Subsequently, a two steps alignment procedure was applied. Pre-processed reads were aligned against the GRCm38 reference genome (Ensembl) with HISAT2 (v.2.2.1) [16] and then the reads left unmapped were subjected to a second alignment round using BOWTIE2 (v.2.3.5.1) [17] with the --local and --very-sensitive local switches turned on.

**Computational analysis**

Downstream analysis of the resulting BAM files was performed with metaseqR2 (v.1.9.2) [18]. Briefly, the raw BAM files, one per sample, were summarized to a 3' UTR reads count table, using the package GenomicRanges (v1.44.0) [19] and Ensembl mouse genome mm10. For the UTR counting, the entire 3' UTR region, with a minimum length of 300 base pairs and 50 base pairs to flank the UTR end, was taken into consideration. In the resulting reads count table, each row represented one 3' UTR region and each column a Quant-Seq sample. Next, reads were summarized per gene and the returned gene count table was normalized using the package EDA-Seq (v.2.26.1) [20] after removing genes having zero reads across all samples. Post-normalization, gene counts were filtered for possible artifacts using default gene filtering options. The filtered gene counts table was subjected to differential expression analysis using sequentially all nine individual statistical analysis methods supported by metaseqR2. Their p-values were then combined by the PANDORA algorithm to account, among others, for the false positives reported. Benjamini-Hochberg corrected PANDORA p-values of less than 0.05 and absolute fold change of at least 1.2 were used as differential expression thresholds. Normalized expression values required for each heatmap were retrieved and standardized across samples. Hierarchical clustering of samples and genes based on calculated Euclidean distance.

**Gene Set Enrichment Analysis**

Differential expression analysis results were sorted by decreasing fold change and used for Gene Set Enrichment Analysis (GSEA) against Gene Ontology terms [21] using the package clusterProfiler (v.4.0.5) [22]. Signed normalized enrichment score (NES) was used to isolate the top of the significantly enriched induced (NES > 0) and suppressed (NES < 0) terms (adjusted p-value < 0.05).

#### Text mining

PubMed 2022 baseline was downloaded from the respective FTP site. XML R package (v.3.99.0.8) was used to create an abstract-based corpus which was then queried with rentrez R package (v.1.2.3) for *IPF[All Fields] OR (\"pulmonary fibrosis\"[MeSH Terms] OR \"pulmonary fibrosis\"[All Fields]) OR (\"lung diseases, interstitial\"[MeSH Terms] OR \"interstitial lung diseases\"[All Fields] OR \"interstitial lung disease\"[All Fields])* containing elements. Subsequently, human HGNC gene symbols atomization was performed using pubmed.mineR package [23] (v.1.0.19) and recovered genes were intersected with the human homologues of the mouse Quant-seq differentially expressed genes. Homologue feature mapping was obtained via biomaRt R package [24] (v. 2.48.3).

#### Transcription factor analysis

Transcription factor analysis was performed using DoRothEA R package [25] (v.1.4.2). All mouse transcription factor-target regulons were queried for those high quality ones ("A" level of confidence) including as targets any of the Quant-seq differentially expressed genes. Subsequent filtering maintained those interaction pairs where both interactors were found significantly deregulated in the bulk sequencing experiment. Based on the mode of regulation (mor) as described in the DoRothEA database and the Quant-seq derived differential expression data, pairs with the same or opposite direction of deregulation were maintained in cases of an activator or repressor transcription factor, respectively.

#### CMap/LINCS analysis

CMap/LINCS database (https://clue.io/query) query was performed using the top 150 up and top 150 down regulated genes as sorted by fold change. 150 consisted of the maximum number of genes supported by the platform. L1000 gene expression data from the Expanded CMap LINCS Resource 2020 (last update 11/23/2021) were queried. Results were processed by the R package cmapR (v.1.4.0). FDR-corrected p-values less than 0.05 were used to isolate those signatures having a statistically important connection with the provided one and signed normalized connectivity score (NCS) was used to discern between similar and opposite signatures. Focus was given on compounds (trt_cp) and peptides (trt_lig) having an already known mechanism of action and known targets. Datasets comprising of only one replicate and/or of a treatment duration other than 24 hours were discarded.
