## Supplementary material for "SRC-mediated and TKS5-enabled podosome formation is an inherent property of IPF fibroblasts, promoting ECM invasion and pulmonary fibrosis": All supplementary Tables

**Table S1.** ***TKS5 mRNA*** **expression in cohorts of IPF patients** as compared with the corresponding respective controls. All re-normalized and re-analyzed datasets are sourced from Fibromine (Fanidis, Moulos et al. 2021).

| PMID | Samples | GEO dataset | Platform | IPF/Ctrl | FC | FDR |
|---|---|---|---|---|---|---|
| 19363140 | Lungs of patients collected during pulmonary transplant compared with control ones derived from cancer free lung margins. | GSE10667 | GPL4133 | 21/14 | 1.29 | 3.43E-02 |
| 30111332 | Fresh frozen lung samples from IPF patients were compared to normal lung tissue flanking pulmonary cancer areas. | GSE110147 | GPL6244 | 22/11 | -1.23 | 1.95E-03 |
| 21241464 | IPF lung tissue originated from LTRC were assessed in comparison to healthy lung tissue of cancer patients and a lung transplant. | GSE21369 | GPL570 | 11/6 | 2.09 | 4.81E-02 |
| 21974901 | IPF lung samples transcriptional profile was compared to that of control samples from lung volume reduction during transplantation. | GSE24206 | GPL570 | 8/6 | 2.10 | 1.08E-02 |
| 26560100 | IPF lung samples were compared to healthy ones collected during an exploratory surgery. | GSE47460 | GPL6480 | 28/15 | 1.23 | 3.69E-03 |
| 26560100 | | GSE47460 | GPL14550 | 84/75 | 1.31 | 1.54E-20 |
| 21360508 | Microarray data from IPF patients were juxtaposed to data from control individuals. | GSE48149 | GPL16221 | 12/5 | 1.56 | 2.79E-03 |
| 25217476 | Differences between IPF and control transcriptomic profile were examined based on lung biopsies or explants. | GSE53845 | GPL6480 | 39/7 | 1.28 | 1.24E-02 |
| 28230051 | LTRC IPF and control samples were examined to identify transcriptional differences between disease and steady state conditions. | GSE92592 | GPL11154 | 19/18 | 1.78 | 4.24E-06 |
| 29329637 | Scarred regions from explanted lungs were compared to healthy control lung tissues. | GSE99621 | GPL16791 | 8/8 | 1.52 | 3.70E-03 |

*PMID: PubMed ID; GEO: gene expression database - datasets are hyperlinked; Platform: the hyperlinked numbers indicate the profiling technology; FC: fold change; FDR: false discovery rate; IPF: idiopathic pulmonary fibrosis; LTRC: Lung Tissue Research Consortium.*

**Table S2. Demographics and clinical characteristics of lung tissue donors.**

| Characteristic | IPF (*n*=20) | Control (*n*=9) | COPD (n=19) |
|---|---|---|---|
| Age (yr)(Mean ± SD) | 64.8 ± 8.3 | 68.8 ± 13.9 | 67.9 ± 14.1 |
| Gender, *n* (%) | | | |
| Males | 13 (65%) | 4 (44.4) | 14 (73.6%) |
| Females | 7 (35%) | 50 (55.6) | 5 (26.4%) |
| Race, *n (%)* | | | |
| White | 19 (95%) | 9 (100%) | 19 (100%) |
| Hispanic | 1 (5%) | 0 | 0 |
| Pulmonary function tests (Mean ± SD) | | | |
| FVC% | 66 ± 14 | 94 ± 16 | 76.4 ± 14.2 |
| DLCO% | 46 ± 15 | 86± 13 | 58.9± 21.2 |
| FEV1% | 73 ± 14 | 93 ± 11 | 53.3 ± 21.2 |

**FVC%: Forced vital capacity percent predicted; DLCO%: Carbon monoxide diffusing capacity percent predicted; FEV1% Forced expiratory volume in 1 second percent predicted.*

**Table S3. Demographics and clinical characteristics of lung fibroblasts donors**

| Characteristic | IPF (*n*=5) | Control (*n*=5) |
|---|---|---|
| Age (yr)(Mean ± SD) | 73.8± 20 | 64.8 ± 10 |
| Gender, *n* (%) | | |
| Males | 4 (57%) | 3 (60%) |
| Females | 3 (43%) | 2 (40%) |
| Pulmonary function tests (Mean ± SD) | | |
| DLCO% | 48 ± 11 | 52 ± 12 |
| FEV1/FVC% | 96 ± 7** | 72 ± 10 |
| Podosomes | | |
| % Cells with podosomes/area | 52 ± 12**** | 10.4 ± 4 |
| Podosomes/cell | 3±1**** | 1±1 |

**FVC%: Forced vital capacity percent predicted, DLCO%: Carbon monoxide diffusing capacity percent predicted. FEV1% Forced expiratory volume in 1 second percent predicted. Statistical significance was assessed between IPF and control groups with unpaired t-test followed by Welch's correction; **/***denote p<0.01/0.0001.*

**Table S4: Deregulated genes upon Tks5 haploinsufficiency and treatment with TGFb**

**Symbol:** HGNC gene symbol
**FDR:** Bonferroni-Hochberg corrected PANDORA p-value
**Fold Change:** KO-TGFb vs WT-TGFb fold change
**Description:** Gene description adapted from the Ensembl-provided one
**TF target?** Whether, according to DoRothEA, the gene participates as a transcription factor target in "A" level confidence interactions with any other gene
**TF ?** Whether, according to DoRothEA, the gene participates as a transcription factor in "A" level confidence interactions with any other gene of this list (
**Text mining:** How many times a gene's human homologue was reported as an HGNC symbol in PubMed baseline 2022 abstracts relative to IPF, pulmona

| Symbol | FDR | FoldChange | Description | TF target? | TF? | Text mining |
|---|---|---|---|---|---|---|
| Saa3 | 3,70E-53 | -154,56 | serum amyloid A 3 | No | No | - |
| Itgax | 1,05E-07 | -114,00 | integrin alpha X | Yes | No | - |
| Cybb | 2,91E-11 | -74,29 | cytochrome b-245, beta polypeptide | No | No | - |
| Lrp2 | 1,09E-06 | -54,00 | low density lipoprotein receptor-related protein 2 | No | No | - |
| Lrg1 | 9,03E-06 | -50,00 | leucine-rich alpha-2-glycoprotein 1 | No | No | - |
| Tnfsf18 | 3,00E-06 | -50,00 | tumor necrosis factor (ligand) superfamily, member 18 | No | No | - |
| C3 | 4,98E-34 | -47,72 | complement component 3 | No | No | 270 |
| Clec7a | 1,55E-06 | -41,50 | C-type lectin domain family 7, member a | No | No | - |
| Krt18 | 1,04E-05 | -38,00 | keratin 18 | Yes | No | - |
| Cd93 | 2,01E-05 | -36,00 | CD93 antigen | No | No | - |
| Hp | 4,97E-28 | -35,87 | haptoglobin | Yes | No | 2282 |
| Gpnmb | 3,67E-19 | -35,83 | glycoprotein (transmembrane) nmb | No | No | - |
| Cd68 | 4,14E-07 | -35,50 | CD68 antigen | No | No | - |
| Pex5l | 1,74E-07 | -33,33 | peroxisomal biogenesis factor 5-like | No | No | - |
| Gm20547 | 1,00E-29 | -30,52 | predicted gene 20547 | No | No | - |
| Gp1bb | 7,49E-07 | -29,00 | glycoprotein Ib, beta polypeptide | No | No | - |
| Cfb | 3,14E-30 | -28,69 | complement factor B | Yes | No | - |
| Gm36963 | 2,13E-05 | -28,50 | predicted gene, 36963 | No | No | - |
| Laptm5 | 1,57E-10 | -28,32 | lysosomal-associated protein transmembrane 5 | No | No | - |
| Nelfe | 1,52E-28 | -27,19 | negative elongation factor complex member E, Rdbp | No | No | - |
| Gm49601 | 5,65E-06 | -26,50 | predicted gene, 49601 | No | No | - |
| Kdr | 1,45E-06 | -26,00 | kinase insert domain protein receptor | No | No | 14 |
| Col15a1 | 7,06E-05 | -25,50 | collagen, type XV, alpha 1 | No | No | 3 |
| C4b | 3,61E-13 | -25,27 | complement component 4B (Chido blood group) | No | No | 4 |
| Ifit3 | 1,66E-06 | -24,83 | interferon-induced protein with tetratricopeptide repeats 3 | Yes | No | - |
| Ntn4 | 7,98E-09 | -22,55 | netrin 4 | No | No | - |
| Rspo1 | 9,49E-07 | -22,00 | R-spondin 1 | No | No | - |
| Cmklr1 | 6,28E-10 | -20,43 | chemokine-like receptor 1 | No | No | - |
| Wfdc17 | 9,40E-06 | -20,40 | WAP four-disulfide core domain 17 | No | No | - |
| Epas1 | 5,71E-06 | -19,00 | endothelial PAS domain protein 1 | No | No | 1 |
| Prl3a1 | 4,66E-04 | -19,00 | prolactin family 3, subfamily a, member 1 | No | No | - |
| Cldn4 | 1,49E-05 | -18,25 | claudin 4 | No | No | - |
| Emcn | 9,37E-05 | -18,00 | endomucin | No | No | - |
| Samd9l | 2,66E-08 | -17,17 | sterile alpha motif domain containing 9-like | No | No | 2 |
| Ifit3b | 3,81E-05 | -16,10 | interferon-induced protein with tetratricopeptide repeats 3B | Yes | No | - |
| Lcp1 | 3,58E-06 | -16,10 | lymphocyte cytosolic protein 1 | No | No | - |
| Gm42790 | 3,50E-06 | -16,02 | predicted gene 42790 | No | No | - |
| Cilp | 4,83E-08 | -16,00 | cartilage intermediate layer protein, nucleotide pyrophosphc | No | No | - |
| Sema3b | 4,60E-05 | -15,25 | sema domain, immunoglobulin domain (Ig), short basic don | No | No | - |
| Ctss | 2,75E-07 | -15,23 | cathepsin S | No | No | 25 |
| Oasl2 | 9,06E-07 | -15,00 | 2'-5' oligoadenylate synthetase-like 2 | No | No | - |
| S100a8 | 6,69E-07 | -14,82 | S100 calcium binding protein A8 (calgranulin A) | No | No | 15 |
| Dync1h1 | 5,97E-33 | -14,49 | dynein cytoplasmic 1 heavy chain 1 | No | No | - |
| Gm6712 | 3,41E-04 | -14,00 | predicted gene 6712 | No | No | - |
| Zfp874a | 3,44E-04 | -14,00 | zinc finger protein 874a | No | No | - |
| Rasip1 | 2,97E-06 | -13,89 | Ras interacting protein 1 | No | No | - |
| Lbp | 1,06E-05 | -13,40 | lipopolysaccharide binding protein | Yes | No | 9 |
| Adamts8 | 6,23E-05 | -13,00 | a disintegrin-like and metallopeptidase (reprolysin type) wit | No | No | - |
| Mmp12 | 9,42E-10 | -13,00 | matrix metallopeptidase 12 | No | No | 18 |
| Cdh6 | 5,42E-07 | -12,88 | cadherin 6 | No | No | - |
| Cpxm2 | 2,48E-06 | -12,67 | carboxypeptidase X 2 (M14 family) | No | No | - |
| Pf4 | 2,37E-04 | -12,67 | platelet factor 4 | No | No | - |
| Slc1a6 | 7,96E-08 | -12,26 | solute carrier family 1 (high affinity aspartate/glutamate trai | No | No | - |
| Mmp8 | 1,48E-17 | -12,22 | matrix metallopeptidase 8 | No | No | 2 |
| Clec4d | 3,32E-06 | -11,88 | C-type lectin domain family 4, member d | No | No | - |
| Eln | 1,83E-15 | -11,29 | elastin | Yes | No | - |
| Cfh | 1,02E-04 | -11,20 | complement component factor h | No | No | - |
| Aldh1a1 | 1,04E-06 | -11,14 | aldehyde dehydrogenase family 1, subfamily A1 | No | No | 1 |
| Efemp1 | 4,90E-07 | -11,14 | epidermal growth factor-containing fibulin-like extracellula | No | No | - |
| Col8a2 | 6,90E-04 | -11,00 | collagen, type VIII, alpha 2 | No | No | - |
| Plce1 | 5,21E-04 | -11,00 | phospholipase C, epsilon 1 | No | No | - |
| Nkx6-1 | 3,26E-04 | -10,80 | NK6 homeobox 1 | No | No | - |
| Trim30a | 1,44E-04 | -10,80 | tripartite motif-containing 30A | No | No | - |
| Ifi44 | 1,48E-04 | -10,78 | interferon-induced protein 44 | No | No | 3 |

| | | | | | | |
|---|---|---|---|---|---|---|
| Gal3st2 | 3,02E-06 | -10,77 | galactose-3-O-sulfotransferase 2 | No | No | - |
| Ifit1 | 3,01E-05 | -10,71 | interferon-induced protein with tetratricopeptide repeats 1 | No | No | - |
| Ifi27l2a | 6,00E-06 | -10,60 | interferon, alpha-inducible protein 27 like 2A | No | No | - |
| Sfn | 3,05E-16 | -10,59 | stratifin | No | No | 36 |
| Lama4 | 3,51E-08 | -10,55 | laminin, alpha 4 | No | No | - |
| Nkd2 | 2,59E-11 | -10,54 | naked cuticle 2 | No | No | - |
| Lpin1 | 1,68E-10 | -10,50 | lipin 1 | No | No | - |
| Cspg4 | 1,34E-07 | -10,45 | chondroitin sulfate proteoglycan 4 | No | No | 1 |
| Slfn5 | 1,94E-07 | -10,45 | schlafen 5 | No | No | - |
| Prelp | 2,09E-08 | -10,18 | proline arginine-rich end leucine-rich repeat | No | No | - |
| Ctsd | 8,12E-16 | -10,18 | cathepsin D | Yes | No | - |
| Lyz2 | 5,38E-07 | -9,63 | lysozyme 2 | No | No | 3 |
| Arap3 | 4,81E-04 | -9,60 | ArfGAP with RhoGAP domain, ankyrin repeat and PH dom | No | No | - |
| Slc7a11 | 2,11E-06 | -9,28 | solute carrier family 7 (cationic amino acid transporter, y+ s | No | No | 4 |
| Pcsk6 | 4,24E-07 | -9,24 | proprotein convertase subtilisin/kexin type 6 | No | No | - |
| Pramef12 | 2,15E-03 | -9,00 | PRAME like 13 | No | No | - |
| Gm5611 | 4,64E-10 | -8,94 | predicted gene 5611 | No | No | - |
| Trim12c | 1,01E-04 | -8,86 | tripartite motif-containing 12C | No | No | - |
| Scn3b | 1,16E-03 | -8,83 | sodium channel, voltage-gated, type III, beta | No | No | - |
| Cp | 4,39E-09 | -8,68 | ceruloplasmin | No | No | 131 |
| Bst2 | 1,98E-14 | -8,63 | bone marrow stromal cell antigen 2 | Yes | No | - |
| Lamb2 | 2,52E-14 | -8,55 | laminin, beta 2 | No | No | 1 |
| Ndor1 | 3,18E-03 | -8,50 | NADPH dependent diflavin oxidoreductase 1 | No | No | - |
| Ndor1 | 1,27E-03 | -8,40 | NADPH dependent diflavin oxidoreductase 1 | No | No | - |
| Col5a3 | 4,14E-15 | -8,39 | collagen, type V, alpha 3 | No | No | - |
| Cdo1 | 1,02E-14 | -8,36 | cysteine dioxygenase 1, cytosolic | No | No | - |
| Sp110 | 6,80E-10 | -8,35 | Sp110 nuclear body protein | No | No | - |
| Aldh2 | 1,73E-17 | -8,31 | aldehyde dehydrogenase 2, mitochondrial | No | No | - |
| Gas6 | 3,34E-19 | -8,30 | growth arrest specific 6 | No | No | - |
| Upk3b | 3,13E-04 | -8,29 | uroplakin 3B | No | No | - |
| Fam171a2 | 1,33E-17 | -7,96 | family with sequence similarity 171, member A2 | No | No | - |
| P2ry6 | 1,54E-06 | -7,94 | pyrimidinergic receptor P2Y, G-protein coupled, 6 | No | No | - |
| Gm4070 | 7,91E-06 | -7,91 | GTPase, very large interferon inducible, family member 2 | No | No | - |
| Gstm1 | 1,42E-13 | -7,87 | glutathione S-transferase, mu 1 | No | No | 30 |
| Grn | 1,07E-17 | -7,82 | granulin | No | No | - |
| Anxa8 | 4,35E-09 | -7,71 | annexin A8 | No | No | - |
| Itga11 | 1,64E-08 | -7,68 | integrin alpha 11 | No | No | - |
| Faah | 5,39E-06 | -7,63 | fatty acid amide hydrolase | No | No | - |
| Tgfbi | 1,90E-04 | -7,62 | transforming growth factor, beta induced | No | No | 1 |
| Galns | 3,43E-05 | -7,62 | galactosamine (N-acetyl)-6-sulfate sulfatase | No | No | - |
| Mphosph8 | 1,88E-07 | -7,57 | M-phase phosphoprotein 8 | No | No | - |
| Usp53 | 6,20E-04 | -7,57 | ubiquitin specific peptidase 53 | No | No | - |
| Fam20c | 8,59E-08 | -7,46 | FAM20C, golgi associated secretory pathway kinase | No | No | - |
| Usp18 | 1,66E-05 | -7,41 | ubiquitin specific peptidase 18 | No | No | 3 |
| Rapgef3 | 3,16E-10 | -7,39 | Rap guanine nucleotide exchange factor (GEF) 3 | No | No | - |
| Podxl | 1,92E-03 | -7,33 | podocalyxin-like | No | No | - |
| Slc39a5 | 5,69E-06 | -7,27 | solute carrier family 39 (metal ion transporter), member 5 | No | No | - |
| Cotl1 | 5,67E-15 | -7,24 | coactosin-like 1 (Dictyostelium) | No | No | - |
| Dnaaf1 | 4,98E-05 | -7,23 | dynein, axonemal assembly factor 1 | No | No | - |
| Clu | 6,51E-06 | -7,13 | clusterin | Yes | No | - |
| Syn2 | 5,42E-07 | -7,11 | synapsin II | Yes | No | - |
| Taf1c | 5,00E-05 | -7,07 | TATA-box binding protein associated factor, RNA polymer | No | No | - |
| Gm19090 | 5,23E-05 | -7,00 | predicted gene, 19090 | No | No | - |
| Ass1 | 1,43E-08 | -6,97 | argininosuccinate synthetase 1 | No | No | 6 |
| Serpina3n | 1,68E-10 | -6,96 | serine (or cysteine) peptidase inhibitor, clade A, member 3N | No | No | 5 |
| Endod1 | 7,69E-09 | -6,95 | endonuclease domain containing 1 | No | No | - |
| Frrs1 | 1,83E-14 | -6,86 | ferric-chelate reductase 1 | No | No | - |
| Rtp4 | 7,95E-06 | -6,82 | receptor transporter protein 4 | No | No | - |
| Ddx3y | 3,15E-14 | -6,79 | DEAD box helicase 3, Y-linked | No | No | - |
| 2610008E11Rik | 5,29E-04 | -6,67 | RIKEN cDNA 2610008E11 gene | No | No | - |
| Mfge8 | 1,87E-17 | -6,62 | milk fat globule EGF and factor V/VIII domain containing | No | No | - |
| Atp2b4 | 4,62E-04 | -6,60 | ATPase, Ca++ transporting, plasma membrane 4 | Yes | No | - |
| Cdk5rap2 | 1,45E-03 | -6,57 | CDK5 regulatory subunit associated protein 2 | No | No | - |
| Slpi | 7,02E-04 | -6,47 | secretory leukocyte peptidase inhibitor | No | No | 44 |
| Pnpla7 | 3,36E-04 | -6,43 | patatin-like phospholipase domain containing 7 | No | No | - |
| Ltbp2 | 3,05E-16 | -6,34 | latent transforming growth factor beta binding protein 2 | No | No | 7 |
| Crebrf | 1,19E-03 | -6,33 | CREB3 regulatory factor | No | No | - |
| Mgll | 3,60E-13 | -6,32 | monoglyceride lipase | No | No | - |
| Gbp2 | 2,42E-03 | -6,25 | guanylate binding protein 2 | No | No | - |
| Ces2g | 1,33E-03 | -6,22 | carboxylesterase 2G | No | No | - |
| Ldlr | 3,67E-19 | -6,20 | low density lipoprotein receptor | Yes | No | 1 |
| Gm15217 | 2,38E-04 | -6,18 | predicted gene 15217 | No | No | - |
| Bmp4 | 3,17E-04 | -6,12 | bone morphogenetic protein 4 | No | No | 4 |

| | | | | | | |
|---|---|---|---|---|---|---|
| Tcn2 | 2,11E-10 | -6,11 | transcobalamin 2 | Yes | No | - |
| Ccl2 | 9,17E-15 | -6,07 | chemokine (C-C motif) ligand 2 | Yes | No | - |
| Neat1 | 2,37E-10 | -6,03 | nuclear paraspeckle assembly transcript 1 (non-protein codi | No | No | - |
| Bcl2a1d | 6,23E-03 | -6,00 | B cell leukemia/lymphoma 2 related protein A1d | No | No | - |
| Npy1r | 3,92E-08 | -6,00 | neuropeptide Y receptor Y1 | No | No | - |
| Rasgrp3 | 2,97E-04 | -6,00 | RAS, guanyl releasing protein 3 | No | No | - |
| Stag2 | 2,51E-04 | -5,95 | stromal antigen 2 | No | No | - |
| Pamr1 | 4,69E-05 | -5,89 | peptidase domain containing associated with muscle regene | No | No | - |
| Cpxm1 | 1,24E-06 | -5,88 | carboxypeptidase X 1 (M14 family) | No | No | - |
| Bgn | 1,83E-15 | -5,87 | biglycan | No | No | 1 |
| Gsta3 | 3,43E-03 | -5,83 | glutathione S-transferase, alpha 3 | No | No | - |
| Tapbp | 5,24E-13 | -5,83 | TAP binding protein | No | No | - |
| Enpp1 | 1,04E-04 | -5,81 | ectonucleotide pyrophosphatase/phosphodiesterase 1 | No | No | - |
| Lat2 | 6,90E-04 | -5,79 | linker for activation of T cells family, member 2 | No | No | - |
| Shroom3 | 1,28E-05 | -5,77 | shroom family member 3 | No | No | - |
| Mysm1 | 3,11E-03 | -5,75 | myb-like, SWIRM and MPN domains 1 | No | No | - |
| Fastkd2 | 1,69E-03 | -5,70 | FAST kinase domains 2 | No | No | - |
| Clec4n | 5,18E-04 | -5,69 | C-type lectin domain family 4, member n | No | No | - |
| Diaph2 | 2,74E-06 | -5,67 | diaphanous related formin 2 | No | No | - |
| Lgals3bp | 2,97E-09 | -5,64 | lectin, galactoside-binding, soluble, 3 binding protein | No | No | - |
| P2ry14 | 1,46E-05 | -5,64 | purinergic receptor P2Y, G-protein coupled, 14 | No | No | - |
| Smg1 | 6,12E-05 | -5,63 | SMG1 nonsense mediated mRNA decay associated PI3K re | No | No | - |
| Uchl1 | 2,47E-09 | -5,59 | ubiquitin carboxy-terminal hydrolase L1 | No | No | 7 |
| Fbn1 | 7,87E-16 | -5,56 | fibrillin 1 | No | No | 3 |
| Adamts5 | 1,26E-07 | -5,54 | a disintegrin-like and metallopeptidase (reprolysin type) wit | No | No | - |
| Stk3 | 2,05E-04 | -5,53 | serine/threonine kinase 3 | No | No | 1 |
| Sgsm1 | 2,64E-04 | -5,50 | small G protein signaling modulator 1 | No | No | - |
| Rhbdf1 | 1,91E-09 | -5,46 | rhomboid 5 homolog 1 | No | No | - |
| Cep57 | 3,95E-05 | -5,45 | centrosomal protein 57 | No | No | - |
| Aplp2 | 1,65E-09 | -5,43 | amyloid beta (A4) precursor-like protein 2 | No | No | - |
| Cd52 | 5,83E-04 | -5,40 | CD52 antigen | No | No | - |
| Gm16973 | 8,26E-04 | -5,38 | predicted gene, 16973 | No | No | - |
| Ssc5d | 1,70E-05 | -5,38 | scavenger receptor cysteine rich family, 5 domains | No | No | - |
| Mvd | 4,28E-15 | -5,37 | mevalonate (diphospho) decarboxylase | No | No | 7 |
| Ubr4 | 1,99E-12 | -5,33 | ubiquitin protein ligase E3 component n-recognin 4 | No | No | - |
| Jag1 | 1,16E-06 | -5,28 | jagged 1 | No | No | 7 |
| Fn1 | 4,14E-15 | -5,27 | fibronectin 1 | Yes | No | 19 |
| Prkg2 | 1,92E-03 | -5,27 | protein kinase, cGMP-dependent, type II | No | No | - |
| Ncor2 | 2,56E-12 | -5,27 | nuclear receptor co-repressor 2 | No | No | - |
| Il17ra | 6,83E-15 | -5,26 | interleukin 17 receptor A | No | No | 7 |
| Lox | 7,49E-14 | -5,26 | lysyl oxidase | No | No | 83 |
| Adcyap1r1 | 1,63E-06 | -5,23 | adenylate cyclase activating polypeptide 1 receptor 1 | No | No | - |
| Serpine2 | 6,53E-12 | -5,23 | serine (or cysteine) peptidase inhibitor, clade E, member 2 | No | No | 5 |
| Lrpprc | 1,11E-12 | -5,21 | leucine-rich PPR-motif containing | No | No | - |
| Pik3c2a | 9,40E-07 | -5,21 | phosphatidylinositol-4-phosphate 3-kinase catalytic subunit | No | No | - |
| Igfbp2 | 2,92E-06 | -5,20 | insulin-like growth factor binding protein 2 | No | No | 2 |
| Pltp | 6,83E-15 | -5,18 | phospholipid transfer protein | No | No | 8 |
| Trim12a | 2,89E-03 | -5,18 | tripartite motif-containing 12A | No | No | - |
| Itga1 | 1,49E-05 | -5,17 | integrin alpha 1 | No | No | - |
| Zc3h11a | 1,54E-12 | -5,14 | zinc finger CCCH type containing 11A | No | No | - |
| Tyrobp | 7,17E-04 | -5,13 | TYRO protein tyrosine kinase binding protein | No | No | - |
| Slc5a3 | 5,68E-05 | -5,13 | solute carrier family 5 (inositol transporters), member 3 | No | No | - |
| Adgrd1 | 3,04E-09 | -5,12 | adhesion G protein-coupled receptor D1 | No | No | - |
| Gm38394 | 2,78E-12 | -5,12 | zinc finger, BED type containing 6 | No | No | - |
| Dop1b | 3,98E-03 | -5,11 | DOP1 leucine zipper like protein B | No | No | - |
| Ift74 | 2,60E-04 | -5,11 | intraflagellar transport 74 | No | No | - |
| Zbed4 | 4,50E-03 | -5,11 | zinc finger, BED type containing 4 | No | No | - |
| Zfp955a | 4,52E-03 | -5,11 | zinc finger protein 955A | No | No | - |
| Parm1 | 6,58E-09 | -5,11 | prostate androgen-regulated mucin-like protein 1 | No | No | - |
| Slit2 | 1,37E-12 | -5,08 | slit guidance ligand 2 | No | No | 4 |
| Ahnak2 | 1,15E-11 | -5,07 | - | No | No | - |
| Hcst | 1,08E-03 | -5,06 | hematopoietic cell signal transducer | No | No | - |
| Ahnak | 2,60E-12 | -5,03 | AHNAK nucleoprotein (desmoyokin) | No | No | - |
| Ero1lb | 9,75E-03 | -5,00 | endoplasmic reticulum oxidoreductase 1 beta | No | No | - |
| Fmod | 1,48E-04 | -5,00 | fibromodulin | No | No | - |
| Plcl1 | 3,90E-04 | -5,00 | phospholipase C-like 1 | No | No | 3 |
| Zbtb20 | 9,51E-06 | -4,97 | zinc finger and BTB domain containing 20 | No | No | - |
| Col4a2 | 8,29E-13 | -4,96 | collagen, type IV, alpha 2 | No | No | - |
| Prrg3 | 1,77E-03 | -4,94 | proline rich Gla (G-carboxyglutamic acid) 3 (transmembran | No | No | - |
| Tlr3 | 1,60E-05 | -4,94 | toll-like receptor 3 | Yes | No | 49 |
| Nmi | 1,24E-03 | -4,92 | N-myc (and STAT) interactor | Yes | No | - |
| Ifitm10 | 6,51E-05 | -4,92 | interferon induced transmembrane protein 10 | No | No | - |
| Tgfb2 | 1,16E-07 | -4,90 | transforming growth factor, beta 2 | Yes | No | - |

| | | | | | | |
|---|---|---|---|---|---|---|
| Prex2 | 5,06E-03 | -4,90 | phosphatidylinositol-3,4,5-trisphosphate-dependent Rac exc | No | No | - |
| Zfp120 | 2,63E-04 | -4,90 | zinc finger protein 120 | No | No | - |
| C1ra | 2,77E-07 | -4,89 | complement component 1, r subcomponent A | Yes | No | - |
| Zc3h11a | 6,73E-12 | -4,89 | zinc finger CCCH type containing 11A | No | No | - |
| Zbp1 | 3,85E-04 | -4,83 | Z-DNA binding protein 1 | No | No | - |
| Ldlrad2 | 8,15E-08 | -4,82 | low density lipoprotein receptor class A domain containing | No | No | - |
| Ada | 2,23E-05 | -4,81 | adenosine deaminase | No | No | 25 |
| BC002059 | 1,76E-04 | -4,81 | cDNA sequence BC002059 | No | No | - |
| Adgrf1 | 5,03E-03 | -4,80 | adhesion G protein-coupled receptor F1 | No | No | - |
| Cpeb1 | 2,95E-06 | -4,80 | cytoplasmic polyadenylation element binding protein 1 | No | No | - |
| Aldh1a3 | 2,51E-09 | -4,79 | aldehyde dehydrogenase family 1, subfamily A3 | No | No | - |
| Mpp5 | 1,28E-06 | -4,79 | protein associated with LIN7 1, MAGUK family member | No | No | - |
| Ncl | 3,29E-12 | -4,78 | nucleolin | No | No | 2 |
| Ctsa | 6,13E-15 | -4,78 | cathepsin A | No | No | - |
| AI987944 | 1,63E-05 | -4,77 | expressed sequence AI987944 | No | No | - |
| Ccl8 | 1,20E-11 | -4,76 | chemokine (C-C motif) ligand 8 | No | No | - |
| Pold2 | 4,39E-09 | -4,72 | polymerase (DNA directed), delta 2, regulatory subunit | No | No | - |
| Sema7a | 1,58E-02 | -4,71 | sema domain, immunoglobulin domain (Ig), and GPI memb | No | No | 11 |
| Slc20a1 | 2,42E-08 | -4,71 | solute carrier family 20, member 1 | No | No | - |
| Tmem176a | 1,30E-08 | -4,71 | transmembrane protein 176A | No | No | - |
| Zfp451 | 4,40E-09 | -4,70 | zinc finger protein 451 | No | No | - |
| Fam160a1 | 3,74E-03 | -4,67 | FHF complex subunit HOOK interacting protein 1A | No | No | - |
| Sp100 | 5,24E-03 | -4,67 | nuclear antigen Sp100 | No | No | - |
| Fth1 | 2,62E-08 | -4,65 | ferritin heavy polypeptide 1 | No | No | - |
| Fcer1g | 2,01E-03 | -4,63 | Fc receptor, IgE, high affinity I, gamma polypeptide | No | No | - |
| Lamp1 | 3,52E-14 | -4,62 | lysosomal-associated membrane protein 1 | No | No | 3 |
| Upk1b | 7,16E-03 | -4,60 | uroplakin 1B | No | No | - |
| Plxdc2 | 2,42E-11 | -4,59 | plexin domain containing 2 | No | No | - |
| Trim16 | 7,86E-04 | -4,59 | tripartite motif-containing 16 | No | No | - |
| Best1 | 3,65E-08 | -4,59 | bestrophin 1 | No | No | - |
| Ism1 | 7,36E-06 | -4,58 | isthmin 1, angiogenesis inhibitor | No | No | - |
| Psen1 | 2,16E-05 | -4,57 | presenilin 1 | No | No | - |
| Slc9a8 | 1,03E-05 | -4,56 | solute carrier family 9 (sodium/hydrogen exchanger), memb | No | No | - |
| P2rx7 | 6,91E-04 | -4,56 | purinergic receptor P2X, ligand-gated ion channel, 7 | No | No | 1 |
| Mapk12 | 5,66E-03 | -4,55 | mitogen-activated protein kinase 12 | No | No | - |
| Mcoln1 | 5,65E-06 | -4,54 | mucolipin 1 | No | No | - |
| Plcb4 | 2,04E-03 | -4,50 | phospholipase C, beta 4 | No | No | 1 |
| Ptpn21 | 1,28E-05 | -4,50 | protein tyrosine phosphatase, non-receptor type 21 | No | No | - |
| Ptpre | 1,60E-02 | -4,50 | protein tyrosine phosphatase, receptor type, E | No | No | - |
| Serping1 | 3,81E-11 | -4,49 | serine (or cysteine) peptidase inhibitor, clade G, member 1 | No | No | - |
| Isg15 | 2,08E-05 | -4,49 | ISG15 ubiquitin-like modifier | Yes | No | 4 |
| Rock2 | 2,45E-12 | -4,49 | Rho-associated coiled-coil containing protein kinase 2 | No | No | 11 |
| Sod3 | 1,48E-10 | -4,48 | superoxide dismutase 3, extracellular | No | No | 12 |
| Clcn5 | 6,79E-07 | -4,45 | chloride channel, voltage-sensitive 5 | No | No | - |
| Mocos | 1,33E-02 | -4,44 | molybdenum cofactor sulfurase | No | No | - |
| 1700006J14Rik | 5,47E-03 | -4,43 | RIKEN cDNA 1700006J14 gene | No | No | - |
| Chpf | 2,60E-13 | -4,43 | chondroitin polymerizing factor | No | No | - |
| Rnf169 | 7,80E-05 | -4,42 | ring finger protein 169 | No | No | - |
| Gria3 | 2,64E-04 | -4,42 | glutamate receptor, ionotropic, AMPA3 (alpha 3) | No | No | - |
| Rgs4 | 9,42E-04 | -4,40 | regulator of G-protein signaling 4 | No | No | - |
| Syne2 | 7,07E-03 | -4,40 | spectrin repeat containing, nuclear envelope 2 | No | No | - |
| Sulf1 | 2,59E-11 | -4,39 | sulfatase 1 | No | No | 6 |
| Aebp1 | 7,43E-09 | -4,38 | AE binding protein 1 | Yes | No | - |
| Prox1 | 2,21E-03 | -4,37 | prospero homeobox 1 | No | No | - |
| Gipc3 | 6,13E-03 | -4,36 | GIPC PDZ domain containing family, member 3 | No | No | - |
| Cdk6 | 6,52E-05 | -4,35 | cyclin-dependent kinase 6 | Yes | No | 2 |
| Tab1 | 1,44E-02 | -4,33 | TGF-beta activated kinase 1/MAP3K7 binding protein 1 | No | No | - |
| Zfp955b | 1,23E-02 | -4,33 | zinc finger protein 955B | No | No | - |
| Svep1 | 1,90E-05 | -4,32 | sushi, von Willebrand factor type A, EGF and pentraxin dor | No | No | - |
| Dgcr8 | 5,33E-05 | -4,32 | DGCR8, microprocessor complex subunit | No | No | 2 |
| Tbxa2r | 3,01E-03 | -4,31 | thromboxane A2 receptor | No | No | - |
| Fnip1 | 2,57E-05 | -4,28 | folliculin interacting protein 1 | No | No | - |
| Scaf1 | 2,22E-03 | -4,26 | SR-related CTD-associated factor 1 | No | No | - |
| Ebf1 | 1,65E-02 | -4,25 | early B cell factor 1 | No | No | - |
| Tmc6 | 9,81E-05 | -4,25 | transmembrane channel-like gene family 6 | No | No | - |
| Zmym2 | 2,48E-06 | -4,25 | zinc finger, MYM-type 2 | No | No | - |
| Asic4 | 8,28E-13 | -4,25 | acid-sensing (proton-gated) ion channel family member 4 | No | No | - |
| Grtp1 | 2,27E-12 | -4,25 | GH regulated TBC protein 1 | No | No | - |
| St6galnac4 | 4,58E-06 | -4,24 | ST6 (alpha-N-acetyl-neuraminyl-2,3-beta-galactosyl-1,3)-N | No | No | - |
| Synpo2 | 2,62E-04 | -4,24 | synaptopodin 2 | No | No | - |
| Naglu | 4,01E-04 | -4,22 | alpha-N-acetylglucosaminidase (Sanfilippo disease IIIB) | No | No | - |
| Rasgrp2 | 1,57E-02 | -4,22 | RAS, guanyl releasing protein 2 | No | No | - |
| Ythdc2 | 1,34E-02 | -4,22 | YTH domain containing 2 | No | No | - |

| | | | | | | |
|---|---|---|---|---|---|---|
| Etv4 | 1,36E-03 | -4,19 | ets variant 4 | No | Yes | - |
| Flrt2 | 2,59E-11 | -4,17 | fibronectin leucine rich transmembrane protein 2 | No | No | - |
| Zfp719 | 3,68E-03 | -4,15 | zinc finger protein 719 | No | No | - |
| Dhcr24 | 2,04E-06 | -4,15 | 24-dehydrocholesterol reductase | No | No | - |
| Stat2 | 5,78E-03 | -4,13 | signal transducer and activator of transcription 2 | Yes | Yes | 2 |
| C77080 | 3,69E-03 | -4,13 | expressed sequence C77080 | No | No | - |
| Cep162 | 1,98E-02 | -4,13 | centrosomal protein 162 | No | No | - |
| Nuak2 | 1,67E-03 | -4,12 | NUAK family, SNF1-like kinase, 2 | No | No | - |
| Ecm1 | 1,97E-08 | -4,11 | extracellular matrix protein 1 | No | No | - |
| Ccl17 | 1,25E-02 | -4,09 | chemokine (C-C motif) ligand 17 | No | No | 51 |
| Mapk7 | 5,92E-08 | -4,09 | mitogen-activated protein kinase 7 | No | No | 3 |
| Trf | 1,86E-05 | -4,09 | transferrin | Yes | No | 87 |
| Apol6 | 3,26E-04 | -4,08 | apolipoprotein L 6 | No | No | - |
| Jph2 | 1,64E-08 | -4,05 | junctophilin 2 | No | No | - |
| Timp3 | 1,14E-10 | -4,04 | tissue inhibitor of metalloproteinase 3 | No | No | 18 |
| Lrrk1 | 3,52E-08 | -4,04 | leucine-rich repeat kinase 1 | No | No | - |
| Msr1 | 1,15E-02 | -4,03 | macrophage scavenger receptor 1 | Yes | No | 4 |
| Smtnl2 | 3,80E-04 | -4,03 | smoothelin-like 2 | No | No | - |
| Nupr1 | 2,30E-10 | -4,03 | nuclear protein transcription regulator 1 | Yes | No | - |
| Mfap4 | 4,50E-08 | -4,02 | microfibrillar-associated protein 4 | No | No | 8 |
| Nop14 | 5,91E-06 | -4,01 | NOP14 nucleolar protein | No | No | - |
| Boc | 3,16E-03 | -4,00 | biregional cell adhesion molecule-related/down-regulated b | No | No | 2 |
| Egr2 | 1,79E-03 | -4,00 | early growth response 2 | No | No | 3 |
| Iqsec1 | 1,13E-06 | -4,00 | IQ motif and Sec7 domain 1 | No | No | - |
| Mdga1 | 1,67E-03 | -4,00 | MAM domain containing glycosylphosphatidylinositol ancl | No | No | - |
| Plxnd1 | 1,81E-02 | -4,00 | plexin D1 | No | No | - |
| Pstpip1 | 7,52E-03 | -4,00 | proline-serine-threonine phosphatase-interacting protein 1 | No | No | - |
| Hbegf | 1,10E-04 | -3,97 | heparin-binding EGF-like growth factor | No | No | 2 |
| Pld3 | 9,83E-12 | -3,96 | phospholipase D family, member 3 | No | No | - |
| Sptbn1 | 4,28E-10 | -3,94 | spectrin beta, non-erythrocytic 1 | No | No | - |
| B3gnt8 | 3,64E-04 | -3,93 | UDP-GlcNAc:betaGal beta-1,3-N-acetylglucosaminyltransf | No | No | - |
| Zc3h12a | 4,66E-04 | -3,93 | zinc finger CCCH type containing 12A | No | No | 2 |
| Fcgr3 | 1,25E-02 | -3,92 | Fc receptor, IgG, low affinity III | No | No | - |
| Itpkc | 1,68E-02 | -3,90 | inositol 1,4,5-trisphosphate 3-kinase C | No | No | - |
| Wnt9a | 3,66E-08 | -3,89 | wingless-type MMTV integration site family, member 9A | No | No | - |
| Ablim1 | 1,95E-04 | -3,89 | actin-binding LIM protein 1 | No | No | - |
| B230354K17Rik | 4,36E-03 | -3,89 | RIKEN cDNA B230354K17 gene | No | No | - |
| Gm44503 | 3,46E-05 | -3,89 | predicted readthrough transcript (NMD candidate), 44503 | No | No | - |
| Slc18a2 | 1,98E-02 | -3,89 | solute carrier family 18 (vesicular monoamine), member 2 | No | No | - |
| Adam15 | 7,88E-11 | -3,89 | a disintegrin and metallopeptidase domain 15 (metargidin) | No | No | - |
| Tgfbr1 | 5,12E-05 | -3,88 | transforming growth factor, beta receptor I | No | No | 6 |
| Nexn | 8,93E-08 | -3,87 | nexilin | No | No | - |
| Trp53inp1 | 4,62E-06 | -3,86 | transformation related protein 53 inducible nuclear protein 1 | No | No | 1 |
| Eif2ak2 | 6,39E-04 | -3,86 | eukaryotic translation initiation factor 2-alpha kinase 2 | Yes | No | - |
| Psap | 3,90E-10 | -3,85 | prosaposin | No | No | 3 |
| Clcn7 | 3,31E-05 | -3,85 | chloride channel, voltage-sensitive 7 | No | No | - |
| Gadd45g | 3,16E-10 | -3,83 | growth arrest and DNA-damage-inducible 45 gamma | No | No | - |
| Nbeal1 | 1,85E-03 | -3,83 | neurobeachin like 1 | No | No | - |
| Ptar1 | 1,23E-04 | -3,82 | protein prenyltransferase alpha subunit repeat containing 1 | No | No | - |
| Aqp1 | 3,97E-04 | -3,82 | aquaporin 1 | No | No | 16 |
| Pwp2 | 5,87E-05 | -3,82 | PWP2 periodic tryptophan protein homolog (yeast) | Yes | No | - |
| Map3k14 | 1,45E-02 | -3,82 | mitogen-activated protein kinase kinase kinase 14 | No | No | - |
| Usp45 | 5,28E-03 | -3,81 | ubiquitin specific petidase 45 | No | No | - |
| Atp6ap1 | 2,59E-11 | -3,81 | ATPase, H+ transporting, lysosomal accessory protein 1 | No | No | - |
| Parva | 3,41E-06 | -3,81 | parvin, alpha | No | No | - |
| Bach1 | 2,12E-02 | -3,80 | BTB and CNC homology 1, basic leucine zipper transcriptio | No | No | - |
| Ppm1l | 2,35E-03 | -3,80 | protein phosphatase 1 (formerly 2C)-like | No | No | - |
| Cdh23 | 2,11E-10 | -3,80 | cadherin 23 (otocadherin) | No | No | 1 |
| Ece1 | 1,14E-08 | -3,78 | endothelin converting enzyme 1 | No | No | - |
| Adamts2 | 1,75E-05 | -3,77 | a disintegrin-like and metallopeptidase (reprolysin type) wit | No | No | - |
| Col4a1 | 5,94E-10 | -3,77 | collagen, type IV, alpha 1 | No | No | - |
| Zfp930 | 3,51E-03 | -3,76 | zinc finger protein 930 | No | No | - |
| Mmp2 | 2,28E-08 | -3,76 | matrix metallopeptidase 2 | Yes | No | 32 |
| Itga9 | 4,65E-05 | -3,75 | integrin alpha 9 | No | No | - |
| Zfp553 | 4,08E-03 | -3,74 | zinc finger protein 553 | No | No | - |
| Cpd | 7,09E-09 | -3,74 | carboxypeptidase D | No | No | 11 |
| Optn | 1,45E-02 | -3,71 | optineurin | No | No | 4 |
| Plec | 2,01E-08 | -3,70 | plectin | No | No | - |
| Hspg2 | 5,42E-07 | -3,69 | perlecan (heparan sulfate proteoglycan 2) | No | No | - |
| Cav2 | 2,96E-05 | -3,67 | caveolin 2 | No | No | - |
| Osr1 | 2,29E-04 | -3,67 | odd-skipped related transcription factor 1 | No | No | 4 |
| Arhgap5 | 7,94E-06 | -3,66 | Rho GTPase activating protein 5 | No | No | - |
| Slc12a4 | 1,51E-05 | -3,65 | solute carrier family 12, member 4 | No | No | - |

| | | | | | | |
|---|---|---|---|---|---|---|
| Tmem184a | 3,29E-05 | -3,65 | transmembrane protein 184a | No | No | - |
| Dlc1 | 2,21E-09 | -3,65 | deleted in liver cancer 1 | No | No | - |
| Gnl2 | 4,09E-05 | -3,64 | guanine nucleotide binding protein-like 2 (nucleolar) | No | No | - |
| BC034090 | 2,22E-02 | -3,64 | cDNA sequence BC034090 | No | No | - |
| Esco1 | 2,66E-02 | -3,64 | establishment of sister chromatid cohesion N-acetyltransfera | No | No | - |
| Il17rd | 3,51E-03 | -3,64 | interleukin 17 receptor D | No | No | - |
| Nrde2 | 1,92E-02 | -3,64 | nrde-2 necessary for RNA interference, domain containing | No | No | - |
| Olr1 | 1,07E-04 | -3,63 | oxidized low density lipoprotein (lectin-like) receptor 1 | No | No | - |
| Upf3b | 1,16E-03 | -3,63 | UPF3 regulator of nonsense transcripts homolog B (yeast) | No | No | - |
| Nckap1 | 1,54E-08 | -3,62 | NCK-associated protein 1 | No | No | - |
| Pmepa1 | 3,07E-12 | -3,61 | prostate transmembrane protein, androgen induced 1 | Yes | No | - |
| Snx19 | 2,08E-04 | -3,61 | sorting nexin 19 | No | No | - |
| Chm | 1,99E-02 | -3,60 | choroidermia (RAB escort protein 1) | No | No | - |
| Fxyd6 | 2,37E-06 | -3,60 | FXYD domain-containing ion transport regulator 6 | No | No | - |
| Erfe | 8,19E-04 | -3,59 | erythroferrone | No | No | - |
| Atrip | 1,06E-05 | -3,59 | ATR interacting protein | No | No | - |
| Adam8 | 8,08E-03 | -3,59 | a disintegrin and metallopeptidase domain 8 | No | No | 5 |
| Nsun2 | 5,81E-10 | -3,59 | NOL1/NOP2/Sun domain family member 2 | No | No | - |
| Ctla2a | 8,10E-08 | -3,59 | cytotoxic T lymphocyte-associated protein 2 alpha | No | No | - |
| Tef | 2,28E-07 | -3,59 | thyrotroph embryonic factor | No | No | 3 |
| Arhgdib | 3,52E-03 | -3,58 | Rho, GDP dissociation inhibitor (GDI) beta | No | No | - |
| Gas7 | 1,87E-06 | -3,57 | growth arrest specific 7 | Yes | No | - |
| Fasn | 1,25E-07 | -3,56 | fatty acid synthase | Yes | No | 7 |
| Slc6a8 | 2,42E-08 | -3,56 | solute carrier family 6 (neurotransmitter transporter, creatin | No | No | - |
| Atf5 | 1,80E-10 | -3,56 | activating transcription factor 5 | No | No | 1 |
| P4ha3 | 2,60E-08 | -3,54 | procollagen-proline, 2-oxoglutarate 4-dioxygenase (proline | No | No | 2 |
| Atp6ap2 | 3,72E-10 | -3,54 | ATPase, H+ transporting, lysosomal accessory protein 2 | No | No | - |
| Luc7l3 | 1,11E-03 | -3,54 | LUC7-like 3 (S. cerevisiae) | No | No | - |
| Erap1 | 4,51E-05 | -3,53 | endoplasmic reticulum aminopeptidase 1 | No | No | - |
| Mafk | 4,13E-05 | -3,53 | v-maf musculoaponeurotic fibrosarcoma oncogene family, p | No | No | - |
| Map6 | 3,84E-05 | -3,52 | microtubule-associated protein 6 | No | No | - |
| Malat1 | 1,05E-02 | -3,52 | metastasis associated lung adenocarcinoma transcript 1 (nor | No | No | - |
| Iqgap1 | 9,57E-08 | -3,51 | IQ motif containing GTPase activating protein 1 | No | No | 13 |
| Pgrmc2 | 4,20E-10 | -3,51 | progesterone receptor membrane component 2 | No | No | - |
| Crim1 | 2,36E-05 | -3,51 | cysteine rich transmembrane BMP regulator 1 (chordin like) | No | No | - |
| Ghr | 1,51E-05 | -3,51 | growth hormone receptor | No | No | 3 |
| Bcl2a1b | 1,27E-02 | -3,50 | B cell leukemia/lymphoma 2 related protein A1b | No | No | 5 |
| Gcn1 | 1,25E-03 | -3,50 | GCN1 activator of EIF2AK4 | No | No | - |
| Prrc2c | 9,13E-04 | -3,50 | proline-rich coiled-coil 2C | No | No | - |
| Tmem204 | 5,14E-03 | -3,50 | transmembrane protein 204 | No | No | - |
| Trex1 | 1,71E-05 | -3,50 | three prime repair exonuclease 1 | No | No | 1 |
| Pigt | 7,34E-06 | -3,50 | phosphatidylinositol glycan anchor biosynthesis, class T | No | No | - |
| Man2b1 | 3,66E-04 | -3,49 | mannosidase 2, alpha B1 | No | No | - |
| Akap11 | 1,56E-08 | -3,48 | A kinase (PRKA) anchor protein 11 | No | No | - |
| Gm6085 | 1,45E-02 | -3,47 | predicted gene 6085 | No | No | - |
| Kidins220 | 6,30E-07 | -3,46 | kinase D-interacting substrate 220 | No | No | - |
| Pbxip1 | 4,62E-10 | -3,46 | pre B cell leukemia transcription factor interacting protein 1 | Yes | No | - |
| Nt5c2 | 1,95E-03 | -3,46 | 5'-nucleotidase, cytosolic II | No | No | - |
| Grina | 5,57E-07 | -3,46 | glutamate receptor, ionotropic, N-methyl D-aspartate-associ | No | No | - |
| Tppp | 3,78E-02 | -3,45 | tubulin polymerization promoting protein | No | No | - |
| Trps1 | 2,85E-02 | -3,45 | transcriptional repressor GATA binding 1 | Yes | No | - |
| Clip2 | 6,87E-04 | -3,44 | CAP-GLY domain containing linker protein 2 | No | No | - |
| Trib1 | 6,67E-08 | -3,44 | tribbles pseudokinase 1 | No | No | - |
| Tnfrsf10b | 3,98E-05 | -3,44 | tumor necrosis factor receptor superfamily, member 10b | No | No | - |
| Galnt10 | 1,35E-07 | -3,43 | polypeptide N-acetylgalactosaminyltransferase 10 | No | No | - |
| Coro2b | 6,20E-03 | -3,43 | coronin, actin binding protein, 2B | No | No | - |
| Smcr8 | 1,50E-02 | -3,43 | Smith-Magenis syndrome chromosome region, candidate 8 | No | No | - |
| Tubb3 | 3,61E-05 | -3,42 | tubulin, beta 3 class III | No | No | - |
| Cxcl16 | 5,37E-08 | -3,42 | chemokine (C-X-C motif) ligand 16 | No | No | - |
| Cachd1 | 2,21E-03 | -3,42 | cache domain containing 1 | No | No | - |
| Prag1 | 2,68E-03 | -3,42 | PEAK1 related kinase activating pseudokinase 1 | No | No | - |
| Usp24 | 1,28E-05 | -3,41 | ubiquitin specific peptidase 24 | No | No | - |
| Psmd4 | 7,13E-09 | -3,40 | proteasome (prosome, macropain) 26S subunit, non-ATPase | No | No | - |
| Icam1 | 2,83E-05 | -3,40 | intercellular adhesion molecule 1 | Yes | No | 3 |
| Fat1 | 1,79E-07 | -3,40 | FAT atypical cadherin 1 | No | No | - |
| Prdm2 | 1,07E-04 | -3,39 | PR domain containing 2, with ZNF domain | No | No | - |
| Tnrc6c | 8,23E-04 | -3,39 | trinucleotide repeat containing 6C | No | No | - |
| Psmd13 | 1,15E-06 | -3,39 | proteasome (prosome, macropain) 26S subunit, non-ATPase | No | No | - |
| Gm45472 | 6,67E-08 | -3,38 | predicted gene 45472 | No | No | - |
| Ephb3 | 1,17E-06 | -3,38 | Eph receptor B3 | No | No | - |
| Col7a1 | 1,17E-03 | -3,38 | collagen, type VII, alpha 1 | No | No | - |
| C1s1 | 1,58E-05 | -3,38 | complement component 1, s subcomponent 1 | No | No | - |
| Mrnip | 6,48E-07 | -3,38 | MRN complex interacting protein | No | No | - |

| | | | | | | |
|---|---|---|---|---|---|---|
| 4930402H24Rik | 6,23E-08 | -3,38 | dynein axonemal assembly factor 9 | No | No | - |
| Spg11 | 1,89E-04 | -3,38 | SPG11, spatacsin vesicle trafficking associated | No | No | - |
| Zfhx3 | 1,76E-05 | -3,37 | zinc finger homeobox 3 | No | No | - |
| Serpina3g | 1,34E-04 | -3,36 | serine (or cysteine) peptidase inhibitor, clade A, member 3C | No | No | - |
| Shisa5 | 1,67E-07 | -3,36 | shisa family member 5 | No | No | - |
| Man1b1 | 8,76E-08 | -3,36 | mannosidase, alpha, class 1B, member 1 | No | No | - |
| Pcsk9 | 8,02E-07 | -3,36 | proprotein convertase subtilisin/kexin type 9 | Yes | No | - |
| Bmpr2 | 1,21E-07 | -3,34 | bone morphogenetic protein receptor, type II (serine/threoni | No | No | 21 |
| Il3ra | 5,83E-04 | -3,34 | interleukin 3 receptor, alpha chain | No | No | - |
| Ago3 | 1,27E-04 | -3,34 | argonaute RISC catalytic subunit 3 | No | No | - |
| Casp4 | 6,66E-03 | -3,33 | caspase 4, apoptosis-related cysteine peptidase | No | No | - |
| Epc2 | 2,71E-04 | -3,33 | enhancer of polycomb homolog 2 | No | No | - |
| Ezr | 1,50E-02 | -3,33 | ezrin | No | No | 2 |
| Sgf29 | 3,30E-03 | -3,33 | SAGA complex associated factor 29 | No | No | - |
| Tsku | 2,15E-03 | -3,33 | tsukushi, small leucine rich proteoglycan | No | No | - |
| Washc5 | 6,76E-07 | -3,33 | WASH complex subunit 5 | No | No | - |
| Treh | 4,97E-05 | -3,32 | trehalase (brush-border membrane glycoprotein) | No | No | - |
| 0610010F05Rik | 7,01E-03 | -3,32 | SANT and BTB domain regulator of CSR | No | No | - |
| Ctsf | 9,37E-04 | -3,31 | cathepsin F | No | No | - |
| Stox2 | 1,34E-03 | -3,31 | storkhead box 2 | No | No | - |
| Nub1 | 6,72E-06 | -3,31 | negative regulator of ubiquitin-like proteins 1 | No | No | - |
| Tkt | 4,20E-07 | -3,31 | transketolase | No | No | - |
| Sash1 | 3,55E-03 | -3,30 | SAM and SH3 domain containing 1 | No | No | - |
| Zbtb46 | 1,32E-02 | -3,30 | zinc finger and BTB domain containing 46 | No | No | - |
| Zfp35 | 6,48E-03 | -3,30 | zinc finger protein 35 | No | No | - |
| Ank | 2,08E-05 | -3,30 | progressive ankylosis | No | No | - |
| Tnip1 | 5,65E-08 | -3,30 | TNFAIP3 interacting protein 1 | No | No | - |
| Ints1 | 3,51E-04 | -3,30 | integrator complex subunit 1 | No | No | - |
| Brsk1 | 1,59E-02 | -3,29 | BR serine/threonine kinase 1 | No | No | - |
| Dnali1 | 3,71E-05 | -3,29 | dynein, axonemal, light intermediate polypeptide 1 | No | No | - |
| Cercam | 1,74E-04 | -3,29 | cerebral endothelial cell adhesion molecule | No | No | - |
| Bckdha | 2,21E-02 | -3,29 | branched chain ketoacid dehydrogenase E1, alpha polypepti | No | No | - |
| Cep250 | 2,39E-02 | -3,29 | centrosomal protein 250 | No | No | - |
| Slc6a20b | 1,59E-04 | -3,28 | solute carrier family 6 (neurotransmitter transporter), memb | No | No | - |
| 5830417I10Rik | 1,07E-04 | -3,28 | RIKEN cDNA 5830417I10 gene | No | No | - |
| Gns | 2,31E-07 | -3,27 | glucosamine (N-acetyl)-6-sulfatase | No | No | - |
| Ddx24 | 6,79E-04 | -3,27 | DEAD box helicase 24 | No | No | - |
| Ano10 | 1,65E-05 | -3,27 | anoctamin 10 | No | No | - |
| Gm16201 | 1,07E-02 | -3,27 | predicted gene 16201 | No | No | - |
| Ccpg1 | 9,31E-06 | -3,27 | cell cycle progression 1 | No | No | - |
| Pigb | 6,72E-06 | -3,27 | phosphatidylinositol glycan anchor biosynthesis, class B | No | No | - |
| Itpr2 | 9,16E-04 | -3,26 | inositol 1,4,5-triphosphate receptor 2 | No | No | - |
| Traf3 | 6,38E-05 | -3,26 | TNF receptor-associated factor 3 | No | No | - |
| Eif4g1 | 7,01E-07 | -3,26 | eukaryotic translation initiation factor 4, gamma 1 | No | No | - |
| Gaa | 4,57E-08 | -3,26 | glucosidase, alpha, acid | No | No | 5 |
| Marf1 | 3,04E-04 | -3,25 | meiosis regulator and mRNA stability 1 | No | No | - |
| Osbpl1a | 7,82E-03 | -3,25 | oxysterol binding protein-like 1A | No | No | - |
| Rpap3 | 1,45E-02 | -3,25 | RNA polymerase II associated protein 3 | No | No | - |
| Tax1bp1 | 2,15E-07 | -3,25 | Tax1 (human T cell leukemia virus type I) binding protein 1 | No | No | - |
| Mdm2 | 7,01E-07 | -3,24 | transformed mouse 3T3 cell double minute 2 | No | No | 9 |
| Ctns | 2,98E-04 | -3,24 | cystinosis, nephropathic | No | No | - |
| Luzp1 | 3,52E-06 | -3,24 | leucine zipper protein 1 | No | No | - |
| Gpatch4 | 1,14E-04 | -3,24 | G patch domain containing 4 | No | No | - |
| Sfrp1 | 7,40E-07 | -3,24 | secreted frizzled-related protein 1 | No | No | 8 |
| Ctcf | 5,65E-06 | -3,23 | CCCTC-binding factor | No | Yes | 1 |
| Cds1 | 2,36E-02 | -3,23 | CDP-diacylglycerol synthase 1 | No | No | - |
| Phldb1 | 5,14E-05 | -3,23 | pleckstrin homology like domain, family B, member 1 | No | No | - |
| Renbp | 1,72E-07 | -3,22 | renin binding protein | No | No | - |
| Kif5b | 5,61E-08 | -3,22 | kinesin family member 5B | No | No | - |
| Hps4 | 2,21E-02 | -3,21 | HPS4, biogenesis of lysosomal organelles complex 3 subun | No | No | 28 |
| Trappc9 | 2,33E-03 | -3,21 | trafficking protein particle complex 9 | No | No | - |
| Slc25a37 | 8,28E-04 | -3,21 | solute carrier family 25, member 37 | No | No | - |
| Cdkl2 | 1,06E-02 | -3,21 | cyclin-dependent kinase-like 2 (CDC2-related kinase) | No | No | - |
| Ift81 | 5,10E-03 | -3,21 | intraflagellar transport 81 | No | No | - |
| Sympk | 1,35E-03 | -3,21 | symplekin | No | No | - |
| Rlf | 1,10E-03 | -3,20 | rearranged L-myc fusion sequence | No | No | 10 |
| Ubc | 4,20E-08 | -3,20 | ubiquitin C | Yes | No | - |
| 2900026A02Rik | 7,06E-03 | -3,20 | RIKEN cDNA 2900026A02 gene | No | No | - |
| Ankrd28 | 3,11E-03 | -3,20 | ankyrin repeat domain 28 | No | No | - |
| Gm28627 | 7,80E-03 | -3,20 | predicted gene 28627 | No | No | - |
| Nsrp1 | 1,94E-02 | -3,20 | nuclear speckle regulatory protein 1 | No | No | - |
| Scrn1 | 4,58E-05 | -3,20 | secernin 1 | No | No | - |
| Tsc2 | 9,09E-07 | -3,20 | TSC complex subunit 2 | No | No | 25 |

| | | | | | | |
|---|---|---|---|---|---|---|
| Smyd5 | 1,68E-03 | -3,19 | SET and MYND domain containing 5 | No | No | - |
| Wdtc1 | 4,67E-05 | -3,19 | WD and tetratricopeptide repeats 1 | No | No | - |
| Asah1 | 1,89E-07 | -3,19 | N-acylsphingosine amidohydrolase 1 | No | No | - |
| Emc1 | 9,10E-07 | -3,19 | ER membrane protein complex subunit 1 | No | No | - |
| H2-D1 | 5,40E-07 | -3,18 | histocompatibility 2, D region locus 1 | No | No | 11 |
| Msn | 2,36E-10 | -3,18 | moesin | No | No | 7 |
| Snx8 | 1,85E-04 | -3,18 | sorting nexin 8 | No | No | - |
| Nfatc4 | 4,34E-05 | -3,18 | nuclear factor of activated T cells, cytoplasmic, calcineurin | No | No | - |
| Mfsd12 | 9,68E-03 | -3,17 | major facilitator superfamily domain containing 12 | No | No | - |
| Pkd1 | 8,10E-08 | -3,17 | polycystin 1, transient receptor potential channel interacting | No | No | 7 |
| Atp13a3 | 1,18E-05 | -3,17 | ATPase type 13A3 | No | No | - |
| Papss2 | 1,81E-04 | -3,17 | 3'-phosphoadenosine 5'-phosphosulfate synthase 2 | No | No | - |
| Rflnb | 1,56E-06 | -3,17 | refilin B | No | No | - |
| Zfp292 | 1,19E-04 | -3,16 | zinc finger protein 292 | No | No | - |
| Itpkb | 1,71E-02 | -3,16 | inositol 1,4,5-trisphosphate 3-kinase B | No | No | - |
| Hook3 | 8,97E-05 | -3,16 | hook microtubule tethering protein 3 | No | No | - |
| Gm9774 | 1,37E-07 | -3,15 | predicted pseudogene 9774 | No | No | - |
| Ppp1r16a | 3,73E-02 | -3,15 | protein phosphatase 1, regulatory subunit 16A | No | No | - |
| Dlgap4 | 3,65E-08 | -3,15 | DLG associated protein 4 | No | No | - |
| Adamts1 | 1,25E-04 | -3,15 | a disintegrin-like and metallopeptidase (reprolysin type) wit | No | No | 6 |
| Ap5z1 | 1,16E-03 | -3,15 | adaptor-related protein complex 5, zeta 1 subunit | No | No | - |
| Tap1 | 3,72E-03 | -3,15 | transporter 1, ATP-binding cassette, sub-family B (MDR/TA | Yes | No | 5 |
| Per3 | 3,03E-02 | -3,14 | period circadian clock 3 | No | No | - |
| Zfp445 | 7,15E-03 | -3,14 | zinc finger protein 445 | No | No | - |
| Zfyve26 | 2,38E-02 | -3,14 | zinc finger, FYVE domain containing 26 | No | No | - |
| Isyna1 | 8,10E-08 | -3,14 | myo-inositol 1-phosphate synthase A1 | No | No | - |
| Lmna | 1,14E-08 | -3,14 | lamin A | No | No | 10 |
| Ubn1 | 6,30E-09 | -3,14 | ubinuclein 1 | No | No | - |
| Kctd9 | 5,64E-04 | -3,14 | potassium channel tetramerisation domain containing 9 | No | No | - |
| Hsp90aa1 | 1,29E-06 | -3,14 | heat shock protein 90, alpha (cytosolic), class A member 1 | No | No | 1 |
| Atg7 | 8,94E-03 | -3,14 | autophagy related 7 | No | No | 8 |
| Ccn5 | 1,65E-05 | -3,14 | cellular communication network factor 5 | No | No | 8 |
| Limch1 | 6,90E-03 | -3,13 | LIM and calponin homology domains 1 | No | No | - |
| Huwe1 | 4,16E-06 | -3,13 | HECT, UBA and WWE domain containing 1 | No | No | - |
| Numa1 | 1,68E-04 | -3,13 | nuclear mitotic apparatus protein 1 | No | No | - |
| Cdc14b | 6,89E-03 | -3,13 | CDC14 cell division cycle 14B | No | No | - |
| Nuak1 | 3,52E-05 | -3,13 | NUAK family, SNF1-like kinase, 1 | No | No | - |
| Washc4 | 2,38E-03 | -3,13 | WASH complex subunit 4 | No | No | - |
| Bcas3 | 2,70E-02 | -3,13 | BCAS3 microtubule associated cell migration factor | No | No | - |
| Ppp4r3b | 3,11E-03 | -3,13 | protein phosphatase 4 regulatory subunit 3B | No | No | - |
| Lss | 2,91E-05 | -3,12 | lanosterol synthase | No | No | - |
| Tmem150b | 2,24E-02 | -3,12 | transmembrane protein 150B | No | No | - |
| App | 1,95E-05 | -3,12 | amyloid beta (A4) precursor protein | Yes | No | 23 |
| Dnm1l | 1,05E-04 | -3,12 | dynamin 1-like | No | No | - |
| Slc26a2 | 5,98E-05 | -3,11 | solute carrier family 26 (sulfate transporter), member 2 | No | No | - |
| Ly6e | 7,89E-05 | -3,11 | lymphocyte antigen 6 complex, locus E | No | No | - |
| Vipas39 | 2,00E-02 | -3,11 | VPS33B interacting protein, apical-basolateral polarity regu | No | No | - |
| Arhgap10 | 1,39E-02 | -3,10 | Rho GTPase activating protein 10 | No | No | - |
| Chst15 | 1,97E-02 | -3,10 | carbohydrate sulfotransferase 15 | No | No | 9 |
| Slc38a7 | 8,17E-03 | -3,10 | solute carrier family 38, member 7 | No | No | - |
| Axl | 7,03E-09 | -3,10 | AXL receptor tyrosine kinase | No | No | 10 |
| Pomgnt2 | 3,10E-05 | -3,10 | protein O-linked mannose beta 1,4-N-acetylglucosaminyltra | No | No | - |
| Tmed5 | 1,35E-05 | -3,09 | transmembrane p24 trafficking protein 5 | No | No | - |
| Dpp7 | 8,13E-07 | -3,09 | dipeptidylpeptidase 7 | No | No | - |
| Dym | 7,69E-03 | -3,09 | dymeclin | No | No | - |
| Urgcp | 8,69E-03 | -3,09 | upregulator of cell proliferation | No | No | - |
| Sec62 | 1,24E-07 | -3,08 | SEC62 homolog (S. cerevisiae) | No | No | - |
| Ro60 | 1,88E-04 | -3,08 | Ro60, Y RNA binding protein | No | No | - |
| Ktn1 | 5,58E-03 | -3,08 | kinectin 1 | No | No | - |
| Ncln | 1,27E-06 | -3,07 | nicalin | No | No | - |
| Phkb | 3,83E-02 | -3,07 | phosphorylase kinase beta | No | No | - |
| Mpst | 4,22E-02 | -3,07 | mercaptopyruvate sulfurtransferase | No | No | - |
| Hmgcs1 | 8,39E-10 | -3,06 | 3-hydroxy-3-methylglutaryl-Coenzyme A synthase 1 | No | No | - |
| Tbc1d22a | 6,59E-05 | -3,06 | TBC1 domain family, member 22a | No | No | - |
| Gabrb1 | 1,60E-03 | -3,05 | gamma-aminobutyric acid (GABA) A receptor, subunit beta | No | No | - |
| Pggt1b | 1,80E-04 | -3,05 | protein geranylgeranyltransferase type I, beta subunit | No | No | - |
| Usp47 | 1,80E-07 | -3,05 | ubiquitin specific peptidase 47 | No | No | - |
| Il11 | 1,95E-05 | -3,05 | interleukin 11 | No | No | 10 |
| Amz1 | 1,28E-02 | -3,05 | archaelysin family metallopeptidase 1 | No | No | - |
| Mga | 1,45E-02 | -3,04 | MAX gene associated | No | No | 4 |
| Yme1l1 | 2,08E-06 | -3,04 | YME1-like 1 (S. cerevisiae) | No | No | - |
| Exoc7 | 5,06E-03 | -3,03 | exocyst complex component 7 | No | No | - |
| Usp31 | 8,77E-03 | -3,03 | ubiquitin specific peptidase 31 | No | No | - |

| | | | | | | |
|---|---|---|---|---|---|---|
| Zfyve9 | 5,30E-03 | -3,03 | zinc finger, FYVE domain containing 9 | No | No | - |
| Cops2 | 1,05E-04 | -3,03 | COP9 signalosome subunit 2 | No | No | - |
| Sqstm1 | 8,52E-07 | -3,03 | sequestosome 1 | No | No | 3 |
| Plekhg2 | 9,13E-05 | -3,02 | pleckstrin homology domain containing, family G (with Rh | No | No | - |
| Egfl7 | 1,32E-05 | -3,00 | EGF-like domain 7 | No | No | - |
| Gm13610 | 2,11E-02 | -3,00 | predicted gene 13610 | No | No | - |
| Gprc5a | 2,31E-02 | -3,00 | G protein-coupled receptor, family C, group 5, member A | No | No | 9 |
| Itprip | 8,82E-03 | -3,00 | inositol 1,4,5-triphosphate receptor interacting protein | No | No | - |
| Macrod1 | 3,77E-03 | -3,00 | mono-ADP ribosylhydrolase 1 | No | No | - |
| Sult1a1 | 1,89E-02 | -3,00 | sulfotransferase family 1A, phenol-preferring, member 1 | No | No | - |
| Tgfb1 | 1,62E-02 | -3,00 | transforming growth factor, beta 1 | Yes | No | 19 |
| Snx30 | 8,14E-05 | -2,99 | sorting nexin family member 30 | No | No | - |
| Nr1d2 | 1,74E-07 | -2,99 | nuclear receptor subfamily 1, group D, member 2 | No | No | - |
| Gm15590 | 4,69E-03 | -2,99 | predicted gene 15590 | No | No | - |
| Rc3h2 | 1,86E-08 | -2,99 | ring finger and CCCH-type zinc finger domains 2 | No | No | - |
| Sgsh | 5,17E-04 | -2,99 | N-sulfoglucosamine sulfohydrolase (sulfamidase) | No | No | - |
| Bptf | 1,59E-04 | -2,99 | bromodomain PHD finger transcription factor | No | No | - |
| Slc24a3 | 6,41E-03 | -2,97 | solute carrier family 24 (sodium/potassium/calcium exchan | No | No | - |
| Prkx | 4,64E-04 | -2,97 | protein kinase, X-linked | No | No | 1 |
| Ranbp2 | 3,69E-04 | -2,97 | RAN binding protein 2 | No | No | - |
| Spns1 | 9,46E-06 | -2,96 | spinster homolog 1 | No | No | - |
| Smoc2 | 5,21E-05 | -2,96 | SPARC related modular calcium binding 2 | No | No | 6 |
| Arhgef18 | 1,30E-03 | -2,96 | rho/rac guanine nucleotide exchange factor (GEF) 18 | No | No | - |
| Dstyk | 9,22E-03 | -2,96 | dual serine/threonine and tyrosine protein kinase | No | No | - |
| Uggt1 | 2,46E-06 | -2,96 | UDP-glucose glycoprotein glucosyltransferase 1 | No | No | - |
| Nom1 | 6,69E-04 | -2,96 | nucleolar protein with MIF4G domain 1 | No | No | - |
| Odr4 | 2,01E-02 | -2,96 | odr4 GPCR localization factor homolog | No | No | - |
| Extl3 | 1,34E-04 | -2,96 | exostosin-like glycosyltransferase 3 | No | No | - |
| Pknox2 | 1,92E-02 | -2,95 | Pbx/knotted 1 homeobox 2 | No | No | - |
| Pgf | 1,67E-03 | -2,95 | placental growth factor | No | No | 9 |
| Dhcr7 | 2,25E-03 | -2,95 | 7-dehydrocholesterol reductase | No | No | - |
| Igf1r | 1,91E-06 | -2,95 | insulin-like growth factor I receptor | Yes | No | 6 |
| Nnt | 6,56E-04 | -2,94 | nicotinamide nucleotide transhydrogenase | No | No | 10 |
| Plscr4 | 2,19E-02 | -2,94 | phospholipid scramblase 4 | No | No | - |
| Atp9a | 5,15E-04 | -2,94 | ATPase, class II, type 9A | No | No | - |
| Mob3a | 8,51E-03 | -2,94 | MOB kinase activator 3A | No | No | - |
| Prkcsh | 9,33E-09 | -2,94 | protein kinase C substrate 80K-H | No | No | - |
| Fam49a | 6,66E-03 | -2,94 | CYFIP related Rac1 interactor A | No | No | - |
| Dync1li2 | 6,78E-09 | -2,94 | dynein, cytoplasmic 1 light intermediate chain 2 | No | No | - |
| Fpgt | 4,57E-02 | -2,94 | fucose-1-phosphate guanylyltransferase | No | No | - |
| Tmem131 | 3,04E-02 | -2,94 | transmembrane protein 131 | No | No | - |
| Tmem176b | 9,24E-05 | -2,93 | transmembrane protein 176B | No | No | - |
| Kctd11 | 1,56E-05 | -2,93 | potassium channel tetramerisation domain containing 11 | No | No | - |
| Prickle2 | 5,55E-03 | -2,93 | prickle planar cell polarity protein 2 | No | No | - |
| Ssbp4 | 7,01E-07 | -2,93 | single stranded DNA binding protein 4 | No | No | - |
| Nox4 | 1,61E-05 | -2,93 | NADPH oxidase 4 | No | No | 151 |
| Fitm2 | 5,14E-03 | -2,93 | fat storage-inducing transmembrane protein 2 | No | No | - |
| Plxna4 | 2,27E-02 | -2,93 | plexin A4 | No | No | 5 |
| Atp6v0a1 | 2,49E-04 | -2,93 | ATPase, H+ transporting, lysosomal V0 subunit A1 | No | No | - |
| Pik3r1 | 2,22E-03 | -2,93 | phosphoinositide-3-kinase regulatory subunit 1 | No | No | 2 |
| Ccdc82 | 8,26E-03 | -2,93 | coiled-coil domain containing 82 | No | No | - |
| Klf4 | 9,13E-04 | -2,92 | Kruppel-like factor 4 (gut) | Yes | Yes | 11 |
| Lgals9 | 9,87E-04 | -2,92 | lectin, galactose binding, soluble 9 | No | No | - |
| Uap1l1 | 2,79E-05 | -2,92 | UDP-N-acteylglucosamine pyrophosphorylase 1-like 1 | No | No | - |
| Sf3a2 | 1,70E-04 | -2,92 | splicing factor 3a, subunit 2 | No | No | - |
| Exosc5 | 4,76E-04 | -2,92 | exosome component 5 | No | No | - |
| Fmnl2 | 8,96E-03 | -2,92 | formin-like 2 | No | No | - |
| Gdf6 | 1,20E-03 | -2,92 | growth differentiation factor 6 | No | No | - |
| Glg1 | 1,43E-04 | -2,92 | golgi apparatus protein 1 | No | No | - |
| Peg13 | 1,19E-02 | -2,92 | paternally expressed 13 | No | No | - |
| Mbnl1 | 1,94E-08 | -2,91 | muscleblind like splicing factor 1 | No | No | - |
| Kmt2a | 1,69E-03 | -2,91 | lysine (K)-specific methyltransferase 2A | No | No | - |
| Ogdhl | 1,38E-02 | -2,91 | oxoglutarate dehydrogenase-like | No | No | - |
| Foxo3 | 4,22E-05 | -2,90 | forkhead box O3 | No | Yes | 5 |
| Ar | 2,22E-03 | -2,90 | androgen receptor | Yes | Yes | 102 |
| C1qa | 4,14E-02 | -2,90 | complement component 1, q subcomponent, alpha polypept | No | No | - |
| Gm6548 | 2,93E-03 | -2,90 | predicted gene 6548 | No | No | - |
| Igip | 3,60E-02 | -2,90 | IgA inducing protein | No | No | - |
| Zfp869 | 8,30E-03 | -2,90 | zinc finger protein 869 | No | No | - |
| Furin | 7,00E-07 | -2,90 | furin (paired basic amino acid cleaving enzyme) | No | No | 2 |
| Bmp1 | 1,15E-07 | -2,90 | bone morphogenetic protein 1 | No | No | - |
| Pcnx | 1,42E-02 | -2,90 | pecanex homolog | No | No | - |
| Agrn | 5,17E-03 | -2,89 | agrin | No | No | - |

| | | | | | | |
|---|---|---|---|---|---|---|
| Tnks2 | 2,40E-07 | -2,89 | tankyrase, TRF1-interacting ankyrin-related ADP-ribose po | No | No | 6 |
| Trim37 | 3,29E-03 | -2,89 | tripartite motif-containing 37 | No | No | - |
| Capg | 7,81E-05 | -2,89 | capping protein (actin filament), gelsolin-like | No | No | - |
| Nkain4 | 4,10E-02 | -2,89 | Na+/K+ transporting ATPase interacting 4 | No | No | - |
| Zmym4 | 5,92E-03 | -2,89 | zinc finger, MYM-type 4 | No | No | - |
| Prkg1 | 3,09E-03 | -2,89 | protein kinase, cGMP-dependent, type I | No | No | - |
| Akap13 | 8,35E-04 | -2,88 | A kinase (PRKA) anchor protein 13 | No | No | 5 |
| Myo7a | 3,81E-04 | -2,88 | myosin VIIA | No | No | - |
| Lrrc8c | 4,69E-05 | -2,88 | leucine rich repeat containing 8 family, member C | No | No | 2 |
| Arhgef19 | 3,39E-02 | -2,88 | Rho guanine nucleotide exchange factor (GEF) 19 | No | No | - |
| Gm15866 | 2,51E-03 | -2,88 | predicted gene 15866 | No | No | - |
| Man2b2 | 3,35E-02 | -2,88 | mannosidase 2, alpha B2 | No | No | - |
| Tmed8 | 4,34E-02 | -2,88 | transmembrane p24 trafficking protein 8 | No | No | - |
| Utp20 | 8,08E-03 | -2,88 | UTP20 small subunit processome component | No | No | - |
| Hsdl1 | 2,61E-02 | -2,87 | hydroxysteroid dehydrogenase like 1 | No | No | - |
| Wdr81 | 2,25E-03 | -2,87 | WD repeat domain 81 | No | No | - |
| Ctsb | 6,10E-07 | -2,87 | cathepsin B | Yes | No | 1 |
| Ppfibp2 | 4,47E-02 | -2,87 | PTPRF interacting protein, binding protein 2 (liprin beta 2) | No | No | - |
| Sacm1l | 3,99E-04 | -2,86 | SAC1 suppressor of actin mutations 1-like (yeast) | No | No | - |
| Pofut1 | 1,05E-04 | -2,86 | protein O-fucosyltransferase 1 | No | No | - |
| Slc38a10 | 9,76E-07 | -2,86 | solute carrier family 38, member 10 | No | No | - |
| Canx | 3,84E-08 | -2,86 | calnexin | No | No | 1 |
| Ccn4 | 1,78E-04 | -2,86 | cellular communication network factor 4 | No | No | - |
| Hspa4l | 2,66E-04 | -2,86 | heat shock protein 4 like | No | No | - |
| Syt13 | 2,55E-02 | -2,86 | synaptotagmin XIII | No | No | - |
| Thoc2 | 7,33E-04 | -2,85 | THO complex 2 | No | No | - |
| Ppl | 3,11E-07 | -2,85 | periplakin | No | No | 13 |
| Pls3 | 2,17E-05 | -2,85 | plastin 3 (T-isoform) | No | No | - |
| Nnt | 8,26E-04 | -2,85 | nicotinamide nucleotide transhydrogenase | No | No | 10 |
| Slc7a5 | 2,07E-05 | -2,85 | solute carrier family 7 (cationic amino acid transporter, y+ s | No | No | - |
| Col14a1 | 1,18E-02 | -2,85 | collagen, type XIV, alpha 1 | No | No | 1 |
| Hmgxb3 | 1,76E-03 | -2,85 | HMG box domain containing 3 | No | No | - |
| Mtmr10 | 2,61E-03 | -2,85 | myotubularin related protein 10 | No | No | - |
| Map2k5 | 8,44E-03 | -2,85 | mitogen-activated protein kinase kinase 5 | No | No | - |
| Tcaf2 | 9,27E-04 | -2,85 | TRPM8 channel-associated factor 2 | No | No | - |
| Strn3 | 1,73E-04 | -2,85 | striatin, calmodulin binding protein 3 | No | No | - |
| Ptk2b | 6,01E-05 | -2,84 | PTK2 protein tyrosine kinase 2 beta | No | No | - |
| Slc9a3r2 | 2,30E-03 | -2,84 | solute carrier family 9 (sodium/hydrogen exchanger), memb | No | No | - |
| Ccdc25 | 3,07E-04 | -2,84 | coiled-coil domain containing 25 | No | No | - |
| Rb1cc1 | 2,70E-03 | -2,84 | RB1-inducible coiled-coil 1 | No | No | - |
| Fgfr1op | 5,58E-03 | -2,83 | centrosomal protein 43 | No | No | - |
| Spen | 5,95E-04 | -2,83 | spen family transcription repressor | No | No | - |
| Stim1 | 4,49E-05 | -2,83 | stromal interaction molecule 1 | No | No | - |
| Syngr1 | 2,87E-02 | -2,83 | synaptogyrin 1 | No | No | - |
| Foxo1 | 5,90E-03 | -2,83 | forkhead box O1 | Yes | Yes | 1 |
| Nfkbil1 | 4,29E-02 | -2,82 | nuclear factor of kappa light polypeptide gene enhancer in E | No | No | - |
| Psd | 1,77E-03 | -2,82 | pleckstrin and Sec7 domain containing | No | No | 2 |
| Sardh | 4,64E-03 | -2,82 | sarcosine dehydrogenase | No | No | - |
| Col12a1 | 1,78E-05 | -2,82 | collagen, type XII, alpha 1 | No | No | - |
| Dicer1 | 9,21E-03 | -2,82 | dicer 1, ribonuclease type III | No | No | - |
| Atrx | 4,74E-04 | -2,82 | ATRX, chromatin remodeler | No | No | - |
| Fastkd5 | 3,02E-02 | -2,82 | FAST kinase domains 5 | No | No | - |
| Foxp4 | 3,07E-02 | -2,82 | forkhead box P4 | No | No | 2 |
| Bicd2 | 6,92E-05 | -2,82 | BICD cargo adaptor 2 | Yes | No | 2 |
| Slu7 | 1,87E-03 | -2,81 | SLU7 splicing factor homolog (S. cerevisiae) | No | No | - |
| Aldh16a1 | 2,16E-02 | -2,81 | aldehyde dehydrogenase 16 family, member A1 | No | No | - |
| Mmp23 | 1,35E-03 | -2,80 | matrix metallopeptidase 23 | No | No | - |
| Ltbr | 5,78E-06 | -2,80 | lymphotoxin B receptor | No | No | 2 |
| Tns1 | 4,26E-07 | -2,80 | tensin 1 | No | No | 10 |
| Flad1 | 2,59E-02 | -2,80 | flavin adenine dinucleotide synthetase 1 | No | No | - |
| Mtus1 | 6,95E-05 | -2,80 | mitochondrial tumor suppressor 1 | No | No | - |
| Denn2b | 9,10E-07 | -2,80 | DENN domain containing 2B | No | No | - |
| Apoe | 1,67E-03 | -2,80 | apolipoprotein E | Yes | No | 13 |
| Maml2 | 5,09E-04 | -2,79 | mastermind like transcriptional coactivator 2 | No | No | - |
| Gnaq | 6,29E-05 | -2,79 | guanine nucleotide binding protein, alpha q polypeptide | No | No | - |
| Aftph | 1,93E-03 | -2,79 | aftiphilin | No | No | - |
| Itpr1 | 4,84E-03 | -2,79 | inositol 1,4,5-trisphosphate receptor 1 | No | No | - |
| Suco | 4,76E-05 | -2,79 | SUN domain containing ossification factor | No | No | - |
| Med16 | 5,52E-03 | -2,79 | mediator complex subunit 16 | Yes | No | - |
| Pear1 | 7,04E-03 | -2,79 | platelet endothelial aggregation receptor 1 | No | No | - |
| Zdhhc18 | 3,43E-03 | -2,79 | zinc finger, DHHC domain containing 18 | No | No | - |
| Fam234a | 3,05E-02 | -2,78 | family with sequence similarity 234, member A | No | No | - |
| Zbtb43 | 3,45E-02 | -2,78 | zinc finger and BTB domain containing 43 | No | No | - |

| | | | | | | |
|---|---|---|---|---|---|---|
| Zc3hav1 | 9,67E-05 | -2,78 | zinc finger CCCH type, antiviral 1 | Yes | No | - |
| Fnip2 | 9,04E-03 | -2,77 | folliculin interacting protein 2 | No | No | - |
| Znhit6 | 4,42E-04 | -2,77 | zinc finger, HIT type 6 | No | No | - |
| Zfp871 | 1,61E-02 | -2,77 | zinc finger protein 871 | No | No | - |
| Exoc4 | 2,97E-04 | -2,77 | exocyst complex component 4 | No | No | - |
| Cep104 | 4,33E-03 | -2,77 | centrosomal protein 104 | No | No | - |
| Mfn1 | 8,71E-04 | -2,77 | mitofusin 1 | No | No | 4 |
| Disp1 | 2,44E-03 | -2,76 | dispatched RND transporter family member 1 | No | No | - |
| Gm7901 | 2,72E-02 | -2,76 | predicted gene 7901 | No | No | - |
| Arhgap21 | 2,05E-03 | -2,76 | Rho GTPase activating protein 21 | No | No | - |
| Gigyf2 | 2,85E-02 | -2,75 | GRB10 interacting GYF protein 2 | No | No | - |
| Lmf2 | 3,55E-03 | -2,75 | lipase maturation factor 2 | No | No | - |
| Prkca | 4,86E-03 | -2,75 | protein kinase C, alpha | No | No | - |
| Traf6 | 4,95E-02 | -2,75 | TNF receptor-associated factor 6 | No | No | 3 |
| Ncam1 | 5,46E-06 | -2,75 | neural cell adhesion molecule 1 | No | No | - |
| Ttc3 | 2,03E-03 | -2,74 | tetratricopeptide repeat domain 3 | No | No | - |
| Gpr39 | 3,69E-04 | -2,74 | G protein-coupled receptor 39 | No | No | - |
| Cc2d1b | 1,93E-03 | -2,74 | coiled-coil and C2 domain containing 1B | No | No | - |
| Cx3cl1 | 2,22E-02 | -2,74 | chemokine (C-X3-C motif) ligand 1 | Yes | No | 43 |
| Nucb2 | 1,14E-04 | -2,74 | nucleobindin 2 | No | No | - |
| Acin1 | 2,61E-03 | -2,74 | apoptotic chromatin condensation inducer 1 | No | No | - |
| Gcc2 | 3,85E-02 | -2,74 | GRIP and coiled-coil domain containing 2 | No | No | - |
| Gm21691 | 3,97E-04 | -2,73 | predicted gene, 21691 | No | No | - |
| Cebpa | 5,17E-03 | -2,73 | CCAAT/enhancer binding protein (C/EBP), alpha | No | Yes | 14 |
| Xrn1 | 1,60E-02 | -2,73 | 5'-3' exoribonuclease 1 | No | No | - |
| Unc45a | 2,18E-04 | -2,73 | unc-45 myosin chaperone A | No | No | - |
| Kif1a | 8,57E-05 | -2,73 | kinesin family member 1A | No | No | - |
| Ddx3x | 2,97E-06 | -2,73 | DEAD box helicase 3, X-linked | No | No | - |
| Eng | 2,70E-04 | -2,72 | endoglin | No | No | 1 |
| Nsdhl | 3,48E-07 | -2,72 | NAD(P) dependent steroid dehydrogenase-like | No | No | - |
| Adamts12 | 1,07E-03 | -2,72 | a disintegrin-like and metallopeptidase (reprolysin type) wit | No | No | - |
| Ehd3 | 3,54E-04 | -2,72 | EH-domain containing 3 | No | No | - |
| Nfkb2 | 1,15E-03 | -2,72 | nuclear factor of kappa light polypeptide gene enhancer in E | No | Yes | - |
| Evc | 3,73E-02 | -2,71 | EvC ciliary complex subunit 1 | No | No | - |
| Fah | 1,30E-02 | -2,71 | fumarylacetoacetate hydrolase | No | No | 6 |
| Tmem63a | 4,53E-07 | -2,71 | transmembrane protein 63a | No | No | - |
| Vps50 | 4,46E-04 | -2,71 | VPS50 EARP/GARPII complex subunit | No | No | - |
| Zfpm1 | 1,28E-04 | -2,71 | zinc finger protein, multitype 1 | No | No | - |
| Gclc | 2,12E-02 | -2,71 | glutamate-cysteine ligase, catalytic subunit | No | No | 6 |
| Enc1 | 2,54E-04 | -2,71 | ectodermal-neural cortex 1 | No | No | - |
| Fbln2 | 4,12E-04 | -2,70 | fibulin 2 | No | No | - |
| Cdkn1a | 1,92E-05 | -2,70 | cyclin-dependent kinase inhibitor 1A (P21) | Yes | No | 5 |
| Slc39a7 | 1,40E-07 | -2,70 | solute carrier family 39 (zinc transporter), member 7 | No | No | - |
| Cyb5b | 3,30E-06 | -2,70 | cytochrome b5 type B | No | No | - |
| Ap2a1 | 5,35E-04 | -2,69 | adaptor-related protein complex 2, alpha 1 subunit | No | No | 1 |
| Ip6k1 | 6,31E-05 | -2,69 | inositol hexaphosphate kinase 1 | No | No | - |
| Creb3l2 | 4,90E-07 | -2,69 | cAMP responsive element binding protein 3-like 2 | No | No | - |
| Gm7072 | 4,47E-02 | -2,69 | predicted gene 7072 | No | No | - |
| Il18bp | 1,31E-03 | -2,69 | interleukin 18 binding protein | Yes | No | - |
| Loxl1 | 4,59E-05 | -2,68 | lysyl oxidase-like 1 | No | No | 13 |
| Tssc4 | 5,87E-05 | -2,68 | tumor-suppressing subchromosomal transferable fragment 4 | No | No | - |
| Kazn | 4,28E-03 | -2,68 | kazrin, periplakin interacting protein | No | No | - |
| Slc7a2 | 8,02E-03 | -2,68 | solute carrier family 7 (cationic amino acid transporter, y+ s | No | No | 1 |
| Sun2 | 4,77E-04 | -2,68 | Sad1 and UNC84 domain containing 2 | No | No | - |
| Nhs | 1,75E-02 | -2,67 | NHS actin remodeling regulator | No | No | 26 |
| Nifk | 1,42E-03 | -2,67 | nucleolar protein interacting with the FHA domain of MKI6 | No | No | - |
| Notch3 | 4,38E-02 | -2,67 | notch 3 | No | No | 7 |
| Sobp | 2,26E-02 | -2,67 | sine oculis binding protein | No | No | - |
| Srgap1 | 2,59E-02 | -2,67 | SLIT-ROBO Rho GTPase activating protein 1 | No | No | - |
| Tbcd | 4,47E-03 | -2,67 | tubulin-specific chaperone d | No | No | - |
| Tpp1 | 3,60E-07 | -2,66 | tripeptidyl peptidase I | No | No | 8 |
| Lrrc27 | 8,44E-03 | -2,66 | leucine rich repeat containing 27 | No | No | - |
| Parp9 | 1,71E-02 | -2,66 | poly (ADP-ribose) polymerase family, member 9 | No | No | - |
| Rasa1 | 2,98E-03 | -2,65 | RAS p21 protein activator 1 | No | No | - |
| Herc2 | 5,27E-03 | -2,65 | HECT and RLD domain containing E3 ubiquitin protein lig | No | No | - |
| Ereg | 6,59E-04 | -2,65 | epiregulin | No | No | - |
| Lrp1 | 4,91E-05 | -2,64 | low density lipoprotein receptor-related protein 1 | No | No | 1 |
| Stat1 | 2,24E-02 | -2,64 | signal transducer and activator of transcription 1 | No | Yes | 112 |
| Tmem104 | 2,12E-02 | -2,64 | transmembrane protein 104 | No | No | - |
| Slc15a3 | 3,17E-02 | -2,64 | solute carrier family 15, member 3 | No | No | - |
| Sos2 | 2,51E-02 | -2,64 | SOS Ras/Rho guanine nucleotide exchange factor 2 | No | No | - |
| Hexa | 6,28E-06 | -2,64 | hexosaminidase A | No | No | - |
| Wwc2 | 8,01E-05 | -2,64 | WW, C2 and coiled-coil domain containing 2 | No | No | - |

| | | | | | | |
|---|---|---|---|---|---|---|
| Ctsz | 4,00E-07 | -2,63 | cathepsin Z | No | No | - |
| Auts2 | 3,71E-02 | -2,63 | autism susceptibility candidate 2 | No | No | - |
| Cln6 | 1,44E-02 | -2,63 | ceroid-lipofuscinosis, neuronal 6 | No | No | - |
| Dusp4 | 5,00E-02 | -2,63 | dual specificity phosphatase 4 | No | No | - |
| Sipa1l2 | 2,92E-03 | -2,63 | signal-induced proliferation-associated 1 like 2 | No | No | - |
| Stub1 | 2,47E-05 | -2,62 | STIP1 homology and U-Box containing protein 1 | No | No | 1 |
| Tln1 | 5,76E-05 | -2,62 | talin 1 | No | No | - |
| B3gntl1 | 3,14E-03 | -2,62 | UDP-GlcNAc:betaGal beta-1,3-N-acetylglucosaminyltransf | No | No | - |
| Zhx3 | 1,37E-04 | -2,62 | zinc fingers and homeoboxes 3 | No | No | - |
| Sparc | 1,57E-05 | -2,61 | secreted acidic cysteine rich glycoprotein | No | No | 93 |
| Aspn | 2,61E-02 | -2,61 | asporin | No | No | 15 |
| Stxbp5 | 3,37E-02 | -2,61 | syntaxin binding protein 5 (tomosyn) | No | No | - |
| Ankle2 | 2,27E-02 | -2,61 | ankyrin repeat and LEM domain containing 2 | No | No | - |
| Timp2 | 3,35E-05 | -2,61 | tissue inhibitor of metalloproteinase 2 | Yes | No | 4 |
| Smc5 | 7,07E-03 | -2,61 | structural maintenance of chromosomes 5 | No | No | - |
| Fbln5 | 3,48E-04 | -2,60 | fibulin 5 | No | No | - |
| Ctr9 | 3,05E-03 | -2,60 | CTR9 homolog, Paf1/RNA polymerase II complex compon | No | No | - |
| Tmcc3 | 5,47E-03 | -2,60 | transmembrane and coiled coil domains 3 | No | No | - |
| Lrsam1 | 5,87E-07 | -2,60 | leucine rich repeat and sterile alpha motif containing 1 | No | No | - |
| Arhgef5 | 3,09E-05 | -2,60 | Rho guanine nucleotide exchange factor (GEF) 5 | No | No | - |
| Btbd19 | 1,34E-03 | -2,60 | BTB (POZ) domain containing 19 | No | No | - |
| Pygb | 3,00E-04 | -2,60 | brain glycogen phosphorylase | No | No | - |
| Tnpo2 | 1,04E-06 | -2,60 | transportin 2 (importin 3, karyopherin beta 2b) | No | No | - |
| Sqle | 9,81E-05 | -2,59 | squalene epoxidase | Yes | No | - |
| Nedd9 | 1,90E-04 | -2,59 | neural precursor cell expressed, developmentally down-regu | No | No | 2 |
| Dgkq | 1,33E-03 | -2,58 | diacylglycerol kinase, theta | No | No | - |
| Ptger4 | 3,18E-02 | -2,58 | prostaglandin E receptor 4 (subtype EP4) | No | No | 2 |
| Zfp316 | 1,42E-02 | -2,58 | zinc finger protein 316 | No | No | - |
| Smurf1 | 2,94E-04 | -2,58 | SMAD specific E3 ubiquitin protein ligase 1 | No | No | - |
| Oip5os1 | 2,26E-03 | -2,58 | Opa interacting protein 5, opposite strand 1 | No | No | - |
| Fam129b | 6,76E-07 | -2,58 | niban apoptosis regulator 2 | No | No | - |
| Bcr | 1,14E-02 | -2,58 | BCR activator of RhoGEF and GTPase | No | No | 15 |
| Itgb5 | 1,92E-06 | -2,57 | integrin beta 5 | No | No | - |
| Ccdc186 | 2,67E-03 | -2,57 | coiled-coil domain containing 186 | No | No | - |
| Ecpas | 1,67E-04 | -2,57 | Ecm29 proteasome adaptor and scaffold | No | No | - |
| Amot | 1,34E-03 | -2,57 | angiomotin | No | No | - |
| Sema3d | 4,34E-03 | -2,57 | sema domain, immunoglobulin domain (Ig), short basic don | No | No | - |
| Nelfcd | 6,95E-06 | -2,57 | negative elongation factor complex member C/D, Th1l | No | No | - |
| Trpm5 | 8,55E-05 | -2,56 | transient receptor potential cation channel, subfamily M, me | No | No | - |
| Ndst1 | 1,56E-06 | -2,56 | N-deacetylase/N-sulfotransferase (heparan glucosaminyl) 1 | No | No | - |
| Smg9 | 2,71E-02 | -2,56 | SMG9 nonsense mediated mRNA decay factor | No | No | - |
| Map1lc3b | 4,48E-07 | -2,55 | microtubule-associated protein 1 light chain 3 beta | No | No | - |
| Acads | 3,30E-04 | -2,55 | acyl-Coenzyme A dehydrogenase, short chain | No | No | - |
| Lins1 | 1,71E-02 | -2,55 | lines homolog 1 | No | No | - |
| Plxna1 | 3,49E-04 | -2,55 | plexin A1 | No | No | - |
| Smc1a | 1,14E-03 | -2,55 | structural maintenance of chromosomes 1A | No | No | - |
| Brwd3 | 1,48E-03 | -2,55 | bromodomain and WD repeat domain containing 3 | No | No | - |
| Rrn3 | 2,24E-04 | -2,55 | RRN3 RNA polymerase I transcription factor homolog (yea | No | No | - |
| Ephx1 | 2,59E-04 | -2,55 | epoxide hydrolase 1, microsomal | No | No | 5 |
| Fbxo31 | 1,11E-02 | -2,55 | F-box protein 31 | No | No | - |
| Klhl18 | 3,21E-02 | -2,55 | kelch-like 18 | No | No | - |
| Zfp579 | 2,98E-02 | -2,55 | zinc finger protein 579 | No | No | - |
| Myh11 | 2,98E-03 | -2,54 | myosin, heavy polypeptide 11, smooth muscle | No | No | 1 |
| Gpat4 | 4,89E-05 | -2,54 | glycerol-3-phosphate acyltransferase 4 | No | No | - |
| Ccdc174 | 6,08E-03 | -2,54 | coiled-coil domain containing 174 | No | No | - |
| Faf1 | 1,16E-02 | -2,54 | Fas-associated factor 1 | No | No | - |
| Irs2 | 1,60E-03 | -2,54 | insulin receptor substrate 2 | No | No | - |
| Tmem175 | 9,18E-04 | -2,54 | transmembrane protein 175 | No | No | - |
| AU020206 | 3,67E-02 | -2,54 | expressed sequence AU020206 | No | No | - |
| Vav2 | 2,56E-03 | -2,54 | vav 2 oncogene | No | No | - |
| Arrb2 | 2,24E-02 | -2,54 | arrestin, beta 2 | No | No | - |
| Plekho2 | 1,36E-04 | -2,54 | pleckstrin homology domain containing, family O member 2 | No | No | - |
| Ctdspl | 1,12E-04 | -2,54 | CTD (carboxy-terminal domain, RNA polymerase II, polyp | No | No | - |
| Pdgfd | 3,06E-02 | -2,53 | platelet-derived growth factor, D polypeptide | No | No | - |
| Appl1 | 9,36E-05 | -2,53 | adaptor protein, phosphotyrosine interaction, PH domain an | No | No | - |
| Aup1 | 4,25E-06 | -2,52 | ancient ubiquitous protein 1 | No | No | - |
| Cep120 | 1,15E-02 | -2,52 | centrosomal protein 120 | No | No | - |
| Gm18584 | 3,23E-02 | -2,52 | predicted gene, 18584 | No | No | - |
| Heatr1 | 1,03E-03 | -2,52 | HEAT repeat containing 1 | No | No | - |
| Csf1 | 6,04E-06 | -2,52 | colony stimulating factor 1 (macrophage) | Yes | No | 5 |
| Fuz | 8,93E-04 | -2,52 | fuzzy planar cell polarity protein | No | No | - |
| Slc7a1 | 7,06E-05 | -2,52 | solute carrier family 7 (cationic amino acid transporter, y+ s | No | No | 1 |
| Skil | 1,65E-04 | -2,52 | SKI-like | No | No | - |

| | | | | | | |
|---|---|---|---|---|---|---|
| Hdac4 | 4,71E-03 | -2,52 | histone deacetylase 4 | No | No | 32 |
| Nlrx1 | 1,48E-02 | -2,52 | NLR family member X1 | No | No | - |
| Pitpnm2 | 2,22E-02 | -2,51 | phosphatidylinositol transfer protein, membrane-associated | No | No | - |
| Sel1l | 2,18E-04 | -2,51 | sel-1 suppressor of lin-12-like (C. elegans) | No | No | - |
| Mgat5 | 1,00E-02 | -2,51 | mannoside acetylglucosaminyltransferase 5 | No | No | - |
| Myh9 | 1,79E-06 | -2,51 | myosin, heavy polypeptide 9, non-muscle | Yes | No | 9 |
| Thbs2 | 6,24E-05 | -2,51 | thrombospondin 2 | No | No | 1 |
| Jup | 4,01E-03 | -2,51 | junction plakoglobin | No | No | - |
| Xpo7 | 1,31E-03 | -2,51 | exportin 7 | No | No | - |
| Gabarapl1 | 2,80E-07 | -2,50 | gamma-aminobutyric acid (GABA) A receptor-associated p | No | No | - |
| Cd40 | 2,46E-02 | -2,50 | CD40 antigen | Yes | No | 80 |
| Cln3 | 3,56E-02 | -2,50 | ceroid lipofuscinosis, neuronal 3, juvenile (Batten, Spielme | No | No | - |
| Enpp4 | 2,18E-02 | -2,50 | ectonucleotide pyrophosphatase/phosphodiesterase 4 | No | No | - |
| Fam3a | 3,39E-02 | -2,50 | FAM3 metabolism regulating signaling molecule A | No | No | - |
| Ptprn | 2,95E-03 | -2,50 | protein tyrosine phosphatase, receptor type, N | No | No | - |
| Tmem132a | 3,96E-02 | -2,50 | transmembrane protein 132A | No | No | - |
| Fus | 4,01E-07 | -2,50 | fused in sarcoma | No | No | - |
| Csde1 | 2,93E-07 | -2,50 | cold shock domain containing E1, RNA binding | No | No | - |
| Sec61a1 | 3,69E-07 | -2,49 | Sec61 alpha 1 subunit (S. cerevisiae) | No | No | - |
| Abl2 | 7,80E-03 | -2,49 | v-abl Abelson murine leukemia viral oncogene 2 (arg, Abel | No | No | - |
| Septin11 | 7,33E-07 | -2,49 | septin 11 | No | No | - |
| Btbd7 | 2,40E-02 | -2,49 | BTB (POZ) domain containing 7 | No | No | - |
| Des | 1,71E-02 | -2,49 | desmin | No | No | 4 |
| Tasor | 3,33E-02 | -2,48 | transcription activation suppressor | No | No | - |
| Trappc12 | 3,88E-02 | -2,48 | trafficking protein particle complex 12 | No | No | - |
| Farsa | 1,10E-06 | -2,48 | phenylalanyl-tRNA synthetase, alpha subunit | No | No | 6 |
| Olfm1 | 1,90E-03 | -2,48 | olfactomedin 1 | No | No | - |
| Tmem245 | 1,28E-02 | -2,48 | transmembrane protein 245 | No | No | - |
| Btbd9 | 4,58E-04 | -2,47 | BTB (POZ) domain containing 9 | No | No | - |
| Kansl3 | 7,70E-04 | -2,47 | KAT8 regulatory NSL complex subunit 3 | No | No | - |
| Cpe | 4,53E-07 | -2,47 | carboxypeptidase E | No | No | 18 |
| Gm9780 | 1,08E-02 | -2,47 | - | No | No | - |
| Jmjd8 | 3,01E-05 | -2,47 | jumonji domain containing 8 | No | No | - |
| Fgf9 | 2,11E-02 | -2,47 | fibroblast growth factor 9 | No | No | 27 |
| Zzef1 | 6,92E-03 | -2,47 | zinc finger, ZZ-type with EF hand domain 1 | No | No | - |
| Smarcd3 | 8,78E-04 | -2,47 | SWI/SNF related, matrix associated, actin dependent regula | No | No | - |
| Atp6v0d1 | 1,49E-06 | -2,47 | ATPase, H+ transporting, lysosomal V0 subunit D1 | No | No | - |
| Opa3 | 9,45E-05 | -2,47 | optic atrophy 3 | No | No | - |
| Golga4 | 2,28E-02 | -2,47 | golgi autoantigen, golgin subfamily a, 4 | No | No | - |
| Sec63 | 2,26E-06 | -2,47 | SEC63-like (S. cerevisiae) | No | No | - |
| Gm4754 | 3,47E-03 | -2,46 | predicted gene 4754 | No | No | - |
| Plbd2 | 4,89E-04 | -2,46 | phospholipase B domain containing 2 | No | No | - |
| Ern1 | 3,70E-02 | -2,46 | endoplasmic reticulum (ER) to nucleus signalling 1 | No | No | - |
| H3c15 | 6,96E-03 | -2,46 | H3 clustered histone 15 | No | No | - |
| Mfsd6 | 4,11E-02 | -2,46 | major facilitator superfamily domain containing 6 | No | No | - |
| Nav1 | 1,59E-03 | -2,46 | neuron navigator 1 | No | No | - |
| Zfyve16 | 4,85E-02 | -2,45 | zinc finger, FYVE domain containing 16 | No | No | - |
| Flna | 2,71E-06 | -2,45 | filamin, alpha | No | No | 30 |
| Pik3r4 | 8,82E-03 | -2,45 | phosphoinositide-3-kinase regulatory subunit 4 | No | No | - |
| Nacc1 | 4,49E-04 | -2,45 | nucleus accumbens associated 1, BEN and BTB (POZ) dom | No | No | - |
| C2cd2 | 1,98E-02 | -2,45 | C2 calcium-dependent domain containing 2 | No | No | - |
| Agtrap | 4,92E-05 | -2,45 | angiotensin II, type I receptor-associated protein | No | No | - |
| Il4i1 | 2,12E-02 | -2,45 | interleukin 4 induced 1 | No | No | - |
| Pml | 1,23E-03 | -2,45 | promyelocytic leukemia | Yes | No | 34 |
| Gsk3b | 5,54E-04 | -2,45 | glycogen synthase kinase 3 beta | Yes | No | 3 |
| Kdm5a | 7,01E-03 | -2,45 | lysine (K)-specific demethylase 5A | No | No | - |
| Rrbp1 | 7,46E-06 | -2,45 | ribosome binding protein 1 | No | No | - |
| Fgf10 | 5,50E-03 | -2,44 | fibroblast growth factor 10 | No | No | 15 |
| Cyld | 4,83E-06 | -2,44 | CYLD lysine 63 deubiquitinase | No | No | - |
| Tapbpl | 4,54E-03 | -2,44 | TAP binding protein-like | No | No | - |
| Rab40c | 1,21E-02 | -2,44 | Rab40C, member RAS oncogene family | No | No | - |
| Csrp1 | 6,86E-05 | -2,44 | cysteine and glycine-rich protein 1 | Yes | No | - |
| Ggnbp2 | 1,91E-04 | -2,44 | gametogenetin binding protein 2 | No | No | - |
| Acaca | 5,14E-03 | -2,44 | acetyl-Coenzyme A carboxylase alpha | Yes | No | - |
| Ptpn14 | 2,15E-03 | -2,43 | protein tyrosine phosphatase, non-receptor type 14 | No | No | - |
| Acat3 | 8,05E-06 | -2,43 | acetyl-Coenzyme A acetyltransferase 3 | No | No | - |
| Stk38 | 2,94E-04 | -2,43 | serine/threonine kinase 38 | No | No | - |
| Slc44a2 | 2,96E-05 | -2,43 | solute carrier family 44, member 2 | No | No | - |
| Snx11 | 6,96E-05 | -2,43 | sorting nexin 11 | No | No | - |
| Zpr1 | 1,14E-04 | -2,43 | ZPR1 zinc finger | No | No | - |
| Bcl2l1 | 8,76E-05 | -2,43 | BCL2-like 1 | No | No | 3 |
| Phlda3 | 6,23E-05 | -2,43 | pleckstrin homology like domain, family A, member 3 | No | No | - |
| Vti1a | 2,21E-02 | -2,43 | vesicle transport through interaction with t-SNAREs 1A | No | No | - |

| | | | | | | |
|---|---|---|---|---|---|---|
| Ltbp4 | 5,92E-03 | -2,43 | latent transforming growth factor beta binding protein 4 | No | No | 2 |
| Masp1 | 1,28E-03 | -2,43 | mannan-binding lectin serine peptidase 1 | No | No | - |
| Chpf2 | 7,06E-04 | -2,42 | chondroitin polymerizing factor 2 | No | No | - |
| Sesn2 | 1,39E-02 | -2,42 | sestrin 2 | No | No | - |
| Tep1 | 2,06E-02 | -2,42 | telomerase associated protein 1 | No | No | - |
| Kif1b | 3,16E-03 | -2,42 | kinesin family member 1B | No | No | - |
| Ccn2 | 2,09E-06 | -2,41 | cellular communication network factor 2 | No | No | 55 |
| Ide | 1,02E-02 | -2,41 | insulin degrading enzyme | No | No | - |
| Ubn2 | 5,93E-03 | -2,41 | ubinuclein 2 | No | No | - |
| Acat2 | 8,39E-05 | -2,41 | acetyl-Coenzyme A acetyltransferase 2 | No | No | - |
| Tsc1 | 4,72E-03 | -2,41 | TSC complex subunit 1 | No | No | 16 |
| Csf1r | 9,38E-03 | -2,41 | colony stimulating factor 1 receptor | No | No | 13 |
| Sgpl1 | 1,27E-06 | -2,40 | sphingosine phosphate lyase 1 | No | No | - |
| Mcm3ap | 1,12E-02 | -2,40 | minichromosome maintenance complex component 3 assoc | No | No | - |
| Tmcc1 | 5,86E-03 | -2,40 | transmembrane and coiled coil domains 1 | No | No | - |
| Adar | 6,59E-03 | -2,40 | adenosine deaminase, RNA-specific | Yes | No | - |
| Clstn1 | 6,44E-04 | -2,40 | calsyntenin 1 | No | No | - |
| Bscl2 | 1,48E-03 | -2,40 | Berardinelli-Seip congenital lipodystrophy 2 (seipin) | No | No | - |
| Lamc1 | 8,26E-03 | -2,40 | laminin, gamma 1 | No | No | - |
| Panx1 | 4,84E-03 | -2,39 | pannexin 1 | No | No | - |
| Bhlhe41 | 1,68E-02 | -2,39 | basic helix-loop-helix family, member e41 | No | No | - |
| Vps41 | 6,76E-03 | -2,39 | VPS41 HOPS complex subunit | No | No | - |
| Hsd17b11 | 3,18E-03 | -2,39 | hydroxysteroid (17-beta) dehydrogenase 11 | Yes | No | - |
| Cavin2 | 4,90E-07 | -2,39 | caveolae associated 2 | No | No | - |
| Col6a6 | 1,13E-02 | -2,39 | collagen, type VI, alpha 6 | No | No | - |
| Fbxw7 | 9,96E-03 | -2,39 | F-box and WD-40 domain protein 7 | No | No | 1 |
| Polr1a | 8,61E-03 | -2,38 | polymerase (RNA) I polypeptide A | No | No | - |
| Il6st | 5,91E-04 | -2,38 | interleukin 6 signal transducer | Yes | No | 1 |
| Scfd1 | 2,89E-03 | -2,38 | Sec1 family domain containing 1 | No | No | - |
| Bak1 | 1,27E-03 | -2,38 | BCL2-antagonist/killer 1 | No | No | - |
| Apom | 3,53E-02 | -2,38 | apolipoprotein M | No | No | - |
| Ascc2 | 2,10E-02 | -2,38 | activating signal cointegrator 1 complex subunit 2 | No | No | - |
| Atp6v0e2 | 1,13E-02 | -2,38 | ATPase, H+ transporting, lysosomal V0 subunit E2 | No | No | - |
| Npr2 | 1,40E-02 | -2,38 | natriuretic peptide receptor 2 | No | No | - |
| Trpv2 | 2,81E-02 | -2,38 | transient receptor potential cation channel, subfamily V, me | No | No | - |
| Ldlrad4 | 9,22E-03 | -2,37 | low density lipoprotein receptor class A domain containing | No | No | - |
| Slco3a1 | 1,29E-03 | -2,37 | solute carrier organic anion transporter family, member 3a1 | No | No | - |
| Scpep1 | 1,66E-05 | -2,37 | serine carboxypeptidase 1 | No | No | - |
| Ccdc24 | 7,00E-03 | -2,37 | coiled-coil domain containing 24 | No | No | - |
| Edf1 | 3,27E-05 | -2,36 | endothelial differentiation-related factor 1 | No | No | - |
| Dgkh | 5,24E-03 | -2,36 | diacylglycerol kinase, eta | No | No | - |
| Gm49388 | 1,68E-02 | -2,36 | predicted gene, 49388 | No | No | - |
| Tecpr1 | 2,31E-03 | -2,36 | tectonin beta-propeller repeat containing 1 | No | No | - |
| Sh3bgrl | 2,33E-03 | -2,36 | SH3-binding domain glutamic acid-rich protein like | No | No | - |
| Nucb1 | 2,72E-04 | -2,36 | nucleobindin 1 | No | No | - |
| Cpeb4 | 1,30E-03 | -2,35 | cytoplasmic polyadenylation element binding protein 4 | No | No | - |
| Natd1 | 2,29E-02 | -2,35 | N-acetyltransferase domain containing 1 | No | No | - |
| Adamtsl5 | 5,04E-03 | -2,35 | ADAMTS-like 5 | No | No | - |
| Gpr107 | 8,01E-05 | -2,35 | G protein-coupled receptor 107 | No | No | - |
| Cnnm3 | 1,21E-02 | -2,35 | cyclin M3 | No | No | - |
| Cttnbp2nl | 2,22E-04 | -2,35 | CTTNBP2 N-terminal like | No | No | - |
| Acly | 7,71E-06 | -2,34 | ATP citrate lyase | Yes | No | - |
| Ppm1b | 1,59E-04 | -2,34 | protein phosphatase 1B, magnesium dependent, beta isoforr | No | No | - |
| Cxcl1 | 5,50E-03 | -2,34 | chemokine (C-X-C motif) ligand 1 | No | No | - |
| Ogt | 3,14E-03 | -2,34 | O-linked N-acetylglucosamine (GlcNAc) transferase (UDP- | No | No | - |
| Glb1 | 2,99E-04 | -2,34 | galactosidase, beta 1 | No | No | - |
| Lama2 | 3,91E-03 | -2,34 | laminin, alpha 2 | No | No | - |
| Ift140 | 1,95E-03 | -2,34 | intraflagellar transport 140 | No | No | - |
| Gbf1 | 2,80E-03 | -2,34 | golgi-specific brefeldin A-resistance factor 1 | No | No | - |
| Otud7b | 6,77E-03 | -2,33 | OTU domain containing 7B | No | No | - |
| Ndst2 | 1,99E-03 | -2,33 | N-deacetylase/N-sulfotransferase (heparan glucosaminyl) 2 | No | No | - |
| Slc6a9 | 8,12E-03 | -2,33 | solute carrier family 6 (neurotransmitter transporter, glycine | No | No | - |
| Usp10 | 1,02E-02 | -2,33 | ubiquitin specific peptidase 10 | No | No | - |
| Gm15459 | 9,54E-04 | -2,33 | predicted gene 15459 | No | No | - |
| Tfe3 | 1,83E-03 | -2,33 | transcription factor E3 | No | No | - |
| Sil1 | 2,90E-03 | -2,33 | endoplasmic reticulum chaperone SIL1 homolog (S. cerevis | No | No | - |
| Wdr7 | 4,70E-02 | -2,33 | WD repeat domain 7 | No | No | - |
| Agap3 | 3,43E-04 | -2,32 | ArfGAP with GTPase domain, ankyrin repeat and PH doma | No | No | - |
| Zbtb38 | 3,99E-04 | -2,32 | zinc finger and BTB domain containing 38 | No | No | - |
| Fgfrl1 | 1,59E-02 | -2,32 | fibroblast growth factor receptor-like 1 | No | No | - |
| Slc8b1 | 1,04E-02 | -2,32 | solute carrier family 8 (sodium/lithium/calcium exchanger), | No | No | - |
| Tmbim1 | 2,66E-06 | -2,32 | transmembrane BAX inhibitor motif containing 1 | No | No | - |
| Rab7b | 2,96E-04 | -2,32 | RAB7B, member RAS oncogene family | No | No | - |

| | | | | | | |
|---|---|---|---|---|---|---|
| Recql | 9,90E-03 | -2,32 | RecQ protein-like | No | No | - |
| Neurl4 | 1,90E-02 | -2,32 | neuralized E3 ubiquitin protein ligase 4 | No | No | - |
| Plcxd2 | 3,85E-02 | -2,32 | phosphatidylinositol-specific phospholipase C, X domain cc | No | No | - |
| Zfp260 | 2,14E-02 | -2,32 | zinc finger protein 260 | No | No | - |
| Dlg5 | 8,24E-04 | -2,32 | discs large MAGUK scaffold protein 5 | No | No | - |
| Zswim8 | 2,05E-03 | -2,31 | zinc finger SWIM-type containing 8 | No | No | - |
| Cad | 1,60E-03 | -2,31 | carbamoyl-phosphate synthetase 2, aspartate transcarbamyla | No | No | 237 |
| Spag8 | 1,48E-02 | -2,31 | sperm associated antigen 8 | No | No | - |
| Gripap1 | 2,89E-03 | -2,31 | GRIP1 associated protein 1 | No | No | - |
| Tmem106a | 4,75E-02 | -2,31 | transmembrane protein 106A | No | No | - |
| Pcyt2 | 2,51E-03 | -2,31 | phosphate cytidylyltransferase 2, ethanolamine | Yes | No | - |
| Coro1c | 1,12E-04 | -2,30 | coronin, actin binding protein 1C | No | No | - |
| Apc | 4,61E-03 | -2,30 | APC, WNT signaling pathway regulator | Yes | No | 53 |
| Rictor | 1,98E-02 | -2,30 | RPTOR independent companion of MTOR, complex 2 | No | No | 1 |
| Pam | 6,63E-04 | -2,30 | peptidylglycine alpha-amidating monooxygenase | No | No | 102 |
| Eif1a | 1,13E-04 | -2,30 | eukaryotic translation initiation factor 1A | No | No | - |
| Itsn1 | 5,37E-04 | -2,30 | intersectin 1 (SH3 domain protein 1A) | No | No | - |
| Exosc10 | 1,85E-03 | -2,30 | exosome component 10 | No | No | - |
| Ssh1 | 9,52E-04 | -2,30 | slingshot protein phosphatase 1 | No | No | - |
| G6pdx | 5,13E-03 | -2,30 | glucose-6-phosphate dehydrogenase X-linked | No | No | 15 |
| Inhba | 1,76E-04 | -2,30 | inhibin beta-A | No | No | 1 |
| Gm2a | 2,38E-03 | -2,30 | GM2 ganglioside activator protein | No | No | 2 |
| Fkbp10 | 7,87E-04 | -2,30 | FK506 binding protein 10 | No | No | 19 |
| Ehd4 | 1,76E-03 | -2,30 | EH-domain containing 4 | No | No | - |
| Eif2b5 | 1,92E-03 | -2,29 | eukaryotic translation initiation factor 2B, subunit 5 epsilon | No | No | - |
| Dgkz | 4,63E-02 | -2,29 | diacylglycerol kinase zeta | No | No | - |
| Stx1b | 2,34E-02 | -2,29 | syntaxin 1B | No | No | - |
| Impact | 1,70E-02 | -2,29 | impact, RWD domain protein | No | No | 6 |
| Mroh1 | 1,70E-03 | -2,29 | maestro heat-like repeat family member 1 | No | No | - |
| Bop1 | 1,52E-03 | -2,29 | block of proliferation 1 | No | No | - |
| Coq10a | 3,72E-02 | -2,29 | coenzyme Q10A | No | No | - |
| Rfc1 | 9,75E-03 | -2,29 | replication factor C (activator 1) 1 | No | No | - |
| Emsy | 1,08E-03 | -2,28 | EMSY, BRCA2-interacting transcriptional repressor | No | No | - |
| Eda2r | 1,10E-02 | -2,28 | ectodysplasin A2 receptor | No | No | - |
| Ftl2-ps | 1,47E-02 | -2,28 | ferritin light polypeptide 2, pseudogene | No | No | - |
| Pyroxd1 | 1,72E-02 | -2,28 | pyridine nucleotide-disulphide oxidoreductase domain 1 | No | No | - |
| Arl5a | 8,68E-04 | -2,28 | ADP-ribosylation factor-like 5A | No | No | - |
| Cgref1 | 1,37E-02 | -2,28 | cell growth regulator with EF hand domain 1 | No | No | - |
| Hsd3b7 | 2,31E-02 | -2,28 | hydroxy-delta-5-steroid dehydrogenase, 3 beta- and steroid | No | No | - |
| Clip3 | 2,17E-03 | -2,28 | CAP-GLY domain containing linker protein 3 | No | No | - |
| B4galt1 | 1,54E-06 | -2,28 | UDP-Gal:betaGlcNAc beta 1,4- galactosyltransferase, polyp | No | No | - |
| Tmem98 | 8,91E-04 | -2,27 | transmembrane protein 98 | No | No | - |
| Pkn1 | 4,70E-03 | -2,27 | protein kinase N1 | No | No | - |
| Dpep1 | 1,17E-03 | -2,27 | dipeptidase 1 | No | No | - |
| Ccdc93 | 2,12E-02 | -2,27 | coiled-coil domain containing 93 | No | No | - |
| Emc3 | 2,07E-04 | -2,27 | ER membrane protein complex subunit 3 | No | No | - |
| Ywhab | 1,07E-04 | -2,27 | tyrosine 3-monooxygenase/tryptophan 5-monooxygenase ac | No | No | - |
| Gm13625 | 1,51E-03 | -2,27 | predicted gene 13625 | No | No | - |
| Cyp51 | 3,59E-06 | -2,26 | cytochrome P450, family 51 | No | No | - |
| Map1a | 4,80E-03 | -2,26 | microtubule-associated protein 1 A | No | No | - |
| Cdk16 | 4,28E-03 | -2,26 | cyclin-dependent kinase 16 | No | No | - |
| Rubcn | 1,93E-02 | -2,26 | RUN domain and cysteine-rich domain containing, Beclin 1 | No | No | - |
| Nrp2 | 9,94E-05 | -2,26 | neuropilin 2 | No | No | - |
| Ddx58 | 9,95E-03 | -2,26 | DEAD/H box helicase 58 | No | No | 1 |
| Stk40 | 1,13E-02 | -2,26 | serine/threonine kinase 40 | No | No | - |
| Ubr3 | 2,03E-03 | -2,26 | ubiquitin protein ligase E3 component n-recognin 3 | No | No | - |
| Zzz3 | 1,12E-02 | -2,26 | zinc finger, ZZ domain containing 3 | No | No | - |
| Tbc1d10a | 6,66E-03 | -2,26 | TBC1 domain family, member 10a | No | No | - |
| Capn1 | 6,69E-03 | -2,26 | calpain 1 | No | No | - |
| Crip2 | 2,83E-04 | -2,26 | cysteine rich protein 2 | No | No | - |
| Eif3c | 4,59E-04 | -2,25 | eukaryotic translation initiation factor 3, subunit C | No | No | - |
| BC024978 | 4,05E-02 | -2,25 | cDNA sequence BC024978 | No | No | - |
| Lpl | 3,70E-02 | -2,25 | lipoprotein lipase | No | No | 4 |
| Otud3 | 3,42E-02 | -2,25 | OTU domain containing 3 | No | No | - |
| Sft2d2 | 1,78E-03 | -2,25 | SFT2 domain containing 2 | No | No | - |
| Lamb1 | 1,51E-05 | -2,25 | laminin B1 | No | No | 12 |
| Clip1 | 5,62E-03 | -2,25 | CAP-GLY domain containing linker protein 1 | No | No | - |
| Klhl26 | 5,71E-03 | -2,25 | kelch-like 26 | No | No | - |
| Ahctf1 | 1,25E-02 | -2,24 | AT hook containing transcription factor 1 | No | No | - |
| Bclaf1 | 2,50E-03 | -2,24 | BCL2-associated transcription factor 1 | No | No | - |
| Hinfp | 3,00E-02 | -2,24 | histone H4 transcription factor | No | No | - |
| Pqlc2 | 3,67E-02 | -2,24 | solute carrier family 66 member 1 | No | No | - |
| Brd3 | 1,38E-03 | -2,24 | bromodomain containing 3 | No | No | - |

| | | | | | | |
|---|---|---|---|---|---|---|
| Eri3 | 2,46E-03 | -2,24 | exoribonuclease 3 | No | No | - |
| Pdgfrl | 7,83E-03 | -2,24 | platelet-derived growth factor receptor-like | No | No | - |
| Sirpa | 6,24E-03 | -2,24 | signal-regulatory protein alpha | No | No | - |
| Asxl1 | 2,31E-02 | -2,24 | ASXL transcriptional regulator 1 | No | No | 1 |
| Itgb3 | 3,39E-02 | -2,24 | integrin beta 3 | No | No | - |
| Cox10 | 4,95E-03 | -2,24 | heme A:farnesyltransferase cytochrome c oxidase assembly | No | No | 3 |
| Tnfrsf26 | 6,24E-03 | -2,24 | tumor necrosis factor receptor superfamily, member 26 | No | No | - |
| Ddr1 | 1,53E-03 | -2,23 | discoidin domain receptor family, member 1 | No | No | 53 |
| Golgb1 | 1,21E-02 | -2,23 | golgi autoantigen, golgin subfamily b, macrogolgin 1 | No | No | - |
| Rcn3 | 4,06E-05 | -2,23 | reticulocalbin 3, EF-hand calcium binding domain | No | No | - |
| Pacs2 | 3,68E-03 | -2,23 | phosphofurin acidic cluster sorting protein 2 | No | No | - |
| Tmem214 | 4,28E-03 | -2,23 | transmembrane protein 214 | No | No | - |
| Fads2 | 1,12E-03 | -2,23 | fatty acid desaturase 2 | No | No | - |
| Map3k20 | 1,60E-03 | -2,23 | mitogen-activated protein kinase kinase kinase 20 | No | No | - |
| Sugp1 | 4,57E-02 | -2,23 | SURP and G patch domain containing 1 | No | No | - |
| Atp9b | 1,43E-02 | -2,23 | ATPase, class II, type 9B | No | No | - |
| Senp2 | 4,99E-03 | -2,23 | SUMO/sentrin specific peptidase 2 | No | No | - |
| Zkscan6 | 4,73E-02 | -2,23 | zinc finger with KRAB and SCAN domains 6 | No | No | - |
| Pakap | 6,09E-04 | -2,22 | paralemmin A kinase anchor protein | No | No | - |
| Usp16 | 8,74E-04 | -2,22 | ubiquitin specific peptidase 16 | No | No | - |
| Pakap | 5,83E-04 | -2,22 | paralemmin A kinase anchor protein | No | No | - |
| D10Wsu102e | 1,19E-03 | -2,22 | NOP protein chaperone 1 | No | No | - |
| Ifi35 | 4,62E-06 | -2,22 | interferon-induced protein 35 | Yes | No | - |
| Tcp1 | 2,35E-05 | -2,22 | t-complex protein 1 | No | No | - |
| Taok1 | 6,16E-03 | -2,22 | TAO kinase 1 | No | No | - |
| Far1 | 1,89E-03 | -2,22 | fatty acyl CoA reductase 1 | No | No | - |
| Bcl2 | 4,11E-02 | -2,22 | B cell leukemia/lymphoma 2 | Yes | No | 14 |
| Ccdc80 | 7,56E-04 | -2,21 | coiled-coil domain containing 80 | No | No | - |
| Peak1 | 1,28E-03 | -2,21 | pseudopodium-enriched atypical kinase 1 | No | No | - |
| Slc25a13 | 1,84E-02 | -2,21 | solute carrier family 25 (mitochondrial carrier, adenine nucl | No | No | - |
| Ttll7 | 2,03E-02 | -2,21 | tubulin tyrosine ligase-like family, member 7 | No | No | - |
| Alad | 2,88E-02 | -2,21 | aminolevulinate, delta-, dehydratase | No | No | - |
| H2-DMb1 | 3,79E-03 | -2,21 | histocompatibility 2, class II, locus Mb1 | Yes | No | - |
| Vps16 | 4,98E-03 | -2,21 | VSP16 CORVET/HOPS core subunit | No | No | - |
| Zfp322a | 2,30E-02 | -2,21 | zinc finger protein 322A | No | No | - |
| Nploc4 | 3,00E-03 | -2,21 | NPL4 homolog, ubiquitin recognition factor | No | No | - |
| Mov10 | 3,77E-02 | -2,21 | Mov10 RISC complex RNA helicase | No | No | - |
| Rnpepl1 | 3,89E-02 | -2,21 | arginyl aminopeptidase (aminopeptidase B)-like 1 | No | No | - |
| Slc8a1 | 2,99E-02 | -2,20 | solute carrier family 8 (sodium/calcium exchanger), membe | No | No | - |
| Orai3 | 5,54E-03 | -2,20 | ORAI calcium release-activated calcium modulator 3 | No | No | - |
| Ppp6r3 | 6,30E-04 | -2,20 | protein phosphatase 6, regulatory subunit 3 | No | No | - |
| Rnf216 | 3,97E-02 | -2,20 | ring finger protein 216 | No | No | - |
| Tpcn1 | 1,36E-02 | -2,20 | two pore channel 1 | No | No | - |
| Rock1 | 3,17E-03 | -2,20 | Rho-associated coiled-coil containing protein kinase 1 | No | No | 24 |
| Gfpt2 | 1,62E-03 | -2,20 | glutamine fructose-6-phosphate transaminase 2 | No | No | - |
| Gapvd1 | 5,35E-04 | -2,20 | GTPase activating protein and VPS9 domains 1 | No | No | - |
| Rabgap1 | 3,54E-03 | -2,20 | RAB GTPase activating protein 1 | No | No | - |
| Hmox1 | 1,40E-02 | -2,19 | heme oxygenase 1 | Yes | No | 7 |
| Chmp1a | 7,35E-04 | -2,19 | charged multivesicular body protein 1A | No | No | - |
| Marchf7 | 1,69E-02 | -2,19 | membrane associated ring-CH-type finger 7 | No | No | - |
| Ttc13 | 4,63E-04 | -2,19 | tetratricopeptide repeat domain 13 | No | No | - |
| Urm1 | 3,13E-03 | -2,19 | ubiquitin related modifier 1 | No | No | - |
| Septin5 | 8,28E-04 | -2,19 | septin 5 | No | No | - |
| Gstm2 | 5,02E-05 | -2,19 | glutathione S-transferase, mu 2 | No | No | 30 |
| Sf3b2 | 9,31E-05 | -2,19 | splicing factor 3b, subunit 2 | No | No | - |
| Dnajc16 | 1,71E-02 | -2,19 | DnaJ heat shock protein family (Hsp40) member C16 | No | No | - |
| Piezo2 | 1,00E-02 | -2,19 | piezo-type mechanosensitive ion channel component 2 | No | No | - |
| Phf21a | 3,67E-03 | -2,19 | PHD finger protein 21A | No | No | - |
| Macf1 | 7,81E-04 | -2,19 | microtubule-actin crosslinking factor 1 | No | No | - |
| Poldip2 | 3,95E-05 | -2,19 | polymerase (DNA-directed), delta interacting protein 2 | No | No | - |
| Elk4 | 3,83E-03 | -2,18 | ELK4, member of ETS oncogene family | No | No | - |
| Foxg1 | 4,38E-02 | -2,18 | forkhead box G1 | No | No | - |
| Nol6 | 2,20E-02 | -2,18 | nucleolar protein family 6 (RNA-associated) | No | No | - |
| Nfat5 | 2,74E-03 | -2,18 | nuclear factor of activated T cells 5 | No | No | - |
| Glul | 1,12E-02 | -2,18 | glutamate-ammonia ligase (glutamine synthetase) | No | No | - |
| Adcy7 | 1,57E-03 | -2,18 | adenylate cyclase 7 | No | No | - |
| Ralgapb | 6,66E-03 | -2,18 | Ral GTPase activating protein, beta subunit (non-catalytic) | No | No | - |
| Chrnb2 | 2,32E-02 | -2,18 | cholinergic receptor, nicotinic, beta polypeptide 2 (neuronal | No | No | - |
| Anapc2 | 4,36E-03 | -2,18 | anaphase promoting complex subunit 2 | No | No | - |
| Orc2 | 2,17E-02 | -2,18 | origin recognition complex, subunit 2 | No | No | - |
| Rbm15 | 4,48E-02 | -2,18 | RNA binding motif protein 15 | No | No | - |
| Man1c1 | 8,26E-04 | -2,18 | mannosidase, alpha, class 1C, member 1 | No | No | - |
| Gpha2 | 8,69E-03 | -2,18 | glycoprotein hormone alpha 2 | No | No | - |

| | | | | | | |
|---|---|---|---|---|---|---|
| Rbms3 | 3,25E-03 | -2,17 | RNA binding motif, single stranded interacting protein | No | No | 1 |
| Asxl2 | 1,72E-02 | -2,17 | ASXL transcriptional regulator 2 | No | No | - |
| Hnrnpul2 | 1,32E-03 | -2,17 | heterogeneous nuclear ribonucleoprotein U-like 2 | No | No | - |
| Wsb2 | 4,41E-04 | -2,17 | WD repeat and SOCS box-containing 2 | No | No | - |
| Pcdh19 | 1,31E-03 | -2,17 | protocadherin 19 | No | No | - |
| Uba1 | 9,70E-05 | -2,17 | ubiquitin-like modifier activating enzyme 1 | No | No | 1 |
| Cltc | 7,67E-06 | -2,17 | clathrin, heavy polypeptide (Hc) | No | No | - |
| Atp1a1 | 1,95E-03 | -2,17 | ATPase, Na+/K+ transporting, alpha 1 polypeptide | No | No | - |
| Cep95 | 3,32E-02 | -2,17 | centrosomal protein 95 | No | No | - |
| Eea1 | 8,61E-03 | -2,17 | early endosome antigen 1 | No | No | 11 |
| Sdccag8 | 3,31E-02 | -2,17 | serologically defined colon cancer antigen 8 | No | No | - |
| Flnb | 1,73E-05 | -2,17 | filamin, beta | No | No | - |
| Peli2 | 1,39E-02 | -2,16 | pellino 2 | No | No | - |
| Ipo5 | 7,91E-05 | -2,16 | importin 5 | No | No | - |
| Foxn3 | 2,78E-03 | -2,16 | forkhead box N3 | No | No | - |
| Aldh4a1 | 2,29E-03 | -2,16 | aldehyde dehydrogenase 4 family, member A1 | No | No | - |
| Hdlbp | 3,70E-04 | -2,16 | high density lipoprotein (HDL) binding protein | No | No | - |
| Ctnnb1 | 9,36E-05 | -2,16 | catenin (cadherin associated protein), beta 1 | Yes | No | 3 |
| Sf3b3 | 5,28E-04 | -2,16 | splicing factor 3b, subunit 3 | No | No | - |
| Akt1s1 | 1,17E-02 | -2,16 | AKT1 substrate 1 (proline-rich) | No | No | - |
| Pdp2 | 4,59E-03 | -2,16 | pyruvate dehyrogenase phosphatase catalytic subunit 2 | No | No | - |
| Epb41 | 1,50E-03 | -2,16 | erythrocyte membrane protein band 4.1 | No | No | - |
| Zfp469 | 3,15E-02 | -2,16 | zinc finger protein 469 | No | No | - |
| Sltm | 2,56E-03 | -2,16 | SAFB-like, transcription modulator | No | No | - |
| Efcab14 | 4,36E-02 | -2,16 | EF-hand calcium binding domain 14 | No | No | - |
| Os9 | 4,35E-02 | -2,16 | amplified in osteosarcoma | No | No | - |
| Tcf7l2 | 7,44E-03 | -2,16 | transcription factor 7 like 2, T cell specific, HMG box | No | Yes | 1 |
| Rpl26 | 9,91E-07 | -2,16 | ribosomal protein L26 | No | No | - |
| Pnma2 | 1,27E-02 | -2,16 | paraneoplastic antigen MA2 | No | No | - |
| Rpp38 | 1,72E-02 | -2,16 | ribonuclease P/MRP 38 subunit | No | No | - |
| Mybbp1a | 1,98E-03 | -2,15 | MYB binding protein (P160) 1a | No | No | - |
| Ap3d1 | 8,22E-04 | -2,15 | adaptor-related protein complex 3, delta 1 subunit | No | No | - |
| Fxr2 | 1,42E-03 | -2,15 | fragile X mental retardation, autosomal homolog 2 | No | No | - |
| Gga2 | 1,32E-03 | -2,15 | golgi associated, gamma adaptin ear containing, ARF bindi | No | No | - |
| Zfp212 | 2,70E-02 | -2,15 | Zinc finger protein 212 | No | No | - |
| Mir100hg | 4,98E-03 | -2,15 | Mir100 Mirlet7a-2 Mir125b-1 cluster host gene | No | No | - |
| Wdfy2 | 1,60E-02 | -2,15 | WD repeat and FYVE domain containing 2 | No | No | - |
| Pdgfrb | 3,34E-03 | -2,15 | platelet derived growth factor receptor, beta polypeptide | No | No | 7 |
| Ate1 | 4,51E-04 | -2,15 | arginyltransferase 1 | No | No | - |
| Arhgap29 | 2,07E-03 | -2,15 | Rho GTPase activating protein 29 | No | No | - |
| Ncf2 | 1,38E-02 | -2,15 | neutrophil cytosolic factor 2 | No | No | 5 |
| Pcyt1a | 7,12E-04 | -2,15 | phosphate cytidylyltransferase 1, choline, alpha isoform | No | No | - |
| Parp3 | 8,64E-06 | -2,15 | poly (ADP-ribose) polymerase family, member 3 | No | No | - |
| Prpf4b | 4,33E-02 | -2,15 | pre-mRNA processing factor 4B | No | No | - |
| Pik3cd | 9,94E-04 | -2,14 | phosphatidylinositol-4,5-bisphosphate 3-kinase catalytic sul | No | No | 2 |
| Pdia3 | 3,70E-04 | -2,14 | protein disulfide isomerase associated 3 | No | No | 12 |
| Tnfaip1 | 5,35E-04 | -2,14 | tumor necrosis factor, alpha-induced protein 1 (endothelial) | No | No | - |
| Gorasp2 | 4,94E-04 | -2,14 | golgi reassembly stacking protein 2 | No | No | - |
| Podn | 2,22E-02 | -2,14 | podocan | No | No | - |
| Abhd17c | 1,04E-03 | -2,14 | abhydrolase domain containing 17C | No | No | - |
| Dnajc10 | 1,67E-03 | -2,14 | DnaJ heat shock protein family (Hsp40) member C10 | No | No | - |
| Dlg1 | 5,33E-03 | -2,14 | discs large MAGUK scaffold protein 1 | No | No | - |
| Bms1 | 2,63E-03 | -2,14 | BMS1, ribosome biogenesis factor | No | No | - |
| Spns2 | 2,31E-03 | -2,14 | spinster homolog 2 | No | No | 2 |
| Rassf1 | 2,25E-03 | -2,14 | Ras association (RalGDS/AF-6) domain family member 1 | No | No | 1 |
| Bnip2 | 1,98E-04 | -2,14 | BCL2/adenovirus E1B interacting protein 2 | No | No | - |
| Atp6v0a2 | 1,25E-03 | -2,14 | ATPase, H+ transporting, lysosomal V0 subunit A2 | No | No | - |
| Pgd | 1,10E-03 | -2,14 | phosphogluconate dehydrogenase | No | No | 169 |
| Supt6 | 4,61E-03 | -2,13 | SPT6, histone chaperone and transcription elongation factor | No | No | - |
| Cped1 | 2,38E-02 | -2,13 | cadherin-like and PC-esterase domain containing 1 | No | No | - |
| Trim56 | 7,26E-03 | -2,13 | tripartite motif-containing 56 | No | No | - |
| Dkk3 | 4,14E-04 | -2,13 | dickkopf WNT signaling pathway inhibitor 3 | No | No | - |
| Zfp652 | 3,16E-02 | -2,13 | zinc finger protein 652 | No | No | - |
| Hyou1 | 2,38E-03 | -2,13 | hypoxia up-regulated 1 | No | No | - |
| Isoc2a | 2,36E-02 | -2,13 | isochorismatase domain containing 2a | No | No | - |
| Preb | 6,99E-03 | -2,13 | prolactin regulatory element binding | No | No | - |
| Slc51a | 8,49E-04 | -2,13 | solute carrier family 51, alpha subunit | No | No | - |
| Brd4 | 3,57E-03 | -2,13 | bromodomain containing 4 | No | No | 22 |
| Ssb | 6,11E-04 | -2,13 | Sjogren syndrome antigen B | No | No | 2 |
| Exoc5 | 3,40E-03 | -2,13 | exocyst complex component 5 | No | No | - |
| Acox3 | 4,13E-03 | -2,13 | acyl-Coenzyme A oxidase 3, pristanoyl | No | No | - |
| Crlf2 | 7,16E-04 | -2,13 | cytokine receptor-like factor 2 | No | No | - |
| Celf2 | 3,08E-03 | -2,12 | CUGBP, Elav-like family member 2 | No | No | - |

| | | | | | | |
|---|---|---|---|---|---|---|
| Ythdf3 | 1,76E-03 | -2,12 | YTH N6-methyladenosine RNA binding protein 3 | No | No | - |
| Synpo | 9,40E-04 | -2,12 | synaptopodin | No | No | - |
| Creld1 | 2,57E-02 | -2,12 | cysteine-rich with EGF-like domains 1 | No | No | - |
| Drosha | 1,19E-02 | -2,12 | drosha, ribonuclease type III | No | No | 7 |
| Fxr1 | 8,56E-03 | -2,12 | fragile X mental retardation gene 1, autosomal homolog | No | No | - |
| Gtpbp4 | 4,84E-02 | -2,12 | GTP binding protein 4 | No | No | - |
| Ppp2r5b | 1,47E-02 | -2,12 | protein phosphatase 2, regulatory subunit B', beta | No | No | - |
| Cacul1 | 3,86E-04 | -2,11 | CDK2 associated, cullin domain 1 | No | No | - |
| Iqsec2 | 1,52E-02 | -2,11 | IQ motif and Sec7 domain 2 | No | No | - |
| Ogdh | 1,64E-02 | -2,11 | oxoglutarate (alpha-ketoglutarate) dehydrogenase (lipoamid | No | No | - |
| Slc30a1 | 2,93E-02 | -2,11 | solute carrier family 30 (zinc transporter), member 1 | No | No | - |
| 1700017B05Rik | 1,64E-03 | -2,11 | RIKEN cDNA 1700017B05 gene | No | No | - |
| Mia3 | 1,43E-03 | -2,11 | MIA SH3 domain ER export factor 3 | No | No | - |
| Atp11a | 7,33E-03 | -2,11 | ATPase, class VI, type 11A | No | No | 1 |
| Ccdc9b | 2,42E-02 | -2,11 | coiled-coil domain containing 9B | No | No | - |
| Ddb1 | 1,46E-04 | -2,11 | damage specific DNA binding protein 1 | No | No | - |
| Pdpn | 4,95E-05 | -2,11 | podoplanin | No | No | - |
| Zfp219 | 3,79E-03 | -2,11 | zinc finger protein 219 | No | No | - |
| Pmaip1 | 9,08E-03 | -2,11 | phorbol-12-myristate-13-acetate-induced protein 1 | Yes | No | - |
| Htra3 | 4,85E-02 | -2,11 | HtrA serine peptidase 3 | No | No | - |
| Tram2 | 7,16E-03 | -2,10 | translocating chain-associating membrane protein 2 | No | No | - |
| Akap8 | 2,52E-02 | -2,10 | A kinase (PRKA) anchor protein 8 | No | No | - |
| Aif1l | 8,72E-03 | -2,10 | allograft inflammatory factor 1-like | No | No | - |
| Lnpep | 1,14E-02 | -2,10 | leucyl/cystinyl aminopeptidase | No | No | - |
| Npm1 | 3,70E-03 | -2,10 | nucleophosmin 1 | No | No | 2 |
| Arrdc1 | 2,10E-02 | -2,10 | arrestin domain containing 1 | No | No | - |
| Tmem248 | 4,78E-04 | -2,10 | transmembrane protein 248 | No | No | - |
| Mmp3 | 6,34E-03 | -2,10 | matrix metallopeptidase 3 | No | No | 21 |
| Adck5 | 1,71E-02 | -2,10 | aarF domain containing kinase 5 | No | No | - |
| Gss | 5,59E-03 | -2,10 | glutathione synthetase | No | No | 2 |
| Rtf2 | 2,04E-03 | -2,10 | replication termination factor 2 | No | No | - |
| Zfp148 | 2,59E-02 | -2,10 | zinc finger protein 148 | No | No | - |
| Fubp1 | 4,75E-04 | -2,10 | far upstream element (FUSE) binding protein 1 | No | No | - |
| Cdc42bpa | 3,88E-03 | -2,10 | CDC42 binding protein kinase alpha | No | No | - |
| Qsox1 | 5,83E-06 | -2,09 | quiescin Q6 sulfhydryl oxidase 1 | Yes | No | - |
| Csgalnact1 | 4,93E-03 | -2,09 | chondroitin sulfate N-acetylgalactosaminyltransferase 1 | No | No | - |
| Camsap2 | 1,36E-02 | -2,09 | calmodulin regulated spectrin-associated protein family, me | No | No | - |
| Ehd2 | 2,51E-03 | -2,09 | EH-domain containing 2 | No | No | - |
| Haus7 | 1,29E-02 | -2,09 | HAUS augmin-like complex, subunit 7 | No | No | - |
| Septin7 | 1,80E-03 | -2,09 | septin 7 | No | No | - |
| Smg5 | 3,60E-02 | -2,09 | SMG5 nonsense mediated mRNA decay factor | No | No | - |
| Neu1 | 3,99E-03 | -2,09 | neuraminidase 1 | No | No | 11 |
| 4930442H23Rik | 3,12E-05 | -2,09 | RIKEN cDNA 4930442H23 gene | No | No | - |
| Cds2 | 3,23E-03 | -2,09 | CDP-diacylglycerol synthase (phosphatidate cytidylyltransf | No | No | - |
| Gpam | 3,26E-02 | -2,09 | glycerol-3-phosphate acyltransferase, mitochondrial | No | No | - |
| Mfap5 | 2,70E-04 | -2,09 | microfibrillar associated protein 5 | No | No | - |
| Grk6 | 4,12E-03 | -2,09 | G protein-coupled receptor kinase 6 | No | No | - |
| Cd248 | 1,76E-03 | -2,08 | CD248 antigen, endosialin | No | No | 12 |
| Orai1 | 6,30E-04 | -2,08 | ORAI calcium release-activated calcium modulator 1 | No | No | - |
| Prpf6 | 3,93E-03 | -2,08 | pre-mRNA splicing factor 6 | No | No | - |
| Sae1 | 4,36E-03 | -2,08 | SUMO1 activating enzyme subunit 1 | No | No | 3 |
| Zfx | 3,15E-02 | -2,08 | zinc finger protein X-linked | No | No | - |
| Atp13a1 | 2,14E-02 | -2,08 | ATPase type 13A1 | No | No | - |
| Zbtb7a | 1,63E-03 | -2,08 | zinc finger and BTB domain containing 7a | No | No | - |
| Elp1 | 4,07E-02 | -2,08 | elongator complex protein 1 | No | No | - |
| Jak2 | 2,05E-02 | -2,08 | Janus kinase 2 | Yes | No | 46 |
| Eef2 | 4,64E-05 | -2,08 | eukaryotic translation elongation factor 2 | No | No | 1 |
| B230219D22Rik | 9,17E-03 | -2,07 | RIKEN cDNA B230219D22 gene | No | No | - |
| Phc3 | 8,22E-03 | -2,07 | polyhomeotic 3 | No | No | - |
| Zmat3 | 3,22E-03 | -2,07 | zinc finger matrin type 3 | No | No | - |
| Abhd1 | 1,27E-02 | -2,07 | abhydrolase domain containing 1 | No | No | - |
| Anapc7 | 1,98E-02 | -2,07 | anaphase promoting complex subunit 7 | No | No | - |
| Polr2a | 4,54E-03 | -2,07 | polymerase (RNA) II (DNA directed) polypeptide A | No | No | - |
| Arhgap35 | 3,67E-03 | -2,07 | Rho GTPase activating protein 35 | No | No | - |
| Slc41a2 | 5,98E-03 | -2,07 | solute carrier family 41, member 2 | No | No | - |
| Ugt1a2 | 1,24E-03 | -2,07 | UDP glucuronosyltransferase 1 family, polypeptide A2 | No | No | - |
| Naa50 | 1,81E-04 | -2,06 | N(alpha)-acetyltransferase 50, NatE catalytic subunit | No | No | - |
| Mast2 | 2,90E-02 | -2,06 | microtubule associated serine/threonine kinase 2 | No | No | - |
| Ski | 5,67E-04 | -2,06 | ski sarcoma viral oncogene homolog (avian) | No | No | 1 |
| Mprip | 2,33E-04 | -2,06 | myosin phosphatase Rho interacting protein | No | No | - |
| Ppp6r1 | 1,02E-02 | -2,06 | protein phosphatase 6, regulatory subunit 1 | No | No | - |
| Junb | 8,16E-04 | -2,06 | jun B proto-oncogene | Yes | No | 1 |
| Rxra | 2,77E-02 | -2,06 | retinoid X receptor alpha | No | No | - |

| | | | | | | |
|---|---|---|---|---|---|---|
| Gm6222 | 4,86E-03 | -2,05 | predicted pseudogene 6222 | No | No | - |
| Fuca1 | 1,17E-03 | -2,05 | fucosidase, alpha-L- 1, tissue | No | No | - |
| Gak | 4,67E-03 | -2,05 | cyclin G associated kinase | No | No | - |
| Abcf1 | 6,30E-03 | -2,05 | ATP-binding cassette, sub-family F (GCN20), member 1 | No | No | - |
| Ugt1a1 | 1,33E-03 | -2,05 | UDP glucuronosyltransferase 1 family, polypeptide A1 | No | No | - |
| Birc6 | 1,71E-02 | -2,05 | baculoviral IAP repeat-containing 6 | No | No | - |
| Kif1c | 1,54E-03 | -2,05 | kinesin family member 1C | No | No | - |
| Mfn2 | 8,88E-03 | -2,05 | mitofusin 2 | No | No | 4 |
| Sbno1 | 2,89E-02 | -2,05 | strawberry notch 1 | No | No | 2 |
| Mvp | 2,97E-02 | -2,05 | major vault protein | Yes | No | 8 |
| Hspa8 | 4,99E-06 | -2,05 | heat shock protein 8 | No | No | - |
| Atg2b | 1,99E-02 | -2,05 | autophagy related 2B | No | No | - |
| Mtss2 | 1,54E-02 | -2,05 | MTSS I-BAR domain containing 2 | No | No | - |
| Telo2 | 8,81E-03 | -2,04 | telomere maintenance 2 | No | No | - |
| Asap1 | 2,96E-03 | -2,04 | ArfGAP with SH3 domain, ankyrin repeat and PH domain1 | No | No | - |
| Rfc5 | 1,31E-03 | -2,04 | replication factor C (activator 1) 5 | No | No | - |
| Fads1 | 8,51E-04 | -2,04 | fatty acid desaturase 1 | No | No | - |
| Arhgef12 | 4,66E-03 | -2,04 | Rho guanine nucleotide exchange factor (GEF) 12 | No | No | - |
| Srprb | 4,17E-03 | -2,04 | signal recognition particle receptor, B subunit | No | No | - |
| Samhd1 | 1,37E-03 | -2,04 | SAM domain and HD domain, 1 | No | No | 2 |
| Myof | 5,86E-03 | -2,04 | myoferlin | No | No | - |
| Pank3 | 5,14E-03 | -2,04 | pantothenate kinase 3 | No | No | - |
| Arih1 | 7,63E-04 | -2,04 | ariadne RBR E3 ubiquitin protein ligase 1 | No | No | - |
| Tgoln1 | 1,48E-03 | -2,04 | trans-golgi network protein | No | No | - |
| Vat1 | 5,58E-04 | -2,03 | vesicle amine transport 1 | No | No | - |
| Paf1 | 1,42E-03 | -2,03 | Paf1, RNA polymerase II complex component | No | No | - |
| Prpf40a | 3,18E-03 | -2,03 | pre-mRNA processing factor 40A | No | No | - |
| Srpk2 | 2,79E-03 | -2,03 | serine/arginine-rich protein specific kinase 2 | No | No | - |
| Parg | 5,15E-03 | -2,03 | poly (ADP-ribose) glycohydrolase | No | No | - |
| Hspb8 | 5,54E-03 | -2,03 | heat shock protein 8 | No | No | - |
| Zc3h4 | 1,05E-02 | -2,03 | zinc finger CCCH-type containing 4 | No | No | 29 |
| Tuba1c | 1,46E-04 | -2,03 | tubulin, alpha 1C | No | No | - |
| Palm | 4,77E-02 | -2,03 | paralemmin | No | No | - |
| Cct8 | 9,54E-04 | -2,03 | chaperonin containing Tcp1, subunit 8 (theta) | No | No | - |
| Gm5835 | 2,72E-02 | -2,03 | predicted gene 5835 | No | No | - |
| Nop2 | 1,17E-02 | -2,03 | NOP2 nucleolar protein | No | No | - |
| Clptm1 | 4,27E-03 | -2,02 | cleft lip and palate associated transmembrane protein 1 | No | No | - |
| Nsun4 | 1,15E-02 | -2,02 | NOL1/NOP2/Sun domain family, member 4 | No | No | - |
| Lyl1 | 8,76E-03 | -2,02 | lymphoblastomic leukemia 1 | No | No | - |
| Irf7 | 1,53E-02 | -2,02 | interferon regulatory factor 7 | Yes | No | 2 |
| Zfp775 | 3,69E-02 | -2,02 | zinc finger protein 775 | No | No | - |
| Wipi2 | 1,07E-03 | -2,02 | WD repeat domain, phosphoinositide interacting 2 | No | No | - |
| Hif1an | 9,75E-04 | -2,02 | hypoxia-inducible factor 1, alpha subunit inhibitor | No | No | - |
| Smc6 | 2,97E-02 | -2,02 | structural maintenance of chromosomes 6 | No | No | - |
| Abcc5 | 6,13E-03 | -2,02 | ATP-binding cassette, sub-family C (CFTR/MRP), member | No | No | - |
| Nceh1 | 2,66E-02 | -2,02 | neutral cholesterol ester hydrolase 1 | No | No | - |
| Hspa5 | 1,04E-05 | -2,02 | heat shock protein 5 | Yes | No | - |
| Ccdc71 | 5,22E-03 | -2,02 | coiled-coil domain containing 71 | No | No | - |
| Anapc5 | 2,52E-04 | -2,02 | anaphase-promoting complex subunit 5 | No | No | - |
| Dcbld2 | 2,01E-03 | -2,01 | discoidin, CUB and LCCL domain containing 2 | No | No | - |
| Sipa1l1 | 1,93E-02 | -2,01 | signal-induced proliferation-associated 1 like 1 | No | No | - |
| Vcp | 5,35E-04 | -2,01 | valosin containing protein | No | No | - |
| Myo6 | 1,62E-03 | -2,01 | myosin VI | No | No | - |
| Capns1 | 5,87E-05 | -2,01 | calpain, small subunit 1 | No | No | - |
| Plaa | 6,00E-03 | -2,01 | phospholipase A2, activating protein | No | No | - |
| Epha1 | 5,09E-03 | -2,01 | Eph receptor A1 | No | No | - |
| Ganab | 4,01E-03 | -2,01 | alpha glucosidase 2 alpha neutral subunit | No | No | - |
| Hsd17b7 | 2,20E-03 | -2,01 | hydroxysteroid (17-beta) dehydrogenase 7 | No | No | - |
| Kmt2e | 3,79E-03 | -2,01 | lysine (K)-specific methyltransferase 2E | No | No | - |
| Tmem151a | 2,90E-03 | -2,01 | transmembrane protein 151A | No | No | - |
| Lats2 | 7,29E-04 | -2,00 | large tumor suppressor 2 | No | No | 1 |
| Ier3 | 7,71E-03 | -2,00 | immediate early response 3 | No | No | - |
| Dst | 9,04E-03 | -2,00 | dystonin | No | No | - |
| Elovl6 | 1,57E-02 | -2,00 | ELOVL family member 6, elongation of long chain fatty ac | No | No | - |
| Exoc1 | 4,09E-03 | -2,00 | exocyst complex component 1 | No | No | - |
| Gdf15 | 3,40E-02 | -2,00 | growth differentiation factor 15 | No | No | 29 |
| Mycbp2 | 1,22E-02 | -2,00 | MYC binding protein 2, E3 ubiquitin protein ligase | No | No | - |
| Ptp4a1 | 3,06E-02 | -2,00 | protein tyrosine phosphatase 4a1 | No | No | - |
| Yars | 2,08E-02 | -2,00 | tyrosyl-tRNA synthetase | No | No | - |
| Wdr26 | 5,18E-03 | -2,00 | WD repeat domain 26 | No | No | - |
| Eif3d | 3,85E-05 | -2,00 | eukaryotic translation initiation factor 3, subunit D | No | No | - |
| Mfsd5 | 2,21E-03 | -2,00 | major facilitator superfamily domain containing 5 | No | No | - |
| Zc3h15 | 2,80E-03 | -2,00 | zinc finger CCCH-type containing 15 | No | No | - |

| | | | | | | |
|---|---|---|---|---|---|---|
| B2m | 1,11E-04 | -1,99 | beta-2 microglobulin | Yes | No | - |
| Ugt1a6b | 1,69E-03 | -1,99 | UDP glucuronosyltransferase 1 family, polypeptide A6B | No | No | - |
| Ubap2l | 5,52E-03 | -1,99 | ubiquitin-associated protein 2-like | No | No | - |
| Ipo7 | 3,14E-03 | -1,99 | importin 7 | No | No | - |
| Ube3b | 6,00E-03 | -1,99 | ubiquitin protein ligase E3B | No | No | - |
| Hipk3 | 3,26E-03 | -1,99 | homeodomain interacting protein kinase 3 | No | No | - |
| Ugt1a7c | 1,75E-03 | -1,99 | UDP glucuronosyltransferase 1 family, polypeptide A7C | No | No | - |
| Ugt1a8 | 1,75E-03 | -1,99 | UDP glucuronosyltransferase 1 family, polypeptide A8 | No | No | - |
| Picalm | 1,90E-04 | -1,99 | phosphatidylinositol binding clathrin assembly protein | No | No | - |
| S100a4 | 1,00E-06 | -1,99 | S100 calcium binding protein A4 | No | No | 62 |
| Irgq | 7,00E-03 | -1,99 | immunity-related GTPase family, Q | No | No | - |
| Ap2s1 | 5,85E-03 | -1,99 | adaptor-related protein complex 2, sigma 1 subunit | No | No | - |
| Parp12 | 4,46E-02 | -1,99 | poly (ADP-ribose) polymerase family, member 12 | No | No | - |
| Plekhg3 | 4,09E-03 | -1,99 | pleckstrin homology domain containing, family G (with Rh | No | No | - |
| Ubap2 | 2,62E-02 | -1,99 | ubiquitin-associated protein 2 | No | No | - |
| Spryd3 | 8,69E-03 | -1,98 | SPRY domain containing 3 | No | No | - |
| Scap | 4,46E-03 | -1,98 | SREBF chaperone | Yes | No | - |
| Avl9 | 3,16E-02 | -1,98 | AVL9 cell migration associated | No | No | - |
| Srebf2 | 6,01E-03 | -1,98 | sterol regulatory element binding factor 2 | Yes | Yes | - |
| Nrbp1 | 1,72E-02 | -1,98 | nuclear receptor binding protein 1 | No | No | - |
| Dido1 | 3,15E-02 | -1,98 | death inducer-obliterator 1 | No | No | - |
| Ugt1a6a | 1,84E-03 | -1,98 | UDP glucuronosyltransferase 1 family, polypeptide A6A | No | No | - |
| Sgta | 7,30E-04 | -1,98 | small glutamine-rich tetratricopeptide repeat (TPR)-containi | No | No | - |
| Hnrnpa3 | 4,37E-02 | -1,98 | heterogeneous nuclear ribonucleoprotein A3 | No | No | - |
| Gtf3c6 | 1,52E-02 | -1,98 | general transcription factor IIIC, polypeptide 6, alpha | No | No | - |
| Zc3h7a | 2,15E-02 | -1,98 | zinc finger CCCH type containing 7 A | No | No | - |
| Ugt1a10 | 1,85E-03 | -1,98 | UDP glycosyltransferase 1 family, polypeptide A10 | No | No | - |
| Hspa13 | 1,85E-02 | -1,98 | heat shock protein 70 family, member 13 | No | No | - |
| Cavin1 | 1,86E-04 | -1,98 | caveolae associated 1 | No | No | - |
| Clic1 | 4,89E-03 | -1,98 | chloride intracellular channel 1 | No | No | - |
| Ddx49 | 1,90E-02 | -1,98 | DEAD box helicase 49 | No | No | - |
| Ugt1a5 | 1,85E-03 | -1,98 | UDP glucuronosyltransferase 1 family, polypeptide A5 | No | No | - |
| Lnpk | 1,84E-02 | -1,98 | lunapark, ER junction formation factor | No | No | - |
| Ifitm3 | 1,53E-05 | -1,98 | interferon induced transmembrane protein 3 | No | No | - |
| Fam166a | 3,71E-04 | -1,98 | family with sequence similarity 166, member A | No | No | - |
| Tnpo3 | 5,11E-03 | -1,97 | transportin 3 | No | No | - |
| St3gal6 | 3,16E-03 | -1,97 | ST3 beta-galactoside alpha-2,3-sialyltransferase 6 | No | No | - |
| Cryab | 2,33E-05 | -1,97 | crystallin, alpha B | No | No | - |
| Ugt1a9 | 1,92E-03 | -1,97 | UDP glucuronosyltransferase 1 family, polypeptide A9 | No | No | - |
| Matr3 | 1,05E-02 | -1,97 | matrin 3 | No | No | - |
| Nmt1 | 4,54E-03 | -1,97 | N-myristoyltransferase 1 | No | No | - |
| Tcerg1 | 2,44E-02 | -1,97 | transcription elongation regulator 1 (CA150) | No | No | - |
| Ap2a2 | 1,97E-03 | -1,97 | adaptor-related protein complex 2, alpha 2 subunit | No | No | - |
| Rora | 3,35E-02 | -1,97 | RAR-related orphan receptor alpha | No | No | 1 |
| Ptgfrn | 1,56E-02 | -1,97 | prostaglandin F2 receptor negative regulator | No | No | 2 |
| Arhgap42 | 2,71E-02 | -1,97 | Rho GTPase activating protein 42 | No | No | 8 |
| Osbpl9 | 1,83E-03 | -1,96 | oxysterol binding protein-like 9 | No | No | - |
| Trub2 | 1,45E-02 | -1,96 | TruB pseudouridine (psi) synthase family member 2 | No | No | - |
| Dnajc7 | 4,05E-02 | -1,96 | DnaJ heat shock protein family (Hsp40) member C7 | No | No | - |
| Tbc1d15 | 3,52E-02 | -1,96 | TBC1 domain family, member 15 | No | No | - |
| Eif3a | 3,02E-03 | -1,96 | eukaryotic translation initiation factor 3, subunit A | No | No | 9 |
| Pde7a | 4,59E-03 | -1,96 | phosphodiesterase 7A | No | No | - |
| Uri1 | 6,66E-03 | -1,96 | URI1, prefoldin-like chaperone | No | No | - |
| Snd1 | 2,63E-03 | -1,96 | staphylococcal nuclease and tudor domain containing 1 | Yes | No | - |
| Ilvbl | 1,03E-03 | -1,96 | ilvB (bacterial acetolactate synthase)-like | No | No | - |
| Mvk | 1,52E-03 | -1,96 | mevalonate kinase | No | No | 2 |
| Cars | 4,28E-03 | -1,96 | cysteinyl-tRNA synthetase | No | No | - |
| Plod3 | 1,34E-04 | -1,96 | procollagen-lysine, 2-oxoglutarate 5-dioxygenase 3 | No | No | 4 |
| Ncor1 | 6,96E-03 | -1,95 | nuclear receptor co-repressor 1 | No | No | - |
| Lrp10 | 6,73E-03 | -1,95 | low-density lipoprotein receptor-related protein 10 | No | No | - |
| Neo1 | 8,69E-03 | -1,95 | neogenin | No | No | - |
| Psmc6 | 2,44E-03 | -1,95 | proteasome (prosome, macropain) 26S subunit, ATPase, 6 | No | No | - |
| Slc39a6 | 5,40E-03 | -1,95 | solute carrier family 39 (metal ion transporter), member 6 | No | No | - |
| Fam50a | 6,90E-03 | -1,95 | family with sequence similarity 50, member A | No | No | - |
| Mindy2 | 1,13E-02 | -1,95 | MINDY lysine 48 deubiquitinase 2 | No | No | - |
| Specc1l | 6,07E-03 | -1,95 | sperm antigen with calponin homology and coiled-coil dom | No | No | - |
| Snrpd3 | 1,57E-02 | -1,95 | small nuclear ribonucleoprotein D3 | No | No | - |
| Vegfb | 1,24E-02 | -1,95 | vascular endothelial growth factor B | No | No | - |
| Zcchc7 | 3,72E-02 | -1,95 | zinc finger, CCHC domain containing 7 | No | No | - |
| Mn1 | 4,93E-02 | -1,95 | meningioma 1 | No | No | - |
| Dhx38 | 1,88E-02 | -1,95 | DEAH (Asp-Glu-Ala-His) box polypeptide 38 | No | No | - |
| Rragc | 2,62E-03 | -1,94 | Ras-related GTP binding C | No | No | - |
| Ptprs | 6,78E-03 | -1,94 | protein tyrosine phosphatase, receptor type, S | No | No | - |

| | | | | | | |
|---|---|---|---|---|---|---|
| Wdr43 | 2,98E-02 | -1,94 | WD repeat domain 43 | No | No | - |
| Cd3eap | 3,78E-03 | -1,94 | CD3E antigen, epsilon polypeptide associated protein | No | No | - |
| Pde2a | 3,44E-02 | -1,94 | phosphodiesterase 2A, cGMP-stimulated | No | No | - |
| Rai14 | 2,85E-02 | -1,94 | retinoic acid induced 14 | No | No | - |
| Gpr108 | 4,41E-03 | -1,94 | G protein-coupled receptor 108 | No | No | - |
| Borcs6 | 7,20E-03 | -1,94 | BLOC-1 related complex subunit 6 | No | No | - |
| Ptpn1 | 3,60E-03 | -1,94 | protein tyrosine phosphatase, non-receptor type 1 | No | No | - |
| Adgrl1 | 1,01E-02 | -1,94 | adhesion G protein-coupled receptor L1 | No | No | - |
| Myo1c | 6,17E-03 | -1,94 | myosin IC | No | No | - |
| Mical2 | 5,98E-03 | -1,94 | microtubule associated monooxygenase, calponin and LIM | No | No | - |
| Ap1g1 | 2,31E-03 | -1,93 | adaptor protein complex AP-1, gamma 1 subunit | No | No | - |
| Piezo1 | 4,46E-03 | -1,93 | piezo-type mechanosensitive ion channel component 1 | No | No | 11 |
| Anapc4 | 3,39E-03 | -1,93 | anaphase promoting complex subunit 4 | No | No | - |
| Vps35 | 9,07E-03 | -1,93 | VPS35 retromer complex component | No | No | - |
| Ctu2 | 6,91E-03 | -1,93 | cytosolic thiouridylase subunit 2 | No | No | - |
| Cul4b | 1,21E-02 | -1,93 | cullin 4B | No | No | - |
| Gars | 2,01E-03 | -1,93 | glycyl-tRNA synthetase | No | No | - |
| Tead1 | 3,78E-03 | -1,93 | TEA domain family member 1 | No | No | - |
| 1110008P14Rik | 2,64E-02 | -1,93 | bublin coiled coil protein | No | No | - |
| Psmd3 | 4,33E-03 | -1,93 | proteasome (prosome, macropain) 26S subunit, non-ATPase | No | No | - |
| Prnp | 1,45E-03 | -1,93 | prion protein | No | No | - |
| Trim25 | 3,03E-02 | -1,93 | tripartite motif-containing 25 | Yes | No | - |
| Gm45837 | 3,59E-02 | -1,93 | predicted gene 45837 | No | No | - |
| Dpagt1 | 6,41E-03 | -1,93 | dolichyl-phosphate (UDP-N-acetylglucosamine) acetylgluc | No | No | - |
| Nod1 | 7,97E-03 | -1,92 | nucleotide-binding oligomerization domain containing 1 | No | No | 4 |
| Etf1 | 4,44E-03 | -1,92 | eukaryotic translation termination factor 1 | No | No | - |
| Cpsf1 | 2,56E-02 | -1,92 | cleavage and polyadenylation specific factor 1 | No | No | - |
| Nab1 | 8,51E-03 | -1,92 | Ngfi-A binding protein 1 | No | No | - |
| Oga | 1,79E-02 | -1,92 | O-GlcNAcase | No | No | - |
| Dag1 | 8,99E-03 | -1,92 | dystroglycan 1 | No | No | - |
| Vax2 | 1,55E-02 | -1,92 | ventral anterior homeobox 2 | No | No | - |
| Cd99l2 | 2,21E-02 | -1,92 | CD99 antigen-like 2 | No | No | - |
| 2200002D01Rik | 7,02E-03 | -1,92 | RIKEN cDNA 2200002D01 gene | No | No | - |
| Fam160b1 | 3,31E-02 | -1,92 | FHF complex subunit HOOK interacting protein 2A | No | No | - |
| Tor3a | 2,88E-02 | -1,92 | torsin family 3, member A | No | No | - |
| Gstt1 | 4,48E-02 | -1,92 | glutathione S-transferase, theta 1 | No | No | - |
| Tns2 | 1,40E-02 | -1,92 | tensin 2 | No | No | - |
| Dcaf7 | 2,10E-02 | -1,92 | DDB1 and CUL4 associated factor 7 | No | No | - |
| Edc3 | 4,56E-03 | -1,92 | enhancer of mRNA decapping 3 | No | No | - |
| Vcl | 1,51E-02 | -1,91 | vinculin | No | No | 3 |
| Dnajc5 | 4,01E-03 | -1,91 | DnaJ heat shock protein family (Hsp40) member C5 | No | No | - |
| Colgalt1 | 3,76E-04 | -1,91 | collagen beta(1-O)galactosyltransferase 1 | No | No | - |
| Raly | 3,18E-02 | -1,91 | hnRNP-associated with lethal yellow | No | No | - |
| Psme4 | 1,60E-02 | -1,91 | proteasome (prosome, macropain) activator subunit 4 | No | No | - |
| Kmt2c | 4,42E-02 | -1,91 | lysine (K)-specific methyltransferase 2C | No | No | - |
| Uqcrc1 | 8,45E-03 | -1,91 | ubiquinol-cytochrome c reductase core protein 1 | No | No | - |
| Rgl1 | 2,41E-02 | -1,91 | ral guanine nucleotide dissociation stimulator,-like 1 | No | No | - |
| Agl | 3,95E-02 | -1,91 | amylo-1,6-glucosidase, 4-alpha-glucanotransferase | No | No | - |
| Csk | 1,53E-02 | -1,91 | c-src tyrosine kinase | No | No | 2 |
| Hspa4 | 9,53E-04 | -1,90 | heat shock protein 4 | Yes | No | - |
| Rpn1 | 1,19E-04 | -1,90 | ribophorin I | No | No | - |
| Tecr | 2,68E-03 | -1,90 | trans-2,3-enoyl-CoA reductase | No | No | - |
| Eif5 | 1,47E-03 | -1,90 | eukaryotic translation initiation factor 5 | No | No | - |
| Birc2 | 2,52E-02 | -1,90 | baculoviral IAP repeat-containing 2 | No | No | - |
| Mgat1 | 1,24E-03 | -1,90 | mannoside acetylglucosaminyltransferase 1 | No | No | - |
| Gpaa1 | 8,57E-03 | -1,90 | GPI anchor attachment protein 1 | No | No | - |
| Amd2 | 3,58E-02 | -1,90 | S-adenosylmethionine decarboxylase 2 | No | No | - |
| Ldlrap1 | 4,04E-03 | -1,90 | low density lipoprotein receptor adaptor protein 1 | No | No | - |
| Gm13680 | 6,13E-03 | -1,90 | predicted gene 13680 | No | No | - |
| Klhl21 | 4,42E-02 | -1,90 | kelch-like 21 | No | No | - |
| Pum2 | 6,29E-03 | -1,90 | pumilio RNA-binding family member 2 | No | No | - |
| Atp6v1c1 | 5,37E-03 | -1,89 | ATPase, H+ transporting, lysosomal V1 subunit C1 | No | No | - |
| Cat | 1,87E-02 | -1,89 | catalase | Yes | No | 142 |
| Tanc1 | 5,50E-03 | -1,89 | tetratricopeptide repeat, ankyrin repeat and coiled-coil conta | No | No | - |
| Tspan2 | 3,83E-02 | -1,89 | tetraspanin 2 | No | No | - |
| Slc4a2 | 9,37E-03 | -1,89 | solute carrier family 4 (anion exchanger), member 2 | No | No | - |
| N4bp1 | 1,59E-02 | -1,89 | NEDD4 binding protein 1 | No | No | - |
| Sptan1 | 6,17E-03 | -1,89 | spectrin alpha, non-erythrocytic 1 | No | No | - |
| Cygb | 2,95E-02 | -1,89 | cytoglobin | No | No | - |
| Dhx29 | 2,58E-02 | -1,89 | DEAH (Asp-Glu-Ala-His) box polypeptide 29 | No | No | - |
| Unc13a | 1,76E-03 | -1,89 | unc-13 homolog A | No | No | - |
| Washc2 | 4,67E-02 | -1,89 | WASH complex subunit 2 | No | No | - |
| Maco1 | 5,13E-03 | -1,89 | macoilin 1 | No | No | - |

| | | | | | | |
|---|---|---|---|---|---|---|
| Purb | 1,24E-02 | -1,89 | purine rich element binding protein B | No | No | - |
| Phf3 | 2,90E-02 | -1,89 | PHD finger protein 3 | No | No | - |
| Syngr2 | 4,24E-03 | -1,89 | synaptogyrin 2 | No | No | - |
| Psmf1 | 3,62E-02 | -1,89 | proteasome (prosome, macropain) inhibitor subunit 1 | No | No | - |
| Mak16 | 3,62E-02 | -1,89 | MAK16 homolog | No | No | - |
| Hgsnat | 4,89E-02 | -1,88 | heparan-alpha-glucosaminide N-acetyltransferase | No | No | - |
| Zfp768 | 2,18E-02 | -1,88 | zinc finger protein 768 | No | No | - |
| Atp6v1b2 | 6,52E-04 | -1,88 | ATPase, H+ transporting, lysosomal V1 subunit B2 | No | No | 1 |
| Rps4x-ps | 3,90E-03 | -1,88 | Rps4x retrotransposed pseudogene | No | No | - |
| Ndufb7 | 3,92E-03 | -1,88 | NADH:ubiquinone oxidoreductase subunit B7 | No | No | - |
| Sox4 | 3,06E-03 | -1,88 | SRY (sex determining region Y)-box 4 | Yes | No | 5 |
| Prpsap1 | 1,17E-02 | -1,88 | phosphoribosyl pyrophosphate synthetase-associated protei | No | No | - |
| Ptp4a1 | 4,42E-02 | -1,88 | protein tyrosine phosphatase 4a1 | No | No | - |
| Krit1 | 3,80E-02 | -1,88 | KRIT1, ankyrin repeat containing | No | No | 2 |
| Actr1a | 1,17E-02 | -1,88 | ARP1 actin-related protein 1A, centractin alpha | No | No | - |
| Tsr1 | 4,98E-03 | -1,88 | TSR1 20S rRNA accumulation | No | No | - |
| Ftsj3 | 8,05E-03 | -1,88 | FtsJ RNA methyltransferase homolog 3 (E. coli) | No | No | - |
| Mbtps1 | 2,17E-03 | -1,88 | membrane-bound transcription factor peptidase, site 1 | No | No | - |
| Arrb1 | 1,37E-02 | -1,88 | arrestin, beta 1 | No | No | - |
| Ccl7 | 6,94E-03 | -1,88 | chemokine (C-C motif) ligand 7 | No | No | - |
| Ddx54 | 3,77E-02 | -1,88 | DEAD box helicase 54 | No | No | - |
| Klhl13 | 2,49E-02 | -1,87 | kelch-like 13 | No | No | - |
| Pxdn | 1,99E-02 | -1,87 | peroxidasin | No | No | - |
| Ifitm2 | 9,08E-03 | -1,87 | interferon induced transmembrane protein 2 | No | No | - |
| Gmppb | 4,40E-03 | -1,87 | GDP-mannose pyrophosphorylase B | No | No | - |
| Eif4g3 | 5,09E-03 | -1,87 | eukaryotic translation initiation factor 4 gamma, 3 | No | No | - |
| Prss57 | 8,76E-03 | -1,87 | protease, serine 57 | No | No | - |
| Fam210b | 8,04E-03 | -1,87 | family with sequence similarity 210, member B | No | No | - |
| Adipor2 | 6,37E-03 | -1,87 | adiponectin receptor 2 | No | No | - |
| Rsrc2 | 6,31E-03 | -1,87 | arginine/serine-rich coiled-coil 2 | No | No | - |
| Heca | 3,23E-02 | -1,86 | hdc homolog, cell cycle regulator | No | No | - |
| Syde1 | 2,68E-03 | -1,86 | synapse defective 1, Rho GTPase, homolog 1 (C. elegans) | No | No | - |
| Fech | 4,17E-02 | -1,86 | ferrochelatase | No | No | - |
| Dusp3 | 3,09E-03 | -1,86 | dual specificity phosphatase 3 (vaccinia virus phosphatase V | No | No | - |
| Vps33a | 2,02E-02 | -1,86 | VPS33A CORVET/HOPS core subunit | No | No | - |
| Fam120a | 5,16E-03 | -1,86 | family with sequence similarity 120, member A | No | No | - |
| Zyx | 9,58E-03 | -1,86 | zyxin | No | No | 1 |
| Ankrd1 | 4,46E-04 | -1,86 | ankyrin repeat domain 1 (cardiac muscle) | No | No | - |
| Whamm | 3,81E-02 | -1,86 | WAS protein homolog associated with actin, golgi membrai | No | No | - |
| Mrtfa | 8,21E-03 | -1,86 | myocardin related transcription factor A | No | No | - |
| Arhgef40 | 5,83E-03 | -1,85 | Rho guanine nucleotide exchange factor (GEF) 40 | No | No | - |
| Ctps | 3,06E-03 | -1,85 | cytidine 5'-triphosphate synthase | No | No | - |
| Gpc4 | 1,29E-02 | -1,85 | glypican 4 | No | No | - |
| Ap1b1 | 1,92E-02 | -1,85 | adaptor protein complex AP-1, beta 1 subunit | No | No | - |
| Rnf123 | 4,35E-03 | -1,85 | ring finger protein 123 | No | No | - |
| Mob4 | 2,77E-03 | -1,85 | MOB family member 4, phocein | No | No | - |
| Nomo1 | 1,06E-02 | -1,85 | nodal modulator 1 | No | No | - |
| Eif2s2 | 4,64E-03 | -1,85 | eukaryotic translation initiation factor 2, subunit 2 (beta) | No | No | - |
| Gm8960 | 1,38E-02 | -1,85 | predicted gene 8960 | No | No | - |
| Lmcd1 | 3,92E-02 | -1,85 | LIM and cysteine-rich domains 1 | No | No | - |
| Hp1bp3 | 1,81E-04 | -1,84 | heterochromatin protein 1, binding protein 3 | No | No | - |
| Trmt2a | 1,83E-02 | -1,84 | TRM2 tRNA methyltransferase 2A | No | No | - |
| Acta2 | 4,99E-03 | -1,84 | actin alpha 2, smooth muscle, aorta | No | No | 15 |
| Smarca4 | 2,01E-02 | -1,84 | SWI/SNF related, matrix associated, actin dependent regula | Yes | No | 6 |
| Arfgap1 | 7,52E-03 | -1,84 | ADP-ribosylation factor GTPase activating protein 1 | No | No | - |
| Arhgef2 | 1,50E-02 | -1,84 | rho/rac guanine nucleotide exchange factor (GEF) 2 | No | No | - |
| Pa2g4 | 8,19E-03 | -1,84 | proliferation-associated 2G4 | No | No | - |
| Rnf20 | 1,52E-02 | -1,84 | ring finger protein 20 | No | No | - |
| Ogfr | 1,53E-02 | -1,84 | opioid growth factor receptor | No | No | - |
| Stip1 | 5,24E-03 | -1,84 | stress-induced phosphoprotein 1 | No | No | - |
| Erlin2 | 3,53E-02 | -1,84 | ER lipid raft associated 2 | No | No | - |
| Bag3 | 1,63E-02 | -1,84 | BCL2-associated athanogene 3 | No | No | - |
| Sulf2 | 3,35E-02 | -1,84 | sulfatase 2 | No | No | - |
| Afap1 | 8,48E-03 | -1,84 | actin filament associated protein 1 | No | No | - |
| Tesk1 | 7,60E-03 | -1,84 | testis specific protein kinase 1 | Yes | No | - |
| Rlim | 5,27E-03 | -1,84 | ring finger protein, LIM domain interacting | No | No | - |
| Rbck1 | 3,06E-02 | -1,84 | RanBP-type and C3HC4-type zinc finger containing 1 | No | No | - |
| Tm9sf4 | 1,67E-02 | -1,83 | transmembrane 9 superfamily member 4 | No | No | - |
| Dnajc18 | 3,34E-02 | -1,83 | DnaJ heat shock protein family (Hsp40) member C18 | No | No | - |
| Uvrag | 3,35E-02 | -1,83 | UV radiation resistance associated gene | No | No | - |
| Tmem259 | 2,83E-02 | -1,83 | transmembrane protein 259 | No | No | - |
| Col16a1 | 5,11E-03 | -1,83 | collagen, type XVI, alpha 1 | No | No | - |
| Tsn | 1,10E-04 | -1,83 | translin | No | No | - |

| | | | | | | |
|---|---|---|---|---|---|---|
| Serinc1 | 8,48E-03 | -1,83 | serine incorporator 1 | No | No | - |
| Myo5a | 2,16E-02 | -1,83 | myosin VA | No | No | - |
| Dnajc3 | 9,24E-03 | -1,82 | DnaJ heat shock protein family (Hsp40) member C3 | No | No | - |
| Ikbkb | 1,85E-02 | -1,82 | inhibitor of kappaB kinase beta | No | No | - |
| Spag9 | 1,25E-02 | -1,82 | sperm associated antigen 9 | No | No | - |
| Cd72 | 8,79E-03 | -1,82 | CD72 antigen | No | No | - |
| Cxcl14 | 7,66E-03 | -1,82 | chemokine (C-X-C motif) ligand 14 | No | No | 28 |
| Actb | 4,44E-04 | -1,82 | actin, beta | No | No | - |
| Tbx5 | 5,84E-03 | -1,82 | T-box 5 | No | No | 2 |
| Tubb4b | 1,30E-04 | -1,82 | tubulin, beta 4B class IVB | No | No | - |
| Fstl3 | 1,31E-02 | -1,82 | follistatin-like 3 | No | No | - |
| Itgav | 1,43E-02 | -1,82 | integrin alpha V | No | No | 3 |
| Gatad2a | 9,97E-03 | -1,82 | GATA zinc finger domain containing 2A | No | No | - |
| Cul7 | 2,60E-02 | -1,82 | cullin 7 | No | No | - |
| Pole3 | 1,10E-02 | -1,81 | polymerase (DNA directed), epsilon 3 (p17 subunit) | No | No | - |
| Kpna4 | 2,28E-02 | -1,81 | karyopherin (importin) alpha 4 | No | No | - |
| Cnppd1 | 1,08E-02 | -1,81 | cyclin Pas1/PHO80 domain containing 1 | No | No | - |
| Ehbp1l1 | 8,77E-03 | -1,81 | EH domain binding protein 1-like 1 | No | No | - |
| Anxa3 | 1,36E-04 | -1,81 | annexin A3 | No | No | 1 |
| Naf1 | 1,69E-02 | -1,81 | nuclear assembly factor 1 ribonucleoprotein | No | No | 11 |
| Pcolce | 6,17E-03 | -1,81 | procollagen C-endopeptidase enhancer protein | No | No | - |
| P3h1 | 7,82E-03 | -1,81 | prolyl 3-hydroxylase 1 | No | No | - |
| Klhdc4 | 1,72E-02 | -1,81 | kelch domain containing 4 | No | No | - |
| Med13l | 3,99E-02 | -1,81 | mediator complex subunit 13-like | No | No | - |
| Mtdh | 2,40E-02 | -1,81 | metadherin | No | No | - |
| Ggcx | 3,17E-02 | -1,81 | gamma-glutamyl carboxylase | No | No | - |
| Snx33 | 4,08E-02 | -1,81 | sorting nexin 33 | No | No | - |
| Ubr2 | 3,89E-02 | -1,81 | ubiquitin protein ligase E3 component n-recognin 2 | No | No | - |
| Zmat2 | 3,07E-02 | -1,81 | zinc finger, matrin type 2 | No | No | - |
| Gm50241 | 1,55E-02 | -1,80 | DERPC proline and glycine rich nuclear protein | No | No | - |
| Atp2a2 | 8,94E-03 | -1,80 | ATPase, Ca++ transporting, cardiac muscle, slow twitch 2 | No | No | - |
| Ankrd50 | 4,14E-03 | -1,80 | ankyrin repeat domain 50 | No | No | - |
| Capn2 | 4,27E-03 | -1,80 | calpain 2 | No | No | - |
| Clcn4 | 2,05E-02 | -1,80 | chloride channel, voltage-sensitive 4 | No | No | - |
| Chtf8 | 1,59E-02 | -1,80 | CTF8, chromosome transmission fidelity factor 8 | No | No | - |
| Grin3b | 3,80E-02 | -1,80 | glutamate receptor, ionotropic, NMDA3B | No | No | - |
| Brox | 2,77E-02 | -1,80 | BRO1 domain and CAAX motif containing | No | No | - |
| Pi4k2a | 2,50E-02 | -1,80 | phosphatidylinositol 4-kinase type 2 alpha | No | No | - |
| Jund | 2,02E-02 | -1,80 | jun D proto-oncogene | No | Yes | - |
| Pigs | 3,55E-02 | -1,80 | phosphatidylinositol glycan anchor biosynthesis, class S | No | No | - |
| Tnpo1 | 3,79E-03 | -1,80 | transportin 1 | No | No | - |
| Sgsm3 | 1,02E-02 | -1,80 | small G protein signaling modulator 3 | No | No | - |
| Ddi2 | 1,14E-02 | -1,79 | DNA-damage inducible protein 2 | No | No | - |
| Rftn2 | 3,37E-02 | -1,79 | raftlin family member 2 | No | No | - |
| Zdhhc9 | 2,21E-02 | -1,79 | zinc finger, DHHC domain containing 9 | No | No | - |
| Nufip2 | 2,59E-02 | -1,79 | nuclear fragile X mental retardation protein interacting prot | No | No | - |
| Cast | 1,67E-02 | -1,79 | calpastatin | No | No | - |
| Synj2 | 3,67E-02 | -1,79 | synaptojanin 2 | No | No | - |
| Letm1 | 3,23E-02 | -1,79 | leucine zipper-EF-hand containing transmembrane protein 1 | No | No | - |
| Eif5b | 4,58E-02 | -1,79 | eukaryotic translation initiation factor 5B | No | No | - |
| Txlna | 3,23E-02 | -1,79 | taxilin alpha | No | No | - |
| Ptges3 | 9,34E-03 | -1,79 | prostaglandin E synthase 3 | No | No | - |
| Ap3m1 | 2,04E-02 | -1,79 | adaptor-related protein complex 3, mu 1 subunit | No | No | - |
| Acaa1a | 1,08E-02 | -1,79 | acetyl-Coenzyme A acyltransferase 1A | No | No | - |
| Hadha | 1,88E-02 | -1,78 | hydroxyacyl-CoA dehydrogenase trifunctional multienzyme | No | No | - |
| Rnf185 | 3,44E-02 | -1,78 | ring finger protein 185 | No | No | - |
| Plcg1 | 2,71E-02 | -1,78 | phospholipase C, gamma 1 | No | No | - |
| Zdhhc16 | 3,50E-02 | -1,78 | zinc finger, DHHC domain containing 16 | No | No | - |
| Creg1 | 3,37E-02 | -1,78 | cellular repressor of E1A-stimulated genes 1 | No | No | - |
| Large1 | 2,31E-02 | -1,78 | LARGE xylosyl- and glucuronyltransferase 1 | No | No | - |
| Cops5 | 9,34E-03 | -1,78 | COP9 signalosome subunit 5 | No | No | - |
| Olfr77 | 3,92E-02 | -1,78 | olfactory receptor 77 | No | No | - |
| Mras | 2,41E-02 | -1,78 | muscle and microspikes RAS | No | No | - |
| Gm15500 | 1,17E-02 | -1,78 | predicted pseudogene 15500 | No | No | - |
| Zc2hc1c | 2,28E-02 | -1,77 | zinc finger, C2HC-type containing 1C | No | No | - |
| Blmh | 3,95E-02 | -1,77 | bleomycin hydrolase | No | No | 11 |
| Arhgef17 | 8,31E-03 | -1,77 | Rho guanine nucleotide exchange factor (GEF) 17 | No | No | - |
| Clk3 | 1,48E-02 | -1,77 | CDC-like kinase 3 | No | No | 1 |
| Parp14 | 2,71E-02 | -1,77 | poly (ADP-ribose) polymerase family, member 14 | Yes | No | - |
| Nfib | 3,87E-02 | -1,77 | nuclear factor I/B | No | No | - |
| Paqr7 | 2,70E-02 | -1,77 | progestin and adipoQ receptor family member VII | No | No | - |
| Scd2 | 1,62E-02 | -1,77 | stearoyl-Coenzyme A desaturase 2 | Yes | No | 54 |
| Reep5 | 8,69E-03 | -1,76 | receptor accessory protein 5 | No | No | - |

| | | | | | | |
|---|---|---|---|---|---|---|
| Zfp513 | 1,90E-02 | -1,76 | zinc finger protein 513 | No | No | - |
| Ptpn23 | 1,71E-02 | -1,76 | protein tyrosine phosphatase, non-receptor type 23 | No | No | - |
| Cul1 | 3,82E-02 | -1,76 | cullin 1 | No | No | - |
| Fastk | 2,02E-02 | -1,75 | Fas-activated serine/threonine kinase | No | No | - |
| Cald1 | 1,45E-02 | -1,75 | caldesmon 1 | No | No | - |
| Dyrk1a | 3,66E-02 | -1,75 | dual-specificity tyrosine-(Y)-phosphorylation regulated kina | No | No | - |
| Bag1 | 1,07E-02 | -1,75 | BCL2-associated athanogene 1 | No | No | - |
| Trmt1 | 2,56E-02 | -1,75 | tRNA methyltransferase 1 | No | No | - |
| Hsp90b1 | 3,57E-03 | -1,75 | heat shock protein 90, beta (Grp94), member 1 | No | No | - |
| Plekha1 | 3,81E-02 | -1,75 | pleckstrin homology domain containing, family A (phospho | No | No | - |
| Bicc1 | 7,03E-03 | -1,75 | BicC family RNA binding protein 1 | No | No | - |
| Ddx42 | 1,56E-02 | -1,75 | DEAD box helicase 42 | No | No | - |
| Actn1 | 4,56E-03 | -1,75 | actinin, alpha 1 | No | No | - |
| Anxa9 | 3,53E-02 | -1,74 | annexin A9 | No | No | - |
| Mfhas1 | 3,13E-02 | -1,74 | malignant fibrous histiocytoma amplified sequence 1 | No | No | - |
| Prmt5 | 2,11E-02 | -1,74 | protein arginine N-methyltransferase 5 | No | No | - |
| Zfp512 | 1,54E-02 | -1,74 | zinc finger protein 512 | No | No | - |
| Ubp1 | 1,93E-02 | -1,74 | upstream binding protein 1 | No | No | - |
| Gng12 | 6,80E-04 | -1,74 | guanine nucleotide binding protein (G protein), gamma 12 | No | No | - |
| Add1 | 2,02E-02 | -1,74 | adducin 1 (alpha) | No | No | - |
| Oaz2 | 6,87E-03 | -1,74 | ornithine decarboxylase antizyme 2 | No | No | - |
| Rex1bd | 8,66E-03 | -1,74 | required for excision 1-B domain containing | No | No | - |
| Foxp1 | 1,83E-02 | -1,74 | forkhead box P1 | No | No | 1 |
| Snx17 | 2,16E-02 | -1,74 | sorting nexin 17 | No | No | - |
| Slc29a3 | 8,69E-03 | -1,74 | solute carrier family 29 (nucleoside transporters), member 3 | No | No | - |
| Prr13 | 9,42E-03 | -1,74 | proline rich 13 | No | No | - |
| Rab3gap1 | 3,35E-02 | -1,73 | RAB3 GTPase activating protein subunit 1 | No | No | - |
| Ubb-ps | 1,01E-02 | -1,73 | ubiquitin B, pseudogene | No | No | - |
| Eif3f | 2,09E-03 | -1,73 | eukaryotic translation initiation factor 3, subunit F | No | No | - |
| Map4k4 | 6,70E-03 | -1,73 | mitogen-activated protein kinase kinase kinase kinase 4 | No | No | - |
| Spsb1 | 3,00E-02 | -1,73 | splA/ryanodine receptor domain and SOCS box containing | No | No | - |
| Slc27a4 | 3,80E-02 | -1,73 | solute carrier family 27 (fatty acid transporter), member 4 | No | No | - |
| Fto | 1,71E-02 | -1,73 | fat mass and obesity associated | No | No | - |
| Tamm41 | 4,86E-02 | -1,73 | TAM41 mitochondrial translocator assembly and maintenar | No | No | - |
| Osbpl5 | 2,46E-02 | -1,73 | oxysterol binding protein-like 5 | No | No | - |
| Tor1b | 1,34E-02 | -1,73 | torsin family 1, member B | No | No | - |
| H13 | 4,26E-03 | -1,73 | histocompatibility 13 | No | No | - |
| Znhit2 | 4,88E-02 | -1,73 | zinc finger, HIT domain containing 2 | No | No | - |
| Sh3bp5l | 2,23E-02 | -1,73 | SH3 binding domain protein 5 like | No | No | - |
| Tsc22d2 | 2,26E-02 | -1,73 | TSC22 domain family, member 2 | No | No | - |
| Flot2 | 4,02E-02 | -1,73 | flotillin 2 | No | No | - |
| Saraf | 6,04E-03 | -1,72 | store-operated calcium entry-associated regulatory factor | No | No | - |
| Vcam1 | 1,06E-02 | -1,72 | vascular cell adhesion molecule 1 | No | No | 10 |
| Copg1 | 1,69E-02 | -1,72 | coatomer protein complex, subunit gamma 1 | No | No | - |
| Fbxl2 | 2,30E-02 | -1,72 | F-box and leucine-rich repeat protein 2 | No | No | - |
| Vamp5 | 2,92E-02 | -1,72 | vesicle-associated membrane protein 5 | No | No | - |
| Ube3a | 2,16E-02 | -1,72 | ubiquitin protein ligase E3A | No | No | - |
| Slc48a1 | 5,72E-04 | -1,72 | solute carrier family 48 (heme transporter), member 1 | No | No | - |
| Pcdhga1 | 2,72E-02 | -1,72 | protocadherin gamma subfamily A, 1 | No | No | - |
| Gm3571 | 4,26E-02 | -1,71 | predicted gene 3571 | No | No | - |
| Ltbp3 | 1,25E-02 | -1,71 | latent transforming growth factor beta binding protein 3 | No | No | 1 |
| Cand1 | 1,73E-02 | -1,71 | cullin associated and neddylation disassociated 1 | No | No | - |
| Mylk | 1,92E-02 | -1,71 | myosin, light polypeptide kinase | No | No | - |
| Pcdhga9 | 2,76E-02 | -1,71 | protocadherin gamma subfamily A, 9 | No | No | - |
| Hsp25-ps1 | 3,01E-02 | -1,71 | heat shock protein 25, pseudogene 1 | No | No | - |
| Sec16a | 4,33E-02 | -1,71 | SEC16 homolog A, endoplasmic reticulum export factor | No | No | - |
| Dnajb11 | 3,01E-02 | -1,71 | DnaJ heat shock protein family (Hsp40) member B11 | No | No | - |
| Pcdhga8 | 2,82E-02 | -1,71 | protocadherin gamma subfamily A, 8 | No | No | - |
| Pcdhgb6 | 2,79E-02 | -1,71 | protocadherin gamma subfamily B, 6 | No | No | - |
| Mindy1 | 4,05E-02 | -1,71 | MINDY lysine 48 deubiquitinase 1 | No | No | - |
| Yif1b | 2,89E-02 | -1,71 | Yip1 interacting factor homolog B (S. cerevisiae) | No | No | - |
| Kank1 | 3,39E-02 | -1,71 | KN motif and ankyrin repeat domains 1 | No | No | - |
| Pcdhgb7 | 2,81E-02 | -1,71 | protocadherin gamma subfamily B, 7 | No | No | - |
| Pcdhga12 | 2,84E-02 | -1,71 | protocadherin gamma subfamily A, 12 | No | No | - |
| Gba | 5,86E-03 | -1,71 | glucosidase, beta, acid | No | No | 3 |
| Pcdhga11 | 2,84E-02 | -1,71 | protocadherin gamma subfamily A, 11 | No | No | - |
| Pcdhga10 | 2,85E-02 | -1,71 | protocadherin gamma subfamily A, 10 | No | No | - |
| Pcdhga7 | 2,84E-02 | -1,71 | protocadherin gamma subfamily A, 7 | No | No | - |
| Pcdhgb5 | 2,86E-02 | -1,70 | protocadherin gamma subfamily B, 5 | No | No | - |
| Slc1a5 | 3,18E-02 | -1,70 | solute carrier family 1 (neutral amino acid transporter), men | No | No | - |
| Pcdhga6 | 2,88E-02 | -1,70 | protocadherin gamma subfamily A, 6 | No | No | - |
| Cyba | 2,81E-02 | -1,70 | cytochrome b-245, alpha polypeptide | No | No | 7 |
| Eif3j1 | 2,85E-02 | -1,70 | eukaryotic translation initiation factor 3, subunit J1 | No | No | - |

| | | | | | | |
|---|---|---|---|---|---|---|
| Gm21982 | 4,04E-02 | -1,70 | predicted gene 21982 | No | No | - |
| Ccng1 | 6,48E-03 | -1,70 | cyclin G1 | No | No | - |
| Pcdhgb4 | 2,96E-02 | -1,70 | protocadherin gamma subfamily B, 4 | No | No | - |
| Gab2 | 8,44E-03 | -1,70 | growth factor receptor bound protein 2-associated protein 2 | No | No | 1 |
| Pcdhgb8 | 2,93E-02 | -1,70 | protocadherin gamma subfamily B, 8 | No | No | - |
| Arpc1b | 1,10E-02 | -1,70 | actin related protein 2/3 complex, subunit 1B | No | No | - |
| Mtmr4 | 1,98E-02 | -1,70 | myotubularin related protein 4 | No | No | - |
| Nbl1 | 1,69E-02 | -1,70 | NBL1, DAN family BMP antagonist | No | No | 1 |
| Lima1 | 3,23E-02 | -1,70 | LIM domain and actin binding 1 | No | No | - |
| Rpn2 | 7,61E-03 | -1,70 | ribophorin II | No | No | - |
| Pcdhgb2 | 2,93E-02 | -1,70 | protocadherin gamma subfamily B, 2 | No | No | - |
| Tmtc3 | 4,28E-02 | -1,70 | transmembrane and tetratricopeptide repeat containing 3 | No | No | - |
| Septin8 | 2,71E-02 | -1,69 | septin 8 | No | No | - |
| Plxnb2 | 1,63E-02 | -1,69 | plexin B2 | No | No | - |
| Nek9 | 3,80E-02 | -1,69 | NIMA (never in mitosis gene a)-related expressed kinase 9 | No | No | - |
| Pcdhga3 | 2,98E-02 | -1,69 | protocadherin gamma subfamily A, 3 | No | No | - |
| Tom1l2 | 2,13E-02 | -1,69 | target of myb1-like 2 (chicken) | No | No | - |
| Mbd6 | 1,19E-02 | -1,69 | methyl-CpG binding domain protein 6 | No | No | - |
| Pard3 | 2,22E-02 | -1,69 | par-3 family cell polarity regulator | No | No | - |
| Pcdhga2 | 3,04E-02 | -1,69 | protocadherin gamma subfamily A, 2 | No | No | - |
| Pcdhga5 | 3,06E-02 | -1,69 | protocadherin gamma subfamily A, 5 | No | No | - |
| Msmo1 | 9,60E-03 | -1,69 | methylsterol monoxygenase 1 | No | No | - |
| Scyl1 | 1,57E-02 | -1,69 | SCY1-like 1 (S. cerevisiae) | No | No | - |
| Psmc5 | 4,95E-04 | -1,69 | protease (prosome, macropain) 26S subunit, ATPase 5 | No | No | - |
| Por | 1,48E-02 | -1,68 | cytochrome p450 oxidoreductase | No | No | 2 |
| Pcdhgb1 | 3,13E-02 | -1,68 | protocadherin gamma subfamily B, 1 | No | No | - |
| Sod1 | 1,39E-02 | -1,68 | superoxide dismutase 1, soluble | Yes | No | 5 |
| Pcdhga4 | 3,15E-02 | -1,68 | protocadherin gamma subfamily A, 4 | No | No | - |
| Crlf1 | 1,36E-02 | -1,68 | cytokine receptor-like factor 1 | No | No | - |
| Rbm42 | 3,89E-02 | -1,68 | RNA binding motif protein 42 | No | No | 1 |
| Fgf2 | 1,25E-02 | -1,68 | fibroblast growth factor 2 | Yes | No | 62 |
| Slc3a2 | 4,28E-02 | -1,68 | solute carrier family 3 (activators of dibasic and neutral ami | No | No | - |
| Dcn | 5,80E-03 | -1,68 | decorin | No | No | 6 |
| Cep170b | 3,02E-02 | -1,68 | centrosomal protein 170B | No | No | - |
| Nfix | 2,10E-02 | -1,67 | nuclear factor I/X | No | No | - |
| Sdc2 | 3,01E-02 | -1,67 | syndecan 2 | No | No | 21 |
| Bax | 2,70E-02 | -1,67 | BCL2-associated X protein | Yes | No | 22 |
| Dctn2 | 1,13E-02 | -1,67 | dynactin 2 | No | No | - |
| Pip5k1a | 4,05E-02 | -1,67 | phosphatidylinositol-4-phosphate 5-kinase, type 1 alpha | No | No | - |
| Tmf1 | 4,45E-02 | -1,67 | TATA element modulatory factor 1 | No | No | - |
| Cggbp1 | 2,59E-02 | -1,67 | CGG triplet repeat binding protein 1 | No | No | - |
| Gm5586 | 4,14E-02 | -1,67 | predicted gene 5586 | No | No | - |
| Golga2 | 1,30E-02 | -1,67 | golgi autoantigen, golgin subfamily a, 2 | No | No | 3 |
| Mtch1 | 1,10E-02 | -1,67 | mitochondrial carrier 1 | No | No | - |
| Ppm1f | 2,90E-02 | -1,67 | protein phosphatase 1F (PP2C domain containing) | No | No | - |
| Ankrd17 | 2,99E-02 | -1,67 | ankyrin repeat domain 17 | No | No | - |
| Swi5 | 4,53E-02 | -1,66 | SWI5 recombination repair homolog (yeast) | No | No | - |
| Slc15a4 | 2,72E-02 | -1,66 | solute carrier family 15, member 4 | No | No | - |
| Trip12 | 2,30E-02 | -1,66 | thyroid hormone receptor interactor 12 | No | No | - |
| Trak1 | 3,68E-02 | -1,66 | trafficking protein, kinesin binding 1 | No | No | 1 |
| Uhmk1 | 3,53E-02 | -1,66 | U2AF homology motif (UHM) kinase 1 | No | No | - |
| Anxa6 | 2,12E-02 | -1,66 | annexin A6 | No | No | - |
| Dcaf12 | 2,30E-02 | -1,66 | DDB1 and CUL4 associated factor 12 | No | No | - |
| Hgs | 4,42E-02 | -1,65 | HGF-regulated tyrosine kinase substrate | No | No | 8 |
| Phldb2 | 2,18E-02 | -1,65 | pleckstrin homology like domain, family B, member 2 | No | No | - |
| Usf2 | 9,35E-04 | -1,65 | upstream transcription factor 2 | No | Yes | - |
| Ptpra | 4,90E-02 | -1,65 | protein tyrosine phosphatase, receptor type, A | No | No | - |
| Cdon | 3,96E-02 | -1,65 | cell adhesion molecule-related/down-regulated by oncogene | No | No | - |
| Vdac2 | 1,63E-02 | -1,65 | voltage-dependent anion channel 2 | No | No | - |
| Stk16 | 1,64E-02 | -1,65 | serine/threonine kinase 16 | No | No | - |
| Otud5 | 3,29E-02 | -1,64 | OTU domain containing 5 | No | No | - |
| Foxs1 | 3,61E-02 | -1,64 | forkhead box S1 | No | No | - |
| Ptbp3 | 3,58E-02 | -1,64 | polypyrimidine tract binding protein 3 | No | No | - |
| Wdr34 | 3,57E-02 | -1,64 | dynein 2 intermediate chain 2 | No | No | - |
| Unc5b | 2,20E-02 | -1,64 | unc-5 netrin receptor B | No | No | - |
| Tcf25 | 1,74E-02 | -1,64 | transcription factor 25 (basic helix-loop-helix) | No | No | - |
| Ehd1 | 1,14E-02 | -1,64 | EH-domain containing 1 | No | No | - |
| Cav1 | 3,33E-02 | -1,64 | caveolin 1, caveolae protein | Yes | No | 19 |
| Arhgdia | 3,71E-02 | -1,63 | Rho GDP dissociation inhibitor (GDI) alpha | No | No | - |
| Uaca | 4,66E-02 | -1,63 | uveal autoantigen with coiled-coil domains and ankyrin repe | No | No | - |
| Rnh1 | 1,42E-02 | -1,63 | ribonuclease/angiogenin inhibitor 1 | No | No | - |
| Wnk1 | 3,92E-02 | -1,63 | WNK lysine deficient protein kinase 1 | No | No | - |
| Eif4a3 | 2,42E-02 | -1,63 | eukaryotic translation initiation factor 4A3 | No | No | - |

| | | | | | | |
|---|---|---|---|---|---|---|
| Cdc20 | 1,24E-02 | -1,63 | cell division cycle 20 | No | No | 1 |
| Igsf8 | 2,77E-02 | -1,62 | immunoglobulin superfamily, member 8 | No | No | - |
| Rab7 | 1,75E-02 | -1,62 | RAB7, member RAS oncogene family | No | No | - |
| Ubb | 7,52E-04 | -1,62 | ubiquitin B | No | No | 9 |
| Hmgcr | 3,38E-02 | -1,62 | 3-hydroxy-3-methylglutaryl-Coenzyme A reductase | Yes | No | 4 |
| Mbnl2 | 4,26E-02 | -1,62 | muscleblind like splicing factor 2 | No | No | - |
| Habp4 | 5,00E-02 | -1,62 | hyaluronic acid binding protein 4 | No | No | - |
| Rabgef1 | 4,60E-02 | -1,62 | RAB guanine nucleotide exchange factor (GEF) 1 | No | No | - |
| Tuba4a | 2,04E-02 | -1,62 | tubulin, alpha 4A | No | No | - |
| Ddr2 | 4,43E-02 | -1,61 | discoidin domain receptor family, member 2 | Yes | No | 14 |
| Usp4 | 3,92E-02 | -1,61 | ubiquitin specific peptidase 4 (proto-oncogene) | No | No | - |
| Mtx1 | 3,97E-02 | -1,61 | metaxin 1 | No | No | - |
| Slc2a10 | 3,44E-02 | -1,61 | solute carrier family 2 (facilitated glucose transporter), mem | No | No | - |
| Tnrc6a | 2,77E-02 | -1,61 | trinucleotide repeat containing 6a | No | No | - |
| Vamp3 | 1,99E-02 | -1,61 | vesicle-associated membrane protein 3 | No | No | - |
| Fnbp1 | 3,94E-02 | -1,61 | formin binding protein 1 | No | No | - |
| Rhoc | 2,06E-02 | -1,60 | ras homolog family member C | No | No | 1 |
| Dhx15 | 3,45E-02 | -1,60 | DEAH (Asp-Glu-Ala-His) box polypeptide 15 | Yes | No | - |
| Pip4k2b | 4,69E-02 | -1,60 | phosphatidylinositol-5-phosphate 4-kinase, type II, beta | No | No | - |
| Atf4 | 3,90E-03 | -1,60 | activating transcription factor 4 | No | Yes | 12 |
| Rhoq | 1,71E-02 | -1,60 | ras homolog family member Q | No | No | - |
| Nrd1 | 2,97E-02 | -1,60 | nardilysin, N-arginine dibasic convertase, NRD convertase | No | No | - |
| Pon2 | 2,03E-02 | -1,60 | paraoxonase 2 | No | No | - |
| Rae1 | 4,05E-02 | -1,60 | ribonucleic acid export 1 | No | No | - |
| Elovl1 | 1,64E-02 | -1,60 | elongation of very long chain fatty acids (FEN1/Elo2, SUR4 | No | No | - |
| Acot9 | 3,56E-02 | -1,60 | acyl-CoA thioesterase 9 | No | No | - |
| Dynlt3 | 3,63E-02 | -1,59 | dynein light chain Tctex-type 3 | No | No | - |
| Tpm1 | 5,15E-03 | -1,59 | tropomyosin 1, alpha | No | No | - |
| Efr3a | 4,58E-02 | -1,59 | EFR3 homolog A | No | No | - |
| Arl8b | 4,91E-02 | -1,58 | ADP-ribosylation factor-like 8B | No | No | - |
| Irf9 | 4,48E-02 | -1,58 | interferon regulatory factor 9 | Yes | No | - |
| Hipk1 | 4,26E-02 | -1,58 | homeodomain interacting protein kinase 1 | No | No | - |
| Lamp2 | 3,31E-02 | -1,58 | lysosomal-associated membrane protein 2 | No | No | 3 |
| Prdx6 | 2,52E-02 | -1,57 | peroxiredoxin 6 | No | No | - |
| Ss18 | 4,05E-02 | -1,57 | SS18, nBAF chromatin remodeling complex subunit | No | No | - |
| Rpl7 | 4,89E-02 | -1,57 | ribosomal protein L7 | No | No | - |
| P3h4 | 3,78E-02 | -1,57 | prolyl 3-hydroxylase family member 4 (non-enzymatic) | No | No | - |
| Tmem120a | 3,62E-02 | -1,56 | transmembrane protein 120A | No | No | - |
| Sh3rf1 | 4,90E-02 | -1,56 | SH3 domain containing ring finger 1 | No | No | - |
| Gm5905 | 4,43E-02 | -1,56 | predicted gene 5905 | No | No | - |
| Csnk1d | 3,99E-02 | -1,56 | casein kinase 1, delta | No | No | - |
| Arf5 | 4,11E-02 | -1,55 | ADP-ribosylation factor 5 | No | No | - |
| Inf2 | 3,24E-02 | -1,55 | inverted formin, FH2 and WH2 domain containing | No | No | - |
| F2r | 4,40E-02 | -1,55 | coagulation factor II (thrombin) receptor | No | No | 1 |
| Arpc4 | 3,38E-02 | -1,55 | actin related protein 2/3 complex, subunit 4 | No | No | - |
| Nudt3 | 3,03E-02 | -1,55 | nudix (nucleotide diphosphate linked moiety X)-type motif | No | No | - |
| Eif3b | 3,19E-02 | -1,55 | eukaryotic translation initiation factor 3, subunit B | No | No | - |
| Pgls | 3,13E-02 | -1,55 | 6-phosphogluconolactonase | No | No | - |
| Serpinf1 | 3,97E-02 | -1,54 | serine (or cysteine) peptidase inhibitor, clade F, member 1 | No | No | - |
| Eva1b | 4,57E-02 | -1,53 | eva-1 homolog B (C. elegans) | No | No | - |
| Sh3glb1 | 3,01E-02 | -1,53 | SH3-domain GRB2-like B1 (endophilin) | No | No | - |
| Mapre1 | 3,17E-02 | -1,53 | microtubule-associated protein, RP/EB family, member 1 | No | No | - |
| Usp50 | 2,94E-02 | -1,53 | ubiquitin specific peptidase 50 | No | No | - |
| Spr | 2,68E-02 | -1,53 | sepiapterin reductase | No | No | 4 |
| Eif4g2 | 4,13E-02 | -1,53 | eukaryotic translation initiation factor 4, gamma 2 | No | No | - |
| Gdi2 | 4,39E-02 | -1,52 | guanosine diphosphate (GDP) dissociation inhibitor 2 | No | No | - |
| Tpm3 | 4,66E-02 | -1,50 | tropomyosin 3, gamma | No | No | - |
| S100a6 | 1,32E-03 | -1,50 | S100 calcium binding protein A6 (calcyclin) | No | No | 7 |
| Mfsd1 | 3,83E-02 | -1,49 | major facilitator superfamily domain containing 1 | No | No | - |
| Puf60 | 3,90E-02 | -1,49 | poly-U binding splicing factor 60 | No | No | - |
| Myl9 | 3,79E-03 | -1,45 | myosin, light polypeptide 9, regulatory | No | No | - |
| Dbi | 4,15E-03 | -1,43 | diazepam binding inhibitor | Yes | No | - |
| P4hb | 3,00E-02 | -1,29 | prolyl 4-hydroxylase, beta polypeptide | No | No | - |
| Cd63 | 3,60E-03 | 1,35 | CD63 antigen | No | No | 18 |
| Laptm4a | 4,84E-03 | 1,37 | lysosomal-associated protein transmembrane 4A | No | No | - |
| Eef1a1 | 4,83E-02 | 1,45 | eukaryotic translation elongation factor 1 alpha 1 | No | No | - |
| Ubl5 | 4,70E-02 | 1,45 | ubiquitin-like 5 | No | No | - |
| Hnrnpa0 | 4,21E-02 | 1,46 | heterogeneous nuclear ribonucleoprotein A0 | No | No | - |
| Eif4b | 4,48E-02 | 1,47 | eukaryotic translation initiation factor 4B | No | No | - |
| H3f3b | 4,05E-02 | 1,47 | H3.3 histone B | No | No | - |
| Rbbp7 | 3,89E-02 | 1,47 | retinoblastoma binding protein 7, chromatin remodeling fac | No | No | - |
| Rps10 | 1,67E-03 | 1,48 | ribosomal protein S10 | No | No | - |
| Ndufb9 | 3,38E-02 | 1,48 | NADH:ubiquinone oxidoreductase subunit B9 | No | No | - |

| | | | | | | |
|---|---|---|---|---|---|---|
| Romo1 | 3,13E-02 | 1,49 | reactive oxygen species modulator 1 | No | No | 3 |
| Rpl11 | 3,53E-02 | 1,49 | ribosomal protein L11 | No | No | - |
| Slirp | 3,30E-02 | 1,50 | SRA stem-loop interacting RNA binding protein | No | No | - |
| Fam32a | 4,32E-02 | 1,50 | family with sequence similarity 32, member A | No | No | - |
| Akirin2 | 4,19E-02 | 1,50 | akirin 2 | No | No | - |
| Rps3 | 1,43E-02 | 1,51 | ribosomal protein S3 | No | No | - |
| Rps29 | 3,46E-02 | 1,51 | ribosomal protein S29 | No | No | - |
| Dctn3 | 3,95E-02 | 1,52 | dynactin 3 | No | No | - |
| Ggn | 3,92E-02 | 1,52 | gametogenetin | No | No | 2 |
| Rpl31 | 3,34E-02 | 1,52 | ribosomal protein L31 | No | No | - |
| Cdc42 | 2,15E-02 | 1,53 | cell division cycle 42 | No | No | 3 |
| Rps26 | 3,53E-02 | 1,53 | ribosomal protein S26 | No | No | - |
| Atp5g3 | 4,39E-02 | 1,53 | ATP synthase, H+ transporting, mitochondrial F0 complex, | No | No | - |
| Cox5a | 2,74E-02 | 1,53 | cytochrome c oxidase subunit 5A | No | No | - |
| Psmd8 | 1,53E-02 | 1,53 | proteasome (prosome, macropain) 26S subunit, non-ATPase | No | No | - |
| Spg21 | 3,80E-02 | 1,54 | SPG21, maspardin | No | No | - |
| Rplp1 | 7,20E-03 | 1,54 | ribosomal protein, large, P1 | No | No | - |
| Rabac1 | 1,82E-02 | 1,54 | Rab acceptor 1 (prenylated) | No | No | - |
| Rpl37 | 7,18E-03 | 1,55 | ribosomal protein L37 | No | No | - |
| Ndufa8 | 2,83E-02 | 1,55 | NADH:ubiquinone oxidoreductase subunit A8 | No | No | - |
| Slc35b2 | 3,59E-02 | 1,55 | solute carrier family 35, member B2 | No | No | - |
| Isca1 | 4,17E-02 | 1,55 | iron-sulfur cluster assembly 1 | No | No | - |
| Atox1 | 3,47E-02 | 1,56 | antioxidant 1 copper chaperone | No | No | - |
| Pgam1 | 3,76E-02 | 1,56 | phosphoglycerate mutase 1 | No | No | - |
| Rab29 | 4,47E-02 | 1,56 | RAB29, member RAS oncogene family | No | No | - |
| Erp44 | 4,46E-02 | 1,56 | endoplasmic reticulum protein 44 | No | No | - |
| Rpl3 | 2,15E-02 | 1,56 | ribosomal protein L3 | No | No | - |
| Clns1a | 3,53E-02 | 1,56 | chloride channel, nucleotide-sensitive, 1A | No | No | - |
| Cst3 | 3,07E-02 | 1,57 | cystatin C | No | No | 2 |
| Gm11478 | 2,90E-02 | 1,57 | predicted gene 11478 | No | No | - |
| Lpcat3 | 4,26E-02 | 1,57 | lysophosphatidylcholine acyltransferase 3 | No | No | - |
| Cdk4 | 4,46E-02 | 1,57 | cyclin-dependent kinase 4 | No | No | 14 |
| Atraid | 4,78E-02 | 1,57 | all-trans retinoic acid induced differentiation factor | No | No | - |
| Mtf1 | 2,92E-02 | 1,58 | metal response element binding transcription factor 1 | No | No | - |
| Nemf | 4,77E-02 | 1,58 | nuclear export mediator factor | No | No | - |
| Vma21 | 3,82E-02 | 1,58 | VMA21 vacuolar H+-ATPase homolog (S. cerevisiae) | No | No | - |
| Yeats4 | 4,43E-02 | 1,58 | YEATS domain containing 4 | No | No | - |
| Gtf2e2 | 2,80E-02 | 1,58 | general transcription factor II E, polypeptide 2 (beta subunit | No | No | - |
| Snhg12 | 2,90E-02 | 1,59 | small nucleolar RNA host gene 12 | No | No | - |
| Ccdc84 | 6,11E-03 | 1,59 | coiled-coil domain containing 84 | No | No | - |
| Mrps7 | 3,52E-02 | 1,59 | mitchondrial ribosomal protein S7 | No | No | - |
| Hnrnpab | 1,41E-02 | 1,59 | heterogeneous nuclear ribonucleoprotein A/B | No | No | 1 |
| Lmod1 | 4,82E-02 | 1,60 | leiomodin 1 (smooth muscle) | No | No | - |
| Dok4 | 2,02E-02 | 1,60 | docking protein 4 | No | No | - |
| Gm3226 | 4,82E-02 | 1,60 | predicted gene 3226 | No | No | - |
| Mmd | 3,80E-02 | 1,60 | monocyte to macrophage differentiation-associated | No | No | 3 |
| Mrpl24 | 2,25E-02 | 1,60 | mitochondrial ribosomal protein L24 | No | No | - |
| Sub1 | 2,40E-02 | 1,60 | SUB1 homolog, transcriptional regulator | No | No | - |
| Etfb | 4,67E-02 | 1,61 | electron transferring flavoprotein, beta polypeptide | No | No | - |
| Rpl23 | 6,40E-04 | 1,61 | ribosomal protein L23 | No | No | - |
| Lsm6 | 2,12E-02 | 1,61 | LSM6 homolog, U6 small nuclear RNA and mRNA degrada | No | No | - |
| Ppp1r18 | 1,68E-02 | 1,61 | protein phosphatase 1, regulatory subunit 18 | No | No | - |
| Polr2c | 1,84E-02 | 1,61 | polymerase (RNA) II (DNA directed) polypeptide C | No | No | - |
| Llgl1 | 2,43E-02 | 1,61 | LLGL1 scribble cell polarity complex component | No | No | - |
| Tpd52l2 | 4,98E-02 | 1,61 | tumor protein D52-like 2 | No | No | - |
| Rbm8a | 1,56E-02 | 1,62 | RNA binding motif protein 8a | No | No | - |
| Trappc5 | 3,96E-02 | 1,62 | trafficking protein particle complex 5 | No | No | - |
| Ldha | 7,87E-03 | 1,62 | lactate dehydrogenase A | No | No | 22 |
| Rpl19 | 2,26E-02 | 1,62 | ribosomal protein L19 | No | No | - |
| F11r | 4,20E-02 | 1,62 | F11 receptor | No | No | - |
| Nkiras1 | 2,71E-02 | 1,62 | NFKB inhibitor interacting Ras-like protein 1 | No | No | - |
| Fkbp4 | 1,43E-02 | 1,62 | FK506 binding protein 4 | No | No | - |
| Atp5e | 3,25E-02 | 1,62 | ATP synthase, H+ transporting, mitochondrial F1 complex, | No | No | - |
| Cdc16 | 2,82E-02 | 1,62 | CDC16 cell division cycle 16 | No | No | - |
| Dnajc8 | 3,01E-02 | 1,62 | DnaJ heat shock protein family (Hsp40) member C8 | No | No | - |
| Grcc10 | 2,71E-02 | 1,62 | gene rich cluster, C10 gene | No | No | - |
| Prdx1 | 3,39E-02 | 1,62 | peroxiredoxin 1 | No | No | - |
| Flii | 2,10E-02 | 1,63 | flightless I actin binding protein | No | No | - |
| Rab2a | 1,39E-02 | 1,63 | RAB2A, member RAS oncogene family | No | No | - |
| Enah | 2,41E-02 | 1,63 | ENAH actin regulator | No | No | - |
| Polr1d | 1,75E-02 | 1,63 | polymerase (RNA) I polypeptide D | No | No | 1 |
| Cct2 | 1,10E-02 | 1,64 | chaperonin containing Tcp1, subunit 2 (beta) | No | No | - |
| Hnrnpa2b1 | 2,58E-02 | 1,64 | heterogeneous nuclear ribonucleoprotein A2/B1 | No | No | - |

| | | | | | | |
|---|---|---|---|---|---|---|
| Commd3 | 3,01E-02 | 1,64 | COMM domain containing 3 | No | No | - |
| Dusp1 | 2,74E-02 | 1,64 | dual specificity phosphatase 1 | No | No | 9 |
| Skap2 | 1,98E-02 | 1,64 | src family associated phosphoprotein 2 | No | No | - |
| Atp5g1 | 1,92E-02 | 1,64 | ATP synthase, H+ transporting, mitochondrial F0 complex, | No | No | - |
| Ero1l | 1,57E-02 | 1,64 | endoplasmic reticulum oxidoreductase 1 alpha | No | No | - |
| Mettl23 | 2,27E-02 | 1,64 | methyltransferase like 23 | No | No | - |
| Rsu1 | 2,90E-02 | 1,65 | Ras suppressor protein 1 | No | No | - |
| Ppib | 4,70E-02 | 1,65 | peptidylprolyl isomerase B | No | No | - |
| Pttg1 | 3,96E-02 | 1,65 | pituitary tumor-transforming gene 1 | No | No | - |
| Gm3375 | 2,22E-02 | 1,65 | predicted gene 3375 | No | No | - |
| Sdc4 | 3,52E-02 | 1,65 | syndecan 4 | No | No | - |
| Mbd2 | 4,87E-02 | 1,65 | methyl-CpG binding domain protein 2 | No | No | 2 |
| Psma7 | 1,93E-02 | 1,65 | proteasome subunit alpha 7 | No | No | - |
| Wdfy1 | 3,53E-02 | 1,65 | WD repeat and FYVE domain containing 1 | No | No | - |
| 0610012G03Rik | 3,39E-02 | 1,65 | RIKEN cDNA 0610012G03 gene | No | No | - |
| Cstb | 1,91E-02 | 1,66 | cystatin B | No | No | - |
| Tvp23b | 3,65E-02 | 1,66 | trans-golgi network vesicle protein 23B | No | No | - |
| Myd88 | 2,48E-02 | 1,66 | myeloid differentiation primary response gene 88 | No | No | 3 |
| Tsc22d1 | 2,59E-02 | 1,66 | TSC22 domain family, member 1 | No | No | - |
| Abracl | 9,62E-03 | 1,66 | ABRA C-terminal like | No | No | - |
| Ndufb8 | 1,26E-02 | 1,66 | NADH:ubiquinone oxidoreductase subunit B8 | No | No | - |
| Gm4735 | 1,09E-02 | 1,66 | predicted gene 4735 | No | No | - |
| Polr2f | 3,78E-02 | 1,66 | polymerase (RNA) II (DNA directed) polypeptide F | No | No | - |
| Gm45713 | 4,26E-03 | 1,66 | predicted gene 45713 | No | No | - |
| Sppl2a | 4,55E-02 | 1,66 | signal peptide peptidase like 2A | No | No | - |
| Tomm7 | 8,79E-03 | 1,66 | translocase of outer mitochondrial membrane 7 | No | No | - |
| Snx1 | 3,52E-02 | 1,66 | sorting nexin 1 | No | No | - |
| Rpl38 | 2,08E-02 | 1,67 | ribosomal protein L38 | No | No | - |
| Slc35f5 | 4,13E-02 | 1,67 | solute carrier family 35, member F5 | No | No | - |
| Cyp20a1 | 2,40E-02 | 1,67 | cytochrome P450, family 20, subfamily a, polypeptide 1 | No | No | - |
| Cdc34 | 1,40E-02 | 1,67 | cell division cycle 34 | No | No | - |
| Lcmt1 | 1,88E-02 | 1,67 | leucine carboxyl methyltransferase 1 | No | No | - |
| Cables2 | 2,22E-04 | 1,67 | CDK5 and Abl enzyme substrate 2 | No | No | - |
| Gm50387 | 2,93E-02 | 1,68 | predicted gene, 50387 | No | No | - |
| Rbx1 | 1,03E-02 | 1,68 | ring-box 1 | No | No | - |
| Gm15542 | 2,30E-02 | 1,68 | predicted gene 15542 | No | No | - |
| Psmd6 | 3,25E-02 | 1,68 | proteasome (prosome, macropain) 26S subunit, non-ATPase | No | No | - |
| Sccpdh | 2,31E-02 | 1,68 | saccharopine dehydrogenase (putative) | No | No | - |
| Fam227b | 4,69E-02 | 1,68 | family with sequence similarity 227, member B | No | No | - |
| Tmem183a | 2,26E-02 | 1,68 | transmembrane protein 183A | No | No | - |
| Ptges | 2,55E-02 | 1,68 | prostaglandin E synthase | No | No | 10 |
| Ist1 | 1,84E-02 | 1,68 | increased sodium tolerance 1 homolog (yeast) | No | No | - |
| Cnn1 | 2,23E-03 | 1,68 | calponin 1 | No | No | 2 |
| Slc2a1 | 9,02E-03 | 1,68 | solute carrier family 2 (facilitated glucose transporter), mem | No | No | - |
| Shmt2 | 2,31E-02 | 1,69 | serine hydroxymethyltransferase 2 (mitochondrial) | No | No | 2 |
| Galnt2l | 1,55E-02 | 1,69 | - | No | No | - |
| Bcas2 | 1,94E-02 | 1,69 | BCAS2 pre-mRNA processing factor | No | No | - |
| Derl2 | 4,43E-02 | 1,69 | Der1-like domain family, member 2 | No | No | - |
| Rps21 | 6,87E-04 | 1,69 | ribosomal protein S21 | No | No | - |
| Bloc1s1 | 1,32E-02 | 1,69 | biogenesis of lysosomal organelles complex-1, subunit 1 | No | No | - |
| Mrpl35 | 2,99E-02 | 1,69 | mitochondrial ribosomal protein L35 | No | No | - |
| Lmo4 | 1,12E-02 | 1,69 | LIM domain only 4 | No | No | - |
| Smco4 | 2,23E-02 | 1,69 | single-pass membrane protein with coiled-coil domains 4 | No | No | - |
| Galnt2 | 1,44E-02 | 1,70 | polypeptide N-acetylgalactosaminyltransferase 2 | No | No | - |
| Vhl | 3,45E-02 | 1,70 | von Hippel-Lindau tumor suppressor | No | No | 7 |
| Gm10288 | 4,60E-03 | 1,70 | predicted gene 10288 | No | No | - |
| Atp5j2 | 1,29E-03 | 1,70 | ATP synthase, H+ transporting, mitochondrial F0 complex, | No | No | - |
| Btg2 | 1,83E-02 | 1,70 | BTG anti-proliferation factor 2 | No | No | - |
| Ccz1 | 2,39E-02 | 1,70 | CCZ1 vacuolar protein trafficking and biogenesis associated | No | No | - |
| Rpl24 | 2,07E-02 | 1,70 | ribosomal protein L24 | No | No | - |
| Eif1ad | 1,51E-02 | 1,70 | eukaryotic translation initiation factor 1A domain containing | No | No | - |
| Rpl13a | 2,31E-03 | 1,70 | ribosomal protein L13A | No | No | - |
| Taf13 | 4,12E-02 | 1,70 | TATA-box binding protein associated factor 13 | No | No | - |
| Cmtm3 | 3,04E-02 | 1,70 | CKLF-like MARVEL transmembrane domain containing 3 | No | No | - |
| Thra | 4,64E-02 | 1,71 | thyroid hormone receptor alpha | No | No | - |
| 2210016F16Rik | 4,73E-02 | 1,71 | RIKEN cDNA 2210016F16 gene | No | No | - |
| S100a10 | 7,29E-03 | 1,71 | S100 calcium binding protein A10 (calpactin) | No | No | - |
| Rps11 | 1,91E-02 | 1,71 | ribosomal protein S11 | No | No | - |
| Nedd8 | 1,65E-04 | 1,71 | neural precursor cell expressed, developmentally down-regu | No | No | 13 |
| Arl6ip1 | 6,13E-03 | 1,71 | ADP-ribosylation factor-like 6 interacting protein 1 | No | No | - |
| Thbd | 3,18E-02 | 1,71 | thrombomodulin | No | No | - |
| Uqcrq | 6,51E-04 | 1,71 | ubiquinol-cytochrome c reductase, complex III subunit VII | No | No | - |
| Fam169a | 2,29E-02 | 1,71 | family with sequence similarity 169, member A | No | No | - |

| | | | | | | |
|---|---|---|---|---|---|---|
| Mrpl55 | 3,99E-02 | 1,71 | mitochondrial ribosomal protein L55 | No | No | - |
| 4930453N24Rik | 4,65E-02 | 1,71 | RIKEN cDNA 4930453N24 gene | No | No | - |
| Sec14l1 | 1,28E-02 | 1,71 | SEC14-like lipid binding 1 | No | No | - |
| Dnajc2 | 1,21E-02 | 1,71 | DnaJ heat shock protein family (Hsp40) member C2 | No | No | - |
| Cwc15 | 1,03E-02 | 1,72 | CWC15 spliceosome-associated protein | No | No | - |
| AL731706.1 | 2,89E-02 | 1,72 | novel protein | No | No | - |
| Gpd2 | 3,32E-02 | 1,72 | glycerol phosphate dehydrogenase 2, mitochondrial | No | No | - |
| Cfl2 | 8,36E-03 | 1,72 | cofilin 2, muscle | No | No | - |
| Slc10a3 | 2,86E-02 | 1,72 | solute carrier family 10 (sodium/bile acid cotransporter fam | No | No | - |
| Serinc4 | 1,34E-02 | 1,72 | serine incorporator 4 | No | No | - |
| Csnk2a1 | 2,77E-02 | 1,72 | casein kinase 2, alpha 1 polypeptide | No | No | - |
| Gnb5 | 2,98E-02 | 1,72 | guanine nucleotide binding protein (G protein), beta 5 | No | No | - |
| Mrps23 | 1,20E-02 | 1,72 | mitochondrial ribosomal protein S23 | No | No | - |
| Gm15427 | 2,03E-02 | 1,72 | predicted pseudogene 15427 | No | No | - |
| Fopnl | 1,93E-02 | 1,73 | centrosomal protein 20 | No | No | - |
| Map3k12 | 9,96E-03 | 1,73 | mitogen-activated protein kinase kinase kinase 12 | No | No | - |
| Ifi27 | 1,25E-03 | 1,73 | interferon, alpha-inducible protein 27 | No | No | - |
| Rfk | 1,14E-02 | 1,73 | riboflavin kinase | No | No | - |
| Rpl27a | 6,47E-03 | 1,73 | ribosomal protein L27A | No | No | - |
| Pfn2 | 3,01E-02 | 1,73 | profilin 2 | No | No | - |
| Spryd7 | 2,93E-02 | 1,73 | SPRY domain containing 7 | No | No | - |
| Ubac1 | 2,67E-02 | 1,73 | ubiquitin associated domain containing 1 | No | No | - |
| Phykpl | 4,17E-03 | 1,73 | 5-phosphohydroxy-L-lysine phospholyase | No | No | - |
| Slc35a1 | 1,97E-02 | 1,73 | solute carrier family 35 (CMP-sialic acid transporter), meml | No | No | - |
| Rps15a | 4,58E-04 | 1,73 | ribosomal protein S15A | No | No | - |
| Ppp1r13b | 1,88E-02 | 1,74 | protein phosphatase 1, regulatory subunit 13B | No | No | 8 |
| Tmem242 | 1,31E-02 | 1,74 | transmembrane protein 242 | No | No | - |
| Snrpd1 | 7,01E-03 | 1,74 | small nuclear ribonucleoprotein D1 | No | No | - |
| Pabpc1 | 2,55E-02 | 1,74 | poly(A) binding protein, cytoplasmic 1 | No | No | - |
| C1qtnf5 | 4,43E-02 | 1,74 | C1q and tumor necrosis factor related protein 5 | No | No | - |
| Brwd1 | 4,83E-02 | 1,74 | bromodomain and WD repeat domain containing 1 | No | No | - |
| Pdia6 | 1,99E-03 | 1,74 | protein disulfide isomerase associated 6 | No | No | - |
| Rab10 | 8,98E-03 | 1,74 | RAB10, member RAS oncogene family | No | No | - |
| Actr3 | 3,20E-03 | 1,74 | ARP3 actin-related protein 3 | Yes | No | - |
| Parn | 5,00E-02 | 1,75 | poly(A)-specific ribonuclease (deadenylation nuclease) | No | No | 56 |
| Nt5c3 | 1,98E-02 | 1,75 | 5'-nucleotidase, cytosolic III | Yes | No | - |
| Cox6b1 | 8,09E-04 | 1,75 | cytochrome c oxidase, subunit 6B1 | No | No | - |
| Prrx1 | 1,01E-02 | 1,75 | paired related homeobox 1 | No | No | - |
| Psma4 | 2,18E-02 | 1,75 | proteasome subunit alpha 4 | No | No | - |
| Gm9816 | 1,53E-02 | 1,75 | predicted pseudogene 9816 | No | No | - |
| Abhd11 | 4,20E-02 | 1,75 | abhydrolase domain containing 11 | No | No | - |
| Timm10 | 3,09E-02 | 1,75 | translocase of inner mitochondrial membrane 10 | No | No | - |
| Dnajc9 | 3,23E-02 | 1,75 | DnaJ heat shock protein family (Hsp40) member C9 | No | No | - |
| Wdr61 | 3,04E-02 | 1,75 | WD repeat domain 61 | No | No | - |
| Mrps16 | 1,36E-02 | 1,75 | mitochondrial ribosomal protein S16 | No | No | - |
| Ulk3 | 1,50E-02 | 1,75 | unc-51-like kinase 3 | No | No | - |
| Atl2 | 3,08E-02 | 1,75 | atlastin GTPase 2 | No | No | - |
| Gde1 | 1,27E-02 | 1,75 | glycerophosphodiester phosphodiesterase 1 | No | No | - |
| Phf10 | 9,36E-03 | 1,76 | PHD finger protein 10 | No | No | - |
| Cript | 1,31E-02 | 1,76 | cysteine-rich PDZ-binding protein | No | No | - |
| Nbdy | 1,73E-02 | 1,76 | negative regulator of P-body association | No | No | - |
| Nxn | 6,90E-03 | 1,76 | nucleoredoxin | No | No | - |
| Txndc15 | 2,67E-02 | 1,76 | thioredoxin domain containing 15 | No | No | - |
| Pcbd2 | 3,03E-02 | 1,76 | pterin 4 alpha carbinolamine dehydratase/dimerization cofa | No | No | - |
| Rbm7 | 2,27E-02 | 1,76 | RNA binding motif protein 7 | No | No | 2 |
| Marchf5 | 5,93E-03 | 1,76 | membrane associated ring-CH-type finger 5 | No | No | - |
| Dbf4 | 4,74E-02 | 1,76 | DBF4 zinc finger | No | No | - |
| Zcchc24 | 1,38E-02 | 1,76 | zinc finger, CCHC domain containing 24 | No | No | - |
| Rab1b | 2,76E-02 | 1,76 | RAB1B, member RAS oncogene family | No | No | - |
| Polr2j | 6,91E-03 | 1,77 | polymerase (RNA) II (DNA directed) polypeptide J | No | No | - |
| Decr1 | 1,40E-02 | 1,77 | 2,4-dienoyl CoA reductase 1, mitochondrial | No | No | - |
| Mmadhc | 3,43E-03 | 1,77 | methylmalonic aciduria (cobalamin deficiency) cblD type, v | No | No | - |
| Selenos | 1,98E-02 | 1,77 | selenoprotein S | No | No | - |
| Slc25a4 | 4,18E-03 | 1,77 | solute carrier family 25 (mitochondrial carrier, adenine nucl | No | No | - |
| Zadh2 | 4,80E-02 | 1,77 | zinc binding alcohol dehydrogenase, domain containing 2 | No | No | - |
| Ssu72 | 1,22E-02 | 1,77 | Ssu72 RNA polymerase II CTD phosphatase homolog (yeas | No | No | - |
| Fam89b | 1,88E-02 | 1,77 | family with sequence similarity 89, member B | No | No | - |
| Commd4 | 1,71E-02 | 1,77 | COMM domain containing 4 | No | No | - |
| Avpi1 | 1,12E-02 | 1,77 | arginine vasopressin-induced 1 | No | No | - |
| Ssr4 | 9,91E-05 | 1,77 | signal sequence receptor, delta | No | No | - |
| Vwa5a | 3,53E-02 | 1,78 | von Willebrand factor A domain containing 5A | No | No | - |
| Zbtb7b | 1,03E-02 | 1,78 | zinc finger and BTB domain containing 7B | No | No | - |
| Rps25 | 6,48E-05 | 1,78 | ribosomal protein S25 | No | No | - |

| | | | | | | |
|---|---|---|---|---|---|---|
| AI413582 | 1,45E-02 | 1,78 | expressed sequence AI413582 | No | No | - |
| Rpl12 | 4,79E-05 | 1,78 | ribosomal protein L12 | No | No | - |
| Ran | 2,15E-03 | 1,78 | RAN, member RAS oncogene family | No | No | - |
| Adra1b | 2,31E-02 | 1,78 | adrenergic receptor, alpha 1b | No | No | - |
| Psme2b | 1,98E-02 | 1,78 | protease (prosome, macropain) activator subunit 2B | No | No | - |
| Hint1 | 1,34E-04 | 1,78 | histidine triad nucleotide binding protein 1 | No | No | - |
| Fam104a | 8,31E-03 | 1,78 | family with sequence similarity 104, member A | No | No | - |
| Tmem18 | 3,16E-02 | 1,78 | transmembrane protein 18 | No | No | - |
| Cnih1 | 4,14E-03 | 1,78 | cornichon family AMPA receptor auxiliary protein 1 | No | No | - |
| Rpl10-ps5 | 8,16E-03 | 1,78 | ribosomal protein L10, pseudogene 5 | No | No | - |
| Pim3 | 2,23E-02 | 1,78 | proviral integration site 3 | No | No | - |
| Mfrp | 4,31E-02 | 1,78 | membrane frizzled-related protein | No | No | - |
| Mrps35 | 4,42E-02 | 1,78 | mitochondrial ribosomal protein S35 | No | No | - |
| Cox7b | 2,55E-03 | 1,79 | cytochrome c oxidase subunit 7B | No | No | - |
| Prdx2 | 2,30E-03 | 1,79 | peroxiredoxin 2 | No | No | - |
| Atp6v1h | 2,72E-02 | 1,79 | ATPase, H+ transporting, lysosomal V1 subunit H | No | No | - |
| Il6 | 3,82E-02 | 1,79 | interleukin 6 | Yes | No | - |
| Hmga1 | 1,51E-02 | 1,79 | high mobility group AT-hook 1 | No | No | - |
| Nus1 | 5,27E-03 | 1,79 | NUS1 dehydrodolichyl diphosphate synthase subunit | No | No | - |
| Pno1 | 2,05E-02 | 1,79 | partner of NOB1 homolog | No | No | 4 |
| D8Ertd738e | 3,37E-03 | 1,79 | DNA segment, Chr 8, ERATO Doi 738, expressed | No | No | - |
| Cyc1 | 2,36E-02 | 1,79 | cytochrome c-1 | No | No | - |
| Pdss1 | 8,69E-03 | 1,79 | prenyl (solanesyl) diphosphate synthase, subunit 1 | No | No | - |
| Pop4 | 3,96E-02 | 1,79 | processing of precursor 4, ribonuclease P/MRP family, (S. c | No | No | - |
| Atl1 | 2,57E-02 | 1,79 | atlastin GTPase 1 | No | No | - |
| Fnta | 1,24E-02 | 1,79 | farnesyltransferase, CAAX box, alpha | No | No | - |
| Anp32a | 3,96E-02 | 1,79 | acidic (leucine-rich) nuclear phosphoprotein 32 family, men | No | No | - |
| Hprt | 5,78E-03 | 1,79 | hypoxanthine guanine phosphoribosyl transferase | No | No | - |
| Chmp3 | 1,52E-02 | 1,79 | charged multivesicular body protein 3 | No | No | - |
| Il1rn | 4,93E-02 | 1,79 | interleukin 1 receptor antagonist | No | No | 13 |
| Ndufs8 | 1,71E-02 | 1,80 | NADH:ubiquinone oxidoreductase core subunit S8 | No | No | - |
| Cand2 | 1,09E-04 | 1,80 | cullin-associated and neddylation-dissociated 2 (putative) | No | No | - |
| Gm49527 | 2,40E-02 | 1,80 | predicted gene, 49527 | No | No | - |
| Rraga | 1,25E-02 | 1,80 | Ras-related GTP binding A | No | No | - |
| Paics | 1,66E-02 | 1,80 | phosphoribosylaminoimidazole carboxylase, phosphoribosy | No | No | - |
| Ube2a | 5,60E-03 | 1,80 | ubiquitin-conjugating enzyme E2A | No | No | - |
| Tubb6 | 5,35E-03 | 1,80 | tubulin, beta 6 class V | No | No | - |
| Gm45234 | 5,87E-03 | 1,80 | predicted gene 45234 | No | No | - |
| Cyth2 | 2,79E-02 | 1,80 | cytohesin 2 | No | No | - |
| Ptgr2 | 4,84E-02 | 1,80 | prostaglandin reductase 2 | No | No | - |
| Ccndbp1 | 4,87E-02 | 1,80 | cyclin D-type binding-protein 1 | No | No | - |
| Mlec | 1,38E-03 | 1,81 | malectin | No | No | 2 |
| Fis1 | 8,82E-03 | 1,81 | fission, mitochondrial 1 | No | No | 3 |
| Pon3 | 2,53E-02 | 1,81 | paraoxonase 3 | No | No | - |
| F3 | 3,77E-02 | 1,81 | coagulation factor III | No | No | 6 |
| Nol7 | 6,12E-03 | 1,81 | nucleolar protein 7 | No | No | - |
| Dhrs3 | 2,86E-02 | 1,81 | dehydrogenase/reductase (SDR family) member 3 | No | No | - |
| Rrnad1 | 4,61E-03 | 1,81 | ribosomal RNA adenine dimethylase domain containing 1 | No | No | - |
| Lrwd1 | 5,39E-03 | 1,81 | leucine-rich repeats and WD repeat domain containing 1 | No | No | - |
| Ghitm | 3,31E-04 | 1,81 | growth hormone inducible transmembrane protein | No | No | - |
| 1700015O11Rik | 2,26E-02 | 1,81 | RIKEN cDNA 1700015O11 gene | No | No | - |
| Mif4gd | 4,87E-02 | 1,81 | MIF4G domain containing | No | No | - |
| Arl3 | 7,20E-03 | 1,81 | ADP-ribosylation factor-like 3 | No | No | - |
| Capza2 | 5,08E-03 | 1,81 | capping protein (actin filament) muscle Z-line, alpha 2 | No | No | - |
| Ngdn | 7,96E-03 | 1,81 | neuroguidin, EIF4E binding protein | No | No | - |
| Ankmy1 | 3,77E-03 | 1,82 | ankyrin repeat and MYND domain containing 1 | No | No | - |
| Ilf3 | 2,32E-02 | 1,82 | interleukin enhancer binding factor 3 | No | No | - |
| Oat | 6,55E-03 | 1,82 | ornithine aminotransferase | No | No | - |
| Pnp | 4,36E-03 | 1,82 | purine-nucleoside phosphorylase | No | No | 7 |
| Kbtbd4 | 4,13E-02 | 1,82 | kelch repeat and BTB (POZ) domain containing 4 | No | No | - |
| Tmem237 | 4,44E-02 | 1,82 | transmembrane protein 237 | No | No | - |
| Coprs | 1,75E-02 | 1,82 | coordinator of PRMT5, differentiation stimulator | No | No | - |
| Fzd1 | 2,97E-02 | 1,82 | frizzled class receptor 1 | No | No | - |
| Fam114a2 | 4,81E-03 | 1,82 | family with sequence similarity 114, member A2 | No | No | - |
| Mrpl3 | 1,83E-02 | 1,82 | mitochondrial ribosomal protein L3 | No | No | - |
| Esd | 1,34E-03 | 1,83 | esterase D/formylglutathione hydrolase | No | No | - |
| Alg14 | 1,09E-02 | 1,83 | asparagine-linked glycosylation 14 | No | No | - |
| Uap1 | 2,77E-02 | 1,83 | UDP-N-acetylglucosamine pyrophosphorylase 1 | No | No | - |
| Mdh1 | 4,77E-04 | 1,83 | malate dehydrogenase 1, NAD (soluble) | No | No | - |
| Pcbp2 | 6,35E-03 | 1,83 | poly(rC) binding protein 2 | No | No | - |
| Nup35 | 4,11E-02 | 1,83 | nucleoporin 35 | No | No | - |
| Msl3 | 3,99E-02 | 1,83 | MSL complex subunit 3 | No | No | - |
| 1700025G04Rik | 4,13E-02 | 1,83 | RIKEN cDNA 1700025G04 gene | No | No | - |

| | | | | | | |
|---|---|---|---|---|---|---|
| Ubald2 | 3,53E-02 | 1,83 | UBA-like domain containing 2 | No | No | - |
| Stt3a | 7,87E-03 | 1,83 | STT3, subunit of the oligosaccharyltransferase complex, ho | No | No | - |
| Rpl32 | 6,23E-05 | 1,83 | ribosomal protein L32 | No | No | - |
| Lyrm2 | 4,08E-02 | 1,84 | LYR motif containing 2 | No | No | - |
| Ube2g1 | 2,04E-02 | 1,84 | ubiquitin-conjugating enzyme E2G 1 | No | No | - |
| Rpl36a | 7,07E-05 | 1,84 | ribosomal protein L36A | No | No | - |
| Csrp2 | 4,63E-03 | 1,84 | cysteine and glycine-rich protein 2 | No | No | - |
| Rpsa-ps7 | 3,23E-02 | 1,84 | ribosomal protein SA, pseudogene 7 | No | No | - |
| Atpif1 | 1,90E-04 | 1,84 | ATPase inhibitory factor 1 | No | No | - |
| Dram2 | 2,70E-02 | 1,84 | DNA-damage regulated autophagy modulator 2 | No | No | - |
| Azi2 | 8,94E-03 | 1,84 | 5-azacytidine induced gene 2 | No | No | - |
| Mex3c | 3,73E-02 | 1,84 | mex3 RNA binding family member C | No | No | - |
| Tmem9 | 7,09E-03 | 1,85 | transmembrane protein 9 | No | No | - |
| Map3k15 | 3,92E-03 | 1,85 | mitogen-activated protein kinase kinase kinase 15 | No | No | - |
| C1d | 2,41E-02 | 1,85 | C1D nuclear receptor co-repressor | No | No | - |
| Dgcr6 | 1,56E-02 | 1,85 | DiGeorge syndrome critical region gene 6 | No | No | - |
| Oaz1-ps | 3,13E-03 | 1,85 | ornithine decarboxylase antizyme 1, pseudogene | No | No | - |
| Lsm14a | 6,08E-03 | 1,86 | LSM14A mRNA processing body assembly factor | No | No | - |
| Atp5h | 2,57E-03 | 1,86 | ATP synthase, H+ transporting, mitochondrial F0 complex, | No | No | - |
| Hat1 | 2,19E-02 | 1,86 | histone aminotransferase 1 | No | No | - |
| Dhrs7 | 1,70E-02 | 1,86 | dehydrogenase/reductase (SDR family) member 7 | No | No | - |
| Btf3l4 | 1,14E-02 | 1,86 | basic transcription factor 3-like 4 | No | No | - |
| Cops3 | 7,99E-03 | 1,86 | COP9 signalosome subunit 3 | No | No | - |
| Mfsd14a | 2,40E-02 | 1,86 | major facilitator superfamily domain containing 14A | No | No | - |
| Snapc1 | 3,98E-03 | 1,86 | small nuclear RNA activating complex, polypeptide 1 | No | No | - |
| Ndufa9 | 1,09E-02 | 1,86 | NADH:ubiquinone oxidoreductase subunit A9 | No | No | - |
| Rprd1b | 3,22E-02 | 1,86 | regulation of nuclear pre-mRNA domain containing 1B | No | No | - |
| Bmper | 5,31E-03 | 1,86 | BMP-binding endothelial regulator | No | No | 5 |
| Mrpl14 | 1,40E-02 | 1,87 | mitochondrial ribosomal protein L14 | No | No | - |
| Thap12 | 1,64E-02 | 1,87 | THAP domain containing 12 | No | No | - |
| Gosr2 | 1,19E-02 | 1,87 | golgi SNAP receptor complex member 2 | No | No | - |
| Bahd1 | 3,15E-02 | 1,87 | bromo adjacent homology domain containing 1 | No | No | - |
| Gm17383 | 3,51E-03 | 1,87 | predicted gene, 17383 | No | No | - |
| Mrps17 | 3,28E-03 | 1,87 | mitochondrial ribosomal protein S17 | No | No | - |
| Abi1 | 4,31E-03 | 1,87 | abl interactor 1 | No | No | - |
| Srsf2 | 4,85E-03 | 1,87 | serine and arginine-rich splicing factor 2 | No | No | 1 |
| Wdr55 | 3,83E-02 | 1,87 | WD repeat domain 55 | No | No | - |
| Eif3i | 3,62E-03 | 1,87 | eukaryotic translation initiation factor 3, subunit I | No | No | - |
| 1600012H06Rik | 3,06E-03 | 1,87 | RIKEN cDNA 1600012H06 gene | No | No | - |
| Msantd3 | 2,15E-02 | 1,87 | Myb/SANT-like DNA-binding domain containing 3 | No | No | - |
| Dynlrb1 | 3,72E-03 | 1,87 | dynein light chain roadblock-type 1 | No | No | - |
| Rasl11b | 1,09E-02 | 1,88 | RAS-like, family 11, member B | No | No | - |
| Tbpl1 | 9,30E-03 | 1,88 | TATA box binding protein-like 1 | No | No | 2 |
| Hexim1 | 2,15E-02 | 1,88 | hexamethylene bis-acetamide inducible 1 | No | No | - |
| Manf | 4,89E-03 | 1,88 | mesencephalic astrocyte-derived neurotrophic factor | No | No | - |
| Pdlim2 | 4,82E-04 | 1,88 | PDZ and LIM domain 2 | No | No | - |
| Jam3 | 2,40E-02 | 1,88 | junction adhesion molecule 3 | No | No | - |
| Ndufb10 | 2,34E-04 | 1,88 | NADH:ubiquinone oxidoreductase subunit B10 | No | No | - |
| Snx22 | 1,19E-02 | 1,88 | sorting nexin 22 | No | No | - |
| Rcc2 | 5,11E-03 | 1,88 | regulator of chromosome condensation 2 | No | No | - |
| Prdx5 | 2,84E-05 | 1,88 | peroxiredoxin 5 | No | No | - |
| Enkd1 | 3,62E-02 | 1,88 | enkurin domain containing 1 | No | No | - |
| Mapkbp1 | 3,35E-02 | 1,88 | mitogen-activated protein kinase binding protein 1 | No | No | - |
| 2300009A05Rik | 1,80E-02 | 1,88 | RIKEN cDNA 2300009A05 gene | No | No | - |
| Ccdc71l | 3,95E-02 | 1,89 | coiled-coil domain containing 71 like | No | No | - |
| Acta1 | 4,33E-02 | 1,89 | actin alpha 1, skeletal muscle | Yes | No | - |
| Nsa2 | 5,92E-03 | 1,89 | NSA2 ribosome biogenesis homolog | No | No | - |
| Tmem123 | 6,55E-03 | 1,89 | transmembrane protein 123 | No | No | - |
| Dera | 8,21E-03 | 1,89 | deoxyribose-phosphate aldolase (putative) | No | No | - |
| Senp3 | 3,42E-02 | 1,89 | SUMO/sentrin specific peptidase 3 | No | No | - |
| Vps36 | 4,65E-03 | 1,89 | vacuolar protein sorting 36 | No | No | - |
| Mrpl58 | 4,45E-02 | 1,89 | mitochondrial ribosomal protein L58 | No | No | - |
| Cox7a2 | 1,42E-04 | 1,89 | cytochrome c oxidase subunit 7A2 | No | No | - |
| Gm21992 | 1,57E-02 | 1,89 | predicted gene 21992 | No | No | - |
| Phgdh | 2,73E-02 | 1,90 | 3-phosphoglycerate dehydrogenase | No | No | 9 |
| Pfdn4 | 1,78E-02 | 1,90 | prefoldin 4 | No | No | - |
| Dusp11 | 5,08E-03 | 1,90 | dual specificity phosphatase 11 (RNA/RNP complex 1-inter | No | No | - |
| Rnaset2b | 1,26E-02 | 1,90 | ribonuclease T2B | No | No | - |
| Rras2 | 1,79E-03 | 1,90 | related RAS viral (r-ras) oncogene 2 | No | No | - |
| Gm49721 | 1,24E-02 | 1,90 | predicted gene, 49721 | No | No | - |
| Ndufs4 | 2,57E-03 | 1,90 | NADH:ubiquinone oxidoreductase core subunit S4 | No | No | - |
| Rdh10 | 3,89E-02 | 1,90 | retinol dehydrogenase 10 (all-trans) | No | No | - |
| Ndufa1 | 3,19E-03 | 1,90 | NADH:ubiquinone oxidoreductase subunit A1 | No | No | - |

| | | | | | | |
|---|---|---|---|---|---|---|
| Cbfb | 7,98E-03 | 1,90 | core binding factor beta | No | No | - |
| Eef1akmt1 | 2,12E-02 | 1,90 | EEF1A alpha lysine methyltransferase 1 | No | No | - |
| Gpt2 | 2,59E-02 | 1,91 | glutamic pyruvate transaminase (alanine aminotransferase) 2 | No | No | - |
| Sri | 2,78E-03 | 1,91 | sorcin | No | No | 1 |
| Sass6 | 2,88E-02 | 1,91 | SAS-6 centriolar assembly protein | No | No | - |
| Cnot8 | 8,94E-03 | 1,91 | CCR4-NOT transcription complex, subunit 8 | No | No | - |
| Mrpl41 | 1,87E-02 | 1,91 | mitochondrial ribosomal protein L41 | No | No | - |
| Pole4 | 3,09E-03 | 1,91 | polymerase (DNA-directed), epsilon 4 (p12 subunit) | No | No | - |
| Mphosph6 | 9,63E-03 | 1,91 | M phase phosphoprotein 6 | No | No | - |
| Iffo1 | 3,10E-02 | 1,91 | intermediate filament family orphan 1 | No | No | - |
| Rita1 | 2,76E-02 | 1,91 | RBPJ interacting and tubulin associated 1 | No | No | - |
| Hint2 | 7,71E-03 | 1,91 | histidine triad nucleotide binding protein 2 | No | No | - |
| Cnpy4 | 8,08E-03 | 1,92 | canopy FGF signaling regulator 4 | No | No | - |
| Lsm3 | 3,39E-02 | 1,92 | LSM3 homolog, U6 small nuclear RNA and mRNA degrada | No | No | - |
| Bex3 | 4,72E-04 | 1,92 | brain expressed X-linked 3 | No | No | - |
| Pard6a | 3,03E-02 | 1,92 | par-6 family cell polarity regulator alpha | No | No | - |
| Dhrs1 | 2,32E-03 | 1,92 | dehydrogenase/reductase (SDR family) member 1 | No | No | - |
| Tango2 | 1,20E-02 | 1,92 | transport and golgi organization 2 | No | No | - |
| Thoc7 | 3,82E-02 | 1,92 | THO complex 7 | No | No | - |
| Eif3e | 1,77E-03 | 1,92 | eukaryotic translation initiation factor 3, subunit E | No | No | - |
| Tpst2 | 1,28E-02 | 1,92 | protein-tyrosine sulfotransferase 2 | No | No | - |
| Gpc1 | 1,62E-03 | 1,92 | glypican 1 | No | No | - |
| Srsf3 | 1,22E-03 | 1,93 | serine and arginine-rich splicing factor 3 | No | No | - |
| Pcif1 | 1,94E-02 | 1,93 | phosphorylated CTD interacting factor 1 | No | No | - |
| Nsmce1 | 5,91E-03 | 1,93 | NSE1 homolog, SMC5-SMC6 complex component | No | No | - |
| Xbp1 | 1,27E-03 | 1,93 | X-box binding protein 1 | No | No | 10 |
| Gapdh | 3,95E-02 | 1,93 | glyceraldehyde-3-phosphate dehydrogenase | No | No | 7 |
| Bcl9 | 4,45E-02 | 1,93 | B cell CLL/lymphoma 9 | No | No | - |
| Rala | 1,65E-03 | 1,93 | v-ral simian leukemia viral oncogene A (ras related) | No | No | - |
| Uqcr11 | 7,98E-03 | 1,93 | ubiquinol-cytochrome c reductase, complex III subunit XI | No | No | - |
| Man2a1 | 1,60E-02 | 1,93 | mannosidase 2, alpha 1 | No | No | - |
| Tmem167 | 1,09E-04 | 1,93 | transmembrane protein 167 | No | No | - |
| Exosc8 | 3,52E-02 | 1,93 | exosome component 8 | No | No | - |
| Aopep | 7,88E-03 | 1,94 | aminopeptidase O | No | No | - |
| Hnrnph3 | 2,66E-02 | 1,94 | heterogeneous nuclear ribonucleoprotein H3 | No | No | - |
| Zfp36l1 | 3,00E-02 | 1,94 | zinc finger protein 36, C3H type-like 1 | No | No | - |
| Pigyl | 4,26E-03 | 1,94 | phosphatidylinositol glycan anchor biosynthesis, class Y-lik | No | No | - |
| Tmem50b | 1,80E-02 | 1,94 | transmembrane protein 50B | No | No | - |
| Cdk2ap1 | 1,33E-03 | 1,94 | CDK2 (cyclin-dependent kinase 2)-associated protein 1 | No | No | - |
| Abhd16a | 2,47E-02 | 1,94 | abhydrolase domain containing 16A | No | No | - |
| Polr3f | 2,30E-02 | 1,94 | polymerase (RNA) III (DNA directed) polypeptide F | No | No | - |
| Trim41 | 2,12E-02 | 1,94 | tripartite motif-containing 41 | No | No | - |
| B4galt3 | 3,24E-02 | 1,94 | UDP-Gal:betaGlcNAc beta 1,4-galactosyltransferase, polyp | No | No | - |
| Cog1 | 3,35E-03 | 1,94 | component of oligomeric golgi complex 1 | No | No | - |
| Sys1 | 4,79E-03 | 1,94 | SYS1 Golgi-localized integral membrane protein homolog ( | No | No | - |
| Zfp593 | 2,08E-02 | 1,94 | zinc finger protein 593 | No | No | - |
| Mif | 6,20E-03 | 1,94 | macrophage migration inhibitory factor (glycosylation-inhib | No | No | 123 |
| 9430015G10Rik | 3,76E-02 | 1,94 | RIKEN cDNA 9430015G10 gene | No | No | - |
| Mrpl16 | 6,58E-03 | 1,94 | mitochondrial ribosomal protein L16 | No | No | - |
| Csnk2a3 | 4,67E-02 | 1,94 | casein kinase 2, alpha 1 polypeptide, pseudogene 3 | No | No | - |
| Micos10 | 8,65E-04 | 1,94 | mitochondrial contact site and cristae organizing system sub | No | No | - |
| Ciao2b | 4,64E-03 | 1,95 | cytosolic iron-sulfur assembly component 2B | No | No | - |
| Hnrnpll | 1,95E-02 | 1,95 | heterogeneous nuclear ribonucleoprotein L-like | No | No | - |
| Gla | 1,86E-05 | 1,95 | galactosidase, alpha | No | No | 24 |
| Mrpl43 | 3,46E-03 | 1,95 | mitochondrial ribosomal protein L43 | No | No | - |
| Twist2 | 8,18E-03 | 1,95 | twist basic helix-loop-helix transcription factor 2 | No | No | - |
| Mrpl13 | 3,88E-03 | 1,95 | mitochondrial ribosomal protein L13 | No | No | - |
| Pak1ip1 | 3,16E-03 | 1,95 | PAK1 interacting protein 1 | No | No | - |
| Iqschfp | 3,01E-02 | 1,95 | Iqcj and Schip1 fusion protein | No | No | - |
| Slc33a1 | 1,25E-02 | 1,95 | solute carrier family 33 (acetyl-CoA transporter), member 1 | No | No | - |
| Rbm4 | 9,34E-03 | 1,95 | RNA binding motif protein 4 | No | No | - |
| Ndufa4l2 | 8,13E-03 | 1,96 | Ndufa4, mitochondrial complex associated like 2 | No | No | - |
| Pgap1 | 4,98E-02 | 1,96 | post-GPI attachment to proteins 1 | No | No | - |
| Tmem256 | 3,64E-04 | 1,96 | transmembrane protein 256 | No | No | - |
| Rheb | 2,11E-03 | 1,96 | Ras homolog enriched in brain | No | No | - |
| Exosc6 | 2,12E-03 | 1,96 | exosome component 6 | No | No | - |
| Nid2 | 3,57E-03 | 1,96 | nidogen 2 | No | No | - |
| Hdac1 | 6,68E-03 | 1,96 | histone deacetylase 1 | No | No | 4 |
| B3galnt1 | 2,40E-02 | 1,96 | UDP-GalNAc:betaGlcNAc beta 1,3-galactosaminyltransfera | No | No | - |
| H1f2 | 2,75E-03 | 1,96 | H1.2 linker histone, cluster member | No | No | - |
| Prps1l3 | 4,83E-02 | 1,96 | phosphoribosyl pyrophosphate synthetase 1-like 3 | No | No | - |
| Dguok | 3,53E-02 | 1,96 | deoxyguanosine kinase | No | No | - |
| Ctsk | 5,50E-03 | 1,96 | cathepsin K | No | No | 2 |

| | | | | | | |
|---|---|---|---|---|---|---|
| Gpr137 | 8,10E-03 | 1,96 | G protein-coupled receptor 137 | No | No | - |
| Setd5 | 4,71E-03 | 1,96 | SET domain containing 5 | No | No | - |
| Dnajb14 | 1,07E-02 | 1,96 | DnaJ heat shock protein family (Hsp40) member B14 | No | No | - |
| Paip1 | 8,77E-03 | 1,96 | polyadenylate binding protein-interacting protein 1 | No | No | - |
| Rpl10-ps3 | 6,11E-04 | 1,96 | ribosomal protein L10, pseudogene 3 | No | No | - |
| Glod4 | 4,26E-03 | 1,97 | glyoxalase domain containing 4 | No | No | - |
| Pmpcb | 1,37E-03 | 1,97 | peptidase (mitochondrial processing) beta | No | No | - |
| Echs1 | 3,16E-03 | 1,97 | enoyl Coenzyme A hydratase, short chain, 1, mitochondrial | No | No | - |
| Krr1 | 5,75E-03 | 1,97 | KRR1, small subunit (SSU) processome component, homol | No | No | - |
| Rassf3 | 1,21E-02 | 1,97 | Ras association (RalGDS/AF-6) domain family member 3 | No | No | - |
| Pdcd5 | 7,47E-03 | 1,97 | programmed cell death 5 | No | No | 2 |
| Ch25h | 3,39E-02 | 1,97 | cholesterol 25-hydroxylase | No | No | 1 |
| Bcl7b | 1,54E-02 | 1,97 | B cell CLL/lymphoma 7B | No | No | - |
| Pgrmc1 | 1,24E-02 | 1,97 | progesterone receptor membrane component 1 | No | No | - |
| Pnrc1 | 4,01E-02 | 1,97 | proline-rich nuclear receptor coactivator 1 | No | No | - |
| Wdr45b | 2,21E-03 | 1,98 | WD repeat domain 45B | No | No | - |
| Gm20390 | 2,51E-05 | 1,98 | predicted gene 20390 | No | No | - |
| Naca | 5,51E-04 | 1,98 | nascent polypeptide-associated complex alpha polypeptide | No | No | - |
| 1110065P20Rik | 9,58E-04 | 1,98 | RIKEN cDNA 1110065P20 gene | No | No | - |
| Myl12a | 9,22E-06 | 1,98 | myosin, light chain 12A, regulatory, non-sarcomeric | No | No | - |
| Tusc2 | 1,47E-02 | 1,98 | tumor suppressor 2, mitochondrial calcium regulator | No | No | - |
| Gtf2h5 | 6,40E-04 | 1,98 | general transcription factor IIH, polypeptide 5 | No | No | - |
| Tra2b | 3,66E-03 | 1,98 | transformer 2 beta | No | No | - |
| H2az2 | 4,32E-03 | 1,98 | H2A.Z histone variant 2 | No | No | - |
| Suclg1 | 1,55E-03 | 1,98 | succinate-CoA ligase, GDP-forming, alpha subunit | No | No | - |
| Marcksl1 | 3,56E-03 | 1,98 | MARCKS-like 1 | No | No | - |
| Psme2 | 8,66E-03 | 1,98 | proteasome (prosome, macropain) activator subunit 2 (PA2 | No | No | - |
| Trappc4 | 6,97E-05 | 1,98 | trafficking protein particle complex 4 | No | No | - |
| Tmem230 | 6,79E-03 | 1,98 | transmembrane protein 230 | No | No | - |
| H2ax | 2,59E-02 | 1,99 | H2A.X variant histone | No | No | 3 |
| Exosc1 | 2,64E-03 | 1,99 | exosome component 1 | No | No | - |
| Arpc5 | 4,75E-05 | 1,99 | actin related protein 2/3 complex, subunit 5 | No | No | - |
| Gm20488 | 1,25E-02 | 1,99 | predicted gene 20488 | No | No | - |
| Snhg1 | 2,94E-03 | 1,99 | small nucleolar RNA host gene 1 | No | No | - |
| Pja1 | 6,65E-03 | 1,99 | praja ring finger ubiquitin ligase 1 | No | No | - |
| Ncbp2 | 1,85E-03 | 2,00 | nuclear cap binding protein subunit 2 | No | No | - |
| Tmem219 | 2,30E-03 | 2,00 | transmembrane protein 219 | No | No | 1 |
| Tceal8 | 1,41E-04 | 2,00 | transcription elongation factor A (SII)-like 8 | No | No | - |
| Ccbe1 | 9,31E-03 | 2,00 | collagen and calcium binding EGF domains 1 | No | No | - |
| Spry2 | 5,06E-03 | 2,00 | sprouty RTK signaling antagonist 2 | No | No | 3 |
| Ostf1 | 7,41E-05 | 2,00 | osteoclast stimulating factor 1 | No | No | - |
| Rpl10a-ps1 | 9,97E-06 | 2,00 | ribosomal protein L10A, pseudogene 1 | No | No | - |
| Tpd52 | 1,21E-02 | 2,01 | tumor protein D52 | No | No | - |
| AC163703.1 | 2,88E-05 | 2,01 | - | No | No | - |
| Ier3ip1 | 4,79E-04 | 2,01 | immediate early response 3 interacting protein 1 | No | No | - |
| Rspry1 | 2,02E-02 | 2,01 | ring finger and SPRY domain containing 1 | No | No | - |
| Dnajc14 | 3,72E-03 | 2,01 | DnaJ heat shock protein family (Hsp40) member C14 | No | No | - |
| Rps17 | 1,25E-05 | 2,02 | ribosomal protein S17 | No | No | - |
| Ndufc1 | 1,57E-05 | 2,02 | NADH:ubiquinone oxidoreductase subunit C1 | No | No | - |
| Zfyve21 | 3,16E-03 | 2,02 | zinc finger, FYVE domain containing 21 | No | No | - |
| Arhgef10l | 2,88E-02 | 2,02 | Rho guanine nucleotide exchange factor (GEF) 10-like | No | No | - |
| Rab11a | 8,93E-04 | 2,02 | RAB11A, member RAS oncogene family | No | No | - |
| Sertad3 | 3,18E-02 | 2,02 | SERTA domain containing 3 | No | No | - |
| Hacd1 | 5,35E-04 | 2,02 | 3-hydroxyacyl-CoA dehydratase 1 | No | No | - |
| Nsg1 | 3,89E-03 | 2,02 | neuron specific gene family member 1 | No | No | - |
| Mrpl34 | 1,05E-03 | 2,02 | mitochondrial ribosomal protein L34 | No | No | - |
| Dcst2 | 4,32E-02 | 2,02 | DC-STAMP domain containing 2 | No | No | - |
| Dhrs7b | 9,04E-03 | 2,02 | dehydrogenase/reductase (SDR family) member 7B | No | No | - |
| Pdcd6 | 1,89E-03 | 2,02 | programmed cell death 6 | No | No | - |
| Ensa | 1,51E-02 | 2,02 | endosulfine alpha | No | No | - |
| Vamp4 | 8,48E-03 | 2,02 | vesicle-associated membrane protein 4 | No | No | 1 |
| Gm11249 | 1,84E-02 | 2,02 | predicted gene 11249 | No | No | - |
| Rtraf | 2,55E-03 | 2,02 | RNA transcription, translation and transport factor | No | No | - |
| Ndufaf4 | 2,87E-03 | 2,02 | NADH:ubiquinone oxidoreductase complex assembly facto | No | No | - |
| A930015D03Rik | 1,90E-02 | 2,02 | RIKEN cDNA A930015D03 gene | No | No | - |
| Cox19 | 4,40E-03 | 2,02 | cytochrome c oxidase assembly protein 19 | No | No | - |
| Cox8a | 7,86E-06 | 2,03 | cytochrome c oxidase subunit 8A | No | No | - |
| Yaf2 | 6,32E-03 | 2,03 | YY1 associated factor 2 | No | No | - |
| Gm21974 | 2,46E-03 | 2,03 | predicted gene 21974 | No | No | - |
| Tomm20l | 5,55E-03 | 2,03 | translocase of outer mitochondrial membrane 20-like | No | No | - |
| Med21 | 2,29E-02 | 2,03 | mediator complex subunit 21 | No | No | - |
| Znrf2 | 2,56E-02 | 2,03 | zinc and ring finger 2 | No | No | - |
| Rfc3 | 2,87E-02 | 2,03 | replication factor C (activator 1) 3 | No | No | - |

| | | | | | | |
|---|---|---|---|---|---|---|
| Cox6c | 1,73E-05 | 2,03 | cytochrome c oxidase subunit 6C | No | No | - |
| Ptma | 9,36E-03 | 2,03 | prothymosin alpha | No | No | - |
| Commd10 | 2,66E-02 | 2,03 | COMM domain containing 10 | No | No | - |
| Dpf2 | 2,55E-02 | 2,03 | D4, zinc and double PHD fingers family 2 | No | No | - |
| Gm7611 | 1,55E-02 | 2,03 | predicted gene 7611 | No | No | - |
| Prpf38a | 4,14E-02 | 2,04 | PRP38 pre-mRNA processing factor 38 (yeast) domain cont | No | No | - |
| Snca | 4,31E-02 | 2,04 | synuclein, alpha | No | No | - |
| Lyar | 3,94E-02 | 2,04 | Ly1 antibody reactive clone | No | No | - |
| Ribc1 | 5,10E-03 | 2,04 | RIB43A domain with coiled-coils 1 | No | No | - |
| Psmb5 | 2,30E-03 | 2,04 | proteasome (prosome, macropain) subunit, beta type 5 | No | No | - |
| Hmbs | 1,81E-02 | 2,04 | hydroxymethylbilane synthase | No | No | - |
| Supt20 | 1,13E-02 | 2,04 | SPT20 SAGA complex component | No | No | - |
| Mnat1 | 1,01E-02 | 2,04 | menage a trois 1 | No | No | - |
| Anapc16 | 3,52E-03 | 2,04 | anaphase promoting complex subunit 16 | No | No | - |
| Get1 | 3,18E-02 | 2,04 | guided entry of tail-anchored proteins factor 1 | No | No | - |
| Utp23 | 1,08E-02 | 2,04 | UTP23 small subunit processome component | No | No | - |
| Nme2 | 1,11E-05 | 2,04 | NME/NM23 nucleoside diphosphate kinase 2 | No | No | - |
| Calm3 | 2,55E-03 | 2,05 | calmodulin 3 | No | No | - |
| Gm43118 | 8,44E-03 | 2,05 | predicted gene 43118 | No | No | - |
| Commd5 | 6,96E-03 | 2,05 | COMM domain containing 5 | No | No | - |
| Mex3a | 3,93E-02 | 2,05 | mex3 RNA binding family member A | No | No | - |
| Mcts1 | 8,46E-04 | 2,05 | malignant T cell amplified sequence 1 | No | No | - |
| Tyms | 4,67E-03 | 2,05 | thymidylate synthase | No | No | - |
| Ndufa7 | 9,76E-06 | 2,05 | NADH:ubiquinone oxidoreductase subunit A7 | No | No | - |
| Hotairm1 | 2,42E-03 | 2,05 | Hoxa transcript antisense RNA, myeloid-specific 1 | No | No | - |
| Ndufa12 | 1,96E-03 | 2,05 | NADH:ubiquinone oxidoreductase subunit A12 | No | No | - |
| Ydjc | 1,72E-02 | 2,05 | YdjC homolog (bacterial) | No | No | - |
| Pkig | 3,68E-03 | 2,05 | protein kinase inhibitor, gamma | No | No | - |
| Cstf2t | 2,33E-03 | 2,05 | cleavage stimulation factor, 3' pre-RNA subunit 2, tau | No | No | - |
| Sgpp1 | 8,31E-03 | 2,05 | sphingosine-1-phosphate phosphatase 1 | No | No | - |
| Hdac2 | 1,04E-02 | 2,05 | histone deacetylase 2 | No | No | 32 |
| Mir22hg | 4,79E-03 | 2,05 | Mir22 host gene (non-protein coding) | No | No | - |
| Mms22l | 1,08E-02 | 2,06 | MMS22-like, DNA repair protein | No | No | - |
| D030056L22Rik | 2,43E-02 | 2,06 | RIKEN cDNA D030056L22 gene | No | No | - |
| Cyb5a | 8,74E-04 | 2,06 | cytochrome b5 type A (microsomal) | No | No | - |
| Eif1b | 1,42E-03 | 2,06 | eukaryotic translation initiation factor 1B | No | No | - |
| Ndufb2 | 2,04E-04 | 2,06 | NADH:ubiquinone oxidoreductase subunit B2 | No | No | - |
| Scrn3 | 3,85E-02 | 2,07 | secernin 3 | No | No | - |
| Bag2 | 7,08E-03 | 2,07 | BCL2-associated athanogene 2 | No | No | - |
| Rpl18 | 4,42E-04 | 2,07 | ribosomal protein L18 | No | No | - |
| 2810025M15Rik | 2,08E-02 | 2,07 | RIKEN cDNA 2810025M15 gene | No | No | - |
| Prr14 | 3,80E-02 | 2,07 | proline rich 14 | No | No | - |
| Coq2 | 1,31E-02 | 2,07 | coenzyme Q2 4-hydroxybenzoate polyprenyltransferase | No | No | - |
| Plpp5 | 8,28E-03 | 2,07 | phospholipid phosphatase 5 | No | No | - |
| Ypel5 | 8,74E-04 | 2,07 | yippee like 5 | No | No | - |
| 2410006H16Rik | 1,64E-04 | 2,07 | RIKEN cDNA 2410006H16 gene | No | No | - |
| Trmt112 | 1,86E-03 | 2,07 | tRNA methyltransferase 11-2 | No | No | - |
| Prdx4 | 1,29E-04 | 2,08 | peroxiredoxin 4 | No | No | 13 |
| Bloc1s3 | 3,32E-02 | 2,08 | biogenesis of lysosomal organelles complex-1, subunit 3 | No | No | 2 |
| Wdr38 | 1,89E-03 | 2,08 | WD repeat domain 38 | No | No | - |
| Tomm22 | 1,12E-03 | 2,08 | translocase of outer mitochondrial membrane 22 | No | No | - |
| Gm11451 | 3,76E-02 | 2,08 | predicted gene 11451 | No | No | - |
| Mzt1 | 9,58E-03 | 2,08 | mitotic spindle organizing protein 1 | No | No | - |
| Ddit4l | 1,19E-02 | 2,08 | DNA-damage-inducible transcript 4-like | No | No | 4 |
| Gm11263 | 6,86E-03 | 2,09 | predicted gene 11263 | No | No | - |
| Tmed7 | 1,87E-02 | 2,09 | transmembrane p24 trafficking protein 7 | No | No | - |
| Ryk | 1,74E-03 | 2,09 | receptor-like tyrosine kinase | No | No | - |
| Pex13 | 3,11E-02 | 2,09 | peroxisomal biogenesis factor 13 | No | No | 1 |
| Gm16580 | 3,62E-02 | 2,09 | predicted gene 16580 | No | No | - |
| Frzb | 1,64E-03 | 2,09 | frizzled-related protein | No | No | 4 |
| Cmc1 | 6,77E-03 | 2,09 | COX assembly mitochondrial protein 1 | No | No | - |
| Idh2 | 7,44E-03 | 2,09 | isocitrate dehydrogenase 2 (NADP+), mitochondrial | No | No | 2 |
| Txnl4a | 1,01E-02 | 2,09 | thioredoxin-like 4A | No | No | - |
| Aurkaip1 | 4,91E-05 | 2,09 | aurora kinase A interacting protein 1 | No | No | - |
| Gm49356 | 6,25E-03 | 2,09 | predicted gene, 49356 | No | No | - |
| Ubtf | 8,99E-03 | 2,09 | upstream binding transcription factor, RNA polymerase I | No | No | - |
| Emc4 | 4,89E-02 | 2,10 | ER membrane protein complex subunit 4 | No | No | - |
| Rbbp9 | 9,58E-03 | 2,10 | retinoblastoma binding protein 9, serine hydrolase | No | No | - |
| Stard10 | 4,42E-02 | 2,10 | START domain containing 10 | No | No | - |
| Cbx4 | 2,69E-02 | 2,10 | chromobox 4 | No | No | - |
| Nqo2 | 1,75E-02 | 2,10 | N-ribosyldihydronicotinamide quinone reductase 2 | No | No | - |
| Cnot7 | 1,45E-03 | 2,10 | CCR4-NOT transcription complex, subunit 7 | No | No | - |
| Arf4 | 9,76E-06 | 2,10 | ADP-ribosylation factor 4 | No | No | - |

| | | | | | | |
|---|---|---|---|---|---|---|
| Ssr2 | 1,16E-04 | 2,10 | signal sequence receptor, beta | No | No | - |
| Timm9 | 3,34E-03 | 2,10 | translocase of inner mitochondrial membrane 9 | No | No | - |
| Chmp1b | 4,28E-03 | 2,10 | charged multivesicular body protein 1B | No | No | - |
| Dqx1 | 4,50E-02 | 2,10 | DEAQ RNA-dependent ATPase | No | No | - |
| Tra2a | 7,48E-03 | 2,10 | transformer 2 alpha | No | No | - |
| Trp53rkb | 4,63E-02 | 2,11 | transformation related protein 53 regulating kinase B | No | No | - |
| Oaz1 | 2,21E-04 | 2,11 | ornithine decarboxylase antizyme 1 | No | No | - |
| Rab15 | 2,97E-02 | 2,11 | RAB15, member RAS oncogene family | No | No | - |
| Esrra | 1,66E-03 | 2,11 | estrogen related receptor, alpha | No | No | - |
| Exoc3l4 | 3,65E-03 | 2,11 | exocyst complex component 3-like 4 | No | No | - |
| Kti12 | 3,48E-02 | 2,11 | KTI12 homolog, chromatin associated | No | No | - |
| Med7 | 7,83E-03 | 2,11 | mediator complex subunit 7 | No | No | - |
| C1qtnf12 | 2,80E-02 | 2,11 | C1q and tumor necrosis factor related 12 | No | No | - |
| C9orf72 | 1,17E-02 | 2,11 | C9orf72, member of C9orf72-SMCR8 complex | No | No | - |
| Gm6863 | 9,15E-05 | 2,11 | predicted gene 6863 | No | No | - |
| C1qtnf7 | 3,70E-02 | 2,11 | C1q and tumor necrosis factor related protein 7 | No | No | - |
| C330018D20Rik | 4,30E-02 | 2,11 | RIKEN cDNA C330018D20 gene | No | No | - |
| Lzts2 | 1,43E-02 | 2,11 | leucine zipper, putative tumor suppressor 2 | No | No | - |
| Chmp5 | 3,55E-06 | 2,11 | charged multivesicular body protein 5 | No | No | - |
| Zeb2os | 3,83E-02 | 2,12 | zinc finger E-box binding homeobox 2, opposite strand | No | No | - |
| Ggh | 5,10E-03 | 2,12 | gamma-glutamyl hydrolase | No | No | - |
| Tmem126a | 1,21E-03 | 2,12 | transmembrane protein 126A | No | No | - |
| Lage3 | 8,35E-04 | 2,12 | L antigen family, member 3 | No | No | - |
| Armc7 | 3,78E-03 | 2,12 | armadillo repeat containing 7 | No | No | - |
| Syf2 | 1,70E-03 | 2,12 | SYF2 homolog, RNA splicing factor (S. cerevisiae) | No | No | - |
| Dennd5b | 7,04E-03 | 2,12 | DENN/MADD domain containing 5B | No | No | - |
| Alkbh5 | 1,25E-02 | 2,12 | alkB homolog 5, RNA demethylase | No | No | - |
| Gm7666 | 1,03E-02 | 2,12 | predicted pseudogene 7666 | No | No | - |
| Rbms2 | 1,40E-02 | 2,12 | RNA binding motif, single stranded interacting protein 2 | No | No | - |
| Shfl | 4,34E-02 | 2,12 | shiftless antiviral inhibitor of ribosomal frameshifting | No | No | - |
| Ormdl2 | 3,00E-03 | 2,12 | ORM1-like 2 (S. cerevisiae) | No | No | - |
| Ndufv3 | 3,10E-03 | 2,12 | NADH:ubiquinone oxidoreductase core subunit V3 | No | No | - |
| Aurka | 7,71E-03 | 2,13 | aurora kinase A | No | No | - |
| Eif1ax | 2,29E-05 | 2,13 | eukaryotic translation initiation factor 1A, X-linked | No | No | - |
| Txn1 | 1,46E-04 | 2,13 | thioredoxin 1 | No | No | - |
| Cxcl12 | 1,18E-06 | 2,13 | chemokine (C-X-C motif) ligand 12 | No | No | 79 |
| Rsl24d1 | 9,34E-03 | 2,13 | ribosomal L24 domain containing 1 | No | No | - |
| Snai1 | 5,57E-03 | 2,13 | snail family zinc finger 1 | No | No | 12 |
| Mrps21 | 1,38E-02 | 2,13 | mitochondrial ribosomal protein S21 | No | No | - |
| Dot1l | 2,03E-02 | 2,13 | DOT1-like, histone H3 methyltransferase (S. cerevisiae) | No | No | - |
| Mcm7 | 4,43E-02 | 2,13 | minichromosome maintenance complex component 7 | No | No | - |
| Rabif | 8,30E-03 | 2,13 | RAB interacting factor | No | No | - |
| Taf9 | 2,95E-03 | 2,13 | TATA-box binding protein associated factor 9 | No | No | - |
| Gm10275 | 2,92E-04 | 2,13 | predicted pseudogene 10275 | No | No | - |
| Morf4l2 | 3,70E-04 | 2,13 | mortality factor 4 like 2 | No | No | - |
| Lap3 | 6,62E-04 | 2,13 | leucine aminopeptidase 3 | No | No | - |
| Galnt11 | 2,11E-02 | 2,13 | polypeptide N-acetylgalactosaminyltransferase 11 | No | No | - |
| Slc25a17 | 1,92E-02 | 2,14 | solute carrier family 25 (mitochondrial carrier, peroxisomal | No | No | - |
| Sdhc | 2,56E-03 | 2,14 | succinate dehydrogenase complex, subunit C, integral meml | No | No | - |
| 0610010K14Rik | 9,24E-03 | 2,14 | RIKEN cDNA 0610010K14 gene | No | No | - |
| Npr3 | 1,06E-02 | 2,14 | natriuretic peptide receptor 3 | No | No | - |
| Kansl1l | 3,98E-02 | 2,14 | KAT8 regulatory NSL complex subunit 1-like | No | No | - |
| Soga1 | 1,92E-02 | 2,14 | suppressor of glucose, autophagy associated 1 | No | No | - |
| Mcfd2 | 1,52E-03 | 2,14 | multiple coagulation factor deficiency 2 | No | No | - |
| Gng11 | 7,93E-04 | 2,14 | guanine nucleotide binding protein (G protein), gamma 11 | No | No | - |
| 1110038F14Rik | 3,72E-04 | 2,14 | RIKEN cDNA 1110038F14 gene | No | No | - |
| Mrpl50 | 2,00E-03 | 2,14 | mitochondrial ribosomal protein L50 | No | No | - |
| Pyurf | 1,52E-02 | 2,14 | Pigy upstream reading frame | No | No | - |
| C230062I16Rik | 1,47E-03 | 2,15 | RIKEN cDNA C230062I16 gene | No | No | - |
| Ube2g2 | 1,37E-03 | 2,15 | ubiquitin-conjugating enzyme E2G 2 | No | No | - |
| Gosr1 | 4,14E-02 | 2,15 | golgi SNAP receptor complex member 1 | No | No | - |
| Gm4786 | 1,00E-03 | 2,15 | predicted gene 4786 | No | No | - |
| Lgals1 | 7,33E-07 | 2,15 | lectin, galactose binding, soluble 1 | No | No | - |
| Plgrkt | 1,30E-02 | 2,16 | plasminogen receptor, C-terminal lysine transmembrane prc | No | No | - |
| Tpt1 | 2,65E-03 | 2,16 | tumor protein, translationally-controlled 1 | No | No | - |
| Foxred1 | 3,95E-02 | 2,16 | FAD-dependent oxidoreductase domain containing 1 | No | No | - |
| Vdac1 | 3,97E-04 | 2,16 | voltage-dependent anion channel 1 | No | No | - |
| H2-T22 | 8,05E-03 | 2,16 | histocompatibility 2, T region locus 22 | No | No | - |
| Mosmo | 1,21E-02 | 2,16 | modulator of smoothened | No | No | - |
| Entpd7 | 1,27E-02 | 2,16 | ectonucleoside triphosphate diphosphohydrolase 7 | No | No | - |
| Cdc23 | 2,94E-03 | 2,16 | CDC23 cell division cycle 23 | No | No | - |
| Bcl2l2 | 2,56E-02 | 2,16 | BCL2-like 2 | No | No | - |
| Gm16853 | 4,99E-03 | 2,17 | predicted gene, 16853 | No | No | - |

| | | | | | | |
|---|---|---|---|---|---|---|
| Rpl35a | 1,95E-04 | 2,17 | ribosomal protein L35A | No | No | - |
| Mrpl22 | 2,69E-02 | 2,17 | mitochondrial ribosomal protein L22 | No | No | - |
| Rpl35 | 8,99E-04 | 2,17 | ribosomal protein L35 | No | No | - |
| Tomm20 | 3,24E-03 | 2,17 | translocase of outer mitochondrial membrane 20 | No | No | - |
| Yars2 | 2,92E-02 | 2,17 | tyrosyl-tRNA synthetase 2 (mitochondrial) | No | No | - |
| Rap1a | 2,56E-04 | 2,17 | RAS-related protein 1a | No | No | - |
| Mrpl36 | 1,44E-03 | 2,17 | mitochondrial ribosomal protein L36 | No | No | - |
| Sat1 | 4,58E-04 | 2,17 | spermidine/spermine N1-acetyl transferase 1 | No | No | - |
| Socs4 | 4,03E-02 | 2,17 | suppressor of cytokine signaling 4 | No | No | - |
| Armc8 | 1,43E-02 | 2,18 | armadillo repeat containing 8 | No | No | - |
| Lbhd1 | 8,35E-04 | 2,18 | LBH domain containing 1 | No | No | - |
| Kyat1 | 3,94E-02 | 2,18 | kynurenine aminotransferase 1 | No | No | - |
| Tmeff1 | 1,44E-04 | 2,18 | transmembrane protein with EGF-like and two follistatin-lik | No | No | - |
| Guk1 | 2,13E-03 | 2,18 | guanylate kinase 1 | No | No | - |
| Aig1 | 1,93E-03 | 2,18 | androgen-induced 1 | No | No | - |
| Jkamp | 1,07E-03 | 2,18 | JNK1/MAPK8-associated membrane protein | No | No | - |
| Gm6467 | 3,86E-02 | 2,18 | predicted gene 6467 | No | No | - |
| Mettl26 | 1,67E-02 | 2,18 | methyltransferase like 26 | No | No | - |
| Smim4 | 2,44E-05 | 2,18 | small integral membrane protein 4 | No | No | - |
| Slc12a6 | 2,86E-02 | 2,19 | solute carrier family 12, member 6 | No | No | - |
| Dpm1 | 7,13E-03 | 2,19 | dolichol-phosphate (beta-D) mannosyltransferase 1 | No | No | - |
| Apopt1 | 4,60E-02 | 2,19 | cytochrome c oxidase assembly factor 8 | No | No | - |
| Gm10036 | 3,16E-04 | 2,19 | predicted gene 10036 | No | No | - |
| Pex7 | 1,61E-02 | 2,19 | peroxisomal biogenesis factor 7 | No | No | - |
| Rps27 | 1,66E-04 | 2,19 | ribosomal protein S27 | No | No | - |
| Eif4ebp1 | 2,04E-05 | 2,20 | eukaryotic translation initiation factor 4E binding protein 1 | No | No | - |
| Polr2i | 4,55E-03 | 2,20 | polymerase (RNA) II (DNA directed) polypeptide I | No | No | - |
| Lbh | 4,50E-04 | 2,20 | limb-bud and heart | No | No | - |
| Mrpl39 | 3,18E-03 | 2,20 | mitochondrial ribosomal protein L39 | No | No | - |
| Rpe | 2,50E-03 | 2,20 | ribulose-5-phosphate-3-epimerase | No | No | 6 |
| Lta4h | 7,87E-04 | 2,20 | leukotriene A4 hydrolase | No | No | 1 |
| Psmg4 | 3,39E-02 | 2,20 | proteasome (prosome, macropain) assembly chaperone 4 | No | No | - |
| Pdzd9 | 4,65E-05 | 2,21 | PDZ domain containing 9 | No | No | - |
| Krcc1 | 2,41E-04 | 2,21 | lysine-rich coiled-coil 1 | No | No | - |
| Pdgfa | 1,67E-03 | 2,21 | platelet derived growth factor, alpha | No | No | 6 |
| Myl6 | 2,14E-05 | 2,21 | myosin, light polypeptide 6, alkali, smooth muscle and non- | No | No | - |
| Rpl17-ps3 | 1,56E-02 | 2,21 | ribosomal protein L17, pseudogene 3 | No | No | - |
| Thoc6 | 6,57E-04 | 2,21 | THO complex 6 | No | No | - |
| Fgf7 | 2,37E-03 | 2,21 | fibroblast growth factor 7 | No | No | 9 |
| Chic2 | 1,52E-06 | 2,22 | cysteine-rich hydrophobic domain 2 | No | No | - |
| Rbm17 | 2,62E-03 | 2,22 | RNA binding motif protein 17 | No | No | - |
| Rpl23a | 9,06E-07 | 2,22 | ribosomal protein L23A | No | No | - |
| Nipa2 | 2,41E-03 | 2,22 | non imprinted in Prader-Willi/Angelman syndrome 2 homol | No | No | - |
| Nt5c | 1,65E-03 | 2,22 | 5',3'-nucleotidase, cytosolic | No | No | - |
| Ehmt1 | 1,57E-04 | 2,22 | euchromatic histone methyltransferase 1 | No | No | - |
| Car13 | 5,17E-03 | 2,22 | carbonic anhydrase 13 | No | No | - |
| Ppil3 | 3,18E-02 | 2,22 | peptidylprolyl isomerase (cyclophilin)-like 3 | No | No | - |
| Hsd17b10 | 1,89E-03 | 2,22 | hydroxysteroid (17-beta) dehydrogenase 10 | No | No | - |
| Akirin1 | 2,32E-03 | 2,22 | akirin 1 | No | No | - |
| Dhx32 | 2,88E-02 | 2,22 | DEAH (Asp-Glu-Ala-His) box polypeptide 32 | No | No | - |
| Ovol3 | 1,43E-02 | 2,22 | ovo like zinc finger 3 | No | No | - |
| Sdhaf1 | 3,79E-03 | 2,22 | succinate dehydrogenase complex assembly factor 1 | No | No | - |
| Pdzd11 | 2,68E-03 | 2,22 | PDZ domain containing 11 | No | No | - |
| Hoxa3 | 4,04E-04 | 2,23 | homeobox A3 | No | No | - |
| Cbr1 | 8,99E-04 | 2,23 | carbonyl reductase 1 | No | No | - |
| Pdcd2l | 4,28E-03 | 2,23 | programmed cell death 2-like | No | No | - |
| Nup107 | 1,48E-02 | 2,23 | nucleoporin 107 | No | No | - |
| Kctd5 | 7,98E-03 | 2,23 | potassium channel tetramerisation domain containing 5 | No | No | - |
| Bccip | 2,71E-02 | 2,23 | BRCA2 and CDKN1A interacting protein | No | No | - |
| Ak6 | 1,32E-03 | 2,23 | adenylate kinase 6 | No | No | - |
| Mrpl15 | 1,64E-03 | 2,23 | mitochondrial ribosomal protein L15 | No | No | - |
| Znhit3 | 1,70E-02 | 2,23 | zinc finger, HIT type 3 | No | No | - |
| Gid8 | 9,21E-04 | 2,24 | GID complex subunit 8 | No | No | - |
| Tfam | 4,63E-02 | 2,24 | transcription factor A, mitochondrial | No | No | - |
| Gm10175 | 1,60E-05 | 2,24 | predicted gene 10175 | No | No | - |
| Mrps18c | 9,14E-04 | 2,24 | mitochondrial ribosomal protein S18C | No | No | - |
| Osgep | 2,54E-03 | 2,24 | O-sialoglycoprotein endopeptidase | No | No | - |
| Use1 | 6,91E-04 | 2,25 | unconventional SNARE in the ER 1 homolog (S. cerevisiae | No | No | - |
| 1700027A07Rik | 2,94E-02 | 2,25 | RIKEN cDNA 1700027A07 gene | No | No | - |
| Bpnt1 | 1,18E-02 | 2,25 | 3'(2'), 5'-bisphosphate nucleotidase 1 | No | No | - |
| Cnih4 | 3,16E-04 | 2,25 | cornichon family AMPA receptor auxiliary protein 4 | No | No | - |
| Bicdl2 | 4,89E-04 | 2,25 | BICD family like cargo adaptor 2 | No | No | - |
| Snx3 | 2,48E-04 | 2,25 | sorting nexin 3 | No | No | - |

| | | | | | | |
|---|---|---|---|---|---|---|
| Tstd3 | 5,75E-04 | 2,25 | thiosulfate sulfurtransferase (rhodanese)-like domain contai | No | No | - |
| Rhebl1 | 1,11E-02 | 2,26 | Ras homolog enriched in brain like 1 | No | No | - |
| Hspe1 | 4,55E-05 | 2,26 | heat shock protein 1 (chaperonin 10) | No | No | - |
| Trappc1 | 8,69E-04 | 2,26 | trafficking protein particle complex 1 | No | No | - |
| Chrac1 | 7,82E-03 | 2,26 | chromatin accessibility complex 1 | No | No | - |
| Fundc2 | 2,14E-05 | 2,26 | FUN14 domain containing 2 | No | No | - |
| Rpsa | 1,55E-06 | 2,26 | ribosomal protein SA | No | No | - |
| Thyn1 | 3,25E-03 | 2,26 | thymocyte nuclear protein 1 | No | No | - |
| Adh5 | 4,62E-04 | 2,26 | alcohol dehydrogenase 5 (class III), chi polypeptide | No | No | 1 |
| Chchd1 | 1,84E-05 | 2,26 | coiled-coil-helix-coiled-coil-helix domain containing 1 | No | No | - |
| Cox20 | 3,14E-03 | 2,26 | cytochrome c oxidase assembly protein 20 | No | No | - |
| Pycrl | 7,52E-03 | 2,26 | pyrroline-5-carboxylate reductase-like | No | No | - |
| Dennd2d | 3,04E-04 | 2,27 | DENN/MADD domain containing 2D | No | No | - |
| Txndc17 | 1,76E-04 | 2,27 | thioredoxin domain containing 17 | No | No | - |
| Ptgr1 | 3,02E-03 | 2,27 | prostaglandin reductase 1 | No | No | - |
| Tlcd1 | 1,33E-06 | 2,27 | TLC domain containing 1 | No | No | - |
| Zfas1 | 4,97E-05 | 2,27 | zinc finger, NFX1-type containing 1, antisense RNA 1 | No | No | - |
| Actr10 | 4,75E-05 | 2,27 | ARP10 actin-related protein 10 | No | No | - |
| Tpgs1 | 1,31E-02 | 2,27 | tubulin polyglutamylase complex subunit 1 | No | No | - |
| Cdkn1b | 4,91E-04 | 2,27 | cyclin-dependent kinase inhibitor 1B | No | No | - |
| Rpp14 | 2,55E-04 | 2,27 | ribonuclease P 14 subunit | No | No | - |
| Cd47 | 1,13E-03 | 2,27 | CD47 antigen (Rh-related antigen, integrin-associated signa | No | No | 23 |
| Timm17a | 4,79E-04 | 2,27 | translocase of inner mitochondrial membrane 17a | No | No | - |
| Gm8242 | 2,39E-02 | 2,28 | predicted gene 8242 | No | No | - |
| Snapc3 | 4,17E-03 | 2,28 | small nuclear RNA activating complex, polypeptide 3 | No | No | - |
| Snrpd2 | 3,60E-04 | 2,28 | small nuclear ribonucleoprotein D2 | No | No | - |
| Eny2 | 6,17E-05 | 2,28 | ENY2 transcription and export complex 2 subunit | No | No | - |
| Ap1ar | 4,39E-04 | 2,28 | adaptor-related protein complex 1 associated regulatory pro | No | No | - |
| Pi4k2b | 7,94E-03 | 2,28 | phosphatidylinositol 4-kinase type 2 beta | No | No | - |
| Gstcd | 4,18E-02 | 2,28 | glutathione S-transferase, C-terminal domain containing | No | No | - |
| Anp32e | 3,42E-04 | 2,28 | acidic (leucine-rich) nuclear phosphoprotein 32 family, men | No | No | - |
| Rab5if | 4,78E-06 | 2,28 | RAB5 interacting factor | No | No | - |
| Tbca | 1,64E-06 | 2,28 | tubulin cofactor A | No | No | - |
| Mtch2 | 8,34E-04 | 2,28 | mitochondrial carrier 2 | No | No | - |
| Eef1e1 | 2,47E-02 | 2,28 | eukaryotic translation elongation factor 1 epsilon 1 | No | No | - |
| B3galt6 | 6,65E-03 | 2,28 | UDP-Gal:betaGal beta 1,3-galactosyltransferase, polypeptid | No | No | - |
| Lrrc75a | 1,24E-06 | 2,28 | leucine rich repeat containing 75A | No | No | - |
| Tmem218 | 3,89E-03 | 2,28 | transmembrane protein 218 | No | No | - |
| Mageh1 | 6,98E-04 | 2,28 | MAGE family member H1 | No | No | - |
| Coa7 | 3,33E-03 | 2,29 | cytochrome c oxidase assembly factor 7 | No | No | - |
| Fkbp7 | 2,16E-04 | 2,29 | FK506 binding protein 7 | No | No | - |
| Snhg5 | 5,76E-04 | 2,29 | small nucleolar RNA host gene 5 | No | No | - |
| Dse | 3,30E-03 | 2,29 | dermatan sulfate epimerase | No | No | 2 |
| Hadh | 4,39E-04 | 2,29 | hydroxyacyl-Coenzyme A dehydrogenase | No | No | - |
| Atp5md | 1,50E-05 | 2,29 | ATP synthase membrane subunit DAPIT | No | No | - |
| Ftsj1 | 2,11E-02 | 2,29 | FtsJ RNA methyltransferase homolog 1 (E. coli) | No | No | - |
| Ufsp2 | 1,64E-04 | 2,29 | UFM1-specific peptidase 2 | No | No | 2 |
| Polr2g | 5,65E-06 | 2,30 | polymerase (RNA) II (DNA directed) polypeptide G | No | No | - |
| Ippk | 3,78E-02 | 2,30 | inositol 1,3,4,5,6-pentakisphosphate 2-kinase | No | No | - |
| Mpv17 | 8,90E-04 | 2,30 | MpV17 mitochondrial inner membrane protein | No | No | - |
| Glipr1 | 8,35E-04 | 2,30 | GLI pathogenesis-related 1 (glioma) | No | No | - |
| Pde6d | 7,57E-03 | 2,30 | phosphodiesterase 6D, cGMP-specific, rod, delta | No | No | 8 |
| Uqcrc2 | 1,01E-05 | 2,30 | ubiquinol cytochrome c reductase core protein 2 | No | No | - |
| Mrps10 | 3,09E-03 | 2,30 | mitochondrial ribosomal protein S10 | No | No | - |
| Stau2 | 1,20E-02 | 2,30 | staufen double-stranded RNA binding protein 2 | No | No | - |
| Msto1 | 3,04E-02 | 2,30 | misato 1, mitochondrial distribution and morphology regula | No | No | - |
| Ethe1 | 3,91E-03 | 2,30 | ethylmalonic encephalopathy 1 | No | No | - |
| Ndufaf8 | 3,14E-04 | 2,31 | NADH:ubiquinone oxidoreductase complex assembly facto | No | No | - |
| Atg16l2 | 3,14E-04 | 2,31 | autophagy related 16-like 2 (S. cerevisiae) | No | No | 4 |
| Kif22 | 3,66E-03 | 2,31 | kinesin family member 22 | No | No | - |
| Gm12669 | 1,15E-02 | 2,31 | predicted gene 12669 | No | No | - |
| Psmb3 | 8,00E-05 | 2,32 | proteasome (prosome, macropain) subunit, beta type 3 | No | No | - |
| Rps27a | 1,64E-06 | 2,32 | ribosomal protein S27A | No | No | - |
| Kif20a | 9,00E-04 | 2,32 | kinesin family member 20A | No | No | - |
| Tars | 1,88E-05 | 2,32 | threonyl-tRNA synthetase | No | No | - |
| Auh | 2,00E-03 | 2,32 | AU RNA binding protein/enoyl-coenzyme A hydratase | No | No | - |
| Fggy | 3,39E-02 | 2,32 | FGGY carbohydrate kinase domain containing | No | No | - |
| Cdk7 | 8,17E-03 | 2,33 | cyclin-dependent kinase 7 | No | No | - |
| Cenph | 2,72E-02 | 2,33 | centromere protein H | No | No | - |
| Cnrip1 | 1,46E-03 | 2,33 | cannabinoid receptor interacting protein 1 | No | No | - |
| Fdx1 | 1,60E-03 | 2,33 | ferredoxin 1 | No | No | - |
| Hagh | 4,01E-02 | 2,33 | hydroxyacyl glutathione hydrolase | No | No | - |
| Impa2 | 4,25E-02 | 2,33 | inositol monophosphatase 2 | No | No | - |

| | | | | | | |
|---|---|---|---|---|---|---|
| Nsfl1c | 5,51E-04 | 2,33 | NSFL1 (p97) cofactor (p47) | No | No | - |
| Prorsd1 | 1,05E-03 | 2,33 | prolyl-tRNA synthetase domain containing 1 | No | No | - |
| Rce1 | 8,23E-03 | 2,33 | Ras converting CAAX endopeptidase 1 | No | No | - |
| Taf5 | 5,00E-06 | 2,33 | TATA-box binding protein associated factor 5 | No | No | - |
| Tmem45a | 4,41E-04 | 2,34 | transmembrane protein 45a | No | No | - |
| E130218I03Rik | 7,09E-03 | 2,34 | RIKEN cDNA E130218I03 gene | No | No | - |
| Snu13 | 2,75E-05 | 2,34 | SNU13 homolog, small nuclear ribonucleoprotein (U4/U6.U | No | No | - |
| Spag7 | 6,25E-05 | 2,34 | sperm associated antigen 7 | No | No | - |
| Gabarapl2 | 3,12E-03 | 2,34 | gamma-aminobutyric acid (GABA) A receptor-associated p | No | No | - |
| Nudt2 | 1,15E-02 | 2,34 | nudix (nucleoside diphosphate linked moiety X)-type motif | No | No | - |
| Tspan6 | 3,88E-05 | 2,34 | tetraspanin 6 | No | No | - |
| Cdk2 | 1,52E-03 | 2,35 | cyclin-dependent kinase 2 | No | No | 4 |
| Med28 | 7,63E-07 | 2,35 | mediator complex subunit 28 | No | No | - |
| Hmgxb4 | 2,19E-02 | 2,35 | HMG box domain containing 4 | No | No | - |
| Slc37a4 | 1,71E-02 | 2,35 | solute carrier family 37 (glucose-6-phosphate transporter), r | No | No | - |
| 4921524J17Rik | 1,80E-02 | 2,35 | RIKEN cDNA 4921524J17 gene | No | No | - |
| Amacr | 2,94E-02 | 2,35 | alpha-methylacyl-CoA racemase | No | No | - |
| Yipf4 | 5,28E-04 | 2,35 | Yip1 domain family, member 4 | No | No | - |
| Fam122a | 3,24E-03 | 2,35 | family with sequence similarity 122, member A | No | No | - |
| Psmb7 | 8,35E-04 | 2,35 | proteasome (prosome, macropain) subunit, beta type 7 | No | No | - |
| Ndufab1 | 3,20E-06 | 2,35 | NADH:ubiquinone oxidoreductase subunit AB1 | No | No | - |
| Dusp22 | 4,27E-02 | 2,35 | dual specificity phosphatase 22 | No | No | - |
| Gpx4 | 1,69E-02 | 2,36 | glutathione peroxidase 4 | No | No | 7 |
| Serpinb6b | 9,11E-04 | 2,36 | serine (or cysteine) peptidase inhibitor, clade B, member 6b | No | No | - |
| Gm44386 | 1,09E-02 | 2,36 | predicted gene, 44386 | No | No | - |
| Gm49594 | 2,01E-03 | 2,36 | predicted gene, 49594 | No | No | - |
| Cavin3 | 3,84E-04 | 2,36 | caveolae associated 3 | No | No | - |
| Rps2 | 7,63E-07 | 2,36 | ribosomal protein S2 | No | No | - |
| Chn1 | 1,49E-02 | 2,36 | chimerin 1 | No | No | - |
| Has2 | 2,29E-02 | 2,36 | hyaluronan synthase 2 | No | No | 20 |
| Slc35b1 | 1,99E-05 | 2,36 | solute carrier family 35, member B1 | No | No | - |
| Ccnh | 5,00E-04 | 2,36 | cyclin H | No | No | - |
| Fibin | 1,50E-03 | 2,37 | fin bud initiation factor homolog (zebrafish) | No | No | - |
| Dbr1 | 1,76E-02 | 2,37 | debranching RNA lariats 1 | No | No | - |
| Insig2 | 2,23E-04 | 2,37 | insulin induced gene 2 | No | No | - |
| Mtarc2 | 7,20E-04 | 2,37 | mitochondrial amidoxime reducing component 2 | No | No | - |
| Cenpl | 2,24E-02 | 2,37 | centromere protein L | No | No | - |
| Mgat2 | 6,63E-04 | 2,37 | mannoside acetylglucosaminyltransferase 2 | No | No | - |
| Gm11100 | 3,58E-03 | 2,37 | predicted gene 11100 | No | No | - |
| Zfp207 | 2,57E-04 | 2,37 | zinc finger protein 207 | No | No | - |
| Angptl6 | 4,63E-02 | 2,38 | angiopoietin-like 6 | No | No | - |
| Cebpzos | 9,67E-03 | 2,38 | CCAAT/enhancer binding protein (C/EBP), zeta, opposite s | No | No | - |
| Tacc3 | 4,45E-02 | 2,38 | transforming, acidic coiled-coil containing protein 3 | No | No | - |
| Rpl29 | 6,44E-04 | 2,38 | ribosomal protein L29 | No | No | - |
| Pgpep1 | 2,43E-04 | 2,38 | pyroglutamyl-peptidase I | No | No | - |
| Acaa2 | 7,41E-03 | 2,38 | acetyl-Coenzyme A acyltransferase 2 (mitochondrial 3-oxoa | No | No | - |
| Pigbos1 | 7,56E-04 | 2,38 | Pigb opposite strand 1 | No | No | - |
| Hoxb7 | 4,39E-03 | 2,38 | homeobox B7 | No | No | - |
| Psma2 | 2,02E-07 | 2,38 | proteasome subunit alpha 2 | No | No | - |
| Mcl1 | 3,84E-04 | 2,38 | myeloid cell leukemia sequence 1 | No | No | 10 |
| Surf1 | 3,99E-05 | 2,38 | surfeit gene 1 | No | No | - |
| Bub3 | 1,78E-05 | 2,38 | BUB3 mitotic checkpoint protein | No | No | - |
| Fmc1 | 1,15E-04 | 2,38 | formation of mitochondrial complex V assembly factor 1 | No | No | - |
| Glrx | 2,55E-03 | 2,38 | glutaredoxin | No | No | 5 |
| Magef1 | 2,08E-02 | 2,38 | MAGE family member F1 | No | No | - |
| Vps29 | 3,32E-06 | 2,39 | VPS29 retromer complex component | No | No | - |
| Serf1 | 4,22E-04 | 2,39 | small EDRK-rich factor 1 | No | No | - |
| Rpl22 | 2,13E-06 | 2,39 | ribosomal protein L22 | No | No | - |
| Rps2-ps10 | 6,86E-03 | 2,39 | ribosomal protein S2, pseudogene 10 | No | No | - |
| Naa38 | 6,36E-07 | 2,39 | N(alpha)-acetyltransferase 38, NatC auxiliary subunit | No | No | - |
| Elp6 | 3,08E-02 | 2,39 | elongator acetyltransferase complex subunit 6 | No | No | - |
| Hspa1a | 7,71E-03 | 2,39 | heat shock protein 1A | No | No | 5 |
| Rer1 | 2,25E-05 | 2,39 | retention in endoplasmic reticulum sorting receptor 1 | No | No | - |
| Dctn6 | 1,47E-04 | 2,39 | dynactin 6 | No | No | - |
| Gm13889 | 1,31E-02 | 2,39 | predicted gene 13889 | No | No | - |
| Urod | 6,99E-04 | 2,39 | uroporphyrinogen decarboxylase | No | No | - |
| Diablo | 7,02E-04 | 2,39 | diablo, IAP-binding mitochondrial protein | No | No | - |
| Cox5b | 9,29E-07 | 2,40 | cytochrome c oxidase subunit 5B | No | No | - |
| Mad2l1 | 2,57E-02 | 2,40 | MAD2 mitotic arrest deficient-like 1 | No | No | - |
| Sms | 4,25E-02 | 2,40 | spermine synthase | No | No | - |
| 1810009A15Rik | 3,36E-04 | 2,40 | RIKEN cDNA 1810009A15 gene | No | No | - |
| Ddit4 | 3,92E-04 | 2,40 | DNA-damage-inducible transcript 4 | No | No | 1 |
| Fau | 4,92E-05 | 2,40 | Finkel-Biskis-Reilly murine sarcoma virus (FBR-MuSV) ub | No | No | - |

| | | | | | | |
|---|---|---|---|---|---|---|
| Wdr33 | 1,37E-03 | 2,40 | WD repeat domain 33 | No | No | - |
| Cinp | 1,31E-03 | 2,40 | cyclin-dependent kinase 2 interacting protein | No | No | - |
| Pcna | 1,09E-03 | 2,40 | proliferating cell nuclear antigen | No | No | 52 |
| Gls2 | 3,09E-02 | 2,41 | glutaminase 2 (liver, mitochondrial) | No | No | - |
| Snrnp25 | 4,61E-04 | 2,41 | small nuclear ribonucleoprotein 25 (U11/U12) | No | No | - |
| Spc24 | 1,21E-03 | 2,41 | SPC24, NDC80 kinetochore complex component, homolog | No | No | - |
| Ppm1m | 4,46E-03 | 2,41 | protein phosphatase 1M | No | No | - |
| Igbp1 | 1,87E-04 | 2,41 | immunoglobulin (CD79A) binding protein 1 | No | No | - |
| Bod1 | 7,07E-04 | 2,41 | biorientation of chromosomes in cell division 1 | No | No | - |
| Lysmd2 | 3,82E-02 | 2,41 | LysM, putative peptidoglycan-binding, domain containing 2 | No | No | - |
| Gm43439 | 4,40E-02 | 2,42 | predicted gene 43439 | No | No | - |
| Tmem121 | 4,08E-02 | 2,42 | transmembrane protein 121 | No | No | - |
| Nkapd1 | 4,36E-06 | 2,43 | NKAP domain containing 1 | No | No | - |
| Dnaaf2 | 1,10E-02 | 2,43 | dynein, axonemal assembly factor 2 | No | No | - |
| Fkbp2 | 8,69E-03 | 2,43 | FK506 binding protein 2 | No | No | - |
| Nmrk1 | 2,59E-02 | 2,43 | nicotinamide riboside kinase 1 | No | No | - |
| Ogn | 4,15E-02 | 2,43 | osteoglycin | No | No | 9 |
| Pgk1-rs7 | 1,01E-05 | 2,43 | phosphoglycerate kinase-1, related sequence-7 | No | No | - |
| Setd6 | 7,59E-04 | 2,43 | SET domain containing 6 | No | No | - |
| Hk2 | 1,31E-03 | 2,43 | hexokinase 2 | No | No | 6 |
| 2010204K13Rik | 4,42E-03 | 2,43 | RIKEN cDNA 2010204K13 gene | No | No | - |
| Crls1 | 6,28E-04 | 2,43 | cardiolipin synthase 1 | No | No | - |
| Nudt9 | 4,53E-03 | 2,43 | nudix (nucleoside diphosphate linked moiety X)-type motif | No | No | - |
| Apip | 1,04E-03 | 2,44 | APAF1 interacting protein | No | No | - |
| Nudt13 | 4,31E-02 | 2,44 | nudix (nucleoside diphosphate linked moiety X)-type motif | No | No | - |
| Pcmtd2 | 2,74E-02 | 2,44 | protein-L-isoaspartate (D-aspartate) O-methyltransferase do | No | No | - |
| Kif2a | 1,51E-03 | 2,44 | kinesin family member 2A | No | No | - |
| Wtip | 9,75E-05 | 2,44 | WT1 interacting protein | No | No | - |
| Sec11a | 4,34E-07 | 2,44 | SEC11 homolog A, signal peptidase complex subunit | No | No | - |
| Pdf | 1,76E-03 | 2,44 | peptide deformylase (mitochondrial) | No | No | 19 |
| Malsu1 | 5,80E-05 | 2,44 | mitochondrial assembly of ribosomal large subunit 1 | No | No | - |
| Map2k1 | 1,46E-05 | 2,44 | mitogen-activated protein kinase kinase 1 | No | No | 73 |
| Pdzd7 | 5,08E-03 | 2,45 | PDZ domain containing 7 | No | No | - |
| Cebpd | 2,74E-04 | 2,45 | CCAAT/enhancer binding protein (C/EBP), delta | No | No | - |
| Sh3bp1 | 1,32E-02 | 2,45 | SH3-domain binding protein 1 | No | No | - |
| Mpc2 | 2,96E-04 | 2,45 | mitochondrial pyruvate carrier 2 | No | No | - |
| B3gnt4 | 4,66E-04 | 2,45 | UDP-GlcNAc:betaGal beta-1,3-N-acetylglucosaminyltransf | No | No | - |
| Ak4 | 1,46E-04 | 2,45 | adenylate kinase 4 | No | No | - |
| Uckl1 | 6,90E-04 | 2,45 | uridine-cytidine kinase 1-like 1 | No | No | - |
| Gtse1 | 1,14E-02 | 2,45 | G two S phase expressed protein 1 | No | No | - |
| Pbk | 2,80E-02 | 2,45 | PDZ binding kinase | No | No | - |
| Hnrnpf | 1,40E-02 | 2,46 | heterogeneous nuclear ribonucleoprotein F | No | No | - |
| Rpl18-ps2 | 1,05E-02 | 2,46 | ribosomal protein L18, pseudogene 2 | No | No | - |
| Rxylt1 | 6,74E-03 | 2,46 | ribitol xylosyltransferase 1 | No | No | - |
| Fam216a | 2,06E-04 | 2,46 | family with sequence similarity 216, member A | No | No | - |
| 1810026B05Rik | 8,21E-03 | 2,46 | CHD2 adjacent suppressive regulatory RNA | No | No | - |
| Sdf2 | 1,22E-04 | 2,46 | stromal cell derived factor 2 | No | No | - |
| Sec22a | 1,30E-03 | 2,46 | SEC22 homolog A, vesicle trafficking protein | No | No | - |
| Gm28036 | 4,45E-03 | 2,47 | predicted gene, 28036 | No | No | - |
| Nasp | 1,41E-02 | 2,47 | nuclear autoantigenic sperm protein (histone-binding) | No | No | - |
| Suclg2 | 3,08E-02 | 2,47 | succinate-Coenzyme A ligase, GDP-forming, beta subunit | No | No | - |
| Tmem216 | 1,94E-02 | 2,47 | transmembrane protein 216 | No | No | - |
| Rnf34 | 3,05E-04 | 2,47 | ring finger protein 34 | No | No | - |
| Hpf1 | 1,17E-03 | 2,47 | histone PARylation factor 1 | No | No | - |
| Lrrc41 | 3,00E-02 | 2,47 | leucine rich repeat containing 41 | No | No | - |
| Rpl27 | 1,22E-07 | 2,48 | ribosomal protein L27 | No | No | - |
| Med10 | 3,83E-04 | 2,48 | mediator complex subunit 10 | No | No | - |
| Rps2-ps13 | 9,53E-04 | 2,48 | ribosomal protein S2, pseudogene 13 | No | No | - |
| Nop10 | 2,39E-04 | 2,48 | NOP10 ribonucleoprotein | No | No | 8 |
| Aamdc | 1,00E-03 | 2,48 | adipogenesis associated Mth938 domain containing | No | No | - |
| Cdc42se1 | 2,53E-04 | 2,48 | CDC42 small effector 1 | No | No | - |
| Svbp | 2,41E-04 | 2,49 | small vasohibin binding protein | No | No | - |
| Tmem88 | 3,01E-06 | 2,49 | transmembrane protein 88 | No | No | - |
| Atp6v1c2 | 2,57E-03 | 2,50 | ATPase, H+ transporting, lysosomal V1 subunit C2 | No | No | - |
| Bloc1s2 | 3,99E-02 | 2,50 | biogenesis of lysosomal organelles complex-1, subunit 2 | No | No | - |
| Cdk5rap1 | 4,10E-02 | 2,50 | CDK5 regulatory subunit associated protein 1 | No | No | - |
| Cdkn2c | 3,24E-02 | 2,50 | cyclin dependent kinase inhibitor 2C | No | No | - |
| Ebpl | 4,73E-03 | 2,50 | emopamil binding protein-like | No | No | - |
| Gm12919 | 1,08E-02 | 2,50 | predicted gene 12919 | No | No | - |
| Gmpr | 3,97E-02 | 2,50 | guanosine monophosphate reductase | No | No | - |
| Hadhb | 1,80E-03 | 2,50 | hydroxyacyl-CoA dehydrogenase trifunctional multienzyme | No | No | - |
| Hoxb3 | 6,89E-03 | 2,50 | homeobox B3 | No | No | 2 |
| Lmnb2 | 4,36E-02 | 2,50 | lamin B2 | No | No | - |

| | | | | | | |
|---|---|---|---|---|---|---|
| Lrfn4 | 1,65E-03 | 2,50 | leucine rich repeat and fibronectin type III domain containir | No | No | - |
| Tsen2 | 3,44E-02 | 2,50 | tRNA splicing endonuclease subunit 2 | No | No | - |
| Ttyh3 | 1,87E-04 | 2,51 | tweety family member 3 | No | No | - |
| Pcsk7 | 3,10E-05 | 2,51 | proprotein convertase subtilisin/kexin type 7 | No | No | - |
| Rps15-ps3 | 3,87E-03 | 2,51 | ribosomal protein S15, pseudogene 3 | No | No | - |
| 1110038B12Rik | 1,16E-06 | 2,51 | RIKEN cDNA 1110038B12 gene | No | No | - |
| Gpx1 | 9,41E-07 | 2,52 | glutathione peroxidase 1 | No | No | 9 |
| 2310009B15Rik | 5,00E-03 | 2,52 | RIKEN cDNA 2310009B15 gene | No | No | - |
| Tfdp1 | 5,00E-03 | 2,52 | transcription factor Dp 1 | No | No | - |
| Ap2m1-ps | 4,04E-02 | 2,52 | adaptor-related protein complex 2, mu 1 subunit, pseudogen | No | No | - |
| Snapin | 7,81E-06 | 2,52 | SNAP-associated protein | No | No | - |
| Myo19 | 6,59E-03 | 2,52 | myosin XIX | No | No | - |
| Ndfip2 | 6,29E-04 | 2,53 | Nedd4 family interacting protein 2 | No | No | - |
| Agpat5 | 1,07E-04 | 2,53 | 1-acylglycerol-3-phosphate O-acyltransferase 5 (lysophospl | No | No | - |
| Exosc7 | 1,40E-02 | 2,53 | exosome component 7 | No | No | - |
| Psmg2 | 6,80E-04 | 2,53 | proteasome (prosome, macropain) assembly chaperone 2 | No | No | 1 |
| Gm49027 | 4,25E-04 | 2,53 | predicted gene, 49027 | No | No | - |
| Nudcd2 | 3,78E-04 | 2,54 | NudC domain containing 2 | No | No | - |
| Cstf3 | 2,61E-03 | 2,54 | cleavage stimulation factor, 3' pre-RNA, subunit 3 | No | No | - |
| Rpl10a | 1,06E-05 | 2,54 | ribosomal protein L10A | No | No | - |
| Ndufb3 | 2,17E-07 | 2,54 | NADH:ubiquinone oxidoreductase subunit B3 | No | No | - |
| Gm19209 | 3,79E-03 | 2,54 | predicted gene, 19209 | No | No | - |
| Rps14 | 6,42E-08 | 2,54 | ribosomal protein S14 | No | No | - |
| Uqcrb | 4,81E-05 | 2,54 | ubiquinol-cytochrome c reductase binding protein | No | No | - |
| Ube2c | 3,99E-04 | 2,55 | ubiquitin-conjugating enzyme E2C | No | No | - |
| Hoxb6 | 4,53E-03 | 2,55 | homeobox B6 | No | No | - |
| Slc25a30 | 8,90E-04 | 2,55 | solute carrier family 25, member 30 | No | No | - |
| Calm2 | 4,31E-05 | 2,55 | calmodulin 2 | No | No | - |
| Hnrnpc | 4,31E-07 | 2,55 | heterogeneous nuclear ribonucleoprotein C | No | No | - |
| Pigf | 3,57E-03 | 2,55 | phosphatidylinositol glycan anchor biosynthesis, class F | No | No | - |
| Psmc3ip | 1,91E-02 | 2,55 | proteasome (prosome, macropain) 26S subunit, ATPase 3, i | No | No | - |
| Slc30a7 | 9,44E-05 | 2,55 | solute carrier family 30 (zinc transporter), member 7 | No | No | - |
| Rpl7a | 9,19E-06 | 2,56 | ribosomal protein L7A | No | No | - |
| Rps18 | 1,56E-06 | 2,56 | ribosomal protein S18 | No | No | - |
| Thap11 | 3,22E-05 | 2,56 | THAP domain containing 11 | No | No | - |
| Ndufb5 | 2,35E-07 | 2,56 | NADH:ubiquinone oxidoreductase subunit B5 | No | No | - |
| Orc6 | 3,08E-02 | 2,56 | origin recognition complex, subunit 6 | No | No | - |
| Pcp2 | 5,09E-03 | 2,56 | Purkinje cell protein 2 (L7) | No | No | - |
| Ilf2 | 9,74E-07 | 2,56 | interleukin enhancer binding factor 2 | No | No | 1 |
| Rad1 | 5,24E-03 | 2,57 | RAD1 checkpoint DNA exonuclease | No | No | - |
| A430018G15Rik | 6,59E-03 | 2,57 | RIKEN cDNA A430018G15 gene | No | No | - |
| Hint3 | 5,79E-03 | 2,57 | histidine triad nucleotide binding protein 3 | No | No | - |
| Eif4a1 | 1,66E-06 | 2,57 | eukaryotic translation initiation factor 4A1 | No | No | - |
| Fam173a | 4,95E-05 | 2,57 | adenine nucleotide translocase lysine methyltransferase | No | No | - |
| Uchl3 | 2,56E-03 | 2,58 | ubiquitin carboxyl-terminal esterase L3 (ubiquitin thiolester | No | No | - |
| Ckap2 | 6,61E-04 | 2,58 | cytoskeleton associated protein 2 | No | No | - |
| Calu | 6,35E-06 | 2,58 | calumenin | No | No | - |
| Psat1 | 1,85E-05 | 2,58 | phosphoserine aminotransferase 1 | No | No | 9 |
| Gm50367 | 4,59E-05 | 2,59 | predicted gene, 50367 | No | No | - |
| Polr1c | 5,83E-04 | 2,59 | polymerase (RNA) I polypeptide C | No | No | - |
| Smim14 | 3,07E-08 | 2,59 | small integral membrane protein 14 | No | No | - |
| Fam118a | 1,59E-02 | 2,59 | family with sequence similarity 118, member A | No | No | - |
| Alyref | 9,10E-05 | 2,59 | Aly/REF export factor | No | No | - |
| Zbtb25 | 4,04E-02 | 2,59 | zinc finger and BTB domain containing 25 | No | No | - |
| Gatad1 | 3,50E-06 | 2,59 | GATA zinc finger domain containing 1 | No | No | - |
| Sf3b5 | 1,56E-06 | 2,59 | splicing factor 3b, subunit 5 | No | No | - |
| Ccdc90b | 3,22E-03 | 2,59 | coiled-coil domain containing 90B | No | No | - |
| Anapc10 | 3,94E-02 | 2,60 | anaphase promoting complex subunit 10 | No | No | - |
| Rmdn1 | 3,86E-02 | 2,60 | regulator of microtubule dynamics 1 | No | No | - |
| Snhg20 | 3,24E-02 | 2,60 | small nucleolar RNA host gene 20 | No | No | - |
| Bcdin3d | 2,29E-02 | 2,61 | BCDIN3 domain containing | No | No | - |
| Wrnip1 | 4,95E-03 | 2,61 | Werner helicase interacting protein 1 | No | No | - |
| Acp1 | 2,76E-06 | 2,61 | acid phosphatase 1, soluble | No | No | - |
| Sptssa | 7,91E-04 | 2,61 | serine palmitoyltransferase, small subunit A | No | No | - |
| Sla2 | 1,77E-07 | 2,61 | Src-like-adaptor 2 | No | No | - |
| Ranbp1 | 3,00E-04 | 2,61 | RAN binding protein 1 | No | No | - |
| Bub1b | 1,78E-02 | 2,62 | BUB1B, mitotic checkpoint serine/threonine kinase | No | No | - |
| Ccdc28a | 3,16E-02 | 2,62 | coiled-coil domain containing 28A | No | No | - |
| Tmem60 | 2,60E-03 | 2,62 | transmembrane protein 60 | No | No | - |
| Dusp7 | 2,26E-03 | 2,62 | dual specificity phosphatase 7 | No | No | - |
| Snrpb2 | 5,09E-06 | 2,62 | U2 small nuclear ribonucleoprotein B | No | No | - |
| Ddo | 6,89E-03 | 2,63 | D-aspartate oxidase | No | No | - |
| Timm8b | 1,56E-06 | 2,63 | translocase of inner mitochondrial membrane 8B | No | No | - |

| | | | | | | |
|---|---|---|---|---|---|---|
| Gm49353 | 4,37E-02 | 2,63 | predicted gene, 49353 | No | No | - |
| Atp5j | 1,12E-04 | 2,63 | ATP synthase, H+ transporting, mitochondrial F0 complex, | No | No | - |
| Nme7 | 3,23E-02 | 2,63 | NME/NM23 family member 7 | No | No | - |
| Rad54l | 2,81E-02 | 2,63 | RAD54 like (S. cerevisiae) | No | No | - |
| Smim11 | 1,22E-04 | 2,63 | small integral membrane protein 11 | No | No | - |
| Ccdc78 | 7,54E-06 | 2,64 | coiled-coil domain containing 78 | No | No | - |
| Shld1 | 4,37E-02 | 2,64 | shieldin complex subunit 1 | No | No | - |
| Pafah1b3 | 9,97E-08 | 2,64 | platelet-activating factor acetylhydrolase, isoform 1b, subun | No | No | - |
| Cpne1 | 3,30E-03 | 2,64 | copine I | No | No | - |
| Hoxb2 | 6,48E-03 | 2,64 | homeobox B2 | No | No | - |
| Tmem59l | 1,40E-02 | 2,65 | transmembrane protein 59-like | No | No | - |
| Cisd2 | 6,67E-05 | 2,65 | CDGSH iron sulfur domain 2 | No | No | - |
| Atp5mpl | 2,41E-08 | 2,65 | ATP synthase membrane subunit 6.8PL | No | No | - |
| Tmem168 | 6,29E-04 | 2,65 | transmembrane protein 168 | No | No | - |
| Opn1sw | 4,17E-06 | 2,65 | opsin 1 (cone pigments), short-wave-sensitive (color blindn | No | No | - |
| Lsm12 | 3,17E-03 | 2,65 | LSM12 homolog | No | No | - |
| Rpsa-ps10 | 8,76E-05 | 2,65 | ribosomal protein SA, pseudogene 10 | No | No | - |
| Mettl9 | 8,48E-05 | 2,65 | methyltransferase like 9 | No | No | - |
| Rrp36 | 8,98E-03 | 2,65 | ribosomal RNA processing 36 | No | No | - |
| Lactb2 | 7,66E-03 | 2,66 | lactamase, beta 2 | No | No | - |
| Atp6v0e | 9,22E-08 | 2,66 | ATPase, H+ transporting, lysosomal V0 subunit E | No | No | - |
| Gskip | 1,07E-03 | 2,66 | GSK3B interacting protein | No | No | - |
| Psmb2 | 2,26E-04 | 2,66 | proteasome (prosome, macropain) subunit, beta type 2 | No | No | 3 |
| Sdhaf4 | 3,63E-04 | 2,66 | succinate dehydrogenase complex assembly factor 4 | No | No | - |
| Cox14 | 2,92E-07 | 2,66 | cytochrome c oxidase assembly protein 14 | No | No | - |
| Mrps24 | 1,89E-04 | 2,66 | mitochondrial ribosomal protein S24 | No | No | - |
| Cnpy2 | 1,92E-05 | 2,67 | canopy FGF signaling regulator 2 | No | No | - |
| Rpl34 | 2,00E-08 | 2,67 | ribosomal protein L34 | No | No | - |
| Gm20604 | 1,58E-02 | 2,67 | predicted gene 20604 | No | No | - |
| Nutf2 | 3,73E-02 | 2,67 | nuclear transport factor 2 | No | No | - |
| Cept1 | 4,54E-06 | 2,67 | choline/ethanolaminephosphotransferase 1 | No | No | - |
| 5730455P16Rik | 1,97E-03 | 2,67 | RIKEN cDNA 5730455P16 gene | No | No | - |
| Dcun1d5 | 3,19E-06 | 2,67 | DCN1, defective in cullin neddylation 1, domain containing | No | No | - |
| Dnmt3l | 9,61E-03 | 2,67 | DNA (cytosine-5-)-methyltransferase 3-like | No | No | - |
| Slc39a1 | 1,28E-04 | 2,68 | solute carrier family 39 (zinc transporter), member 1 | No | No | - |
| Emc6 | 2,05E-06 | 2,68 | ER membrane protein complex subunit 6 | No | No | - |
| Gm5559 | 5,21E-04 | 2,68 | predicted gene 5559 | No | No | - |
| Plaat3 | 2,31E-06 | 2,69 | phospholipase A and acyltransferase 3 | No | No | - |
| Bcl10 | 2,89E-03 | 2,69 | B cell leukemia/lymphoma 10 | No | No | - |
| Pirb | 1,28E-03 | 2,69 | paired Ig-like receptor B | No | No | - |
| Sumo1 | 5,49E-05 | 2,69 | small ubiquitin-like modifier 1 | No | No | - |
| Rpl7a-ps5 | 1,38E-04 | 2,70 | ribosomal protein L7A, pseudogene 5 | No | No | - |
| Mcm9 | 1,63E-02 | 2,70 | minichromosome maintenance 9 homologous recombinatio | No | No | - |
| Tagln | 1,35E-05 | 2,71 | transgelin | Yes | No | 4 |
| Rbm3 | 1,81E-04 | 2,71 | RNA binding motif (RNP1, RRM) protein 3 | No | No | - |
| Hmgb1 | 2,57E-02 | 2,71 | high mobility group box 1 | No | No | - |
| Timmdc1 | 6,97E-04 | 2,71 | translocase of inner mitochondrial membrane domain contai | No | No | - |
| Prim2 | 2,73E-03 | 2,71 | DNA primase, p58 subunit | No | No | - |
| Cisd1 | 2,62E-06 | 2,71 | CDGSH iron sulfur domain 1 | No | No | - |
| Ndufv2 | 1,42E-04 | 2,71 | NADH:ubiquinone oxidoreductase core subunit V2 | No | No | - |
| Ndufb4 | 4,21E-06 | 2,71 | NADH:ubiquinone oxidoreductase subunit B4 | No | No | - |
| 2310010J17Rik | 1,38E-02 | 2,71 | RIKEN cDNA 2310010J17 gene | No | No | - |
| Actr3b | 1,01E-02 | 2,71 | ARP3 actin-related protein 3B | No | No | - |
| H1f0 | 1,24E-05 | 2,71 | H1.0 linker histone | No | No | - |
| Mxi1 | 7,04E-03 | 2,72 | MAX interactor 1, dimerization protein | No | No | - |
| Sin3b | 3,98E-08 | 2,72 | transcriptional regulator, SIN3B (yeast) | No | No | - |
| Ms4a4d | 8,26E-04 | 2,72 | membrane-spanning 4-domains, subfamily A, member 4D | No | No | - |
| Vdac3 | 4,31E-07 | 2,72 | voltage-dependent anion channel 3 | No | No | - |
| Rpl6 | 2,96E-05 | 2,72 | ribosomal protein L6 | No | No | - |
| Bckdhb | 6,49E-03 | 2,73 | branched chain ketoacid dehydrogenase E1, beta polypeptid | No | No | - |
| Cda | 2,81E-02 | 2,73 | cytidine deaminase | No | No | 9 |
| Atp1b1 | 2,66E-02 | 2,73 | ATPase, Na+/K+ transporting, beta 1 polypeptide | No | No | - |
| Btbd1 | 2,75E-03 | 2,73 | BTB (POZ) domain containing 1 | No | No | - |
| Coq3 | 3,10E-05 | 2,73 | coenzyme Q3 methyltransferase | No | No | - |
| Nfil3 | 7,84E-03 | 2,73 | nuclear factor, interleukin 3, regulated | No | No | - |
| Mrpl11 | 3,75E-05 | 2,73 | mitochondrial ribosomal protein L11 | No | No | - |
| Pgk1 | 3,48E-07 | 2,74 | phosphoglycerate kinase 1 | No | No | - |
| Mrpl27 | 2,48E-03 | 2,74 | mitochondrial ribosomal protein L27 | No | No | - |
| Qtrt1 | 2,07E-02 | 2,74 | queuine tRNA-ribosyltransferase catalytic subunit 1 | No | No | - |
| Gm6252 | 2,90E-02 | 2,75 | predicted gene 6252 | No | No | - |
| Gpalpp1 | 1,72E-02 | 2,75 | GPALPP motifs containing 1 | No | No | - |
| Igflr1 | 3,68E-02 | 2,75 | IGF-like family receptor 1 | No | No | - |
| Nudt6 | 1,37E-02 | 2,75 | nudix (nucleoside diphosphate linked moiety X)-type motif | No | No | - |

| | | | | | | |
|---|---|---|---|---|---|---|
| Hscb | 4,61E-04 | 2,76 | HscB iron-sulfur cluster co-chaperone | No | No | - |
| Eno1 | 1,49E-03 | 2,76 | enolase 1, alpha non-neuron | No | No | - |
| Cops9 | 1,58E-07 | 2,76 | COP9 signalosome subunit 9 | No | No | - |
| Dusp28 | 2,32E-03 | 2,76 | dual specificity phosphatase 28 | No | No | - |
| Zfp938 | 1,25E-03 | 2,76 | zinc finger protein 938 | No | No | - |
| Slc35a5 | 3,16E-02 | 2,76 | solute carrier family 35, member A5 | No | No | - |
| Ankrd37 | 2,51E-05 | 2,76 | ankyrin repeat domain 37 | No | No | - |
| Ccnyl1 | 1,99E-03 | 2,77 | cyclin Y-like 1 | No | No | - |
| Fam136a | 3,68E-03 | 2,77 | family with sequence similarity 136, member A | No | No | - |
| Tmem128 | 1,89E-03 | 2,77 | transmembrane protein 128 | No | No | - |
| Snrnp27 | 9,58E-06 | 2,77 | small nuclear ribonucleoprotein 27 (U4/U6.U5) | No | No | - |
| Mrpl20 | 6,84E-07 | 2,77 | mitochondrial ribosomal protein L20 | No | No | - |
| 5033417F24Rik | 4,03E-03 | 2,77 | RIKEN cDNA 5033417F24 gene | No | No | - |
| Tmco1 | 3,41E-04 | 2,77 | transmembrane and coiled-coil domains 1 | No | No | - |
| Hibadh | 1,38E-04 | 2,78 | 3-hydroxyisobutyrate dehydrogenase | No | No | - |
| Prepl | 7,87E-03 | 2,78 | prolyl endopeptidase-like | No | No | - |
| Uqcc2 | 8,93E-05 | 2,79 | ubiquinol-cytochrome c reductase complex assembly factor | No | No | - |
| Gtf2h2 | 1,16E-03 | 2,79 | general transcription factor II H, polypeptide 2 | No | No | - |
| Steap1 | 6,41E-06 | 2,80 | six transmembrane epithelial antigen of the prostate 1 | No | No | - |
| Gm6136 | 1,24E-05 | 2,80 | predicted pseudogene 6136 | No | No | - |
| Tex30 | 4,28E-03 | 2,80 | testis expressed 30 | No | No | - |
| Cklf | 7,37E-05 | 2,80 | chemokine-like factor | No | No | 1 |
| Hivep3 | 8,86E-03 | 2,81 | human immunodeficiency virus type I enhancer binding pro | No | No | - |
| Spryd4 | 1,02E-02 | 2,81 | SPRY domain containing 4 | No | No | - |
| Tpst1 | 3,36E-04 | 2,81 | protein-tyrosine sulfotransferase 1 | No | No | - |
| Ss18l2 | 2,05E-04 | 2,81 | SS18, nBAF chromatin remodeling complex subunit like 2 | No | No | - |
| Cryzl1 | 2,57E-03 | 2,81 | crystallin, zeta (quinone reductase)-like 1 | No | No | - |
| Mblac1 | 2,70E-04 | 2,81 | metallo-beta-lactamase domain containing 1 | No | No | - |
| Rnft1 | 1,71E-04 | 2,81 | ring finger protein, transmembrane 1 | No | No | - |
| Dnm3os | 2,43E-04 | 2,81 | dynamin 3, opposite strand | No | No | - |
| Hoxa4 | 2,75E-05 | 2,82 | homeobox A4 | No | No | - |
| Snai2 | 2,53E-02 | 2,82 | snail family zinc finger 2 | No | No | 4 |
| Ndufb1-ps | 1,72E-06 | 2,82 | NADH:ubiquinone oxidoreductase subunit B1 | No | No | - |
| Nfyb | 9,72E-05 | 2,82 | nuclear transcription factor-Y beta | No | No | - |
| Rgs10 | 4,98E-04 | 2,82 | regulator of G-protein signalling 10 | No | No | - |
| Tmem258 | 2,97E-07 | 2,82 | transmembrane protein 258 | No | No | - |
| Rrs1 | 9,42E-07 | 2,82 | ribosome biogenesis regulator 1 | No | No | - |
| Trim59 | 3,31E-04 | 2,83 | tripartite motif-containing 59 | No | No | - |
| Rpl15 | 1,02E-05 | 2,83 | ribosomal protein L15 | No | No | - |
| Psmd5 | 5,48E-05 | 2,83 | proteasome (prosome, macropain) 26S subunit, non-ATPase | No | No | - |
| Mrpl21 | 3,09E-04 | 2,83 | mitochondrial ribosomal protein L21 | No | No | - |
| Fam174a | 1,12E-04 | 2,83 | family with sequence similarity 174, member A | No | No | - |
| Smim27 | 4,20E-07 | 2,83 | small integral membrane protein 27 | No | No | - |
| Gemin2 | 1,39E-02 | 2,83 | gem nuclear organelle associated protein 2 | No | No | - |
| H2aw | 4,99E-06 | 2,83 | H2A.W histone | No | No | - |
| Spata24 | 1,24E-02 | 2,83 | spermatogenesis associated 24 | No | No | - |
| Id2 | 3,92E-04 | 2,83 | inhibitor of DNA binding 2 | No | No | 2 |
| Ptger2 | 3,00E-02 | 2,84 | prostaglandin E receptor 2 (subtype EP2) | No | No | 5 |
| Nfu1 | 2,97E-07 | 2,84 | NFU1 iron-sulfur cluster scaffold | No | No | - |
| Rps7 | 2,74E-08 | 2,85 | ribosomal protein S7 | No | No | - |
| Banf1 | 1,28E-07 | 2,85 | BAF nuclear assembly factor 1 | No | No | - |
| Nfs1 | 1,24E-05 | 2,85 | nitrogen fixation gene 1 (S. cerevisiae) | No | No | - |
| Aasdhppt | 5,70E-04 | 2,85 | aminoadipate-semialdehyde dehydrogenase-phosphopantetł | No | No | - |
| Epb41l4aos | 1,75E-04 | 2,85 | erythrocyte membrane protein band 4.1 like 4a, opposite str | No | No | - |
| Dbndd2 | 9,95E-04 | 2,85 | dysbindin (dystrobrevin binding protein 1) domain containi | No | No | - |
| Mrps33 | 8,17E-05 | 2,86 | mitochondrial ribosomal protein S33 | No | No | - |
| Snrpe | 9,54E-07 | 2,86 | small nuclear ribonucleoprotein E | No | No | - |
| Pigx | 7,87E-06 | 2,86 | phosphatidylinositol glycan anchor biosynthesis, class X | No | No | - |
| Gm6311 | 1,64E-04 | 2,86 | predicted gene 6311 | No | No | - |
| Ndufb6 | 4,16E-06 | 2,86 | NADH:ubiquinone oxidoreductase subunit B6 | No | No | - |
| Errfi1 | 3,31E-04 | 2,86 | ERBB receptor feedback inhibitor 1 | No | No | - |
| Lyrm1 | 6,05E-03 | 2,87 | LYR motif containing 1 | No | No | - |
| Alkbh2 | 1,36E-02 | 2,87 | alkB homolog 2, alpha-ketoglutarate-dependent dioxygenas | No | No | - |
| Cmc2 | 1,38E-04 | 2,87 | COX assembly mitochondrial protein 2 | No | No | - |
| Acadm | 4,36E-03 | 2,88 | acyl-Coenzyme A dehydrogenase, medium chain | No | No | - |
| Gm12989 | 1,52E-02 | 2,88 | predicted gene 12989 | No | No | - |
| Pex12 | 1,20E-02 | 2,88 | peroxisomal biogenesis factor 12 | No | No | - |
| Mrpl44 | 5,87E-03 | 2,88 | mitochondrial ribosomal protein L44 | No | No | - |
| Ccdc28b | 1,73E-03 | 2,88 | coiled coil domain containing 28B | No | No | - |
| Grpel1 | 5,22E-06 | 2,88 | GrpE-like 1, mitochondrial | No | No | - |
| Mettl21a | 2,94E-04 | 2,88 | methyltransferase like 21A | No | No | - |
| Trappc2 | 3,08E-03 | 2,88 | trafficking protein particle complex 2 | No | No | - |
| Gm16437 | 3,06E-05 | 2,89 | predicted pseudogene 16437 | No | No | - |

| | | | | | | |
|---|---|---|---|---|---|---|
| Ccdc59 | 1,21E-03 | 2,89 | coiled-coil domain containing 59 | No | No | - |
| Sra1 | 6,84E-07 | 2,89 | steroid receptor RNA activator 1 | No | No | - |
| Gtpbp3 | 9,26E-03 | 2,90 | GTP binding protein 3 | No | No | - |
| Cdca8 | 7,94E-04 | 2,90 | cell division cycle associated 8 | No | No | - |
| Tiprl | 2,54E-05 | 2,90 | TIP41, TOR signalling pathway regulator-like (S. cerevisiae | No | No | - |
| Ppia | 9,06E-07 | 2,90 | peptidylprolyl isomerase A | No | No | - |
| Acot2 | 4,53E-03 | 2,90 | acyl-CoA thioesterase 2 | No | No | - |
| Cep76 | 1,71E-02 | 2,90 | centrosomal protein 76 | No | No | - |
| Tmem38b | 1,37E-03 | 2,90 | transmembrane protein 38B | No | No | - |
| Tmem53 | 1,99E-02 | 2,90 | transmembrane protein 53 | No | No | - |
| Gm9493 | 3,35E-07 | 2,90 | predicted gene 9493 | No | No | - |
| Rpl18a | 2,93E-07 | 2,91 | ribosomal protein L18A | No | No | - |
| Dctpp1 | 1,51E-03 | 2,91 | dCTP pyrophosphatase 1 | No | No | - |
| Ptrhd1 | 2,03E-04 | 2,91 | peptidyl-tRNA hydrolase domain containing 1 | No | No | - |
| Chchd7 | 8,06E-05 | 2,91 | coiled-coil-helix-coiled-coil-helix domain containing 7 | No | No | - |
| Gm13461 | 6,49E-05 | 2,91 | predicted gene 13461 | No | No | - |
| Mrpl45 | 6,83E-04 | 2,92 | mitochondrial ribosomal protein L45 | No | No | - |
| Adarb1 | 5,89E-03 | 2,92 | adenosine deaminase, RNA-specific, B1 | No | No | - |
| Zmym3 | 2,78E-03 | 2,92 | zinc finger, MYM-type 3 | No | No | - |
| Tmem9b | 1,79E-07 | 2,92 | TMEM9 domain family, member B | No | No | - |
| Imp3 | 1,73E-05 | 2,93 | IMP3, U3 small nucleolar ribonucleoprotein | No | No | - |
| Lamtor3 | 9,26E-08 | 2,93 | late endosomal/lysosomal adaptor, MAPK and MTOR activ | No | No | - |
| Epdr1 | 1,19E-03 | 2,93 | ependymin related protein 1 (zebrafish) | No | No | - |
| Rpl9-ps4 | 4,40E-02 | 2,93 | ribosomal protein L9, pseudogene 4 | No | No | - |
| Ern2 | 3,90E-07 | 2,93 | endoplasmic reticulum (ER) to nucleus signalling 2 | No | No | 3 |
| Gm4691 | 4,13E-02 | 2,93 | predicted gene 4691 | No | No | - |
| Stard3nl | 5,00E-03 | 2,93 | STARD3 N-terminal like | No | No | - |
| Zcchc9 | 7,82E-03 | 2,94 | zinc finger, CCHC domain containing 9 | No | No | - |
| Txndc12 | 4,76E-05 | 2,94 | thioredoxin domain containing 12 (endoplasmic reticulum) | No | No | - |
| Cfap36 | 1,19E-04 | 2,94 | cilia and flagella associated protein 36 | No | No | - |
| BC003965 | 4,76E-05 | 2,94 | cDNA sequence BC003965 | No | No | - |
| Fez1 | 3,09E-04 | 2,95 | fasciculation and elongation protein zeta 1 (zygin I) | No | No | - |
| Trnau1ap | 5,32E-05 | 2,95 | tRNA selenocysteine 1 associated protein 1 | No | No | - |
| Asrgl1 | 1,94E-02 | 2,95 | asparaginase like 1 | No | No | - |
| Gm49384 | 1,27E-02 | 2,96 | predicted gene, 49384 | No | No | - |
| 1810037I17Rik | 1,04E-08 | 2,96 | RIKEN cDNA 1810037I17 gene | No | No | - |
| Psenen | 1,33E-02 | 2,96 | presenilin enhancer gamma secretase subunit | No | No | - |
| Smim3 | 3,19E-05 | 2,96 | small integral membrane protein 3 | No | No | - |
| Rnf130 | 2,35E-03 | 2,96 | ring finger protein 130 | No | No | - |
| Peg3 | 1,27E-02 | 2,96 | paternally expressed 3 | No | No | - |
| Prdx3 | 1,89E-06 | 2,97 | peroxiredoxin 3 | No | No | - |
| BC028528 | 1,14E-02 | 2,97 | cDNA sequence BC028528 | No | No | - |
| Taf11 | 1,17E-04 | 2,97 | TATA-box binding protein associated factor 11 | No | No | - |
| Rpl30 | 6,67E-09 | 2,98 | ribosomal protein L30 | No | No | - |
| Stbd1 | 1,62E-06 | 2,98 | starch binding domain 1 | No | No | - |
| Fabp4 | 1,59E-07 | 2,98 | fatty acid binding protein 4, adipocyte | No | No | 7 |
| Srp19 | 2,80E-07 | 2,98 | signal recognition particle 19 | No | No | - |
| Acer3 | 1,86E-04 | 2,98 | alkaline ceramidase 3 | No | No | - |
| Unc50 | 1,49E-05 | 2,99 | unc-50 homolog | No | No | - |
| Acyp2 | 1,98E-02 | 3,00 | acylphosphatase 2, muscle type | No | No | - |
| Adcy1 | 1,44E-02 | 3,00 | adenylate cyclase 1 | No | No | - |
| Avpr1a | 2,41E-04 | 3,00 | arginine vasopressin receptor 1A | No | No | - |
| Cenpf | 1,15E-04 | 3,00 | centromere protein F | No | No | - |
| Dync2li1 | 3,73E-02 | 3,00 | dynein cytoplasmic 2 light intermediate chain 1 | No | No | - |
| Eid2b | 1,36E-02 | 3,00 | EP300 interacting inhibitor of differentiation 2B | No | No | - |
| Gm6139 | 2,77E-02 | 3,00 | predicted gene 6139 | No | No | - |
| Ppp1cc | 1,44E-02 | 3,00 | protein phosphatase 1 catalytic subunit gamma | No | No | - |
| Tk1 | 6,91E-03 | 3,00 | thymidine kinase 1 | No | No | - |
| Tubd1 | 4,42E-02 | 3,00 | tubulin, delta 1 | No | No | - |
| Ppic | 1,42E-07 | 3,00 | peptidylprolyl isomerase C | No | No | - |
| H2bc4 | 6,30E-07 | 3,00 | H2B clustered histone 4 | No | No | - |
| Txnip | 2,62E-05 | 3,01 | thioredoxin interacting protein | Yes | No | 9 |
| 5430416N02Rik | 6,90E-04 | 3,02 | RIKEN cDNA 5430416N02 gene | No | No | - |
| Srsf7 | 2,12E-05 | 3,02 | serine and arginine-rich splicing factor 7 | No | No | - |
| Mcee | 1,20E-04 | 3,02 | methylmalonyl CoA epimerase | No | No | - |
| Commd6 | 9,06E-04 | 3,02 | COMM domain containing 6 | No | No | - |
| Immp1l | 4,50E-05 | 3,03 | IMP1 inner mitochondrial membrane peptidase-like (S. cere | No | No | - |
| Gm42420 | 1,53E-07 | 3,03 | predicted gene, 42420 | No | No | - |
| Mpc1 | 5,90E-03 | 3,03 | mitochondrial pyruvate carrier 1 | No | No | - |
| Ube2b | 1,20E-06 | 3,04 | ubiquitin-conjugating enzyme E2B | No | No | - |
| Erlin1 | 5,25E-03 | 3,04 | ER lipid raft associated 1 | No | No | - |
| Sec61b | 7,71E-08 | 3,04 | Sec61 beta subunit | No | No | - |
| Tmbim4 | 1,38E-06 | 3,04 | transmembrane BAX inhibitor motif containing 4 | No | No | - |

| | | | | | | |
|---|---|---|---|---|---|---|
| Gm18924 | 5,36E-04 | 3,04 | predicted gene, 18924 | No | No | - |
| Arrdc4 | 2,80E-04 | 3,05 | arrestin domain containing 4 | No | No | - |
| Spcs2 | 1,13E-08 | 3,05 | signal peptidase complex subunit 2 homolog (S. cerevisiae) | No | No | - |
| Mlx | 1,80E-03 | 3,05 | MAX-like protein X | No | No | - |
| Atp5g2 | 1,19E-03 | 3,06 | ATP synthase, H+ transporting, mitochondrial F0 complex, | No | No | - |
| Ntan1 | 2,57E-02 | 3,06 | N-terminal Asn amidase | No | No | - |
| Ing2 | 3,79E-04 | 3,06 | inhibitor of growth family, member 2 | No | No | - |
| Rpl13-ps3 | 6,02E-04 | 3,06 | ribosomal protein L13, pseudogene 3 | No | No | - |
| Plk1 | 4,14E-07 | 3,06 | polo like kinase 1 | No | No | 2 |
| Mrpl23 | 3,04E-03 | 3,06 | mitochondrial ribosomal protein L23 | No | No | - |
| Grb14 | 1,18E-02 | 3,07 | growth factor receptor bound protein 14 | No | No | - |
| Zbed3 | 3,86E-02 | 3,07 | zinc finger, BED type containing 3 | No | No | - |
| Akr1e1 | 3,41E-04 | 3,07 | aldo-keto reductase family 1, member E1 | No | No | - |
| Emx2 | 4,35E-04 | 3,07 | empty spiracles homeobox 2 | No | No | - |
| Prrx2 | 1,77E-04 | 3,08 | paired related homeobox 2 | No | No | - |
| Notch1 | 1,75E-02 | 3,08 | notch 1 | No | No | 18 |
| Cox17 | 1,13E-09 | 3,08 | cytochrome c oxidase assembly protein 17, copper chaperon | No | No | - |
| Ndufs5 | 3,79E-05 | 3,08 | NADH:ubiquinone oxidoreductase core subunit S5 | No | No | - |
| Ctbs | 6,27E-03 | 3,08 | chitobiase | No | No | - |
| A130010J15Rik | 2,31E-02 | 3,09 | RIKEN cDNA A130010J15 gene | No | No | - |
| Ech1 | 4,69E-05 | 3,09 | enoyl coenzyme A hydratase 1, peroxisomal | No | No | - |
| Fahd1 | 1,18E-03 | 3,09 | fumarylacetoacetate hydrolase domain containing 1 | No | No | - |
| Hoxaas2 | 8,49E-04 | 3,09 | Hoxa cluster antisense RNA 2 | No | No | - |
| 9130008F23Rik | 1,02E-02 | 3,10 | RIKEN cDNA 9130008F23 gene | No | No | - |
| Dpf1 | 1,06E-02 | 3,10 | D4, zinc and double PHD fingers family 1 | No | No | - |
| Rbm4b | 8,50E-03 | 3,10 | RNA binding motif protein 4B | No | No | - |
| Zcchc10 | 4,13E-04 | 3,10 | zinc finger, CCHC domain containing 10 | No | No | - |
| Atp5c1 | 4,31E-07 | 3,11 | ATP synthase, H+ transporting, mitochondrial F1 complex, | No | No | - |
| Srp14 | 2,96E-07 | 3,11 | signal recognition particle 14 | No | No | - |
| Btg3 | 1,84E-02 | 3,11 | BTG anti-proliferation factor 3 | No | No | - |
| St6galnac2 | 1,17E-02 | 3,11 | ST6 (alpha-N-acetyl-neuraminyl-2,3-beta-galactosyl-1,3)-N | No | No | - |
| Gm20458 | 4,19E-04 | 3,11 | predicted gene 20458 | No | No | - |
| Selenok | 6,83E-08 | 3,11 | selenoprotein K | No | No | - |
| Lamtor5 | 1,09E-09 | 3,12 | late endosomal/lysosomal adaptor, MAPK and MTOR activ | No | No | - |
| Irx3 | 1,69E-02 | 3,13 | Iroquois related homeobox 3 | No | No | - |
| Snupn | 2,16E-04 | 3,13 | snurportin 1 | No | No | - |
| Exosc2 | 5,39E-04 | 3,13 | exosome component 2 | No | No | - |
| Cyp26b1 | 6,71E-04 | 3,13 | cytochrome P450, family 26, subfamily b, polypeptide 1 | No | No | - |
| Gm5537 | 5,02E-09 | 3,14 | predicted gene 5537 | No | No | - |
| Gamt | 1,62E-02 | 3,14 | guanidinoacetate methyltransferase | No | No | - |
| Hhex | 7,68E-04 | 3,14 | hematopoietically expressed homeobox | No | No | - |
| Iqcc | 5,52E-03 | 3,14 | IQ motif containing C | No | No | - |
| Rnf113a1 | 1,44E-02 | 3,15 | ring finger protein 113A1 | No | No | - |
| Ybx1 | 7,01E-08 | 3,15 | Y box protein 1 | No | No | 2 |
| Rps7-ps3 | 4,11E-07 | 3,15 | ribosomal protein S7, pseudogene 3 | No | No | - |
| 2610001J05Rik | 5,65E-06 | 3,16 | small integral membrane protein 30 | No | No | - |
| E130311K13Rik | 1,41E-02 | 3,16 | RIKEN cDNA E130311K13 gene | No | No | - |
| Gtf2a2 | 1,74E-05 | 3,16 | general transcription factor II A, 2 | No | No | - |
| Npm3-ps1 | 2,30E-02 | 3,16 | nucleoplasmin 3, pseudogene 1 | No | No | - |
| Arl6 | 4,40E-03 | 3,16 | ADP-ribosylation factor-like 6 | No | No | - |
| Cdc42ep3 | 1,06E-04 | 3,16 | CDC42 effector protein (Rho GTPase binding) 3 | No | No | - |
| Dynll1 | 6,30E-09 | 3,16 | dynein light chain LC8-type 1 | No | No | - |
| Nipsnap3b | 3,21E-05 | 3,16 | nipsnap homolog 3B | No | No | - |
| Ube2w | 4,89E-05 | 3,17 | ubiquitin-conjugating enzyme E2W (putative) | No | No | - |
| Mrpl53 | 7,43E-09 | 3,17 | mitochondrial ribosomal protein L53 | No | No | - |
| Tsen15 | 1,76E-03 | 3,17 | tRNA splicing endonuclease subunit 15 | No | No | - |
| Gm10184 | 1,81E-03 | 3,18 | predicted pseudogene 10184 | No | No | - |
| Glipr2 | 8,43E-09 | 3,18 | GLI pathogenesis-related 2 | No | No | - |
| Pdhb | 6,40E-07 | 3,18 | pyruvate dehydrogenase (lipoamide) beta | No | No | - |
| 3110070M22Rik | 4,73E-02 | 3,18 | RIKEN cDNA 3110070M22 gene | No | No | - |
| Gm11110 | 1,26E-02 | 3,18 | predicted gene 11110 | No | No | - |
| Hacd2 | 5,08E-03 | 3,19 | 3-hydroxyacyl-CoA dehydratase 2 | No | No | - |
| Sec11c | 8,95E-07 | 3,19 | SEC11 homolog C, signal peptidase complex subunit | No | No | - |
| Fam117a | 2,11E-03 | 3,20 | family with sequence similarity 117, member A | No | No | - |
| Septin9 | 9,21E-03 | 3,20 | septin 9 | No | No | - |
| Sem1 | 9,24E-07 | 3,20 | SEM1, 26S proteasome complex subunit | No | No | - |
| Atp5l | 1,53E-09 | 3,20 | ATP synthase, H+ transporting, mitochondrial F0 complex, | No | No | - |
| Fbxl12 | 1,42E-04 | 3,20 | F-box and leucine-rich repeat protein 12 | No | No | - |
| Spcs1 | 3,24E-09 | 3,21 | signal peptidase complex subunit 1 homolog (S. cerevisiae) | No | No | - |
| Emc7 | 8,39E-08 | 3,21 | ER membrane protein complex subunit 7 | No | No | - |
| Ugdh | 2,76E-05 | 3,21 | UDP-glucose dehydrogenase | No | No | - |
| Prpsap2 | 2,51E-02 | 3,21 | phosphoribosyl pyrophosphate synthetase-associated protei | No | No | - |
| Sike1 | 7,16E-08 | 3,22 | suppressor of IKBKE 1 | No | No | - |

| | | | | | | |
|---|---|---|---|---|---|---|
| Steap3 | 1,76E-05 | 3,22 | STEAP family member 3 | No | No | - |
| Ift43 | 8,48E-05 | 3,22 | intraflagellar transport 43 | No | No | - |
| Gm42688 | 8,03E-09 | 3,23 | predicted gene 42688 | No | No | - |
| Fbxo5 | 4,59E-03 | 3,23 | F-box protein 5 | No | No | - |
| Gm12918 | 2,70E-05 | 3,23 | predicted gene 12918 | No | No | - |
| Vmp1 | 3,24E-06 | 3,23 | vacuole membrane protein 1 | No | No | - |
| Siva1 | 9,40E-08 | 3,24 | SIVA1, apoptosis-inducing factor | No | No | - |
| Ccne1 | 3,57E-05 | 3,24 | cyclin E1 | No | No | - |
| Ccnd1 | 3,78E-05 | 3,24 | cyclin D1 | Yes | No | 2 |
| Mat2b | 7,89E-05 | 3,24 | methionine adenosyltransferase II, beta | No | No | - |
| Abt1 | 2,41E-03 | 3,25 | activator of basal transcription 1 | No | No | - |
| Dusp19 | 3,75E-04 | 3,25 | dual specificity phosphatase 19 | No | No | - |
| Gm6286 | 8,95E-03 | 3,26 | predicted gene 6286 | No | No | - |
| Hsdl2 | 2,38E-03 | 3,27 | hydroxysteroid dehydrogenase like 2 | No | No | - |
| Gm4204 | 1,63E-04 | 3,27 | predicted gene 4204 | No | No | - |
| Cdk5r1 | 3,43E-02 | 3,27 | cyclin-dependent kinase 5, regulatory subunit 1 (p35) | No | No | - |
| Rpl10 | 3,38E-09 | 3,27 | ribosomal protein L10 | No | No | - |
| Casp6 | 3,98E-05 | 3,28 | caspase 6 | No | No | 1 |
| Pi15 | 3,68E-03 | 3,28 | peptidase inhibitor 15 | No | No | - |
| Hspe1-rs1 | 1,63E-05 | 3,28 | heat shock protein 1 (chaperonin 10), related sequence 1 | No | No | - |
| Ap3s1 | 2,00E-03 | 3,29 | adaptor-related protein complex 3, sigma 1 subunit | No | No | - |
| Nat14 | 1,28E-02 | 3,29 | N-acetyltransferase 14 | No | No | - |
| Dusp12 | 2,66E-05 | 3,29 | dual specificity phosphatase 12 | No | No | - |
| Pex2 | 5,19E-03 | 3,29 | peroxisomal biogenesis factor 2 | No | No | - |
| Nudt1 | 5,51E-05 | 3,30 | nudix (nucleoside diphosphate linked moiety X)-type motif | No | No | - |
| Rps27l | 5,48E-10 | 3,30 | ribosomal protein S27-like | No | No | - |
| Dancr | 9,49E-03 | 3,30 | differentiation antagonizing non-protein coding RNA | No | No | - |
| Eci1 | 1,24E-03 | 3,30 | enoyl-Coenzyme A delta isomerase 1 | No | No | - |
| Fn3krp | 3,78E-02 | 3,30 | fructosamine 3 kinase related protein | No | No | - |
| Slc30a6 | 9,12E-04 | 3,30 | solute carrier family 30 (zinc transporter), member 6 | No | No | - |
| Trim13 | 5,23E-03 | 3,30 | tripartite motif-containing 13 | No | No | - |
| Car5b | 2,03E-02 | 3,31 | carbonic anhydrase 5b, mitochondrial | No | No | - |
| Mzt2 | 3,16E-03 | 3,31 | mitotic spindle organizing protein 2 | No | No | - |
| St3gal5 | 3,85E-07 | 3,31 | ST3 beta-galactoside alpha-2,3-sialyltransferase 5 | No | No | - |
| Eid1 | 6,30E-09 | 3,31 | EP300 interacting inhibitor of differentiation 1 | No | No | - |
| Apba3 | 4,24E-03 | 3,31 | amyloid beta (A4) precursor protein-binding, family A, mer | No | No | - |
| Ccng2 | 5,76E-06 | 3,31 | cyclin G2 | No | No | - |
| Gm20503 | 3,88E-09 | 3,31 | predicted gene 20503 | No | No | - |
| Dusp14 | 5,77E-03 | 3,32 | dual specificity phosphatase 14 | No | No | - |
| Nr1h2 | 2,40E-05 | 3,32 | nuclear receptor subfamily 1, group H, member 2 | No | No | - |
| 1190007I07Rik | 1,04E-03 | 3,32 | RIKEN cDNA 1190007I07 gene | No | No | - |
| Lrrc57 | 1,22E-03 | 3,32 | leucine rich repeat containing 57 | No | No | - |
| Mcts2 | 7,16E-05 | 3,32 | malignant T cell amplified sequence 2 | No | No | - |
| Sft2d3 | 9,43E-04 | 3,32 | SFT2 domain containing 3 | No | No | - |
| Naa10 | 6,01E-05 | 3,33 | N(alpha)-acetyltransferase 10, NatA catalytic subunit | No | No | - |
| Prxl2b | 4,90E-05 | 3,33 | peroxiredoxin like 2B | No | No | - |
| Bloc1s4 | 4,91E-08 | 3,33 | biogenesis of lysosomal organelles complex-1, subunit 4, ca | No | No | - |
| 2210408I21Rik | 1,40E-02 | 3,33 | RIKEN cDNA 2210408I21 gene | No | No | - |
| Dnase1l1 | 1,92E-09 | 3,33 | deoxyribonuclease 1-like 1 | No | No | - |
| Gm3379 | 1,92E-05 | 3,33 | predicted gene 3379 | No | No | - |
| Lyrm4 | 3,46E-04 | 3,33 | LYR motif containing 4 | No | No | - |
| Mycbp | 9,16E-07 | 3,33 | MYC binding protein | No | No | - |
| Pex10 | 2,30E-02 | 3,33 | peroxisomal biogenesis factor 10 | No | No | - |
| Acat1 | 6,65E-08 | 3,34 | acetyl-Coenzyme A acetyltransferase 1 | No | No | - |
| Smim8 | 2,46E-03 | 3,34 | small integral membrane protein 8 | No | No | - |
| Gm5552 | 1,80E-03 | 3,35 | predicted gene 5552 | No | No | - |
| Ccdc62 | 1,65E-02 | 3,36 | coiled-coil domain containing 62 | No | No | - |
| Tusc3 | 4,25E-09 | 3,36 | - | No | No | - |
| Styx | 5,75E-08 | 3,37 | serine/threonine/tyrosine interaction protein | No | No | - |
| Exo5 | 1,58E-03 | 3,37 | exonuclease 5 | No | No | - |
| Cfap298 | 1,09E-02 | 3,37 | cilia and flagella associate protien 298 | No | No | - |
| Lzic | 1,52E-04 | 3,37 | leucine zipper and CTNNBIP1 domain containing | No | No | - |
| Erich1 | 8,48E-03 | 3,38 | glutamate rich 1 | No | No | - |
| Harbi1 | 2,69E-02 | 3,38 | harbinger transposase derived 1 | No | No | - |
| Scp2 | 4,39E-09 | 3,39 | sterol carrier protein 2, liver | No | No | - |
| Polr3k | 1,13E-06 | 3,40 | polymerase (RNA) III (DNA directed) polypeptide K | No | No | - |
| Vta1 | 5,07E-06 | 3,40 | vesicle (multivesicular body) trafficking 1 | No | No | - |
| Stc1 | 8,42E-05 | 3,40 | stanniocalcin 1 | No | No | 24 |
| Bhlhe22 | 2,46E-04 | 3,41 | basic helix-loop-helix family, member e22 | No | No | - |
| Tctex1d2 | 8,31E-07 | 3,42 | dynein light chain Tctex-type 2B | No | No | - |
| Nqo1 | 1,05E-03 | 3,42 | NAD(P)H dehydrogenase, quinone 1 | No | No | 16 |
| Commd2 | 1,30E-06 | 3,42 | COMM domain containing 2 | Yes | No | - |
| Psmg1 | 2,51E-03 | 3,42 | proteasome (prosome, macropain) assembly chaperone 1 | No | No | - |

| | | | | | | |
|---|---|---|---|---|---|---|
| Spsb2 | 3,16E-11 | 3,43 | splA/ryanodine receptor domain and SOCS box containing 2 | No | No | - |
| Msrb2 | 1,48E-02 | 3,43 | methionine sulfoxide reductase B2 | No | No | - |
| Magoh | 9,26E-08 | 3,44 | mago homolog, exon junction complex core component | No | No | - |
| Gng10 | 1,19E-09 | 3,44 | guanine nucleotide binding protein (G protein), gamma 10 | No | No | - |
| Tmem70 | 1,91E-07 | 3,45 | transmembrane protein 70 | No | No | - |
| Tmem165 | 3,35E-08 | 3,45 | transmembrane protein 165 | No | No | - |
| Uqcrfs1 | 2,82E-07 | 3,46 | ubiquinol-cytochrome c reductase, Rieske iron-sulfur polyp | No | No | - |
| Gemin6 | 1,17E-04 | 3,46 | gem nuclear organelle associated protein 6 | No | No | - |
| Chrna1os | 9,28E-04 | 3,47 | cholinergic receptor, nicotinic, alpha polypeptide 1 (muscle) | No | No | - |
| Ccnb2 | 1,42E-04 | 3,47 | cyclin B2 | Yes | No | - |
| Lsm1 | 3,73E-04 | 3,47 | LSM1 homolog, mRNA degradation associated | No | No | - |
| Med6 | 1,11E-04 | 3,47 | mediator complex subunit 6 | No | No | - |
| Sec13 | 1,59E-09 | 3,50 | SEC13 homolog, nuclear pore and COPII coat complex con | No | No | - |
| Pclaf | 2,42E-06 | 3,50 | PCNA clamp associated factor | No | No | - |
| Elfn1 | 1,98E-03 | 3,50 | leucine rich repeat and fibronectin type III, extracellular 1 | No | No | - |
| Il2 | 5,10E-04 | 3,50 | interleukin 2 | No | No | 11 |
| Lsm7 | 8,30E-03 | 3,50 | LSM7 homolog, U6 small nuclear RNA and mRNA degrad | No | No | - |
| Mrps28 | 1,21E-04 | 3,50 | mitochondrial ribosomal protein S28 | No | No | - |
| Snhg16 | 2,76E-02 | 3,50 | small nucleolar RNA host gene 16 | No | No | - |
| Timm23 | 1,09E-05 | 3,50 | translocase of inner mitochondrial membrane 23 | No | No | - |
| Tjp3 | 2,51E-03 | 3,50 | tight junction protein 3 | No | No | - |
| Tomm5 | 6,79E-05 | 3,50 | translocase of outer mitochondrial membrane 5 | No | No | - |
| Tmsb4x | 2,58E-13 | 3,50 | thymosin, beta 4, X chromosome | No | No | - |
| Srp9 | 7,49E-07 | 3,50 | signal recognition particle 9 | No | No | - |
| Ndufc2 | 7,46E-10 | 3,50 | NADH:ubiquinone oxidoreductase subunit C2 | No | No | - |
| Cfap20 | 3,84E-05 | 3,51 | cilia and flagella associated protein 20 | No | No | - |
| Fam83d | 6,45E-05 | 3,52 | family with sequence similarity 83, member D | No | No | - |
| Mthfsl | 6,06E-03 | 3,53 | 5, 10-methenyltetrahydrofolate synthetase-like | No | No | - |
| Tes | 4,06E-10 | 3,54 | testin LIM domain protein | No | No | - |
| 1700123O20Rik | 1,36E-07 | 3,54 | RIKEN cDNA 1700123O20 gene | No | No | - |
| Tpi1 | 5,96E-11 | 3,55 | triosephosphate isomerase 1 | No | No | - |
| Cox7c | 2,47E-10 | 3,55 | cytochrome c oxidase subunit 7C | No | No | - |
| Washc3 | 6,01E-05 | 3,56 | WASH complex subunit 3 | No | No | - |
| Nabp1 | 1,22E-05 | 3,56 | nucleic acid binding protein 1 | No | No | - |
| Gm13436 | 1,72E-05 | 3,56 | predicted gene 13436 | No | No | - |
| 1700120C14Rik | 2,00E-08 | 3,57 | RIKEN cDNA 1700120C14 gene | No | No | - |
| Churc1 | 4,33E-04 | 3,57 | churchill domain containing 1 | No | No | - |
| Rnf7 | 1,64E-08 | 3,57 | ring finger protein 7 | No | No | - |
| Ndufa6 | 1,19E-09 | 3,57 | NADH:ubiquinone oxidoreductase subunit A6 | No | No | - |
| Hmgn2 | 9,81E-03 | 3,57 | high mobility group nucleosomal binding domain 2 | No | No | - |
| Calhm5 | 4,39E-04 | 3,58 | calcium homeostasis modulator family member 5 | No | No | - |
| Snrpg | 2,02E-09 | 3,59 | small nuclear ribonucleoprotein polypeptide G | No | No | - |
| Dph3 | 6,23E-08 | 3,59 | diphthamine biosynthesis 3 | No | No | - |
| 1500011B03Rik | 2,86E-06 | 3,60 | RIKEN cDNA 1500011B03 gene | No | No | - |
| Hikeshi | 1,34E-06 | 3,60 | heat shock protein nuclear import factor | No | No | - |
| Letm2 | 2,29E-02 | 3,60 | leucine zipper-EF-hand containing transmembrane protein 2 | No | No | - |
| Tcfl5 | 2,12E-03 | 3,60 | transcription factor-like 5 (basic helix-loop-helix) | No | No | - |
| Gins2 | 1,04E-06 | 3,60 | GINS complex subunit 2 (Psf2 homolog) | No | No | - |
| Rab28 | 1,59E-07 | 3,63 | RAB28, member RAS oncogene family | No | No | - |
| Rfesd | 5,33E-03 | 3,63 | Rieske (Fe-S) domain containing | No | No | - |
| Dipk1b | 2,65E-03 | 3,64 | divergent protein kinase domain 1B | No | No | - |
| Prxl2c | 6,03E-04 | 3,64 | peroxiredoxin like 2C | No | No | - |
| Gse1 | 6,38E-07 | 3,65 | genetic suppressor element 1, coiled-coil protein | No | No | - |
| 1110004F10Rik | 4,70E-06 | 3,65 | RIKEN cDNA 1110004F10 gene | No | No | - |
| Psma5 | 1,11E-07 | 3,65 | proteasome subunit alpha 5 | No | No | - |
| 1110059G10Rik | 6,20E-05 | 3,65 | RIKEN cDNA 1110059G10 gene | No | No | - |
| 2410004B18Rik | 1,78E-05 | 3,67 | RIKEN cDNA 2410004B18 gene | No | No | - |
| Ccnq | 1,88E-03 | 3,67 | cyclin Q | No | No | - |
| Gm9803 | 6,17E-05 | 3,67 | presequence translocase associated motor 16 like | No | No | - |
| Pnp2 | 3,16E-03 | 3,67 | purine-nucleoside phosphorylase 2 | No | No | 7 |
| Egln1 | 2,31E-07 | 3,67 | egl-9 family hypoxia-inducible factor 1 | No | No | - |
| Agbl5 | 4,24E-07 | 3,67 | ATP/GTP binding protein-like 5 | No | No | - |
| Fbxo4 | 4,68E-05 | 3,67 | F-box protein 4 | No | No | - |
| Chchd3 | 3,60E-07 | 3,68 | coiled-coil-helix-coiled-coil-helix domain containing 3 | No | No | - |
| Dynlt1b | 8,93E-04 | 3,69 | dynein light chain Tctex-type 1B | No | No | - |
| Gng5 | 2,90E-06 | 3,69 | guanine nucleotide binding protein (G protein), gamma 5 | No | No | - |
| Gm13835 | 4,66E-08 | 3,69 | predicted gene 13835 | No | No | - |
| Naa20 | 2,57E-04 | 3,70 | N(alpha)-acetyltransferase 20, NatB catalytic subunit | No | No | - |
| Amn1 | 3,08E-03 | 3,70 | antagonist of mitotic exit network 1 | No | No | - |
| Hoxb8 | 2,51E-03 | 3,70 | homeobox B8 | No | No | - |
| Srpx | 2,32E-03 | 3,70 | sushi-repeat-containing protein | No | No | - |
| Gm21987 | 1,07E-07 | 3,70 | predicted gene 21987 | No | No | - |
| Cyp7b1 | 1,12E-03 | 3,71 | cytochrome P450, family 7, subfamily b, polypeptide 1 | No | No | - |

| | | | | | | |
|---|---|---|---|---|---|---|
| Gng2 | 4,92E-05 | 3,71 | guanine nucleotide binding protein (G protein), gamma 2 | No | No | - |
| Mbip | 1,11E-02 | 3,71 | MAP3K12 binding inhibitory protein 1 | No | No | - |
| Dnajc19 | 1,49E-05 | 3,72 | DnaJ heat shock protein family (Hsp40) member C19 | No | No | - |
| Cox16 | 7,83E-09 | 3,72 | cytochrome c oxidase assembly protein 16 | No | No | - |
| Gm7432 | 4,89E-04 | 3,73 | predicted gene 7432 | No | No | - |
| Fbxl14 | 1,65E-04 | 3,73 | F-box and leucine-rich repeat protein 14 | No | No | - |
| Rwdd1 | 4,40E-03 | 3,74 | RWD domain containing 1 | No | No | - |
| Gm13339 | 4,98E-03 | 3,75 | predicted gene 13339 | No | No | - |
| Tor1a | 9,93E-06 | 3,76 | torsin family 1, member A (torsin A) | No | No | - |
| Gnpnat1 | 4,48E-08 | 3,76 | glucosamine-phosphate N-acetyltransferase 1 | No | No | - |
| Cby1 | 2,31E-02 | 3,78 | chibby family member 1, beta catenin antagonist | No | No | - |
| Gm12366 | 1,67E-04 | 3,78 | predicted gene 12366 | No | No | - |
| Arl6ip6 | 5,78E-06 | 3,78 | ADP-ribosylation factor-like 6 interacting protein 6 | No | No | - |
| Gm49188 | 5,32E-03 | 3,79 | predicted gene, 49188 | No | No | - |
| Snx7 | 1,92E-05 | 3,79 | sorting nexin 7 | No | No | - |
| Inafm2 | 2,08E-04 | 3,79 | InaF motif containing 2 | No | No | - |
| Ngrn | 3,69E-08 | 3,80 | neugrin, neurite outgrowth associated | No | No | - |
| Rps2-ps8 | 2,25E-03 | 3,80 | ribosomal protein S2, pseudogene 8 | No | No | - |
| Ybx1-ps2 | 9,92E-04 | 3,80 | Y box protein 1, pseudogene 2 | No | No | - |
| Socs3 | 3,30E-04 | 3,80 | suppressor of cytokine signaling 3 | No | No | 22 |
| Cnep1r1 | 5,45E-06 | 3,82 | CTD nuclear envelope phosphatase 1 regulatory subunit 1 | No | No | - |
| Trdmt1 | 1,27E-02 | 3,83 | tRNA aspartic acid methyltransferase 1 | No | No | - |
| Tmem69 | 1,08E-02 | 3,85 | transmembrane protein 69 | No | No | - |
| Cldnd1 | 8,48E-05 | 3,85 | claudin domain containing 1 | No | No | - |
| Iscu | 3,66E-05 | 3,85 | iron-sulfur cluster assembly enzyme | No | No | - |
| 4930579G24Rik | 8,64E-03 | 3,86 | RIKEN cDNA 4930579G24 gene | No | No | - |
| Nr4a1 | 7,39E-03 | 3,86 | nuclear receptor subfamily 4, group A, member 1 | No | No | 8 |
| Polr2d | 5,26E-03 | 3,86 | polymerase (RNA) II (DNA directed) polypeptide D | No | No | - |
| Fam204a | 2,51E-05 | 3,87 | family with sequence similarity 204, member A | No | No | - |
| Pfn1 | 4,04E-04 | 3,87 | profilin 1 | No | No | - |
| Nxt1 | 1,86E-05 | 3,88 | NTF2-related export protein 1 | No | No | - |
| 4930506C21Rik | 5,89E-03 | 3,88 | RIKEN cDNA 4930506C21 gene | No | No | - |
| Dcps | 1,57E-06 | 3,89 | decapping enzyme, scavenger | No | No | - |
| Bola3 | 4,72E-07 | 3,89 | bolA-like 3 (E. coli) | No | No | - |
| Ccdc175 | 2,13E-06 | 3,89 | coiled-coil domain containing 175 | No | No | - |
| Tmem179b | 5,82E-09 | 3,90 | transmembrane protein 179B | No | No | - |
| Haus4 | 4,71E-03 | 3,90 | HAUS augmin-like complex, subunit 4 | No | No | - |
| Slc25a5 | 6,67E-08 | 3,91 | solute carrier family 25 (mitochondrial carrier, adenine nucl | No | No | - |
| Mis18a | 1,30E-05 | 3,92 | MIS18 kinetochore protein A | No | No | - |
| Dact1 | 8,84E-05 | 3,92 | dishevelled-binding antagonist of beta-catenin 1 | No | No | - |
| Rpl39 | 5,44E-12 | 3,92 | ribosomal protein L39 | No | No | - |
| Srf | 8,84E-05 | 3,93 | serum response factor | No | Yes | 44 |
| Umad1 | 4,97E-05 | 3,93 | UMAP1-MVP12 associated (UMA) domain containing 1 | No | No | - |
| Magohb | 2,46E-05 | 3,93 | mago homolog B, exon junction complex core component | No | No | - |
| Ccdc115 | 5,22E-06 | 3,93 | coiled-coil domain containing 115 | No | No | - |
| Ost4 | 6,38E-07 | 3,94 | oligosaccharyltransferase complex subunit 4 (non-catalytic) | No | No | - |
| Tarbp1 | 2,34E-04 | 3,95 | TAR RNA binding protein 1 | No | No | - |
| Cmpk1 | 1,09E-08 | 3,96 | cytidine monophosphate (UMP-CMP) kinase 1 | No | No | - |
| Birc5 | 3,11E-07 | 3,97 | baculoviral IAP repeat-containing 5 | Yes | No | - |
| Mydgf | 3,17E-11 | 3,97 | myeloid derived growth factor | No | No | - |
| Ccnb1 | 1,32E-04 | 3,97 | cyclin B1 | Yes | No | - |
| Zfp410 | 1,51E-04 | 3,98 | zinc finger protein 410 | No | No | - |
| Bet1 | 1,59E-06 | 3,98 | Bet1 golgi vesicular membrane trafficking protein | No | No | - |
| Id1 | 1,59E-07 | 3,98 | inhibitor of DNA binding 1, HLH protein | No | No | 6 |
| Mtln | 4,65E-09 | 3,99 | mitoregulin | No | No | - |
| Cck | 8,13E-09 | 4,00 | cholecystokinin | No | No | - |
| 2310039H08Rik | 1,06E-05 | 4,00 | RIKEN cDNA 2310039H08 gene | No | No | - |
| Ccdc34 | 5,62E-04 | 4,00 | coiled-coil domain containing 34 | No | No | - |
| Gm17300 | 8,50E-03 | 4,00 | predicted gene, 17300 | No | No | - |
| Ndufa4 | 5,44E-09 | 4,02 | Ndufa4, mitochondrial complex associated | No | No | - |
| Thy1 | 1,95E-09 | 4,02 | thymus cell antigen 1, theta | No | No | 15 |
| Nip7 | 6,95E-10 | 4,03 | NIP7, nucleolar pre-rRNA processing protein | No | No | - |
| Ppih | 1,01E-05 | 4,04 | peptidyl prolyl isomerase H | No | No | - |
| Nit2 | 2,35E-03 | 4,04 | nitrilase family, member 2 | No | No | - |
| Snx24 | 1,95E-03 | 4,05 | sorting nexing 24 | No | No | - |
| Gm32282 | 1,82E-06 | 4,05 | predicted gene, 32282 | No | No | - |
| Hoxa5 | 1,43E-09 | 4,05 | homeobox A5 | No | No | 1 |
| Psma3 | 4,91E-13 | 4,05 | proteasome subunit alpha 3 | No | No | - |
| Gm47626 | 6,48E-05 | 4,07 | predicted gene, 47626 | No | No | - |
| Slc35g2 | 2,02E-03 | 4,07 | solute carrier family 35, member G2 | No | No | - |
| Gm29430 | 1,73E-05 | 4,08 | predicted gene 29430 | No | No | - |
| Ccdc51 | 9,14E-03 | 4,09 | coiled-coil domain containing 51 | No | No | - |
| Col9a3 | 4,37E-04 | 4,12 | collagen, type IX, alpha 3 | No | No | - |

| | | | | | | |
|---|---|---|---|---|---|---|
| Coa6 | 1,79E-04 | 4,13 | cytochrome c oxidase assembly factor 6 | No | No | - |
| Dut | 2,60E-08 | 4,13 | deoxyuridine triphosphatase | No | No | - |
| 3110040N11Rik | 1,76E-04 | 4,14 | RIKEN cDNA 3110040N11 gene | No | No | - |
| Upp2 | 5,19E-03 | 4,15 | uridine phosphorylase 2 | No | No | - |
| Eloc | 1,05E-07 | 4,17 | elongin C | No | No | - |
| 1700016D06Rik | 8,62E-03 | 4,19 | RIKEN cDNA 1700016D06 gene | No | No | - |
| Ramac | 2,17E-07 | 4,19 | RNA guanine-7 methyltransferase activating subunit | No | No | - |
| Sycp3 | 2,19E-04 | 4,19 | synaptonemal complex protein 3 | No | No | - |
| Ager | 1,74E-05 | 4,19 | advanced glycosylation end product-specific receptor | No | No | 10 |
| Shc4 | 4,66E-04 | 4,20 | SHC (Src homology 2 domain containing) family, member | No | No | - |
| Pstk | 2,28E-05 | 4,21 | phosphoseryl-tRNA kinase | No | No | - |
| Sf3b6 | 4,20E-12 | 4,22 | splicing factor 3B, subunit 6 | No | No | - |
| Chpt1 | 1,13E-05 | 4,22 | choline phosphotransferase 1 | No | No | - |
| Pdxp | 2,98E-03 | 4,24 | pyridoxal (pyridoxine, vitamin B6) phosphatase | No | No | - |
| Rps23 | 1,95E-06 | 4,24 | ribosomal protein S23 | No | No | - |
| Tm4sf1 | 2,83E-11 | 4,24 | transmembrane 4 superfamily member 1 | No | No | - |
| Fgf11 | 2,04E-02 | 4,25 | fibroblast growth factor 11 | No | No | - |
| Gm13056 | 1,84E-02 | 4,25 | predicted gene 13056 | No | No | - |
| Ppie | 1,70E-02 | 4,25 | peptidylprolyl isomerase E (cyclophilin E) | No | No | 3 |
| Ccdc58 | 2,22E-06 | 4,26 | coiled-coil domain containing 58 | No | No | - |
| Slc35b3 | 6,18E-05 | 4,28 | solute carrier family 35, member B3 | No | No | - |
| Znrd1 | 6,68E-05 | 4,29 | RNA polymerase I subunit H | No | No | - |
| Tmem223 | 1,73E-09 | 4,29 | transmembrane protein 223 | No | No | - |
| Moap1 | 9,86E-03 | 4,30 | modulator of apoptosis 1 | No | No | - |
| Gm15421 | 1,15E-10 | 4,32 | predicted gene 15421 | No | No | - |
| Itpa | 5,19E-07 | 4,32 | inosine triphosphatase (nucleoside triphosphate pyrophosph | No | No | - |
| Gm20498 | 1,87E-10 | 4,32 | predicted gene 20498 | No | No | - |
| Maip1 | 1,03E-03 | 4,33 | matrix AAA peptidase interacting protein 1 | No | No | - |
| Ociad2 | 3,52E-04 | 4,33 | OCIA domain containing 2 | No | No | - |
| Pam16 | 1,87E-10 | 4,37 | presequence translocase-asssociated motor 16 | No | No | - |
| Ube2d1 | 1,38E-02 | 4,38 | ubiquitin-conjugating enzyme E2D 1 | No | No | - |
| Apbb3 | 7,89E-03 | 4,40 | amyloid beta (A4) precursor protein-binding, family B, men | No | No | - |
| Cenpp | 1,08E-02 | 4,40 | centromere protein P | No | No | 1 |
| Dtd2 | 8,77E-03 | 4,40 | D-tyrosyl-tRNA deacylase 2 | No | No | - |
| Med31 | 4,35E-04 | 4,40 | mediator complex subunit 31 | No | No | - |
| Pcna-ps2 | 1,30E-03 | 4,40 | proliferating cell nuclear antigen pseudogene 2 | No | No | - |
| Cdca3 | 3,65E-07 | 4,41 | cell division cycle associated 3 | No | No | - |
| Igfbp5 | 2,61E-03 | 4,43 | insulin-like growth factor binding protein 5 | No | No | 5 |
| Gtf2b | 2,49E-06 | 4,43 | general transcription factor IIB | No | No | - |
| H4c9 | 7,07E-05 | 4,43 | H4 clustered histone 9 | No | No | - |
| Crem | 2,81E-05 | 4,44 | cAMP responsive element modulator | No | No | - |
| Sec61g | 1,14E-07 | 4,44 | SEC61, gamma subunit | No | No | - |
| Isca2 | 5,87E-07 | 4,45 | iron-sulfur cluster assembly 2 | No | No | - |
| Rida | 4,47E-04 | 4,45 | reactive intermediate imine deaminase A homolog | No | No | - |
| Apmap | 1,14E-04 | 4,46 | adipocyte plasma membrane associated protein | No | No | - |
| Dnajc12 | 2,76E-03 | 4,46 | DnaJ heat shock protein family (Hsp40) member C12 | No | No | - |
| Gm5529 | 1,62E-06 | 4,49 | predicted pseudogene 5529 | No | No | - |
| Actr3-ps | 6,71E-03 | 4,50 | actin related protein 3, pseudogene | No | No | - |
| Gm10320 | 1,43E-02 | 4,50 | predicted pseudogene 10320 | No | No | - |
| Pigp | 1,13E-04 | 4,50 | phosphatidylinositol glycan anchor biosynthesis, class P | No | No | - |
| Ube2t | 3,54E-03 | 4,50 | ubiquitin-conjugating enzyme E2T | No | No | - |
| Ppp1r14b | 3,86E-11 | 4,52 | protein phosphatase 1, regulatory inhibitor subunit 14B | No | No | - |
| Dimt1 | 8,68E-04 | 4,52 | DIM1 dimethyladenosine transferase 1-like (S. cerevisiae) | No | No | - |
| Nudt21 | 4,51E-05 | 4,54 | nudix (nucleoside diphosphate linked moiety X)-type motif | No | No | 8 |
| Knstrn | 2,86E-05 | 4,54 | kinetochore-localized astrin/SPAG5 binding | No | No | - |
| Rnf5 | 1,18E-05 | 4,54 | ring finger protein 5 | No | No | - |
| Dnajc24 | 1,57E-03 | 4,55 | DnaJ heat shock protein family (Hsp40) member C24 | No | No | - |
| 4933423P22Rik | 1,75E-03 | 4,56 | RIKEN cDNA 4933423P22 gene | No | No | - |
| Rpl10-ps6 | 1,20E-08 | 4,57 | ribosomal protein L10, pseudogene 6 | No | No | - |
| Tma7 | 9,76E-07 | 4,57 | translational machinery associated 7 | No | No | - |
| Cetn4 | 5,08E-03 | 4,60 | centrin 4 | No | No | - |
| Fndc4 | 5,50E-03 | 4,60 | fibronectin type III domain containing 4 | No | No | - |
| Pdk1 | 5,46E-09 | 4,66 | pyruvate dehydrogenase kinase, isoenzyme 1 | No | No | 4 |
| Lsm8 | 6,28E-10 | 4,66 | LSM8 homolog, U6 small nuclear RNA associated | No | No | - |
| Macrod2 | 4,28E-03 | 4,67 | mono-ADP ribosylhydrolase 2 | No | No | - |
| Dtymk | 8,40E-08 | 4,68 | deoxythymidylate kinase | No | No | - |
| Mplkip | 3,66E-05 | 4,69 | M-phase specific PLK1 intereacting protein | No | No | - |
| Cks1b | 2,33E-10 | 4,69 | CDC28 protein kinase 1b | No | No | - |
| Mettl15 | 5,98E-05 | 4,69 | methyltransferase like 15 | No | No | - |
| Tmem215 | 2,76E-03 | 4,71 | transmembrane protein 215 | No | No | - |
| Siah1a | 4,12E-05 | 4,71 | siah E3 ubiquitin protein ligase 1A | No | No | - |
| Psph | 2,83E-11 | 4,72 | phosphoserine phosphatase | No | No | - |
| Hspb11 | 2,22E-04 | 4,73 | heat shock protein family B (small), member 11 | No | No | - |

| | | | | | | |
|---|---|---|---|---|---|---|
| Oard1 | 3,06E-03 | 4,73 | O-acyl-ADP-ribose deacylase 1 | No | No | - |
| Mrps14 | 3,99E-11 | 4,74 | mitochondrial ribosomal protein S14 | No | No | - |
| AV356131 | 9,58E-03 | 4,75 | expressed sequence AV356131 | No | No | - |
| Msl3l2 | 9,02E-03 | 4,75 | MSL3 like 2 | No | No | - |
| Zswim7 | 2,60E-07 | 4,75 | zinc finger SWIM-type containing 7 | No | No | - |
| Mrps36 | 5,35E-05 | 4,78 | mitochondrial ribosomal protein S36 | No | No | - |
| Oip5 | 2,42E-03 | 4,79 | Opa interacting protein 5 | No | No | - |
| Sdhaf3 | 2,05E-05 | 4,79 | succinate dehydrogenase complex assembly factor 3 | No | No | - |
| Gramd4 | 3,43E-05 | 4,80 | GRAM domain containing 4 | No | No | - |
| Nr4a2 | 2,09E-04 | 4,80 | nuclear receptor subfamily 4, group A, member 2 | No | No | - |
| Rbmxl1 | 5,93E-05 | 4,81 | RNA binding motif protein, X-linked like-1 | No | No | - |
| Hoxa2 | 1,46E-05 | 4,81 | homeobox A2 | No | No | - |
| Selenof | 3,35E-10 | 4,84 | selenoprotein F | No | No | - |
| Mtx2 | 1,56E-09 | 4,84 | metaxin 2 | No | No | - |
| Fbxo8 | 3,93E-07 | 4,86 | F-box protein 8 | No | No | - |
| Rnf113a2 | 8,35E-04 | 4,88 | ring finger protein 113A2 | No | No | - |
| Oxnad1 | 9,37E-08 | 4,88 | oxidoreductase NAD-binding domain containing 1 | No | No | - |
| Epm2a | 3,32E-03 | 4,90 | epilepsy, progressive myoclonic epilepsy, type 2 gene alpha | No | No | - |
| Dynlt1f | 2,75E-03 | 4,91 | dynein light chain Tctex-type 1F | No | No | - |
| Anln | 1,76E-03 | 4,92 | anillin, actin binding protein | No | No | - |
| Asf1a | 6,86E-04 | 4,94 | anti-silencing function 1A histone chaperone | No | No | - |
| Sdhd | 7,19E-06 | 4,97 | succinate dehydrogenase complex, subunit D, integral mem | No | No | - |
| Mrpl32 | 1,33E-06 | 4,98 | mitochondrial ribosomal protein L32 | No | No | - |
| Clcn2 | 8,37E-05 | 5,00 | chloride channel, voltage-sensitive 2 | No | No | - |
| Gm15050 | 1,07E-08 | 5,00 | predicted gene 15050 | No | No | - |
| Gm7488 | 3,66E-03 | 5,00 | predicted gene 7488 | No | No | - |
| Gmnn | 4,21E-04 | 5,00 | geminin | No | No | - |
| H3f3a | 5,48E-09 | 5,01 | H3.3 histone A | No | No | - |
| Higd1a | 2,85E-12 | 5,02 | HIG1 domain family, member 1A | No | No | - |
| Cryz | 2,80E-07 | 5,03 | crystallin, zeta | No | No | - |
| Smim26 | 1,67E-06 | 5,04 | small integral membrane protein 26 | No | No | - |
| Lgi3 | 2,54E-04 | 5,06 | leucine-rich repeat LGI family, member 3 | No | No | - |
| Fam131a | 7,78E-05 | 5,08 | family with sequence similarity 131, member A | No | No | - |
| Pdcl3 | 7,32E-10 | 5,09 | phosducin-like 3 | No | No | - |
| Nat2 | 1,07E-04 | 5,10 | N-acetyltransferase 2 (arylamine N-acetyltransferase) | No | No | 1 |
| Cplx2 | 8,20E-03 | 5,14 | complexin 2 | No | No | - |
| Gm27167 | 5,38E-04 | 5,15 | predicted gene 27167 | No | No | - |
| Mirg | 5,83E-06 | 5,19 | miRNA containing gene | No | No | - |
| Dpy30 | 1,13E-08 | 5,20 | dpy-30, histone methyltransferase complex regulatory subu | No | No | - |
| Igfbp4 | 1,54E-12 | 5,23 | insulin-like growth factor binding protein 4 | No | No | - |
| H2-Ab1 | 1,07E-03 | 5,23 | histocompatibility 2, class II antigen A, beta 1 | No | No | 27 |
| Tcim | 7,13E-05 | 5,26 | transcriptional and immune response regulator | No | No | - |
| Rpl22l1 | 7,63E-14 | 5,27 | ribosomal protein L22 like 1 | No | No | - |
| Wdr5b | 1,98E-15 | 5,29 | WD repeat domain 5B | No | No | - |
| Acyp1 | 4,05E-03 | 5,30 | acylphosphatase 1, erythrocyte (common) type | No | No | - |
| Cdc26 | 8,09E-07 | 5,30 | cell division cycle 26 | No | No | - |
| Tram1l1 | 3,06E-03 | 5,33 | translocation associated membrane protein 1-like 1 | No | No | - |
| Snhg8 | 1,60E-14 | 5,37 | small nucleolar RNA host gene 8 | No | No | - |
| Nme4 | 3,09E-06 | 5,38 | NME/NM23 nucleoside diphosphate kinase 4 | No | No | - |
| Cdc45 | 8,66E-04 | 5,38 | cell division cycle 45 | No | No | - |
| Gm7041 | 3,68E-09 | 5,39 | predicted gene 7041 | No | No | - |
| Camta1 | 6,30E-07 | 5,39 | calmodulin binding transcription activator 1 | No | No | - |
| Syce2 | 1,75E-03 | 5,40 | synaptonemal complex central element protein 2 | No | No | - |
| Polr2k | 1,52E-06 | 5,40 | polymerase (RNA) II (DNA directed) polypeptide K | No | No | - |
| Snhg4 | 2,91E-08 | 5,43 | small nucleolar RNA host gene 4 | No | No | - |
| Memo1 | 3,22E-09 | 5,44 | mediator of cell motility 1 | No | No | - |
| Commd1 | 1,36E-07 | 5,44 | COMM domain containing 1 | No | No | - |
| Stoml2 | 3,73E-08 | 5,46 | stomatin (Epb7.2)-like 2 | No | No | - |
| Arl5c | 2,17E-12 | 5,49 | ADP-ribosylation factor-like 5C | No | No | - |
| Gm6576 | 8,00E-04 | 5,50 | predicted gene 6576 | No | No | - |
| Nemp1 | 2,80E-03 | 5,56 | nuclear envelope integral membrane protein 1 | No | No | - |
| Fam162a | 5,33E-16 | 5,58 | family with sequence similarity 162, member A | No | No | - |
| Exosc3 | 8,74E-04 | 5,60 | exosome component 3 | No | No | - |
| Tcf21 | 1,98E-03 | 5,60 | transcription factor 21 | No | No | - |
| Net1 | 1,28E-03 | 5,64 | neuroepithelial cell transforming gene 1 | No | No | - |
| Hilpda | 5,70E-17 | 5,71 | hypoxia inducible lipid droplet associated | No | No | - |
| Ciao2a | 1,59E-09 | 5,75 | cytosolic iron-sulfur assembly component 2A | No | No | - |
| A430005L14Rik | 1,79E-09 | 5,76 | RIKEN cDNA A430005L14 gene | No | No | - |
| Selenbp1 | 4,42E-03 | 5,78 | selenium binding protein 1 | No | No | - |
| Gm45716 | 6,48E-05 | 5,78 | small integral membrane protein 36 | No | No | - |
| Bnip3 | 1,19E-11 | 5,79 | BCL2/adenovirus E1B interacting protein 3 | No | No | - |
| Snhg6 | 3,31E-09 | 5,80 | small nucleolar RNA host gene 6 | No | No | - |
| Dynlt1c | 2,51E-04 | 5,81 | dynein light chain Tctex-type 1C | No | No | - |

| | | | | | | |
|---|---|---|---|---|---|---|
| Ska1 | 5,11E-06 | 5,84 | spindle and kinetochore associated complex subunit 1 | No | No | - |
| Cenpa | 2,37E-10 | 5,85 | centromere protein A | No | No | 2 |
| En1 | 2,35E-03 | 5,88 | engrailed 1 | No | No | - |
| 9430038I01Rik | 3,42E-06 | 5,91 | RIKEN cDNA 9430038I01 gene | No | No | - |
| Mcmdc2 | 2,58E-13 | 6,04 | minichromosome maintenance domain containing 2 | No | No | - |
| Mmp13 | 5,83E-06 | 6,15 | matrix metallopeptidase 13 | Yes | No | 19 |
| Gm40155 | 1,07E-03 | 6,22 | predicted gene, 40155 | No | No | - |
| Gm33051 | 6,86E-18 | 6,26 | predicted gene, 33051 | No | No | - |
| H2az1 | 1,18E-13 | 6,28 | H2A.Z variant histone 1 | No | No | - |
| Tm2d1 | 3,41E-06 | 6,50 | TM2 domain containing 1 | No | No | - |
| Prcp | 3,10E-05 | 6,53 | prolylcarboxypeptidase (angiotensinase C) | No | No | - |
| Cdkn2d | 1,20E-14 | 6,64 | cyclin dependent kinase inhibitor 2D | No | No | 1 |
| Nup37 | 2,54E-07 | 6,82 | nucleoporin 37 | No | No | - |
| Cenpw | 7,69E-16 | 6,84 | centromere protein W | No | No | - |
| Mrps31 | 8,74E-05 | 6,86 | mitochondrial ribosomal protein S31 | No | No | - |
| Lockd | 3,46E-04 | 6,89 | lncRNA downstream of Cdkn1b | No | No | - |
| Tnfsf8 | 4,61E-04 | 6,89 | tumor necrosis factor (ligand) superfamily, member 8 | No | No | - |
| Cthrc1 | 1,96E-12 | 7,02 | collagen triple helix repeat containing 1 | No | No | 4 |
| D630033A02Rik | 7,22E-06 | 7,10 | RIKEN cDNA D630033A02 gene | No | No | - |
| Fgl2 | 1,45E-03 | 7,33 | fibrinogen-like protein 2 | No | No | - |
| G0s2 | 2,61E-06 | 7,33 | G0/G1 switch gene 2 | No | No | - |
| Rgcc | 4,62E-05 | 7,33 | regulator of cell cycle | No | No | 9 |
| Gm7984 | 9,60E-08 | 7,38 | predicted gene 7984 | No | No | - |
| Satb1 | 6,52E-04 | 7,50 | special AT-rich sequence binding protein 1 | No | No | - |
| Cd24a | 6,89E-08 | 7,51 | CD24a antigen | No | No | - |
| 0610009B22Rik | 3,24E-09 | 7,65 | RIKEN cDNA 0610009B22 gene | No | No | - |
| Ncald | 1,26E-03 | 7,67 | neurocalcin delta | No | No | - |
| Snhg3 | 3,15E-14 | 7,71 | small nucleolar RNA host gene 3 | No | No | - |
| Cdk1 | 5,33E-16 | 7,73 | cyclin-dependent kinase 1 | No | No | 1 |
| Ccdc38 | 2,43E-10 | 7,82 | coiled-coil domain containing 38 | No | No | - |
| Tceal1 | 4,76E-05 | 7,85 | transcription elongation factor A (SII)-like 1 | No | No | - |
| Twist1 | 2,17E-07 | 7,94 | twist basic helix-loop-helix transcription factor 1 | No | No | 5 |
| Plac8 | 3,05E-16 | 8,07 | placenta-specific 8 | No | No | - |
| Nup43 | 1,67E-03 | 8,20 | nucleoporin 43 | No | No | - |
| Sap30 | 1,42E-04 | 8,33 | sin3 associated polypeptide | No | No | - |
| Nt5e | 2,46E-08 | 8,35 | 5' nucleotidase, ecto | No | No | - |
| Dtwd1 | 2,40E-08 | 8,38 | DTW domain containing 1 | No | No | - |
| Mgl2 | 2,44E-05 | 8,40 | macrophage galactose N-acetyl-galactosamine specific lecti | No | No | - |
| Alkal1 | 2,07E-11 | 8,43 | ALK and LTK ligand 1 | No | No | - |
| Prelid2 | 6,44E-16 | 8,52 | PRELI domain containing 2 | No | No | - |
| Slc38a4 | 2,54E-04 | 8,55 | solute carrier family 38, member 4 | No | No | - |
| Esm1 | 4,66E-08 | 8,56 | endothelial cell-specific molecule 1 | No | No | - |
| Lyplal1 | 1,42E-04 | 8,63 | lysophospholipase-like 1 | No | No | - |
| Med30 | 2,94E-06 | 8,73 | mediator complex subunit 30 | No | No | - |
| Olfr12 | 1,37E-03 | 8,80 | olfactory receptor 12 | No | No | - |
| AA465934 | 9,33E-04 | 9,00 | expressed sequence AA465934 | No | No | - |
| Egln3 | 1,85E-09 | 9,14 | egl-9 family hypoxia-inducible factor 3 | No | No | - |
| 2500002B13Rik | 6,33E-04 | 9,20 | RIKEN cDNA 2500002B13 gene | No | No | - |
| Kcnj2 | 1,60E-03 | 9,25 | potassium inwardly-rectifying channel, subfamily J, membe | No | No | 9 |
| Nrep | 1,83E-15 | 9,31 | neuronal regeneration related protein | No | No | - |
| Cth | 6,77E-12 | 9,34 | cystathionase (cystathionine gamma-lyase) | No | No | 2 |
| Snrpf | 1,39E-09 | 9,36 | small nuclear ribonucleoprotein polypeptide F | No | No | - |
| Rpa3 | 1,73E-05 | 9,38 | replication protein A3 | No | No | - |
| Dkk2 | 6,16E-05 | 9,50 | dickkopf WNT signaling pathway inhibitor 2 | No | No | - |
| Rps16 | 1,25E-14 | 9,59 | ribosomal protein S16 | No | No | - |
| Id3 | 5,69E-19 | 10,13 | inhibitor of DNA binding 3 | No | No | 3 |
| Tpd52l1 | 9,13E-04 | 10,50 | tumor protein D52-like 1 | No | No | - |
| Gm15640 | 7,49E-14 | 10,69 | predicted gene 15640 | No | No | - |
| Raet1e | 5,18E-10 | 11,06 | retinoic acid early transcript 1E | No | No | - |
| Etfbkmt | 5,61E-07 | 12,42 | electron transfer flavoprotein beta subunit lysine methyltran | No | No | - |
| Selenoh | 2,78E-20 | 12,69 | selenoprotein H | No | No | - |
| Pcdh10 | 3,30E-06 | 13,00 | protocadherin 10 | No | No | - |
| Adora2b | 1,82E-09 | 13,08 | adenosine A2b receptor | No | No | 18 |
| Sft2d1 | 2,94E-04 | 13,50 | SFT2 domain containing 1 | No | No | - |
| Gm10420 | 6,41E-06 | 13,63 | predicted gene 10420 | No | No | - |
| Mgarp | 4,09E-17 | 13,74 | mitochondria localized glutamic acid rich protein | No | No | - |
| Mmp10 | 7,05E-18 | 14,27 | matrix metallopeptidase 10 | No | No | - |
| S100a7a | 4,51E-05 | 15,22 | S100 calcium binding protein A7A | No | No | - |
| Raet1d | 9,16E-07 | 15,60 | retinoic acid early transcript delta | No | No | - |
| Gm15247 | 3,82E-08 | 18,33 | predicted gene 15247 | No | No | - |
| Gas1 | 2,20E-09 | 18,93 | growth arrest specific 1 | No | No | - |
| Bambi | 6,79E-09 | 23,50 | BMP and activin membrane-bound inhibitor | No | No | 3 |
| Thap8 | 9,06E-07 | 24,00 | RIKEN cDNA 1700020L13 gene | No | No | - |

| | | | | | | |
|---|---|---|---|---|---|---|
| Mcpt8 | 1,80E-10 | 26,00 | mast cell protease 8 | No | No | - |
| Nap1l5 | 5,04E-06 | 26,33 | nucleosome assembly protein 1-like 5 | No | No | - |
| Rasl11a | 5,27E-07 | 29,33 | RAS-like, family 11, member A | No | No | - |
| Spon2 | 2,19E-15 | 33,75 | spondin 2, extracellular matrix protein | No | No | - |
| Sult4a1 | 1,05E-07 | 35,00 | sulfotransferase family 4A, member 1 | No | No | - |
| Lsm5 | 8,25E-07 | 52,00 | LSM5 homolog, U6 small nuclear RNA and mRNA degrada | No | No | - |
| Nrn1 | 1,88E-10 | 187,00 | neuritin 1 | No | No | - |

**Table S5: Deregulated pathways upon Tks5 haploinsufficiency and treatment with TGFb**

| Ontology | GO ID | Description | setSize | NES | p.adjust |
|---|---|---|---|---|---|
| CC | 62023 | collagen-containing extracellular matrix | 203 | -2,71 | 5,19E-18 |
| MF | 5201 | extracellular matrix structural constituent | 83 | -2,55 | 1,59E-09 |
| CC | 30312 | external encapsulating structure | 246 | -2,54 | 2,52E-16 |
| CC | 31012 | extracellular matrix | 246 | -2,54 | 2,52E-16 |
| MF | 8201 | heparin binding | 81 | -2,53 | 1,78E-09 |
| CC | 5604 | basement membrane | 65 | -2,49 | 5,25E-08 |
| MF | 5539 | glycosaminoglycan binding | 106 | -2,47 | 1,69E-09 |
| BP | 71621 | granulocyte chemotaxis | 54 | -2,35 | 2,78E-06 |
| BP | 97530 | granulocyte migration | 65 | -2,33 | 2,76E-06 |
| MF | 1901681 | sulfur compound binding | 142 | -2,31 | 4,96E-09 |
| BP | 32103 | positive regulation of response to external stimulus | 212 | -2,29 | 6,80E-10 |
| BP | 43277 | apoptotic cell clearance | 24 | -2,26 | 9,98E-05 |
| BP | 30198 | extracellular matrix organization | 178 | -2,26 | 7,88E-09 |
| BP | 43062 | extracellular structure organization | 178 | -2,26 | 7,88E-09 |
| BP | 45229 | external encapsulating structure organization | 178 | -2,26 | 7,88E-09 |
| BP | 97242 | amyloid-beta clearance | 22 | -2,26 | 1,87E-04 |
| CC | 9986 | cell surface | 345 | -2,26 | 2,99E-13 |
| BP | 45087 | innate immune response | 316 | -2,25 | 8,75E-13 |
| MF | 50840 | extracellular matrix binding | 37 | -2,25 | 4,39E-05 |
| MF | 4867 | serine-type endopeptidase inhibitor activity | 24 | -2,23 | 1,60E-04 |
| BP | 6935 | chemotaxis | 277 | -2,22 | 4,02E-11 |
| BP | 1568 | blood vessel development | 460 | -2,22 | 1,93E-15 |
| MF | 5509 | calcium ion binding | 261 | -2,21 | 4,65E-10 |
| BP | 30595 | leukocyte chemotaxis | 103 | -2,21 | 2,32E-06 |
| CC | 34358 | plasma lipoprotein particle | 15 | -2,20 | 2,56E-04 |
| CC | 1990777 | lipoprotein particle | 15 | -2,20 | 2,56E-04 |
| BP | 30316 | osteoclast differentiation | 64 | -2,20 | 3,26E-05 |
| BP | 98542 | defense response to other organism | 428 | -2,20 | 3,77E-14 |

| | | | | | |
|---|---|---|---|---|---|
| BP | 1944 | vasculature development | 484 | -2,19 | 4,49E-15 |
| BP | 6953 | acute-phase response | 12 | -2,19 | 2,12E-04 |
| BP | 42330 | taxis | 280 | -2,19 | 2,53E-10 |
| BP | 2062 | chondrocyte differentiation | 68 | -2,18 | 3,26E-05 |
| BP | 60326 | cell chemotaxis | 144 | -2,18 | 4,78E-07 |
| MF | 43394 | proteoglycan binding | 23 | -2,17 | 3,21E-04 |
| MF | 5178 | integrin binding | 85 | -2,16 | 2,45E-05 |
| BP | 1525 | angiogenesis | 333 | -2,16 | 2,55E-11 |
| CC | 9925 | basal plasma membrane | 121 | -2,16 | 7,96E-07 |
| BP | 2526 | acute inflammatory response | 41 | -2,15 | 2,37E-04 |
| BP | 48514 | blood vessel morphogenesis | 397 | -2,15 | 3,47E-12 |
| MF | 50839 | cell adhesion molecule binding | 145 | -2,15 | 6,60E-07 |
| MF | 30546 | signaling receptor activator activity | 150 | -2,14 | 2,21E-07 |
| BP | 50729 | positive regulation of inflammatory response | 77 | -2,14 | 3,41E-05 |
| MF | 48018 | receptor ligand activity | 146 | -2,14 | 5,05E-07 |
| BP | 19221 | cytokine-mediated signaling pathway | 180 | -2,14 | 2,35E-07 |
| BP | 1990266 | neutrophil migration | 51 | -2,13 | 2,00E-04 |
| CC | 45178 | basal part of cell | 132 | -2,13 | 7,59E-07 |
| BP | 71622 | regulation of granulocyte chemotaxis | 24 | -2,12 | 7,81E-04 |
| BP | 97529 | myeloid leukocyte migration | 102 | -2,12 | 2,35E-05 |
| BP | 30335 | positive regulation of cell migration | 360 | -2,12 | 4,02E-11 |
| BP | 1945 | lymph vessel development | 20 | -2,12 | 8,50E-04 |
| BP | 6954 | inflammatory response | 336 | -2,12 | 1,21E-10 |
| BP | 30593 | neutrophil chemotaxis | 41 | -2,11 | 4,85E-04 |
| BP | 43410 | positive regulation of MAPK cascade | 274 | -2,10 | 4,71E-09 |
| CC | 43235 | receptor complex | 155 | -2,10 | 1,60E-06 |
| BP | 48771 | tissue remodeling | 111 | -2,10 | 1,11E-05 |
| BP | 1901342 | regulation of vasculature development | 181 | -2,10 | 7,78E-07 |
| BP | 40017 | positive regulation of locomotion | 372 | -2,10 | 7,63E-11 |
| BP | 6956 | complement activation | 18 | -2,10 | 8,27E-04 |
| BP | 50817 | coagulation | 77 | -2,10 | 6,48E-05 |

| | | | | | |
|---|---|---|---|---|---|
| MF | 5044 | scavenger receptor activity | 16 | -2,09 | 1,13E-03 |
| BP | 45765 | regulation of angiogenesis | 179 | -2,09 | 1,41E-06 |
| CC | 32994 | protein-lipid complex | 18 | -2,09 | 8,93E-04 |
| BP | 16322 | neuron remodeling | 10 | -2,09 | 6,82E-04 |
| BP | 43087 | regulation of GTPase activity | 193 | -2,09 | 4,94E-07 |
| BP | 2000147 | positive regulation of cell motility | 367 | -2,08 | 2,53E-10 |
| CC | 45177 | apical part of cell | 200 | -2,08 | 6,60E-07 |
| MF | 4713 | protein tyrosine kinase activity | 64 | -2,07 | 2,26E-04 |
| BP | 1903555 | regulation of tumor necrosis factor superfamily cytokine pr | 97 | -2,07 | 1,95E-05 |
| CC | 5903 | brush border | 58 | -2,07 | 3,89E-04 |
| MF | 30545 | receptor regulator activity | 163 | -2,07 | 4,10E-06 |
| BP | 7596 | blood coagulation | 75 | -2,07 | 1,11E-04 |
| BP | 51272 | positive regulation of cellular component movement | 378 | -2,07 | 2,61E-10 |
| BP | 50921 | positive regulation of chemotaxis | 81 | -2,06 | 9,96E-05 |
| BP | 2252 | immune effector process | 371 | -2,06 | 4,67E-10 |
| BP | 42060 | wound healing | 197 | -2,06 | 8,44E-07 |
| BP | 71706 | tumor necrosis factor superfamily cytokine production | 102 | -2,06 | 5,72E-05 |
| MF | 19199 | transmembrane receptor protein kinase activity | 35 | -2,06 | 9,80E-04 |
| CC | 31594 | neuromuscular junction | 50 | -2,06 | 4,65E-04 |
| BP | 2573 | myeloid leukocyte differentiation | 118 | -2,06 | 1,42E-05 |
| BP | 60840 | artery development | 68 | -2,06 | 2,09E-04 |
| BP | 90630 | activation of GTPase activity | 61 | -2,06 | 3,37E-04 |
| CC | 98862 | cluster of actin-based cell projections | 79 | -2,06 | 1,46E-04 |
| CC | 5765 | lysosomal membrane | 100 | -2,06 | 5,12E-05 |
| CC | 98852 | lytic vacuole membrane | 100 | -2,06 | 5,12E-05 |
| BP | 31349 | positive regulation of defense response | 132 | -2,05 | 5,40E-06 |
| BP | 9615 | response to virus | 167 | -2,05 | 2,09E-06 |
| BP | 42552 | myelination | 86 | -2,05 | 1,07E-04 |
| CC | 16323 | basolateral plasma membrane | 106 | -2,05 | 3,29E-05 |
| BP | 45766 | positive regulation of angiogenesis | 109 | -2,05 | 1,67E-05 |
| BP | 1904018 | positive regulation of vasculature development | 109 | -2,05 | 1,67E-05 |

| | | | | | |
|---|---|---|---|---|---|
| BP | 35455 | response to interferon-alpha | 13 | -2,05 | 1,37E-03 |
| BP | 10758 | regulation of macrophage chemotaxis | 16 | -2,05 | 2,04E-03 |
| BP | 32680 | regulation of tumor necrosis factor production | 96 | -2,05 | 1,10E-04 |
| MF | 45499 | chemorepellent activity | 12 | -2,05 | 2,65E-03 |
| BP | 2690 | positive regulation of leukocyte chemotaxis | 47 | -2,05 | 4,38E-04 |
| BP | 7599 | hemostasis | 77 | -2,05 | 1,41E-04 |
| BP | 33209 | tumor necrosis factor-mediated signaling pathway | 45 | -2,05 | 8,85E-04 |
| BP | 50900 | leukocyte migration | 163 | -2,04 | 7,75E-06 |
| BP | 1952 | regulation of cell-matrix adhesion | 87 | -2,04 | 8,74E-05 |
| BP | 30501 | positive regulation of bone mineralization | 22 | -2,04 | 4,01E-03 |
| MF | 15020 | glucuronosyltransferase activity | 18 | -2,04 | 1,62E-03 |
| MF | 43236 | laminin binding | 19 | -2,04 | 2,87E-03 |
| BP | 2683 | negative regulation of immune system process | 202 | -2,04 | 2,32E-06 |
| BP | 45670 | regulation of osteoclast differentiation | 37 | -2,04 | 1,11E-03 |
| BP | 8347 | glial cell migration | 42 | -2,04 | 1,02E-03 |
| BP | 1817 | regulation of cytokine production | 356 | -2,03 | 1,23E-09 |
| BP | 32640 | tumor necrosis factor production | 101 | -2,03 | 1,07E-04 |
| BP | 2888 | positive regulation of myeloid leukocyte mediated immunit | 13 | -2,03 | 1,96E-03 |
| CC | 16324 | apical plasma membrane | 146 | -2,03 | 7,75E-06 |
| BP | 7178 | transmembrane receptor protein serine/threonine kinase sig | 221 | -2,03 | 1,05E-06 |
| BP | 43900 | regulation of multi-organism process | 20 | -2,03 | 2,07E-03 |
| BP | 32801 | receptor catabolic process | 22 | -2,03 | 5,04E-03 |
| BP | 50731 | positive regulation of peptidyl-tyrosine phosphorylation | 91 | -2,02 | 1,60E-04 |
| CC | 42641 | actomyosin | 72 | -2,02 | 3,97E-04 |
| BP | 50920 | regulation of chemotaxis | 117 | -2,02 | 6,93E-05 |
| BP | 9593 | detection of chemical stimulus | 12 | -2,02 | 3,64E-03 |
| BP | 45071 | negative regulation of viral genome replication | 26 | -2,02 | 2,44E-03 |
| BP | 7160 | cell-matrix adhesion | 137 | -2,02 | 2,24E-05 |
| CC | 5925 | focal adhesion | 120 | -2,02 | 6,83E-05 |
| BP | 71711 | basement membrane organization | 20 | -2,02 | 2,29E-03 |
| BP | 71675 | regulation of mononuclear cell migration | 60 | -2,01 | 6,35E-04 |

| | | | | | |
|---|---|---|---|---|---|
| BP | 31644 | regulation of nervous system process | 65 | -2,01 | 6,28E-04 |
| BP | 1905521 | regulation of macrophage migration | 23 | -2,01 | 2,11E-03 |
| BP | 6898 | receptor-mediated endocytosis | 137 | -2,01 | 2,47E-05 |
| BP | 3279 | cardiac septum development | 79 | -2,01 | 2,56E-04 |
| BP | 150115 | cell-substrate junction organization | 79 | -2,01 | 2,66E-04 |
| BP | 32101 | regulation of response to external stimulus | 464 | -2,01 | 1,21E-10 |
| BP | 7272 | ensheathment of neurons | 88 | -2,01 | 1,41E-04 |
| BP | 8366 | axon ensheathment | 88 | -2,01 | 1,41E-04 |
| BP | 10759 | positive regulation of macrophage chemotaxis | 12 | -2,01 | 4,16E-03 |
| BP | 1774 | microglial cell activation | 23 | -2,01 | 2,29E-03 |
| BP | 61098 | positive regulation of protein tyrosine kinase activity | 20 | -2,00 | 2,56E-03 |
| BP | 2761 | regulation of myeloid leukocyte differentiation | 64 | -2,00 | 6,25E-04 |
| BP | 71347 | cellular response to interleukin-1 | 49 | -2,00 | 1,75E-03 |
| CC | 31526 | brush border membrane | 27 | -2,00 | 4,24E-03 |
| BP | 51607 | defense response to virus | 138 | -2,00 | 4,00E-05 |
| BP | 140546 | defense response to symbiont | 138 | -2,00 | 4,00E-05 |
| BP | 48015 | phosphatidylinositol-mediated signaling | 86 | -2,00 | 2,66E-04 |
| CC | 32432 | actin filament bundle | 70 | -1,99 | 9,71E-04 |
| CC | 42734 | presynaptic membrane | 72 | -1,99 | 5,74E-04 |
| BP | 7229 | integrin-mediated signaling pathway | 58 | -1,99 | 1,29E-03 |
| BP | 31589 | cell-substrate adhesion | 215 | -1,99 | 4,85E-06 |
| CC | 5901 | caveola | 60 | -1,99 | 8,85E-04 |
| CC | 30055 | cell-substrate junction | 125 | -1,98 | 6,60E-05 |
| BP | 71674 | mononuclear cell migration | 85 | -1,98 | 4,67E-04 |
| BP | 150077 | regulation of neuroinflammatory response | 15 | -1,98 | 5,83E-03 |
| BP | 30155 | regulation of cell adhesion | 384 | -1,98 | 6,37E-09 |
| MF | 4866 | endopeptidase inhibitor activity | 60 | -1,98 | 9,42E-04 |
| BP | 1905523 | positive regulation of macrophage migration | 15 | -1,98 | 5,88E-03 |
| BP | 51056 | regulation of small GTPase mediated signal transduction | 169 | -1,98 | 7,92E-06 |
| BP | 48017 | inositol lipid-mediated signaling | 87 | -1,98 | 2,75E-04 |
| BP | 1903557 | positive regulation of tumor necrosis factor superfamily cyt | 63 | -1,98 | 8,27E-04 |

| | | | | | |
|---|---|---|---|---|---|
| BP | 34612 | response to tumor necrosis factor | 108 | -1,98 | 1,56E-04 |
| BP | 31641 | regulation of myelination | 34 | -1,98 | 4,04E-03 |
| BP | 2688 | regulation of leukocyte chemotaxis | 58 | -1,98 | 1,46E-03 |
| BP | 9617 | response to bacterium | 262 | -1,98 | 6,64E-07 |
| BP | 32606 | type I interferon production | 62 | -1,97 | 9,41E-04 |
| BP | 71356 | cellular response to tumor necrosis factor | 94 | -1,97 | 8,85E-04 |
| CC | 5887 | integral component of plasma membrane | 400 | -1,97 | 7,31E-09 |
| BP | 32479 | regulation of type I interferon production | 56 | -1,97 | 1,81E-03 |
| MF | 31593 | polyubiquitin modification-dependent protein binding | 42 | -1,97 | 2,79E-03 |
| BP | 45453 | bone resorption | 47 | -1,97 | 1,46E-03 |
| BP | 90092 | regulation of transmembrane receptor protein serine/threoni | 154 | -1,97 | 2,29E-05 |
| BP | 50878 | regulation of body fluid levels | 149 | -1,97 | 2,45E-05 |
| BP | 2684 | positive regulation of immune system process | 384 | -1,97 | 1,06E-08 |
| CC | 9897 | external side of plasma membrane | 131 | -1,96 | 4,69E-05 |
| BP | 3205 | cardiac chamber development | 107 | -1,96 | 1,11E-04 |
| CC | 44853 | plasma membrane raft | 72 | -1,96 | 7,81E-04 |
| BP | 1990000 | amyloid fibril formation | 20 | -1,96 | 3,76E-03 |
| CC | 5767 | secondary lysosome | 11 | -1,96 | 3,17E-03 |
| CC | 16328 | lateral plasma membrane | 33 | -1,96 | 4,67E-03 |
| CC | 45111 | intermediate filament cytoskeleton | 63 | -1,96 | 1,15E-03 |
| CC | 31253 | cell projection membrane | 135 | -1,96 | 7,05E-05 |
| BP | 32835 | glomerulus development | 35 | -1,96 | 3,81E-03 |
| MF | 19838 | growth factor binding | 92 | -1,96 | 3,66E-04 |
| BP | 60037 | pharyngeal system development | 11 | -1,96 | 3,25E-03 |
| BP | 7411 | axon guidance | 106 | -1,96 | 1,60E-04 |
| BP | 97485 | neuron projection guidance | 106 | -1,96 | 1,60E-04 |
| BP | 46660 | female sex differentiation | 68 | -1,96 | 8,86E-04 |
| BP | 42327 | positive regulation of phosphorylation | 496 | -1,95 | 6,63E-10 |
| BP | 1819 | positive regulation of cytokine production | 225 | -1,95 | 5,71E-06 |
| MF | 15081 | sodium ion transmembrane transporter activity | 39 | -1,95 | 3,99E-03 |
| CC | 97440 | apical dendrite | 15 | -1,95 | 7,45E-03 |

| | | | | | |
|---|---|---|---|---|---|
| BP | 48844 | artery morphogenesis | 49 | -1,95 | 3,17E-03 |
| BP | 52695 | cellular glucuronidation | 10 | -1,95 | 5,80E-03 |
| BP | 1816 | cytokine production | 400 | -1,95 | 1,86E-08 |
| BP | 2763 | positive regulation of myeloid leukocyte differentiation | 27 | -1,94 | 6,82E-03 |
| BP | 51965 | positive regulation of synapse assembly | 26 | -1,94 | 5,50E-03 |
| BP | 46849 | bone remodeling | 59 | -1,94 | 1,38E-03 |
| BP | 36303 | lymph vessel morphogenesis | 15 | -1,94 | 7,79E-03 |
| BP | 50766 | positive regulation of phagocytosis | 31 | -1,94 | 4,64E-03 |
| BP | 60976 | coronary vasculature development | 45 | -1,94 | 3,17E-03 |
| BP | 32760 | positive regulation of tumor necrosis factor production | 62 | -1,94 | 1,44E-03 |
| BP | 50727 | regulation of inflammatory response | 184 | -1,94 | 2,00E-05 |
| BP | 3208 | cardiac ventricle morphogenesis | 40 | -1,94 | 4,17E-03 |
| BP | 9611 | response to wounding | 265 | -1,93 | 1,34E-06 |
| BP | 44057 | regulation of system process | 265 | -1,93 | 1,34E-06 |
| BP | 1541 | ovarian follicle development | 35 | -1,93 | 5,17E-03 |
| BP | 34330 | cell junction organization | 382 | -1,93 | 1,48E-08 |
| BP | 45672 | positive regulation of osteoclast differentiation | 14 | -1,93 | 8,12E-03 |
| BP | 90287 | regulation of cellular response to growth factor stimulus | 186 | -1,93 | 5,91E-05 |
| MF | 30414 | peptidase inhibitor activity | 65 | -1,93 | 1,87E-03 |
| BP | 97435 | supramolecular fiber organization | 454 | -1,93 | 4,60E-09 |
| BP | 35904 | aorta development | 43 | -1,93 | 5,64E-03 |
| BP | 43408 | regulation of MAPK cascade | 391 | -1,93 | 2,11E-08 |
| CC | 5905 | clathrin-coated pit | 54 | -1,93 | 2,42E-03 |
| CC | 98858 | actin-based cell projection | 113 | -1,93 | 1,99E-04 |
| BP | 7044 | cell-substrate junction assembly | 73 | -1,93 | 8,93E-04 |
| BP | 35023 | regulation of Rho protein signal transduction | 79 | -1,93 | 7,95E-04 |
| BP | 70167 | regulation of biomineral tissue development | 50 | -1,93 | 2,62E-03 |
| BP | 110149 | regulation of biomineralization | 50 | -1,93 | 2,62E-03 |
| BP | 61035 | regulation of cartilage development | 42 | -1,93 | 4,48E-03 |
| BP | 3014 | renal system process | 41 | -1,92 | 3,94E-03 |
| BP | 165 | MAPK cascade | 426 | -1,92 | 1,06E-08 |

| | | | | | |
|---|---|---|---|---|---|
| BP | 71260 | cellular response to mechanical stimulus | 24 | -1,92 | 7,53E-03 |
| BP | 70848 | response to growth factor | 392 | -1,92 | 8,38E-08 |
| BP | 19915 | lipid storage | 52 | -1,92 | 1,81E-03 |
| BP | 1902105 | regulation of leukocyte differentiation | 145 | -1,92 | 1,07E-04 |
| BP | 43547 | positive regulation of GTPase activity | 133 | -1,92 | 1,74E-04 |
| CC | 1726 | ruffle | 108 | -1,92 | 4,38E-04 |
| BP | 3158 | endothelium development | 78 | -1,92 | 8,92E-04 |
| BP | 98609 | cell-cell adhesion | 348 | -1,92 | 4,04E-07 |
| BP | 45778 | positive regulation of ossification | 30 | -1,92 | 5,88E-03 |
| MF | 15179 | L-amino acid transmembrane transporter activity | 29 | -1,92 | 6,82E-03 |
| BP | 42551 | neuron maturation | 25 | -1,92 | 7,68E-03 |
| MF | 38024 | cargo receptor activity | 31 | -1,92 | 6,35E-03 |
| CC | 30667 | secretory granule membrane | 42 | -1,91 | 4,67E-03 |
| CC | 30139 | endocytic vesicle | 138 | -1,91 | 2,00E-04 |
| BP | 8360 | regulation of cell shape | 99 | -1,91 | 4,65E-04 |
| BP | 30100 | regulation of endocytosis | 142 | -1,91 | 7,83E-05 |
| CC | 30175 | filopodium | 58 | -1,91 | 2,75E-03 |
| CC | 98857 | membrane microdomain | 224 | -1,91 | 1,11E-05 |
| BP | 6958 | complement activation, classical pathway | 10 | -1,91 | 8,46E-03 |
| CC | 323 | lytic vacuole | 315 | -1,91 | 5,05E-07 |
| CC | 5764 | lysosome | 315 | -1,91 | 5,05E-07 |
| BP | 2685 | regulation of leukocyte migration | 102 | -1,91 | 5,08E-04 |
| CC | 45121 | membrane raft | 223 | -1,91 | 1,45E-05 |
| BP | 2687 | positive regulation of leukocyte migration | 71 | -1,91 | 7,95E-04 |
| BP | 48246 | macrophage chemotaxis | 21 | -1,91 | 1,46E-02 |
| MF | 4715 | non-membrane spanning protein tyrosine kinase activity | 22 | -1,91 | 1,62E-02 |
| BP | 46339 | diacylglycerol metabolic process | 13 | -1,91 | 8,88E-03 |
| MF | 51015 | actin filament binding | 116 | -1,91 | 6,28E-04 |
| BP | 71363 | cellular response to growth factor stimulus | 383 | -1,91 | 5,98E-08 |
| BP | 10656 | negative regulation of muscle cell apoptotic process | 39 | -1,91 | 6,51E-03 |
| BP | 1953 | negative regulation of cell-matrix adhesion | 26 | -1,91 | 7,90E-03 |

| | | | | | |
|---|---|---|---|---|---|
| BP | 2001259 | positive regulation of cation channel activity | 34 | -1,91 | 7,87E-03 |
| BP | 6909 | phagocytosis | 111 | -1,91 | 2,17E-04 |
| BP | 10862 | positive regulation of pathway-restricted SMAD protein ph | 28 | -1,91 | 6,54E-03 |
| BP | 1934 | positive regulation of protein phosphorylation | 455 | -1,91 | 7,88E-09 |
| MF | 30020 | extracellular matrix structural constituent conferring tensile | 22 | -1,90 | 1,71E-02 |
| BP | 1974 | blood vessel remodeling | 33 | -1,90 | 7,78E-03 |
| MF | 42169 | SH2 domain binding | 26 | -1,90 | 7,96E-03 |
| BP | 10171 | body morphogenesis | 48 | -1,90 | 3,55E-03 |
| CC | 1725 | stress fiber | 64 | -1,90 | 2,08E-03 |
| CC | 97517 | contractile actin filament bundle | 64 | -1,90 | 2,08E-03 |
| MF | 19903 | protein phosphatase binding | 120 | -1,90 | 3,98E-04 |
| CC | 5769 | early endosome | 208 | -1,90 | 4,56E-05 |
| BP | 2281 | macrophage activation involved in immune response | 11 | -1,90 | 6,43E-03 |
| MF | 30695 | GTPase regulator activity | 257 | -1,90 | 3,43E-06 |
| BP | 6063 | uronic acid metabolic process | 11 | -1,90 | 6,43E-03 |
| BP | 19585 | glucuronate metabolic process | 11 | -1,90 | 6,43E-03 |
| CC | 31226 | intrinsic component of plasma membrane | 431 | -1,90 | 2,55E-08 |
| BP | 1818 | negative regulation of cytokine production | 132 | -1,90 | 1,12E-04 |
| BP | 46718 | viral entry into host cell | 45 | -1,90 | 4,81E-03 |
| MF | 5518 | collagen binding | 45 | -1,90 | 4,81E-03 |
| BP | 21879 | forebrain neuron differentiation | 26 | -1,90 | 8,01E-03 |
| BP | 31347 | regulation of defense response | 297 | -1,90 | 1,62E-06 |
| BP | 44409 | entry into host | 48 | -1,90 | 3,76E-03 |
| BP | 9791 | post-embryonic development | 82 | -1,90 | 8,38E-04 |
| BP | 6821 | chloride transport | 32 | -1,90 | 9,13E-03 |
| BP | 45807 | positive regulation of endocytosis | 79 | -1,90 | 1,07E-03 |
| BP | 3148 | outflow tract septum morphogenesis | 20 | -1,90 | 7,37E-03 |
| BP | 18108 | peptidyl-tyrosine phosphorylation | 160 | -1,90 | 4,83E-05 |
| CC | 70161 | anchoring junction | 347 | -1,90 | 3,46E-07 |
| BP | 3231 | cardiac ventricle development | 83 | -1,89 | 1,13E-03 |
| BP | 15698 | inorganic anion transport | 52 | -1,89 | 2,18E-03 |

| | | | | | |
|---|---|---|---|---|---|
| BP | 70555 | response to interleukin-1 | 64 | -1,89 | 2,39E-03 |
| BP | 2474 | antigen processing and presentation of peptide antigen via M | 18 | -1,89 | 7,96E-03 |
| MF | 8237 | metallopeptidase activity | 88 | -1,89 | 7,29E-04 |
| MF | 4714 | transmembrane receptor protein tyrosine kinase activity | 25 | -1,89 | 9,47E-03 |
| BP | 42063 | gliogenesis | 189 | -1,89 | 6,99E-05 |
| CC | 5774 | vacuolar membrane | 143 | -1,89 | 1,02E-04 |
| CC | 30141 | secretory granule | 156 | -1,89 | 1,10E-04 |
| BP | 1961 | positive regulation of cytokine-mediated signaling pathway | 32 | -1,89 | 1,03E-02 |
| BP | 46578 | regulation of Ras protein signal transduction | 150 | -1,89 | 9,96E-05 |
| BP | 6959 | humoral immune response | 59 | -1,89 | 2,56E-03 |
| BP | 21872 | forebrain generation of neurons | 30 | -1,89 | 7,58E-03 |
| BP | 30199 | collagen fibril organization | 41 | -1,89 | 5,78E-03 |
| BP | 2250 | adaptive immune response | 183 | -1,89 | 1,55E-04 |
| BP | 50865 | regulation of cell activation | 280 | -1,89 | 1,06E-05 |
| BP | 1905906 | regulation of amyloid fibril formation | 13 | -1,89 | 1,06E-02 |
| BP | 32608 | interferon-beta production | 40 | -1,88 | 6,37E-03 |
| BP | 9267 | cellular response to starvation | 107 | -1,88 | 3,91E-04 |
| BP | 1902043 | positive regulation of extrinsic apoptotic signaling pathway | 12 | -1,88 | 1,75E-02 |
| BP | 32330 | regulation of chondrocyte differentiation | 33 | -1,88 | 9,63E-03 |
| BP | 50866 | negative regulation of cell activation | 96 | -1,88 | 1,25E-03 |
| BP | 16485 | protein processing | 127 | -1,88 | 2,51E-04 |
| BP | 71559 | response to transforming growth factor beta | 151 | -1,88 | 1,87E-04 |
| BP | 8361 | regulation of cell size | 129 | -1,88 | 4,80E-04 |
| BP | 6897 | endocytosis | 313 | -1,88 | 7,23E-07 |
| BP | 7169 | transmembrane receptor protein tyrosine kinase signaling pa | 349 | -1,88 | 5,86E-07 |
| MF | 5125 | cytokine activity | 65 | -1,88 | 4,08E-03 |
| BP | 1904036 | negative regulation of epithelial cell apoptotic process | 32 | -1,87 | 1,20E-02 |
| BP | 50764 | regulation of phagocytosis | 47 | -1,87 | 4,63E-03 |
| BP | 48002 | antigen processing and presentation of peptide antigen | 26 | -1,87 | 1,06E-02 |
| BP | 55010 | ventricular cardiac muscle tissue morphogenesis | 27 | -1,87 | 1,21E-02 |
| BP | 1903706 | regulation of hemopoiesis | 199 | -1,87 | 5,28E-05 |

| | | | | | |
|---|---|---|---|---|---|
| BP | 7009 | plasma membrane organization | 69 | -1,87 | 2,93E-03 |
| BP | 45088 | regulation of innate immune response | 91 | -1,87 | 1,25E-03 |
| MF | 140030 | modification-dependent protein binding | 113 | -1,87 | 3,98E-04 |
| BP | 32648 | regulation of interferon-beta production | 38 | -1,87 | 6,29E-03 |
| BP | 30225 | macrophage differentiation | 23 | -1,87 | 9,78E-03 |
| CC | 30424 | axon | 376 | -1,87 | 4,81E-07 |
| BP | 32615 | interleukin-12 production | 24 | -1,87 | 1,37E-02 |
| BP | 8543 | fibroblast growth factor receptor signaling pathway | 40 | -1,86 | 7,78E-03 |
| BP | 97305 | response to alcohol | 75 | -1,86 | 1,64E-03 |
| BP | 2253 | activation of immune response | 119 | -1,86 | 4,52E-04 |
| MF | 15085 | calcium ion transmembrane transporter activity | 54 | -1,86 | 4,66E-03 |
| BP | 31643 | positive regulation of myelination | 11 | -1,86 | 1,15E-02 |
| BP | 7266 | Rho protein signal transduction | 106 | -1,86 | 7,33E-04 |
| MF | 38023 | signaling receptor activity | 247 | -1,86 | 1,85E-05 |
| MF | 60089 | molecular transducer activity | 247 | -1,86 | 1,85E-05 |
| BP | 1902041 | regulation of extrinsic apoptotic signaling pathway via deat | 34 | -1,86 | 1,12E-02 |
| BP | 3206 | cardiac chamber morphogenesis | 80 | -1,86 | 3,17E-03 |
| CC | 31674 | I band | 69 | -1,86 | 3,14E-03 |
| BP | 31669 | cellular response to nutrient levels | 127 | -1,86 | 3,39E-04 |
| CC | 31252 | cell leading edge | 256 | -1,86 | 1,53E-05 |
| BP | 6949 | syncytium formation | 36 | -1,86 | 9,46E-03 |
| BP | 14002 | astrocyte development | 20 | -1,85 | 1,07E-02 |
| BP | 2438 | acute inflammatory response to antigenic stimulus | 11 | -1,85 | 1,25E-02 |
| MF | 4888 | transmembrane signaling receptor activity | 178 | -1,85 | 1,81E-04 |
| BP | 36035 | osteoclast development | 11 | -1,85 | 1,25E-02 |
| CC | 98552 | side of membrane | 243 | -1,85 | 2,68E-05 |
| CC | 99503 | secretory vesicle | 272 | -1,85 | 6,50E-06 |
| BP | 32612 | interleukin-1 production | 51 | -1,85 | 5,88E-03 |
| MF | 31267 | small GTPase binding | 208 | -1,85 | 1,11E-04 |
| MF | 61135 | endopeptidase regulator activity | 71 | -1,85 | 1,69E-03 |
| BP | 2532 | production of molecular mediator involved in inflammatory | 41 | -1,85 | 7,86E-03 |

| | | | | | |
|---|---|---|---|---|---|
| BP | 18212 | peptidyl-tyrosine modification | 163 | -1,85 | 2,66E-04 |
| BP | 30099 | myeloid cell differentiation | 238 | -1,85 | 5,84E-05 |
| BP | 31341 | regulation of cell killing | 45 | -1,85 | 7,46E-03 |
| BP | 42116 | macrophage activation | 55 | -1,85 | 8,25E-03 |
| BP | 110110 | positive regulation of animal organ morphogenesis | 14 | -1,85 | 2,01E-02 |
| BP | 30168 | platelet activation | 40 | -1,85 | 9,08E-03 |
| BP | 43954 | cellular component maintenance | 41 | -1,85 | 7,99E-03 |
| BP | 30500 | regulation of bone mineralization | 42 | -1,85 | 7,79E-03 |
| BP | 43090 | amino acid import | 29 | -1,85 | 1,37E-02 |
| CC | 44754 | autolysosome | 10 | -1,84 | 1,86E-02 |
| BP | 34329 | cell junction assembly | 203 | -1,84 | 1,66E-04 |
| BP | 32655 | regulation of interleukin-12 production | 22 | -1,84 | 2,97E-02 |
| BP | 1903428 | positive regulation of reactive oxygen species biosynthetic | 35 | -1,84 | 1,17E-02 |
| BP | 32652 | regulation of interleukin-1 production | 46 | -1,84 | 9,89E-03 |
| BP | 31667 | response to nutrient levels | 185 | -1,84 | 2,82E-04 |
| MF | 70530 | K63-linked polyubiquitin modification-dependent protein b | 17 | -1,84 | 2,17E-02 |
| BP | 50885 | neuromuscular process controlling balance | 37 | -1,84 | 1,26E-02 |
| BP | 45637 | regulation of myeloid cell differentiation | 119 | -1,84 | 6,56E-04 |
| BP | 45824 | negative regulation of innate immune response | 32 | -1,84 | 1,71E-02 |
| BP | 42554 | superoxide anion generation | 19 | -1,84 | 2,17E-02 |
| BP | 50691 | regulation of defense response to virus by host | 24 | -1,84 | 1,76E-02 |
| BP | 2831 | regulation of response to biotic stimulus | 155 | -1,84 | 2,56E-04 |
| BP | 70169 | positive regulation of biomineral tissue development | 25 | -1,84 | 1,55E-02 |
| BP | 110151 | positive regulation of biomineralization | 25 | -1,84 | 1,55E-02 |
| CC | 30140 | trans-Golgi network transport vesicle | 22 | -1,84 | 3,15E-02 |
| BP | 30036 | actin cytoskeleton organization | 444 | -1,84 | 3,46E-07 |
| BP | 46460 | neutral lipid biosynthetic process | 26 | -1,84 | 1,47E-02 |
| BP | 46463 | acylglycerol biosynthetic process | 26 | -1,84 | 1,47E-02 |
| BP | 50807 | regulation of synapse organization | 125 | -1,84 | 7,42E-04 |
| BP | 62207 | regulation of pattern recognition receptor signaling pathway | 56 | -1,84 | 7,96E-03 |
| BP | 97284 | hepatocyte apoptotic process | 13 | -1,84 | 1,89E-02 |

| | | | | | |
|---|---|---|---|---|---|
| BP | 2695 | negative regulation of leukocyte activation | 85 | -1,84 | 4,45E-03 |
| BP | 8585 | female gonad development | 58 | -1,83 | 6,54E-03 |
| BP | 768 | syncytium formation by plasma membrane fusion | 34 | -1,83 | 1,41E-02 |
| BP | 140253 | cell-cell fusion | 34 | -1,83 | 1,41E-02 |
| BP | 21782 | glial cell development | 63 | -1,83 | 6,57E-03 |
| BP | 32480 | negative regulation of type I interferon production | 15 | -1,83 | 2,34E-02 |
| BP | 46545 | development of primary female sexual characteristics | 59 | -1,83 | 4,60E-03 |
| BP | 45639 | positive regulation of myeloid cell differentiation | 53 | -1,83 | 1,07E-02 |
| CC | 5773 | vacuole | 378 | -1,83 | 1,87E-06 |
| BP | 43491 | protein kinase B signaling | 119 | -1,83 | 7,44E-04 |
| BP | 72010 | glomerular epithelium development | 10 | -1,83 | 2,20E-02 |
| BP | 70374 | positive regulation of ERK1 and ERK2 cascade | 107 | -1,83 | 7,81E-04 |
| MF | 3779 | actin binding | 241 | -1,83 | 3,73E-05 |
| MF | 4175 | endopeptidase activity | 178 | -1,83 | 2,76E-04 |
| BP | 1904019 | epithelial cell apoptotic process | 77 | -1,83 | 2,55E-03 |
| BP | 50803 | regulation of synapse structure or activity | 126 | -1,83 | 5,14E-04 |
| MF | 42562 | hormone binding | 32 | -1,82 | 1,84E-02 |
| CC | 16363 | nuclear matrix | 70 | -1,82 | 6,80E-03 |
| BP | 70831 | basement membrane assembly | 10 | -1,82 | 2,35E-02 |
| BP | 2694 | regulation of leukocyte activation | 258 | -1,82 | 3,66E-05 |
| BP | 97191 | extrinsic apoptotic signaling pathway | 156 | -1,82 | 3,21E-04 |
| BP | 7162 | negative regulation of cell adhesion | 159 | -1,82 | 4,49E-04 |
| BP | 62208 | positive regulation of pattern recognition receptor signaling | 23 | -1,82 | 1,31E-02 |
| MF | 5096 | GTPase activator activity | 146 | -1,82 | 2,96E-04 |
| CC | 44297 | cell body | 422 | -1,82 | 7,85E-07 |
| BP | 9886 | post-embryonic animal morphogenesis | 10 | -1,82 | 2,41E-02 |
| BP | 1902903 | regulation of supramolecular fiber organization | 255 | -1,82 | 4,03E-05 |
| BP | 904 | cell morphogenesis involved in differentiation | 444 | -1,82 | 6,41E-07 |
| BP | 46827 | positive regulation of protein export from nucleus | 15 | -1,82 | 2,66E-02 |
| MF | 5217 | intracellular ligand-gated ion channel activity | 11 | -1,82 | 1,86E-02 |
| BP | 60389 | pathway-restricted SMAD protein phosphorylation | 42 | -1,82 | 1,01E-02 |

| | | | | | |
|---|---|---|---|---|---|
| CC | 48471 | perinuclear region of cytoplasm | 451 | -1,82 | 2,91E-07 |
| BP | 61635 | regulation of protein complex stability | 10 | -1,82 | 2,58E-02 |
| MF | 4197 | cysteine-type endopeptidase activity | 59 | -1,82 | 5,68E-03 |
| BP | 71560 | cellular response to transforming growth factor beta stimulu | 148 | -1,82 | 5,12E-04 |
| BP | 71496 | cellular response to external stimulus | 167 | -1,82 | 3,10E-04 |
| CC | 30018 | Z disc | 65 | -1,82 | 7,45E-03 |
| BP | 30278 | regulation of ossification | 72 | -1,82 | 4,84E-03 |
| BP | 19882 | antigen processing and presentation | 54 | -1,82 | 7,96E-03 |
| BP | 50776 | regulation of immune response | 302 | -1,81 | 3,06E-05 |
| MF | 30507 | spectrin binding | 15 | -1,81 | 2,95E-02 |
| MF | 15108 | chloride transmembrane transporter activity | 25 | -1,81 | 1,92E-02 |
| BP | 2675 | positive regulation of acute inflammatory response | 12 | -1,81 | 2,99E-02 |
| BP | 2088 | lens development in camera-type eye | 51 | -1,81 | 8,55E-03 |
| MF | 19902 | phosphatase binding | 150 | -1,81 | 3,59E-04 |
| CC | 43025 | neuronal cell body | 364 | -1,81 | 2,46E-06 |
| BP | 85029 | extracellular matrix assembly | 30 | -1,81 | 1,55E-02 |
| BP | 2000379 | positive regulation of reactive oxygen species metabolic pro | 67 | -1,81 | 6,27E-03 |
| BP | 60393 | regulation of pathway-restricted SMAD protein phosphoryl | 39 | -1,81 | 1,77E-02 |
| BP | 71526 | semaphorin-plexin signaling pathway | 21 | -1,81 | 2,97E-02 |
| BP | 8203 | cholesterol metabolic process | 77 | -1,81 | 3,10E-03 |
| BP | 1775 | cell activation | 486 | -1,81 | 1,35E-07 |
| BP | 1904645 | response to amyloid-beta | 22 | -1,81 | 3,68E-02 |
| BP | 35458 | cellular response to interferon-beta | 17 | -1,81 | 2,79E-02 |
| BP | 60760 | positive regulation of response to cytokine stimulus | 37 | -1,81 | 1,71E-02 |
| BP | 55094 | response to lipoprotein particle | 17 | -1,81 | 2,79E-02 |
| BP | 51480 | regulation of cytosolic calcium ion concentration | 143 | -1,81 | 4,60E-04 |
| MF | 46873 | metal ion transmembrane transporter activity | 130 | -1,81 | 6,28E-04 |
| BP | 50730 | regulation of peptidyl-tyrosine phosphorylation | 133 | -1,81 | 1,13E-03 |
| MF | 16298 | lipase activity | 45 | -1,81 | 1,07E-02 |
| BP | 1900120 | regulation of receptor binding | 12 | -1,81 | 3,13E-02 |
| BP | 10817 | regulation of hormone levels | 232 | -1,81 | 1,07E-04 |

| | | | | | |
|---|---|---|---|---|---|
| BP | 60411 | cardiac septum morphogenesis | 54 | -1,81 | 8,57E-03 |
| BP | 33674 | positive regulation of kinase activity | 296 | -1,81 | 3,57E-05 |
| BP | 51963 | regulation of synapse assembly | 43 | -1,81 | 1,56E-02 |
| MF | 22836 | gated channel activity | 75 | -1,80 | 3,46E-03 |
| BP | 72676 | lymphocyte migration | 50 | -1,80 | 7,32E-03 |
| BP | 10591 | regulation of lamellipodium assembly | 27 | -1,80 | 2,27E-02 |
| BP | 71402 | cellular response to lipoprotein particle stimulus | 18 | -1,80 | 1,89E-02 |
| BP | 51493 | regulation of cytoskeleton organization | 344 | -1,80 | 1,64E-05 |
| BP | 45471 | response to ethanol | 30 | -1,80 | 1,71E-02 |
| BP | 51767 | nitric-oxide synthase biosynthetic process | 18 | -1,80 | 1,89E-02 |
| BP | 51769 | regulation of nitric-oxide synthase biosynthetic process | 18 | -1,80 | 1,89E-02 |
| CC | 30136 | clathrin-coated vesicle | 73 | -1,80 | 4,45E-03 |
| BP | 32607 | interferon-alpha production | 23 | -1,80 | 1,61E-02 |
| BP | 30029 | actin filament-based process | 482 | -1,80 | 5,43E-07 |
| BP | 3007 | heart morphogenesis | 155 | -1,80 | 4,52E-04 |
| BP | 2521 | leukocyte differentiation | 280 | -1,80 | 6,46E-05 |
| MF | 5262 | calcium channel activity | 43 | -1,80 | 1,67E-02 |
| BP | 48260 | positive regulation of receptor-mediated endocytosis | 37 | -1,80 | 1,80E-02 |
| BP | 1902652 | secondary alcohol metabolic process | 86 | -1,80 | 4,77E-03 |
| BP | 2548 | monocyte chemotaxis | 26 | -1,79 | 2,21E-02 |
| BP | 2478 | antigen processing and presentation of exogenous peptide a | 16 | -1,79 | 2,28E-02 |
| BP | 10001 | glial cell differentiation | 129 | -1,79 | 2,08E-03 |
| BP | 1903036 | positive regulation of response to wounding | 44 | -1,79 | 1,93E-02 |
| BP | 2221 | pattern recognition receptor signaling pathway | 94 | -1,79 | 7,87E-03 |
| MF | 4843 | thiol-dependent ubiquitin-specific protease activity | 74 | -1,79 | 5,80E-03 |
| BP | 2064 | epithelial cell development | 135 | -1,79 | 1,31E-03 |
| CC | 30670 | phagocytic vesicle membrane | 17 | -1,79 | 3,22E-02 |
| BP | 32689 | negative regulation of interferon-gamma production | 14 | -1,79 | 3,53E-02 |
| BP | 7422 | peripheral nervous system development | 35 | -1,79 | 1,98E-02 |
| BP | 1946 | lymphangiogenesis | 13 | -1,79 | 3,12E-02 |
| BP | 1990868 | response to chemokine | 33 | -1,79 | 2,20E-02 |

| | | | | | |
|---|---|---|---|---|---|
| BP | 1990869 | cellular response to chemokine | 33 | -1,79 | 2,20E-02 |
| BP | 7249 | I-kappaB kinase/NF-kappaB signaling | 134 | -1,79 | 1,30E-03 |
| BP | 1901215 | negative regulation of neuron death | 150 | -1,79 | 5,07E-04 |
| BP | 45446 | endothelial cell differentiation | 70 | -1,79 | 1,00E-02 |
| BP | 2274 | myeloid leukocyte activation | 109 | -1,79 | 1,34E-03 |
| BP | 34097 | response to cytokine | 466 | -1,79 | 6,64E-07 |
| BP | 1903978 | regulation of microglial cell activation | 10 | -1,79 | 3,62E-02 |
| BP | 14911 | positive regulation of smooth muscle cell migration | 45 | -1,79 | 1,23E-02 |
| BP | 22604 | regulation of cell morphogenesis | 216 | -1,79 | 2,00E-04 |
| BP | 48661 | positive regulation of smooth muscle cell proliferation | 80 | -1,79 | 6,82E-03 |
| BP | 43393 | regulation of protein binding | 151 | -1,79 | 6,35E-04 |
| BP | 51216 | cartilage development | 118 | -1,79 | 1,62E-03 |
| BP | 22407 | regulation of cell-cell adhesion | 200 | -1,79 | 3,89E-04 |
| BP | 51960 | regulation of nervous system development | 275 | -1,78 | 4,03E-05 |
| BP | 14910 | regulation of smooth muscle cell migration | 64 | -1,78 | 8,55E-03 |
| MF | 101005 | ubiquitinyl hydrolase activity | 75 | -1,78 | 4,41E-03 |
| BP | 1902107 | positive regulation of leukocyte differentiation | 75 | -1,78 | 4,41E-03 |
| BP | 1903708 | positive regulation of hemopoiesis | 75 | -1,78 | 4,41E-03 |
| BP | 10172 | embryonic body morphogenesis | 13 | -1,78 | 3,20E-02 |
| BP | 90303 | positive regulation of wound healing | 32 | -1,78 | 2,35E-02 |
| BP | 61572 | actin filament bundle organization | 113 | -1,78 | 1,41E-03 |
| BP | 2931 | response to ischemia | 25 | -1,78 | 2,41E-02 |
| MF | 5126 | cytokine receptor binding | 130 | -1,78 | 8,92E-04 |
| BP | 1905517 | macrophage migration | 29 | -1,78 | 2,39E-02 |
| BP | 7179 | transforming growth factor beta receptor signaling pathway | 126 | -1,78 | 9,42E-04 |
| MF | 5547 | phosphatidylinositol-3,4,5-trisphosphate binding | 26 | -1,78 | 2,59E-02 |
| MF | 4252 | serine-type endopeptidase activity | 44 | -1,78 | 2,24E-02 |
| BP | 6814 | sodium ion transport | 85 | -1,78 | 8,33E-03 |
| BP | 7264 | small GTPase mediated signal transduction | 277 | -1,78 | 6,68E-05 |
| BP | 34121 | regulation of toll-like receptor signaling pathway | 38 | -1,78 | 1,45E-02 |
| BP | 1902003 | regulation of amyloid-beta formation | 22 | -1,78 | 4,33E-02 |

| | | | | | |
|---|---|---|---|---|---|
| BP | 32944 | regulation of mononuclear cell proliferation | 101 | -1,78 | 4,44E-03 |
| BP | 3281 | ventricular septum development | 56 | -1,77 | 1,34E-02 |
| BP | 21884 | forebrain neuron development | 18 | -1,77 | 2,28E-02 |
| BP | 120039 | plasma membrane bounded cell projection morphogenesis | 388 | -1,77 | 6,19E-06 |
| BP | 45780 | positive regulation of bone resorption | 17 | -1,77 | 3,35E-02 |
| BP | 42594 | response to starvation | 123 | -1,77 | 1,04E-03 |
| BP | 48812 | neuron projection morphogenesis | 377 | -1,77 | 7,53E-06 |
| BP | 72337 | modified amino acid transport | 15 | -1,77 | 3,64E-02 |
| BP | 30900 | forebrain development | 201 | -1,77 | 4,64E-04 |
| BP | 1901224 | positive regulation of NIK/NF-kappaB signaling | 45 | -1,77 | 1,38E-02 |
| BP | 2224 | toll-like receptor signaling pathway | 65 | -1,77 | 1,24E-02 |
| BP | 60759 | regulation of response to cytokine stimulus | 79 | -1,77 | 4,36E-03 |
| BP | 46850 | regulation of bone remodeling | 34 | -1,77 | 2,10E-02 |
| BP | 60603 | mammary gland duct morphogenesis | 25 | -1,77 | 2,56E-02 |
| BP | 6929 | substrate-dependent cell migration | 18 | -1,77 | 2,36E-02 |
| MF | 5216 | ion channel activity | 114 | -1,77 | 4,10E-03 |
| CC | 16342 | catenin complex | 12 | -1,77 | 4,41E-02 |
| BP | 48858 | cell projection morphogenesis | 392 | -1,77 | 9,58E-06 |
| BP | 60395 | SMAD protein signal transduction | 50 | -1,77 | 1,07E-02 |
| BP | 43255 | regulation of carbohydrate biosynthetic process | 70 | -1,77 | 1,17E-02 |
| BP | 46473 | phosphatidic acid metabolic process | 15 | -1,77 | 3,72E-02 |
| BP | 1901343 | negative regulation of vasculature development | 63 | -1,77 | 1,23E-02 |
| BP | 2000181 | negative regulation of blood vessel morphogenesis | 63 | -1,77 | 1,23E-02 |
| BP | 61028 | establishment of endothelial barrier | 33 | -1,77 | 2,72E-02 |
| BP | 90100 | positive regulation of transmembrane receptor protein serin | 78 | -1,77 | 6,02E-03 |
| BP | 10466 | negative regulation of peptidase activity | 103 | -1,77 | 4,39E-03 |
| BP | 2920 | regulation of humoral immune response | 12 | -1,77 | 4,46E-02 |
| BP | 2286 | T cell activation involved in immune response | 49 | -1,77 | 1,61E-02 |
| BP | 1901890 | positive regulation of cell junction assembly | 49 | -1,76 | 1,61E-02 |
| BP | 1901617 | organic hydroxy compound biosynthetic process | 121 | -1,76 | 1,72E-03 |
| BP | 46777 | protein autophosphorylation | 154 | -1,76 | 7,11E-04 |

| | | | | | |
|---|---|---|---|---|---|
| MF | 15175 | neutral amino acid transmembrane transporter activity | 17 | -1,76 | 3,59E-02 |
| BP | 50808 | synapse organization | 234 | -1,76 | 2,49E-04 |
| BP | 22408 | negative regulation of cell-cell adhesion | 100 | -1,76 | 4,12E-03 |
| CC | 98797 | plasma membrane protein complex | 207 | -1,76 | 4,65E-04 |
| BP | 34103 | regulation of tissue remodeling | 45 | -1,76 | 1,46E-02 |
| MF | 4222 | metalloendopeptidase activity | 49 | -1,76 | 1,62E-02 |
| BP | 51606 | detection of stimulus | 66 | -1,76 | 1,10E-02 |
| BP | 2833 | positive regulation of response to biotic stimulus | 74 | -1,76 | 8,01E-03 |
| BP | 42698 | ovulation cycle | 31 | -1,76 | 2,24E-02 |
| BP | 1822 | kidney development | 160 | -1,76 | 5,82E-04 |
| BP | 50777 | negative regulation of immune response | 74 | -1,76 | 8,01E-03 |
| BP | 43254 | regulation of protein-containing complex assembly | 300 | -1,76 | 4,00E-05 |
| BP | 7040 | lysosome organization | 60 | -1,76 | 1,46E-02 |
| BP | 80171 | lytic vacuole organization | 60 | -1,76 | 1,46E-02 |
| CC | 32587 | ruffle membrane | 45 | -1,76 | 1,46E-02 |
| BP | 90160 | Golgi to lysosome transport | 11 | -1,76 | 3,33E-02 |
| BP | 60443 | mammary gland morphogenesis | 35 | -1,76 | 2,51E-02 |
| BP | 44089 | positive regulation of cellular component biogenesis | 329 | -1,76 | 4,03E-05 |
| BP | 16339 | calcium-dependent cell-cell adhesion via plasma membrane | 13 | -1,76 | 3,81E-02 |
| BP | 1903409 | reactive oxygen species biosynthetic process | 75 | -1,76 | 6,47E-03 |
| CC | 5581 | collagen trimer | 40 | -1,76 | 1,71E-02 |
| BP | 14909 | smooth muscle cell migration | 71 | -1,76 | 5,68E-03 |
| MF | 72341 | modified amino acid binding | 53 | -1,76 | 1,96E-02 |
| BP | 50801 | ion homeostasis | 370 | -1,76 | 2,31E-05 |
| BP | 90066 | regulation of anatomical structure size | 320 | -1,76 | 3,66E-05 |
| BP | 14065 | phosphatidylinositol 3-kinase signaling | 67 | -1,76 | 1,11E-02 |
| BP | 150146 | cell junction disassembly | 13 | -1,76 | 3,86E-02 |
| MF | 15171 | amino acid transmembrane transporter activity | 37 | -1,75 | 2,34E-02 |
| BP | 50670 | regulation of lymphocyte proliferation | 99 | -1,75 | 3,99E-03 |
| BP | 6812 | cation transport | 482 | -1,75 | 3,41E-06 |
| BP | 98771 | inorganic ion homeostasis | 349 | -1,75 | 2,29E-05 |

| | | | | | |
|---|---|---|---|---|---|
| BP | 61912 | selective autophagy | 54 | -1,75 | 1,37E-02 |
| BP | 48608 | reproductive structure development | 244 | -1,75 | 3,65E-04 |
| BP | 32331 | negative regulation of chondrocyte differentiation | 16 | -1,75 | 3,31E-02 |
| BP | 1900015 | regulation of cytokine production involved in inflammatory | 23 | -1,75 | 2,28E-02 |
| BP | 72001 | renal system development | 171 | -1,75 | 8,09E-04 |
| BP | 10092 | specification of animal organ identity | 14 | -1,75 | 4,38E-02 |
| MF | 140375 | immune receptor activity | 32 | -1,75 | 2,78E-02 |
| BP | 7041 | lysosomal transport | 80 | -1,75 | 9,68E-03 |
| BP | 7409 | axonogenesis | 255 | -1,75 | 1,87E-04 |
| BP | 51017 | actin filament bundle assembly | 112 | -1,75 | 4,01E-03 |
| BP | 45321 | leukocyte activation | 428 | -1,75 | 7,04E-06 |
| BP | 2263 | cell activation involved in immune response | 134 | -1,75 | 2,15E-03 |
| MF | 61134 | peptidase regulator activity | 95 | -1,75 | 5,59E-03 |
| BP | 89718 | amino acid import across plasma membrane | 24 | -1,75 | 3,12E-02 |
| MF | 15267 | channel activity | 128 | -1,75 | 3,47E-03 |
| MF | 22803 | passive transmembrane transporter activity | 128 | -1,75 | 3,47E-03 |
| BP | 1655 | urogenital system development | 198 | -1,75 | 6,38E-04 |
| BP | 70098 | chemokine-mediated signaling pathway | 25 | -1,75 | 2,95E-02 |
| MF | 4553 | hydrolase activity, hydrolyzing O-glycosyl compounds | 51 | -1,75 | 1,83E-02 |
| BP | 7507 | heart development | 365 | -1,75 | 2,43E-05 |
| BP | 1902106 | negative regulation of leukocyte differentiation | 58 | -1,75 | 1,32E-02 |
| BP | 32989 | cellular component morphogenesis | 449 | -1,75 | 3,50E-06 |
| BP | 1903707 | negative regulation of hemopoiesis | 59 | -1,75 | 1,04E-02 |
| BP | 32956 | regulation of actin cytoskeleton organization | 233 | -1,75 | 3,66E-04 |
| BP | 3013 | circulatory system process | 245 | -1,74 | 3,16E-04 |
| BP | 55080 | cation homeostasis | 344 | -1,74 | 7,09E-05 |
| CC | 15629 | actin cytoskeleton | 304 | -1,74 | 1,06E-04 |
| BP | 32147 | activation of protein kinase activity | 136 | -1,74 | 1,82E-03 |
| BP | 30097 | hemopoiesis | 492 | -1,74 | 4,81E-06 |
| BP | 45124 | regulation of bone resorption | 29 | -1,74 | 3,09E-02 |
| BP | 6873 | cellular ion homeostasis | 313 | -1,74 | 2,71E-05 |

| | | | | | |
|---|---|---|---|---|---|
| BP | 32990 | cell part morphogenesis | 409 | -1,74 | 1,13E-05 |
| BP | 61448 | connective tissue development | 160 | -1,74 | 6,83E-04 |
| BP | 70663 | regulation of leukocyte proliferation | 110 | -1,74 | 4,44E-03 |
| BP | 42742 | defense response to bacterium | 77 | -1,74 | 6,29E-03 |
| BP | 55082 | cellular chemical homeostasis | 403 | -1,74 | 1,99E-05 |
| BP | 10660 | regulation of muscle cell apoptotic process | 66 | -1,74 | 1,28E-02 |
| BP | 10719 | negative regulation of epithelial to mesenchymal transition | 23 | -1,74 | 2,28E-02 |
| BP | 48535 | lymph node development | 15 | -1,74 | 4,74E-02 |
| BP | 2697 | regulation of immune effector process | 197 | -1,74 | 6,14E-04 |
| BP | 3170 | heart valve development | 35 | -1,74 | 2,66E-02 |
| BP | 35456 | response to interferon-beta | 23 | -1,74 | 2,28E-02 |
| BP | 22602 | ovulation cycle process | 23 | -1,74 | 2,28E-02 |
| BP | 31128 | developmental induction | 16 | -1,74 | 3,52E-02 |
| BP | 9991 | response to extracellular stimulus | 203 | -1,74 | 9,80E-04 |
| BP | 1894 | tissue homeostasis | 129 | -1,74 | 4,21E-03 |
| BP | 2437 | inflammatory response to antigenic stimulus | 25 | -1,74 | 3,03E-02 |
| BP | 31214 | biomineral tissue development | 85 | -1,74 | 1,07E-02 |
| BP | 110148 | biomineralization | 85 | -1,74 | 1,07E-02 |
| BP | 16125 | sterol metabolic process | 82 | -1,74 | 5,31E-03 |
| BP | 60562 | epithelial tube morphogenesis | 221 | -1,74 | 5,82E-04 |
| BP | 30224 | monocyte differentiation | 16 | -1,74 | 3,54E-02 |
| BP | 36230 | granulocyte activation | 17 | -1,74 | 4,78E-02 |
| BP | 1903829 | positive regulation of cellular protein localization | 198 | -1,74 | 7,10E-04 |
| BP | 10592 | positive regulation of lamellipodium assembly | 23 | -1,74 | 2,37E-02 |
| BP | 1904407 | positive regulation of nitric oxide metabolic process | 29 | -1,74 | 3,15E-02 |
| BP | 2001236 | regulation of extrinsic apoptotic signaling pathway | 107 | -1,74 | 2,92E-03 |
| CC | 8305 | integrin complex | 17 | -1,74 | 4,90E-02 |
| BP | 10951 | negative regulation of endopeptidase activity | 86 | -1,74 | 8,87E-03 |
| BP | 3229 | ventricular cardiac muscle tissue development | 33 | -1,74 | 3,47E-02 |
| BP | 60977 | coronary vasculature morphogenesis | 20 | -1,74 | 2,85E-02 |
| BP | 43112 | receptor metabolic process | 117 | -1,73 | 5,34E-03 |

| | | | | | |
|---|---|---|---|---|---|
| BP | 71222 | cellular response to lipopolysaccharide | 98 | -1,73 | 7,09E-03 |
| BP | 7252 | I-kappaB phosphorylation | 14 | -1,73 | 4,63E-02 |
| CC | 5791 | rough endoplasmic reticulum | 50 | -1,73 | 1,37E-02 |
| BP | 150076 | neuroinflammatory response | 19 | -1,73 | 4,44E-02 |
| BP | 48660 | regulation of smooth muscle cell proliferation | 113 | -1,73 | 2,74E-03 |
| BP | 30003 | cellular cation homeostasis | 308 | -1,73 | 1,07E-04 |
| MF | 8236 | serine-type peptidase activity | 56 | -1,73 | 1,99E-02 |
| BP | 10810 | regulation of cell-substrate adhesion | 147 | -1,73 | 8,93E-04 |
| CC | 31045 | dense core granule | 13 | -1,73 | 4,64E-02 |
| BP | 97062 | dendritic spine maintenance | 15 | -1,73 | 4,95E-02 |
| BP | 32535 | regulation of cellular component size | 249 | -1,73 | 3,48E-04 |
| BP | 6875 | cellular metal ion homeostasis | 276 | -1,73 | 1,48E-04 |
| BP | 7159 | leukocyte cell-cell adhesion | 152 | -1,73 | 1,69E-03 |
| CC | 43198 | dendritic shaft | 32 | -1,73 | 3,11E-02 |
| BP | 8015 | blood circulation | 224 | -1,73 | 3,74E-04 |
| BP | 8406 | gonad development | 107 | -1,73 | 3,16E-03 |
| BP | 45137 | development of primary sexual characteristics | 108 | -1,73 | 5,86E-03 |
| BP | 40013 | negative regulation of locomotion | 185 | -1,73 | 1,85E-03 |
| MF | 5085 | guanyl-nucleotide exchange factor activity | 117 | -1,73 | 6,02E-03 |
| BP | 7548 | sex differentiation | 131 | -1,73 | 2,14E-03 |
| BP | 16525 | negative regulation of angiogenesis | 62 | -1,73 | 1,57E-02 |
| CC | 12506 | vesicle membrane | 214 | -1,73 | 5,88E-04 |
| BP | 70527 | platelet aggregation | 24 | -1,73 | 3,59E-02 |
| MF | 4672 | protein kinase activity | 338 | -1,73 | 3,06E-05 |
| CC | 30666 | endocytic vesicle membrane | 28 | -1,73 | 3,49E-02 |
| BP | 61458 | reproductive system development | 247 | -1,73 | 3,94E-04 |
| BP | 7015 | actin filament organization | 277 | -1,73 | 2,17E-04 |
| BP | 2366 | leukocyte activation involved in immune response | 130 | -1,72 | 2,09E-03 |
| BP | 61037 | negative regulation of cartilage development | 20 | -1,72 | 3,12E-02 |
| BP | 2673 | regulation of acute inflammatory response | 19 | -1,72 | 4,79E-02 |
| BP | 1905330 | regulation of morphogenesis of an epithelium | 36 | -1,72 | 2,58E-02 |

| | | | | | |
|---|---|---|---|---|---|
| MF | 5154 | epidermal growth factor receptor binding | 20 | -1,72 | 3,15E-02 |
| BP | 61564 | axon development | 283 | -1,72 | 2,05E-04 |
| BP | 70168 | negative regulation of biomineral tissue development | 13 | -1,72 | 4,96E-02 |
| BP | 110150 | negative regulation of biomineralization | 13 | -1,72 | 4,96E-02 |
| BP | 32647 | regulation of interferon-alpha production | 19 | -1,72 | 4,90E-02 |
| BP | 71216 | cellular response to biotic stimulus | 123 | -1,72 | 2,42E-03 |
| BP | 71774 | response to fibroblast growth factor | 50 | -1,72 | 1,64E-02 |
| BP | 50778 | positive regulation of immune response | 219 | -1,72 | 6,80E-04 |
| BP | 55065 | metal ion homeostasis | 310 | -1,72 | 1,31E-04 |
| CC | 31901 | early endosome membrane | 44 | -1,72 | 3,08E-02 |
| BP | 48729 | tissue morphogenesis | 374 | -1,72 | 4,03E-05 |
| BP | 32720 | negative regulation of tumor necrosis factor production | 32 | -1,72 | 3,43E-02 |
| BP | 1903556 | negative regulation of tumor necrosis factor superfamily cy | 32 | -1,72 | 3,43E-02 |
| BP | 44344 | cellular response to fibroblast growth factor stimulus | 48 | -1,72 | 1,82E-02 |
| BP | 2285 | lymphocyte activation involved in immune response | 92 | -1,72 | 8,87E-03 |
| BP | 72503 | cellular divalent inorganic cation homeostasis | 223 | -1,72 | 5,08E-04 |
| BP | 98586 | cellular response to virus | 44 | -1,72 | 3,12E-02 |
| BP | 30032 | lamellipodium assembly | 47 | -1,72 | 1,72E-02 |
| BP | 9612 | response to mechanical stimulus | 76 | -1,72 | 8,54E-03 |
| BP | 48659 | smooth muscle cell proliferation | 116 | -1,72 | 8,12E-03 |
| CC | 70062 | extracellular exosome | 50 | -1,72 | 1,66E-02 |
| BP | 8625 | extrinsic apoptotic signaling pathway via death domain rece | 57 | -1,72 | 2,47E-02 |
| BP | 1902743 | regulation of lamellipodium organization | 33 | -1,72 | 3,69E-02 |
| BP | 10657 | muscle cell apoptotic process | 69 | -1,72 | 1,72E-02 |
| CC | 5911 | cell-cell junction | 254 | -1,72 | 6,28E-04 |
| BP | 31668 | cellular response to extracellular stimulus | 143 | -1,72 | 1,65E-03 |
| BP | 1901888 | regulation of cell junction assembly | 106 | -1,72 | 3,97E-03 |
| BP | 98739 | import across plasma membrane | 70 | -1,72 | 1,79E-02 |
| BP | 71219 | cellular response to molecule of bacterial origin | 103 | -1,72 | 8,01E-03 |
| BP | 14068 | positive regulation of phosphatidylinositol 3-kinase signalir | 31 | -1,71 | 3,04E-02 |
| BP | 43123 | positive regulation of I-kappaB kinase/NF-kappaB signalin | 74 | -1,71 | 1,16E-02 |

| | | | | | |
|---|---|---|---|---|---|
| BP | 1903522 | regulation of blood circulation | 107 | -1,71 | 3,70E-03 |
| BP | 71677 | positive regulation of mononuclear cell migration | 35 | -1,71 | 3,52E-02 |
| BP | 33619 | membrane protein proteolysis | 40 | -1,71 | 2,23E-02 |
| MF | 60589 | nucleoside-triphosphatase regulator activity | 295 | -1,71 | 1,95E-04 |
| BP | 6816 | calcium ion transport | 192 | -1,71 | 1,38E-03 |
| CC | 5912 | adherens junction | 79 | -1,71 | 7,79E-03 |
| BP | 35821 | modulation of process of other organism | 65 | -1,71 | 2,01E-02 |
| BP | 10959 | regulation of metal ion transport | 137 | -1,71 | 3,10E-03 |
| MF | 8233 | peptidase activity | 299 | -1,71 | 1,60E-04 |
| BP | 42445 | hormone metabolic process | 84 | -1,71 | 7,87E-03 |
| BP | 8277 | regulation of G protein-coupled receptor signaling pathway | 59 | -1,71 | 1,57E-02 |
| BP | 31343 | positive regulation of cell killing | 27 | -1,71 | 4,60E-02 |
| BP | 31664 | regulation of lipopolysaccharide-mediated signaling pathwa | 11 | -1,71 | 4,53E-02 |
| BP | 55008 | cardiac muscle tissue morphogenesis | 33 | -1,71 | 3,74E-02 |
| BP | 2009 | morphogenesis of an epithelium | 327 | -1,71 | 1,02E-04 |
| MF | 5261 | cation channel activity | 88 | -1,71 | 8,01E-03 |
| BP | 16242 | negative regulation of macroautophagy | 28 | -1,71 | 3,92E-02 |
| MF | 15276 | ligand-gated ion channel activity | 29 | -1,71 | 3,74E-02 |
| MF | 17171 | serine hydrolase activity | 57 | -1,71 | 2,63E-02 |
| BP | 51701 | biological process involved in interaction with host | 94 | -1,71 | 1,71E-02 |
| BP | 46661 | male sex differentiation | 73 | -1,71 | 1,04E-02 |
| BP | 42493 | response to drug | 108 | -1,71 | 7,58E-03 |
| BP | 6639 | acylglycerol metabolic process | 64 | -1,71 | 1,83E-02 |
| BP | 1902745 | positive regulation of lamellipodium organization | 26 | -1,71 | 4,07E-02 |
| BP | 80164 | regulation of nitric oxide metabolic process | 44 | -1,71 | 3,52E-02 |
| BP | 7265 | Ras protein signal transduction | 226 | -1,70 | 6,80E-04 |
| BP | 2000463 | positive regulation of excitatory postsynaptic potential | 16 | -1,70 | 4,53E-02 |
| BP | 48168 | regulation of neuronal synaptic plasticity | 30 | -1,70 | 2,99E-02 |
| BP | 2000241 | regulation of reproductive process | 83 | -1,70 | 9,66E-03 |
| MF | 51020 | GTPase binding | 239 | -1,70 | 8,75E-04 |
| BP | 6805 | xenobiotic metabolic process | 30 | -1,70 | 2,99E-02 |

| | | | | | |
|---|---|---|---|---|---|
| BP | 60444 | branching involved in mammary gland duct morphogenesis | 18 | -1,70 | 3,74E-02 |
| BP | 2000106 | regulation of leukocyte apoptotic process | 46 | -1,70 | 2,63E-02 |
| CC | 30659 | cytoplasmic vesicle membrane | 200 | -1,70 | 1,43E-03 |
| CC | 99699 | integral component of synaptic membrane | 64 | -1,70 | 1,92E-02 |
| CC | 45335 | phagocytic vesicle | 79 | -1,70 | 9,13E-03 |
| BP | 45732 | positive regulation of protein catabolic process | 178 | -1,70 | 2,29E-03 |
| BP | 51896 | regulation of protein kinase B signaling | 101 | -1,70 | 9,58E-03 |
| BP | 21953 | central nervous system neuron differentiation | 105 | -1,70 | 6,29E-03 |
| CC | 42383 | sarcolemma | 87 | -1,70 | 9,47E-03 |
| BP | 31663 | lipopolysaccharide-mediated signaling pathway | 31 | -1,70 | 3,30E-02 |
| MF | 42379 | chemokine receptor binding | 23 | -1,70 | 2,97E-02 |
| BP | 44403 | biological process involved in symbiotic interaction | 257 | -1,70 | 3,89E-04 |
| BP | 16579 | protein deubiquitination | 87 | -1,70 | 9,79E-03 |
| BP | 7173 | epidermal growth factor receptor signaling pathway | 63 | -1,70 | 2,30E-02 |
| BP | 60402 | calcium ion transport into cytosol | 77 | -1,70 | 1,02E-02 |
| BP | 30510 | regulation of BMP signaling pathway | 59 | -1,70 | 1,76E-02 |
| BP | 7176 | regulation of epidermal growth factor-activated receptor act | 18 | -1,70 | 4,01E-02 |
| BP | 7186 | G protein-coupled receptor signaling pathway | 225 | -1,70 | 1,82E-03 |
| BP | 7417 | central nervous system development | 489 | -1,70 | 5,85E-06 |
| CC | 99055 | integral component of postsynaptic membrane | 40 | -1,70 | 2,59E-02 |
| BP | 61097 | regulation of protein tyrosine kinase activity | 45 | -1,70 | 2,37E-02 |
| BP | 3151 | outflow tract morphogenesis | 53 | -1,70 | 2,92E-02 |
| BP | 51092 | positive regulation of NF-kappaB transcription factor activi | 88 | -1,69 | 9,45E-03 |
| BP | 21700 | developmental maturation | 158 | -1,69 | 1,99E-03 |
| BP | 32496 | response to lipopolysaccharide | 137 | -1,69 | 3,72E-03 |
| CC | 99056 | integral component of presynaptic membrane | 36 | -1,69 | 3,25E-02 |
| BP | 72006 | nephron development | 66 | -1,69 | 2,18E-02 |
| BP | 70372 | regulation of ERK1 and ERK2 cascade | 163 | -1,69 | 3,26E-03 |
| BP | 98751 | bone cell development | 23 | -1,69 | 3,11E-02 |
| MF | 99094 | ligand-gated cation channel activity | 26 | -1,69 | 4,31E-02 |
| BP | 71695 | anatomical structure maturation | 125 | -1,69 | 4,84E-03 |

| | | | | | |
|---|---|---|---|---|---|
| BP | 31638 | zymogen activation | 26 | -1,69 | 4,31E-02 |
| MF | 51019 | mitogen-activated protein kinase binding | 33 | -1,69 | 4,19E-02 |
| BP | 31344 | regulation of cell projection organization | 434 | -1,69 | 3,21E-05 |
| CC | 43230 | extracellular organelle | 56 | -1,69 | 2,75E-02 |
| BP | 48259 | regulation of receptor-mediated endocytosis | 73 | -1,69 | 1,08E-02 |
| BP | 1959 | regulation of cytokine-mediated signaling pathway | 70 | -1,69 | 2,20E-02 |
| BP | 45785 | positive regulation of cell adhesion | 229 | -1,69 | 7,46E-04 |
| BP | 1901655 | cellular response to ketone | 49 | -1,69 | 2,59E-02 |
| BP | 22612 | gland morphogenesis | 90 | -1,69 | 1,84E-02 |
| BP | 30336 | negative regulation of cell migration | 166 | -1,69 | 2,42E-03 |
| BP | 18958 | phenol-containing compound metabolic process | 44 | -1,69 | 3,78E-02 |
| BP | 48675 | axon extension | 89 | -1,69 | 1,17E-02 |
| CC | 44306 | neuron projection terminus | 81 | -1,69 | 1,27E-02 |
| BP | 3179 | heart valve morphogenesis | 28 | -1,69 | 4,63E-02 |
| BP | 72507 | divalent inorganic cation homeostasis | 234 | -1,69 | 1,22E-03 |
| CC | 5884 | actin filament | 84 | -1,69 | 1,09E-02 |
| MF | 16773 | phosphotransferase activity, alcohol group as acceptor | 403 | -1,68 | 9,10E-05 |
| BP | 70371 | ERK1 and ERK2 cascade | 174 | -1,68 | 2,58E-03 |
| BP | 48871 | multicellular organismal homeostasis | 251 | -1,68 | 8,52E-04 |
| BP | 48708 | astrocyte differentiation | 45 | -1,68 | 2,49E-02 |
| BP | 45599 | negative regulation of fat cell differentiation | 40 | -1,68 | 2,83E-02 |
| BP | 48010 | vascular endothelial growth factor receptor signaling pathw | 38 | -1,68 | 2,84E-02 |
| BP | 43405 | regulation of MAP kinase activity | 177 | -1,68 | 1,87E-03 |
| BP | 30001 | metal ion transport | 233 | -1,68 | 1,11E-03 |
| BP | 1569 | branching involved in blood vessel morphogenesis | 26 | -1,68 | 4,62E-02 |
| BP | 60390 | regulation of SMAD protein signal transduction | 18 | -1,68 | 4,37E-02 |
| BP | 120035 | regulation of plasma membrane bounded cell projection org | 425 | -1,68 | 6,68E-05 |
| BP | 6874 | cellular calcium ion homeostasis | 207 | -1,68 | 1,82E-03 |
| BP | 14037 | Schwann cell differentiation | 26 | -1,68 | 4,65E-02 |
| BP | 51962 | positive regulation of nervous system development | 180 | -1,68 | 2,57E-03 |
| BP | 51249 | regulation of lymphocyte activation | 211 | -1,68 | 2,54E-03 |

| | | | | | |
|---|---|---|---|---|---|
| BP | 32970 | regulation of actin filament-based process | 252 | -1,68 | 4,55E-04 |
| BP | 46580 | negative regulation of Ras protein signal transduction | 40 | -1,68 | 2,92E-02 |
| BP | 72593 | reactive oxygen species metabolic process | 179 | -1,68 | 3,74E-03 |
| BP | 6638 | neutral lipid metabolic process | 65 | -1,68 | 2,59E-02 |
| CC | 30425 | dendrite | 377 | -1,68 | 8,32E-05 |
| BP | 48525 | negative regulation of viral process | 53 | -1,68 | 3,50E-02 |
| CC | 31256 | leading edge membrane | 71 | -1,68 | 1,56E-02 |
| BP | 45428 | regulation of nitric oxide biosynthetic process | 42 | -1,68 | 3,22E-02 |
| BP | 7204 | positive regulation of cytosolic calcium ion concentration | 127 | -1,68 | 4,82E-03 |
| BP | 1763 | morphogenesis of a branching structure | 132 | -1,68 | 3,47E-03 |
| BP | 35725 | sodium ion transmembrane transport | 52 | -1,68 | 2,00E-02 |
| CC | 97447 | dendritic tree | 378 | -1,68 | 1,44E-04 |
| BP | 43122 | regulation of I-kappaB kinase/NF-kappaB signaling | 112 | -1,68 | 8,25E-03 |
| BP | 48469 | cell maturation | 87 | -1,68 | 1,29E-02 |
| BP | 30203 | glycosaminoglycan metabolic process | 50 | -1,68 | 2,18E-02 |
| BP | 48041 | focal adhesion assembly | 64 | -1,68 | 2,34E-02 |
| BP | 45927 | positive regulation of growth | 184 | -1,68 | 2,13E-03 |
| BP | 51250 | negative regulation of lymphocyte activation | 73 | -1,67 | 1,24E-02 |
| BP | 2762 | negative regulation of myeloid leukocyte differentiation | 29 | -1,67 | 4,70E-02 |
| BP | 51235 | maintenance of location | 203 | -1,67 | 2,42E-03 |
| BP | 45089 | positive regulation of innate immune response | 56 | -1,67 | 3,06E-02 |
| BP | 2534 | cytokine production involved in inflammatory response | 25 | -1,67 | 4,56E-02 |
| BP | 2000146 | negative regulation of cell motility | 172 | -1,67 | 4,17E-03 |
| BP | 61138 | morphogenesis of a branching epithelium | 125 | -1,67 | 6,29E-03 |
| BP | 1903900 | regulation of viral life cycle | 89 | -1,67 | 1,47E-02 |
| BP | 43500 | muscle adaptation | 76 | -1,67 | 1,35E-02 |
| BP | 6509 | membrane protein ectodomain proteolysis | 31 | -1,67 | 4,57E-02 |
| BP | 51347 | positive regulation of transferase activity | 358 | -1,67 | 1,41E-04 |
| BP | 6911 | phagocytosis, engulfment | 25 | -1,67 | 4,61E-02 |
| MF | 15318 | inorganic molecular entity transmembrane transporter activ | 288 | -1,67 | 6,14E-04 |
| BP | 30574 | collagen catabolic process | 23 | -1,67 | 3,81E-02 |

| | | | | | |
|---|---|---|---|---|---|
| BP | 51897 | positive regulation of protein kinase B signaling | 68 | -1,67 | 1,71E-02 |
| BP | 45165 | cell fate commitment | 107 | -1,67 | 6,57E-03 |
| BP | 50905 | neuromuscular process | 67 | -1,67 | 2,56E-02 |
| BP | 48732 | gland development | 257 | -1,67 | 7,01E-04 |
| MF | 1786 | phosphatidylserine binding | 30 | -1,67 | 3,82E-02 |
| BP | 71887 | leukocyte apoptotic process | 62 | -1,67 | 2,83E-02 |
| MF | 8324 | cation transmembrane transporter activity | 230 | -1,67 | 2,07E-03 |
| BP | 2000134 | negative regulation of G1/S transition of mitotic cell cycle | 44 | -1,66 | 4,64E-02 |
| BP | 10977 | negative regulation of neuron projection development | 93 | -1,66 | 1,42E-02 |
| BP | 48667 | cell morphogenesis involved in neuron differentiation | 349 | -1,66 | 1,84E-04 |
| CC | 5902 | microvillus | 54 | -1,66 | 3,08E-02 |
| BP | 30282 | bone mineralization | 67 | -1,66 | 2,58E-02 |
| BP | 45860 | positive regulation of protein kinase activity | 265 | -1,66 | 6,45E-04 |
| BP | 19725 | cellular homeostasis | 495 | -1,66 | 3,98E-05 |
| BP | 52126 | movement in host environment | 67 | -1,66 | 2,58E-02 |
| BP | 16032 | viral process | 222 | -1,66 | 1,98E-03 |
| BP | 3300 | cardiac muscle hypertrophy | 66 | -1,66 | 2,79E-02 |
| BP | 10595 | positive regulation of endothelial cell migration | 85 | -1,66 | 2,09E-02 |
| BP | 31532 | actin cytoskeleton reorganization | 62 | -1,66 | 2,94E-02 |
| BP | 6826 | iron ion transport | 30 | -1,66 | 4,17E-02 |
| MF | 15291 | secondary active transmembrane transporter activity | 81 | -1,66 | 1,59E-02 |
| BP | 150116 | regulation of cell-substrate junction organization | 53 | -1,66 | 4,17E-02 |
| CC | 30133 | transport vesicle | 209 | -1,66 | 3,55E-03 |
| BP | 1903038 | negative regulation of leukocyte cell-cell adhesion | 68 | -1,66 | 1,87E-02 |
| CC | 98802 | plasma membrane signaling receptor complex | 71 | -1,66 | 1,71E-02 |
| CC | 98889 | intrinsic component of presynaptic membrane | 40 | -1,66 | 3,28E-02 |
| BP | 32611 | interleukin-1 beta production | 40 | -1,66 | 3,28E-02 |
| BP | 43242 | negative regulation of protein-containing complex disassem | 59 | -1,66 | 2,53E-02 |
| BP | 60348 | bone development | 132 | -1,65 | 4,67E-03 |
| BP | 1901880 | negative regulation of protein depolymerization | 53 | -1,65 | 4,23E-02 |
| BP | 7626 | locomotory behavior | 116 | -1,65 | 1,62E-02 |

| | | | | | |
|---|---|---|---|---|---|
| BP | 2698 | negative regulation of immune effector process | 66 | -1,65 | 2,91E-02 |
| BP | 90101 | negative regulation of transmembrane receptor protein serir | 76 | -1,65 | 1,70E-02 |
| BP | 19058 | viral life cycle | 153 | -1,65 | 5,54E-03 |
| BP | 17015 | regulation of transforming growth factor beta receptor signa | 87 | -1,65 | 1,71E-02 |
| BP | 6809 | nitric oxide biosynthetic process | 49 | -1,65 | 3,23E-02 |
| BP | 55074 | calcium ion homeostasis | 213 | -1,65 | 2,87E-03 |
| BP | 46824 | positive regulation of nucleocytoplasmic transport | 54 | -1,65 | 3,24E-02 |
| BP | 71345 | cellular response to cytokine stimulus | 410 | -1,65 | 1,11E-04 |
| BP | 48638 | regulation of developmental growth | 236 | -1,65 | 2,09E-03 |
| BP | 32963 | collagen metabolic process | 72 | -1,65 | 2,48E-02 |
| BP | 2000377 | regulation of reactive oxygen species metabolic process | 133 | -1,65 | 8,89E-03 |
| BP | 60249 | anatomical structure homeostasis | 239 | -1,65 | 2,19E-03 |
| BP | 1903034 | regulation of response to wounding | 90 | -1,65 | 2,38E-02 |
| BP | 1667 | ameboidal-type cell migration | 285 | -1,65 | 6,14E-04 |
| BP | 2237 | response to molecule of bacterial origin | 145 | -1,64 | 7,57E-03 |
| BP | 6110 | regulation of glycolytic process | 30 | -1,64 | 4,68E-02 |
| BP | 50877 | nervous system process | 400 | -1,64 | 1,32E-04 |
| BP | 61041 | regulation of wound healing | 65 | -1,64 | 3,62E-02 |
| BP | 42391 | regulation of membrane potential | 173 | -1,64 | 3,17E-03 |
| BP | 16236 | macroautophagy | 175 | -1,64 | 6,29E-03 |
| BP | 1903426 | regulation of reactive oxygen species biosynthetic process | 64 | -1,64 | 2,79E-02 |
| BP | 46209 | nitric oxide metabolic process | 52 | -1,64 | 2,68E-02 |
| CC | 99240 | intrinsic component of synaptic membrane | 71 | -1,64 | 1,91E-02 |
| BP | 1903844 | regulation of cellular response to transforming growth facto | 88 | -1,64 | 1,41E-02 |
| BP | 42110 | T cell activation | 234 | -1,64 | 2,67E-03 |
| BP | 55076 | transition metal ion homeostasis | 79 | -1,64 | 1,67E-02 |
| BP | 2000401 | regulation of lymphocyte migration | 30 | -1,64 | 4,78E-02 |
| BP | 1902893 | regulation of pri-miRNA transcription by RNA polymerase | 40 | -1,64 | 3,59E-02 |
| BP | 30307 | positive regulation of cell growth | 127 | -1,64 | 7,32E-03 |
| BP | 71772 | response to BMP | 92 | -1,64 | 2,00E-02 |
| BP | 71773 | cellular response to BMP stimulus | 92 | -1,64 | 2,00E-02 |

| | | | | | |
|---|---|---|---|---|---|
| BP | 8584 | male gonad development | 60 | -1,64 | 3,75E-02 |
| BP | 43010 | camera-type eye development | 180 | -1,64 | 4,12E-03 |
| CC | 1903561 | extracellular vesicle | 54 | -1,64 | 3,49E-02 |
| BP | 70661 | leukocyte proliferation | 146 | -1,64 | 5,81E-03 |
| BP | 19079 | viral genome replication | 79 | -1,64 | 1,67E-02 |
| BP | 60322 | head development | 399 | -1,64 | 2,56E-04 |
| BP | 43502 | regulation of muscle adaptation | 65 | -1,64 | 3,77E-02 |
| BP | 98657 | import into cell | 108 | -1,64 | 1,42E-02 |
| BP | 48754 | branching morphogenesis of an epithelial tube | 102 | -1,64 | 1,42E-02 |
| BP | 1901222 | regulation of NIK/NF-kappaB signaling | 64 | -1,64 | 3,16E-02 |
| BP | 46546 | development of primary male sexual characteristics | 61 | -1,64 | 3,13E-02 |
| BP | 1904035 | regulation of epithelial cell apoptotic process | 60 | -1,63 | 3,75E-02 |
| BP | 50806 | positive regulation of synaptic transmission | 81 | -1,63 | 2,00E-02 |
| BP | 51271 | negative regulation of cellular component movement | 179 | -1,63 | 7,32E-03 |
| CC | 97060 | synaptic membrane | 184 | -1,63 | 4,38E-03 |
| MF | 44325 | ion channel binding | 82 | -1,63 | 1,70E-02 |
| CC | 70382 | exocytic vesicle | 147 | -1,63 | 4,48E-03 |
| BP | 38127 | ERBB signaling pathway | 69 | -1,63 | 3,35E-02 |
| MF | 5215 | transporter activity | 443 | -1,63 | 1,92E-04 |
| BP | 1902904 | negative regulation of supramolecular fiber organization | 112 | -1,63 | 1,36E-02 |
| MF | 22890 | inorganic cation transmembrane transporter activity | 206 | -1,63 | 4,36E-03 |
| BP | 10876 | lipid localization | 244 | -1,63 | 3,28E-03 |
| BP | 45216 | cell-cell junction organization | 100 | -1,63 | 1,72E-02 |
| BP | 43588 | skin development | 136 | -1,63 | 8,76E-03 |
| BP | 8016 | regulation of heart contraction | 79 | -1,63 | 1,83E-02 |
| BP | 45596 | negative regulation of cell differentiation | 382 | -1,63 | 2,00E-04 |
| BP | 60284 | regulation of cell development | 315 | -1,63 | 7,33E-04 |
| BP | 14070 | response to organic cyclic compound | 371 | -1,63 | 6,54E-04 |
| BP | 43524 | negative regulation of neuron apoptotic process | 104 | -1,62 | 2,05E-02 |
| BP | 14812 | muscle cell migration | 80 | -1,62 | 3,07E-02 |
| BP | 46165 | alcohol biosynthetic process | 86 | -1,62 | 2,54E-02 |

| | | | | | |
|---|---|---|---|---|---|
| BP | 38061 | NIK/NF-kappaB signaling | 76 | -1,62 | 2,20E-02 |
| BP | 34446 | substrate adhesion-dependent cell spreading | 73 | -1,62 | 1,80E-02 |
| BP | 30509 | BMP signaling pathway | 91 | -1,62 | 2,20E-02 |
| BP | 32874 | positive regulation of stress-activated MAPK cascade | 68 | -1,62 | 2,59E-02 |
| BP | 14066 | regulation of phosphatidylinositol 3-kinase signaling | 48 | -1,62 | 3,54E-02 |
| MF | 8242 | omega peptidase activity | 78 | -1,62 | 2,34E-02 |
| BP | 1903131 | mononuclear cell differentiation | 207 | -1,62 | 4,13E-03 |
| BP | 1906 | cell killing | 73 | -1,62 | 1,80E-02 |
| BP | 6029 | proteoglycan metabolic process | 48 | -1,62 | 3,57E-02 |
| BP | 60541 | respiratory system development | 156 | -1,62 | 7,78E-03 |
| BP | 1990778 | protein localization to cell periphery | 245 | -1,62 | 2,98E-03 |
| BP | 32635 | interleukin-6 production | 88 | -1,62 | 1,81E-02 |
| BP | 51924 | regulation of calcium ion transport | 132 | -1,62 | 7,32E-03 |
| BP | 32675 | regulation of interleukin-6 production | 82 | -1,62 | 1,84E-02 |
| BP | 51058 | negative regulation of small GTPase mediated signal transd | 46 | -1,62 | 4,29E-02 |
| BP | 31122 | cytoplasmic microtubule organization | 46 | -1,62 | 4,29E-02 |
| MF | 19955 | cytokine binding | 61 | -1,61 | 3,52E-02 |
| BP | 14896 | muscle hypertrophy | 69 | -1,61 | 3,64E-02 |
| BP | 48593 | camera-type eye morphogenesis | 73 | -1,61 | 1,89E-02 |
| BP | 43549 | regulation of kinase activity | 477 | -1,61 | 1,74E-04 |
| BP | 46649 | lymphocyte activation | 350 | -1,61 | 7,95E-04 |
| BP | 1903037 | regulation of leukocyte cell-cell adhesion | 137 | -1,61 | 1,00E-02 |
| BP | 2001057 | reactive nitrogen species metabolic process | 55 | -1,61 | 4,94E-02 |
| BP | 2292 | T cell differentiation involved in immune response | 38 | -1,61 | 4,05E-02 |
| BP | 98742 | cell-cell adhesion via plasma-membrane adhesion molecule | 69 | -1,61 | 3,74E-02 |
| BP | 7610 | behavior | 318 | -1,61 | 5,38E-04 |
| BP | 7600 | sensory perception | 169 | -1,61 | 5,31E-03 |
| BP | 3018 | vascular process in circulatory system | 109 | -1,61 | 1,37E-02 |
| BP | 45580 | regulation of T cell differentiation | 70 | -1,61 | 3,61E-02 |
| BP | 7416 | synapse assembly | 71 | -1,61 | 2,59E-02 |
| BP | 8544 | epidermis development | 156 | -1,61 | 8,31E-03 |

| | | | | | |
|---|---|---|---|---|---|
| BP | 7033 | vacuole organization | 149 | -1,61 | 1,14E-02 |
| BP | 70304 | positive regulation of stress-activated protein kinase signali | 70 | -1,60 | 3,66E-02 |
| BP | 35150 | regulation of tube size | 77 | -1,60 | 2,50E-02 |
| BP | 35296 | regulation of tube diameter | 77 | -1,60 | 2,50E-02 |
| BP | 97746 | blood vessel diameter maintenance | 77 | -1,60 | 2,50E-02 |
| BP | 50863 | regulation of T cell activation | 144 | -1,60 | 1,31E-02 |
| MF | 5516 | calmodulin binding | 93 | -1,60 | 2,35E-02 |
| BP | 9914 | hormone transport | 159 | -1,60 | 8,22E-03 |
| BP | 32943 | mononuclear cell proliferation | 132 | -1,60 | 8,85E-03 |
| BP | 1901214 | regulation of neuron death | 232 | -1,60 | 4,45E-03 |
| BP | 48588 | developmental cell growth | 165 | -1,60 | 1,11E-02 |
| CC | 150034 | distal axon | 204 | -1,60 | 5,73E-03 |
| BP | 42982 | amyloid precursor protein metabolic process | 51 | -1,60 | 4,63E-02 |
| BP | 2460 | adaptive immune response based on somatic recombination | 142 | -1,60 | 9,47E-03 |
| CC | 43292 | contractile fiber | 117 | -1,60 | 2,59E-02 |
| BP | 16441 | posttranscriptional gene silencing | 68 | -1,60 | 3,52E-02 |
| CC | 99081 | supramolecular polymer | 493 | -1,60 | 8,64E-05 |
| BP | 7420 | brain development | 361 | -1,60 | 4,31E-04 |
| BP | 7219 | Notch signaling pathway | 96 | -1,60 | 2,34E-02 |
| CC | 30016 | myofibril | 107 | -1,59 | 1,72E-02 |
| BP | 31346 | positive regulation of cell projection organization | 263 | -1,59 | 3,13E-03 |
| CC | 30017 | sarcomere | 93 | -1,59 | 2,51E-02 |
| BP | 32092 | positive regulation of protein binding | 75 | -1,59 | 3,01E-02 |
| BP | 30879 | mammary gland development | 85 | -1,59 | 3,35E-02 |
| BP | 14897 | striated muscle hypertrophy | 68 | -1,59 | 3,52E-02 |
| BP | 72659 | protein localization to plasma membrane | 210 | -1,59 | 5,41E-03 |
| BP | 10522 | regulation of calcium ion transport into cytosol | 58 | -1,59 | 3,87E-02 |
| BP | 187 | activation of MAPK activity | 60 | -1,59 | 4,78E-02 |
| BP | 1903364 | positive regulation of cellular protein catabolic process | 122 | -1,59 | 1,67E-02 |
| BP | 110053 | regulation of actin filament organization | 183 | -1,59 | 1,70E-02 |
| CC | 34399 | nuclear periphery | 91 | -1,58 | 3,10E-02 |

| | | | | | |
|---|---|---|---|---|---|
| BP | 10506 | regulation of autophagy | 189 | -1,58 | 7,36E-03 |
| MF | 8234 | cysteine-type peptidase activity | 105 | -1,58 | 2,05E-02 |
| BP | 33993 | response to lipid | 374 | -1,58 | 1,04E-03 |
| BP | 6914 | autophagy | 325 | -1,58 | 1,73E-03 |
| BP | 61919 | process utilizing autophagic mechanism | 325 | -1,58 | 1,73E-03 |
| BP | 10975 | regulation of neuron projection development | 316 | -1,58 | 1,81E-03 |
| BP | 30217 | T cell differentiation | 132 | -1,58 | 1,02E-02 |
| BP | 140352 | export from cell | 470 | -1,58 | 4,67E-04 |
| BP | 97581 | lamellipodium organization | 61 | -1,58 | 4,38E-02 |
| MF | 5543 | phospholipid binding | 262 | -1,58 | 3,55E-03 |
| CC | 10008 | endosome membrane | 180 | -1,58 | 8,93E-03 |
| BP | 46651 | lymphocyte proliferation | 130 | -1,58 | 1,30E-02 |
| BP | 21537 | telencephalon development | 128 | -1,58 | 1,90E-02 |
| BP | 51098 | regulation of binding | 270 | -1,58 | 2,90E-03 |
| BP | 46879 | hormone secretion | 158 | -1,57 | 1,09E-02 |
| BP | 42129 | regulation of T cell proliferation | 77 | -1,57 | 2,96E-02 |
| BP | 10720 | positive regulation of cell development | 206 | -1,57 | 8,89E-03 |
| BP | 48167 | regulation of synaptic plasticity | 101 | -1,57 | 2,92E-02 |
| CC | 99512 | supramolecular fiber | 490 | -1,57 | 3,32E-04 |
| BP | 60401 | cytosolic calcium ion transport | 95 | -1,57 | 3,12E-02 |
| BP | 3012 | muscle system process | 200 | -1,57 | 9,34E-03 |
| BP | 48589 | developmental growth | 443 | -1,57 | 6,80E-04 |
| BP | 33135 | regulation of peptidyl-serine phosphorylation | 86 | -1,57 | 3,53E-02 |
| BP | 1901615 | organic hydroxy compound metabolic process | 261 | -1,57 | 5,07E-03 |
| BP | 61387 | regulation of extent of cell growth | 78 | -1,57 | 3,70E-02 |
| CC | 19898 | extrinsic component of membrane | 189 | -1,57 | 8,51E-03 |
| MF | 8083 | growth factor activity | 61 | -1,57 | 4,77E-02 |
| BP | 10942 | positive regulation of cell death | 432 | -1,57 | 9,19E-04 |
| BP | 32940 | secretion by cell | 456 | -1,57 | 3,16E-04 |
| CC | 98793 | presynapse | 286 | -1,56 | 4,38E-03 |
| MF | 8238 | exopeptidase activity | 50 | -1,56 | 4,55E-02 |

| | | | | | |
|---|---|---|---|---|---|
| BP | 30516 | regulation of axon extension | 71 | -1,56 | 3,85E-02 |
| BP | 1501 | skeletal system development | 300 | -1,56 | 3,10E-03 |
| BP | 45861 | negative regulation of proteolysis | 174 | -1,56 | 1,27E-02 |
| MF | 15075 | ion transmembrane transporter activity | 350 | -1,56 | 1,83E-03 |
| MF | 70851 | growth factor receptor binding | 77 | -1,56 | 3,22E-02 |
| MF | 22857 | transmembrane transporter activity | 382 | -1,55 | 8,92E-04 |
| BP | 50867 | positive regulation of cell activation | 162 | -1,55 | 1,98E-02 |
| CC | 5938 | cell cortex | 180 | -1,55 | 1,37E-02 |
| CC | 30027 | lamellipodium | 129 | -1,55 | 2,38E-02 |
| BP | 48880 | sensory system development | 211 | -1,55 | 1,61E-02 |
| BP | 150063 | visual system development | 211 | -1,55 | 1,61E-02 |
| BP | 48872 | homeostasis of number of cells | 220 | -1,55 | 1,23E-02 |
| MF | 8289 | lipid binding | 408 | -1,55 | 7,95E-04 |
| BP | 9896 | positive regulation of catabolic process | 343 | -1,55 | 3,21E-03 |
| BP | 31333 | negative regulation of protein-containing complex assembly | 100 | -1,55 | 3,50E-02 |
| BP | 10634 | positive regulation of epithelial cell migration | 116 | -1,55 | 3,74E-02 |
| CC | 8021 | synaptic vesicle | 138 | -1,54 | 2,37E-02 |
| BP | 50770 | regulation of axonogenesis | 104 | -1,54 | 3,90E-02 |
| BP | 10594 | regulation of endothelial cell migration | 124 | -1,54 | 3,89E-02 |
| BP | 34762 | regulation of transmembrane transport | 262 | -1,54 | 7,78E-03 |
| BP | 30855 | epithelial cell differentiation | 309 | -1,54 | 4,23E-03 |
| BP | 50767 | regulation of neurogenesis | 237 | -1,54 | 1,23E-02 |
| BP | 9913 | epidermal cell differentiation | 92 | -1,54 | 4,57E-02 |
| BP | 1936 | regulation of endothelial cell proliferation | 84 | -1,54 | 3,48E-02 |
| BP | 51345 | positive regulation of hydrolase activity | 318 | -1,54 | 2,12E-03 |
| BP | 6887 | exocytosis | 199 | -1,54 | 1,20E-02 |
| BP | 60627 | regulation of vesicle-mediated transport | 333 | -1,54 | 3,10E-03 |
| BP | 45598 | regulation of fat cell differentiation | 93 | -1,54 | 3,62E-02 |
| BP | 1990138 | neuron projection extension | 120 | -1,54 | 3,89E-02 |
| BP | 16331 | morphogenesis of embryonic epithelium | 116 | -1,54 | 4,05E-02 |
| BP | 9100 | glycoprotein metabolic process | 209 | -1,54 | 1,89E-02 |

| | | | | | |
|---|---|---|---|---|---|
| BP | 1654 | eye development | 210 | -1,53 | 9,92E-03 |
| BP | 33002 | muscle cell proliferation | 158 | -1,53 | 1,89E-02 |
| MF | 42277 | peptide binding | 157 | -1,53 | 2,50E-02 |
| BP | 50804 | modulation of chemical synaptic transmission | 229 | -1,53 | 9,21E-03 |
| BP | 43903 | regulation of biological process involved in symbiotic inter: | 112 | -1,53 | 3,31E-02 |
| BP | 45859 | regulation of protein kinase activity | 427 | -1,53 | 2,45E-03 |
| BP | 8202 | steroid metabolic process | 153 | -1,53 | 2,37E-02 |
| BP | 43406 | positive regulation of MAP kinase activity | 125 | -1,53 | 2,89E-02 |
| BP | 31331 | positive regulation of cellular catabolic process | 295 | -1,53 | 6,55E-03 |
| BP | 34765 | regulation of ion transmembrane transport | 258 | -1,52 | 8,51E-03 |
| MF | 5319 | lipid transporter activity | 71 | -1,52 | 4,78E-02 |
| BP | 99177 | regulation of trans-synaptic signaling | 230 | -1,52 | 1,37E-02 |
| BP | 43068 | positive regulation of programmed cell death | 394 | -1,52 | 1,90E-03 |
| BP | 1935 | endothelial cell proliferation | 95 | -1,52 | 4,90E-02 |
| BP | 1903827 | regulation of cellular protein localization | 407 | -1,52 | 1,98E-03 |
| BP | 7254 | JNK cascade | 101 | -1,52 | 4,82E-02 |
| BP | 2768 | immune response-regulating cell surface receptor signaling | 84 | -1,52 | 3,90E-02 |
| BP | 1990845 | adaptive thermogenesis | 83 | -1,52 | 4,90E-02 |
| BP | 30324 | lung development | 143 | -1,52 | 2,03E-02 |
| BP | 3015 | heart process | 110 | -1,52 | 3,83E-02 |
| BP | 10721 | negative regulation of cell development | 105 | -1,52 | 3,31E-02 |
| BP | 50792 | regulation of viral process | 105 | -1,51 | 3,35E-02 |
| BP | 43542 | endothelial cell migration | 157 | -1,51 | 2,87E-02 |
| BP | 46328 | regulation of JNK cascade | 77 | -1,51 | 4,11E-02 |
| BP | 43270 | positive regulation of ion transport | 393 | -1,51 | 2,30E-03 |
| BP | 51403 | stress-activated MAPK cascade | 147 | -1,51 | 2,20E-02 |
| BP | 30323 | respiratory tube development | 146 | -1,51 | 2,55E-02 |
| BP | 1901699 | cellular response to nitrogen compound | 357 | -1,51 | 2,98E-03 |
| BP | 70646 | protein modification by small protein removal | 102 | -1,51 | 4,60E-02 |
| CC | 30426 | growth cone | 143 | -1,51 | 2,21E-02 |
| BP | 19932 | second-messenger-mediated signaling | 132 | -1,50 | 2,29E-02 |

| | | | | | |
|---|---|---|---|---|---|
| BP | 30098 | lymphocyte differentiation | 185 | -1,50 | 3,52E-02 |
| BP | 51604 | protein maturation | 172 | -1,50 | 2,96E-02 |
| BP | 51099 | positive regulation of binding | 143 | -1,50 | 2,28E-02 |
| BP | 7268 | chemical synaptic transmission | 295 | -1,50 | 9,36E-03 |
| BP | 98916 | anterograde trans-synaptic signaling | 295 | -1,50 | 9,36E-03 |
| BP | 99537 | trans-synaptic signaling | 300 | -1,50 | 8,01E-03 |
| BP | 71417 | cellular response to organonitrogen compound | 321 | -1,49 | 5,74E-03 |
| BP | 43523 | regulation of neuron apoptotic process | 165 | -1,49 | 3,64E-02 |
| BP | 2696 | positive regulation of leukocyte activation | 154 | -1,49 | 2,30E-02 |
| BP | 6109 | regulation of carbohydrate metabolic process | 121 | -1,49 | 3,49E-02 |
| BP | 98655 | cation transmembrane transport | 310 | -1,48 | 1,18E-02 |
| BP | 40008 | regulation of growth | 445 | -1,48 | 3,87E-03 |
| BP | 10631 | epithelial cell migration | 212 | -1,48 | 3,21E-02 |
| BP | 90132 | epithelium migration | 212 | -1,48 | 3,21E-02 |
| BP | 8285 | negative regulation of cell population proliferation | 409 | -1,48 | 5,83E-03 |
| BP | 48639 | positive regulation of developmental growth | 126 | -1,48 | 2,95E-02 |
| BP | 52547 | regulation of peptidase activity | 215 | -1,48 | 2,88E-02 |
| BP | 31345 | negative regulation of cell projection organization | 119 | -1,48 | 4,05E-02 |
| BP | 90130 | tissue migration | 213 | -1,48 | 3,04E-02 |
| CC | 30427 | site of polarized growth | 146 | -1,47 | 3,45E-02 |
| BP | 70507 | regulation of microtubule cytoskeleton organization | 105 | -1,47 | 4,81E-02 |
| BP | 60560 | developmental growth involved in morphogenesis | 162 | -1,47 | 4,17E-02 |
| BP | 43086 | negative regulation of catalytic activity | 401 | -1,47 | 4,12E-03 |
| BP | 50769 | positive regulation of neurogenesis | 163 | -1,47 | 4,37E-02 |
| MF | 33218 | amide binding | 206 | -1,47 | 2,79E-02 |
| BP | 10632 | regulation of epithelial cell migration | 168 | -1,47 | 4,53E-02 |
| MF | 4674 | protein serine/threonine kinase activity | 271 | -1,47 | 1,92E-02 |
| BP | 51346 | negative regulation of hydrolase activity | 195 | -1,46 | 3,28E-02 |
| MF | 44389 | ubiquitin-like protein ligase binding | 259 | -1,46 | 2,02E-02 |
| BP | 1904951 | positive regulation of establishment of protein localization | 215 | -1,46 | 3,22E-02 |
| MF | 16301 | kinase activity | 445 | -1,46 | 5,68E-03 |

| | | | | | |
|---|---|---|---|---|---|
| BP | 16049 | cell growth | 334 | -1,46 | 7,80E-03 |
| BP | 98660 | inorganic ion transmembrane transport | 292 | -1,46 | 1,68E-02 |
| CC | 5770 | late endosome | 162 | -1,46 | 4,64E-02 |
| BP | 71396 | cellular response to lipid | 272 | -1,46 | 1,45E-02 |
| BP | 71214 | cellular response to abiotic stimulus | 177 | -1,46 | 3,22E-02 |
| BP | 104004 | cellular response to environmental stimulus | 177 | -1,46 | 3,22E-02 |
| BP | 30162 | regulation of proteolysis | 430 | -1,46 | 6,86E-03 |
| BP | 9725 | response to hormone | 390 | -1,45 | 7,80E-03 |
| BP | 1901652 | response to peptide | 244 | -1,45 | 3,38E-02 |
| BP | 35265 | organ growth | 127 | -1,45 | 3,97E-02 |
| BP | 32271 | regulation of protein polymerization | 147 | -1,45 | 4,07E-02 |
| MF | 3700 | DNA-binding transcription factor activity | 495 | -1,45 | 3,99E-03 |
| BP | 6869 | lipid transport | 210 | -1,44 | 2,97E-02 |
| BP | 22411 | cellular component disassembly | 278 | -1,44 | 1,78E-02 |
| BP | 51222 | positive regulation of protein transport | 202 | -1,44 | 3,52E-02 |
| BP | 70997 | neuron death | 256 | -1,44 | 2,17E-02 |
| BP | 51051 | negative regulation of transport | 266 | -1,44 | 2,00E-02 |
| BP | 1903793 | positive regulation of anion transport | 302 | -1,44 | 2,10E-02 |
| BP | 3006 | developmental process involved in reproduction | 467 | -1,44 | 6,43E-03 |
| BP | 23061 | signal release | 255 | -1,44 | 2,72E-02 |
| BP | 99536 | synaptic signaling | 313 | -1,43 | 1,64E-02 |
| MF | 31625 | ubiquitin protein ligase binding | 246 | -1,43 | 2,44E-02 |
| BP | 10035 | response to inorganic substance | 262 | -1,43 | 3,19E-02 |
| BP | 43065 | positive regulation of apoptotic process | 387 | -1,43 | 1,54E-02 |
| BP | 42176 | regulation of protein catabolic process | 305 | -1,42 | 2,37E-02 |
| BP | 42692 | muscle cell differentiation | 239 | -1,42 | 3,11E-02 |
| MF | 16887 | ATPase activity | 254 | -1,42 | 3,15E-02 |
| BP | 6066 | alcohol metabolic process | 192 | -1,42 | 4,63E-02 |
| MF | 981 | DNA-binding transcription factor activity, RNA polymeras | 452 | -1,42 | 1,12E-02 |
| BP | 1558 | regulation of cell growth | 282 | -1,42 | 1,71E-02 |
| BP | 71900 | regulation of protein serine/threonine kinase activity | 279 | -1,41 | 2,92E-02 |

| | | | | | |
|---|---|---|---|---|---|
| BP | 1902532 | negative regulation of intracellular signal transduction | 350 | -1,41 | 1,74E-02 |
| BP | 10563 | negative regulation of phosphorus metabolic process | 329 | -1,41 | 2,55E-02 |
| BP | 45936 | negative regulation of phosphate metabolic process | 329 | -1,41 | 2,55E-02 |
| CC | 98794 | postsynapse | 375 | -1,41 | 1,67E-02 |
| CC | 98984 | neuron to neuron synapse | 226 | -1,41 | 4,30E-02 |
| BP | 90087 | regulation of peptide transport | 361 | -1,41 | 1,65E-02 |
| BP | 7423 | sensory organ development | 300 | -1,40 | 2,43E-02 |
| BP | 45862 | positive regulation of proteolysis | 243 | -1,40 | 3,44E-02 |
| BP | 98662 | inorganic cation transmembrane transport | 277 | -1,40 | 3,89E-02 |
| BP | 31334 | positive regulation of protein-containing complex assembly | 160 | -1,40 | 4,79E-02 |
| BP | 31400 | negative regulation of protein modification process | 360 | -1,40 | 2,84E-02 |
| BP | 70201 | regulation of establishment of protein localization | 369 | -1,40 | 1,82E-02 |
| BP | 61061 | muscle structure development | 390 | -1,39 | 1,98E-02 |
| BP | 42326 | negative regulation of phosphorylation | 275 | -1,38 | 3,71E-02 |
| BP | 61024 | membrane organization | 449 | -1,36 | 2,46E-02 |
| BP | 51223 | regulation of protein transport | 348 | -1,36 | 4,42E-02 |
| CC | 99513 | polymeric cytoskeletal fiber | 375 | -1,35 | 3,34E-02 |
| MF | 8270 | zinc ion binding | 420 | -1,35 | 3,33E-02 |
| BP | 48598 | embryonic morphogenesis | 368 | -1,35 | 2,67E-02 |
| BP | 72657 | protein localization to membrane | 394 | -1,35 | 3,11E-02 |
| BP | 80135 | regulation of cellular response to stress | 450 | -1,34 | 3,48E-02 |
| BP | 10638 | positive regulation of organelle organization | 411 | -1,33 | 4,93E-02 |
| BP | 51129 | negative regulation of cellular component organization | 484 | -1,32 | 3,32E-02 |
| BP | 43043 | peptide biosynthetic process | 495 | 1,40 | 5,83E-03 |
| BP | 6091 | generation of precursor metabolites and energy | 275 | 1,41 | 1,76E-02 |
| BP | 6412 | translation | 484 | 1,44 | 2,43E-03 |
| MF | 16829 | lyase activity | 107 | 1,45 | 3,66E-02 |
| BP | 7059 | chromosome segregation | 211 | 1,45 | 2,59E-02 |
| BP | 1901657 | glycosyl compound metabolic process | 79 | 1,45 | 4,74E-02 |
| BP | 71824 | protein-DNA complex subunit organization | 129 | 1,46 | 4,58E-02 |
| BP | 6839 | mitochondrial transport | 171 | 1,46 | 2,78E-02 |

| | | | | | |
|---|---|---|---|---|---|
| BP | 7005 | mitochondrion organization | 394 | 1,50 | 3,51E-03 |
| CC | 61695 | transferase complex, transferring phosphorus-containing gr | 182 | 1,50 | 2,20E-02 |
| BP | 1901293 | nucleoside phosphate biosynthetic process | 134 | 1,51 | 2,46E-02 |
| BP | 46390 | ribose phosphate biosynthetic process | 112 | 1,51 | 2,59E-02 |
| BP | 6164 | purine nucleotide biosynthetic process | 108 | 1,51 | 2,99E-02 |
| BP | 9165 | nucleotide biosynthetic process | 133 | 1,52 | 2,51E-02 |
| BP | 9152 | purine ribonucleotide biosynthetic process | 98 | 1,54 | 4,05E-02 |
| BP | 72522 | purine-containing compound biosynthetic process | 112 | 1,54 | 1,89E-02 |
| CC | 120114 | Sm-like protein family complex | 74 | 1,54 | 3,11E-02 |
| CC | 5684 | U2-type spliceosomal complex | 82 | 1,54 | 2,20E-02 |
| BP | 70585 | protein localization to mitochondrion | 83 | 1,55 | 2,44E-02 |
| BP | 51304 | chromosome separation | 61 | 1,55 | 3,72E-02 |
| BP | 9260 | ribonucleotide biosynthetic process | 106 | 1,56 | 2,01E-02 |
| CC | 31970 | organelle envelope lumen | 73 | 1,56 | 2,66E-02 |
| BP | 1902099 | regulation of metaphase/anaphase transition of cell cycle | 41 | 1,57 | 4,31E-02 |
| BP | 8033 | tRNA processing | 87 | 1,57 | 2,29E-02 |
| BP | 30071 | regulation of mitotic metaphase/anaphase transition | 40 | 1,58 | 3,86E-02 |
| BP | 9116 | nucleoside metabolic process | 65 | 1,58 | 3,53E-02 |
| CC | 97525 | spliceosomal snRNP complex | 55 | 1,59 | 4,41E-02 |
| MF | 9055 | electron transfer activity | 67 | 1,60 | 3,47E-02 |
| BP | 9144 | purine nucleoside triphosphate metabolic process | 68 | 1,60 | 3,31E-02 |
| CC | 775 | chromosome, centromeric region | 144 | 1,61 | 1,37E-02 |
| BP | 9141 | nucleoside triphosphate metabolic process | 80 | 1,62 | 2,11E-02 |
| BP | 44784 | metaphase/anaphase transition of cell cycle | 43 | 1,62 | 4,76E-02 |
| CC | 98573 | intrinsic component of mitochondrial membrane | 74 | 1,62 | 1,66E-02 |
| BP | 9199 | ribonucleoside triphosphate metabolic process | 64 | 1,62 | 2,99E-02 |
| BP | 7091 | metaphase/anaphase transition of mitotic cell cycle | 42 | 1,63 | 3,56E-02 |
| BP | 1990542 | mitochondrial transmembrane transport | 73 | 1,63 | 1,46E-02 |
| CC | 1990204 | oxidoreductase complex | 89 | 1,64 | 1,61E-02 |
| BP | 72655 | establishment of protein localization to mitochondrion | 80 | 1,64 | 1,84E-02 |
| BP | 45333 | cellular respiration | 120 | 1,64 | 1,06E-02 |

| | | | | | |
|---|---|---|---|---|---|
| BP | 7080 | mitotic metaphase plate congression | 34 | 1,64 | 4,62E-02 |
| CC | 46540 | U4/U6 x U5 tri-snRNP complex | 29 | 1,65 | 4,19E-02 |
| BP | 6626 | protein targeting to mitochondrion | 63 | 1,66 | 3,10E-02 |
| BP | 61982 | meiosis I cell cycle process | 58 | 1,66 | 1,93E-02 |
| CC | 70069 | cytochrome complex | 24 | 1,66 | 3,18E-02 |
| BP | 7127 | meiosis I | 55 | 1,66 | 2,43E-02 |
| BP | 45841 | negative regulation of mitotic metaphase/anaphase transitio | 26 | 1,66 | 4,81E-02 |
| BP | 6635 | fatty acid beta-oxidation | 54 | 1,67 | 2,85E-02 |
| CC | 5681 | spliceosomal complex | 166 | 1,67 | 1,57E-03 |
| MF | 16251 | RNA polymerase II general transcription initiation factor ac | 23 | 1,67 | 3,08E-02 |
| CC | 5758 | mitochondrial intermembrane space | 65 | 1,67 | 1,89E-02 |
| BP | 6270 | DNA replication initiation | 25 | 1,68 | 3,95E-02 |
| MF | 61650 | ubiquitin-like protein conjugating enzyme activity | 34 | 1,68 | 3,49E-02 |
| MF | 16836 | hydro-lyase activity | 33 | 1,68 | 4,01E-02 |
| BP | 65004 | protein-DNA complex assembly | 99 | 1,68 | 6,82E-03 |
| CC | 32592 | integral component of mitochondrial membrane | 72 | 1,68 | 1,54E-02 |
| MF | 4129 | cytochrome-c oxidase activity | 16 | 1,69 | 4,45E-02 |
| MF | 15002 | heme-copper terminal oxidase activity | 16 | 1,69 | 4,45E-02 |
| MF | 16676 | oxidoreductase activity, acting on a heme group of donors, | 16 | 1,69 | 4,45E-02 |
| BP | 6342 | chromatin silencing | 42 | 1,69 | 2,05E-02 |
| CC | 31304 | intrinsic component of mitochondrial inner membrane | 45 | 1,70 | 2,35E-02 |
| BP | 9201 | ribonucleoside triphosphate biosynthetic process | 50 | 1,70 | 1,70E-02 |
| CC | 31305 | integral component of mitochondrial inner membrane | 44 | 1,71 | 2,49E-02 |
| BP | 33617 | mitochondrial cytochrome c oxidase assembly | 18 | 1,71 | 4,62E-02 |
| MF | 16655 | oxidoreductase activity, acting on NAD(P)H, quinone or si | 27 | 1,71 | 4,47E-02 |
| CC | 22627 | cytosolic small ribosomal subunit | 42 | 1,72 | 1,71E-02 |
| CC | 46658 | anchored component of plasma membrane | 25 | 1,72 | 3,39E-02 |
| CC | 779 | condensed chromosome, centromeric region | 34 | 1,72 | 2,93E-02 |
| CC | 97526 | spliceosomal tri-snRNP complex | 30 | 1,73 | 2,69E-02 |
| CC | 5744 | TIM23 mitochondrial import inner membrane translocase c | 11 | 1,73 | 3,90E-02 |
| BP | 17004 | cytochrome complex assembly | 29 | 1,73 | 2,43E-02 |

| | | | | | |
|---|---|---|---|---|---|
| CC | 71005 | U2-type precatalytic spliceosome | 45 | 1,73 | 1,81E-02 |
| BP | 9205 | purine ribonucleoside triphosphate metabolic process | 61 | 1,73 | 9,95E-03 |
| MF | 4298 | threonine-type endopeptidase activity | 16 | 1,75 | 2,93E-02 |
| CC | 5839 | proteasome core complex | 16 | 1,75 | 2,93E-02 |
| CC | 30880 | RNA polymerase complex | 81 | 1,76 | 6,29E-03 |
| CC | 5689 | U12-type spliceosomal complex | 22 | 1,78 | 2,59E-02 |
| CC | 5669 | transcription factor TFIID complex | 24 | 1,79 | 1,64E-02 |
| MF | 51539 | 4 iron, 4 sulfur cluster binding | 26 | 1,79 | 2,12E-02 |
| MF | 4550 | nucleoside diphosphate kinase activity | 12 | 1,79 | 3,58E-02 |
| BP | 9147 | pyrimidine nucleoside triphosphate metabolic process | 15 | 1,79 | 2,37E-02 |
| MF | 19205 | nucleobase-containing compound kinase activity | 27 | 1,79 | 2,34E-02 |
| BP | 9219 | pyrimidine deoxyribonucleotide metabolic process | 15 | 1,80 | 2,28E-02 |
| CC | 42788 | polysomal ribosome | 28 | 1,80 | 1,45E-02 |
| CC | 428 | DNA-directed RNA polymerase complex | 79 | 1,80 | 2,17E-03 |
| BP | 34508 | centromere complex assembly | 23 | 1,82 | 1,07E-02 |
| CC | 776 | kinetochore | 98 | 1,82 | 2,56E-03 |
| BP | 6220 | pyrimidine nucleotide metabolic process | 30 | 1,82 | 1,15E-02 |
| BP | 9142 | nucleoside triphosphate biosynthetic process | 56 | 1,83 | 7,37E-03 |
| CC | 55029 | nuclear DNA-directed RNA polymerase complex | 78 | 1,83 | 2,14E-03 |
| MF | 70003 | threonine-type peptidase activity | 17 | 1,83 | 2,92E-02 |
| CC | 45277 | respiratory chain complex IV | 14 | 1,83 | 1,92E-02 |
| CC | 5751 | mitochondrial respiratory chain complex IV | 10 | 1,84 | 1,61E-02 |
| CC | 778 | condensed nuclear chromosome kinetochore | 11 | 1,84 | 1,62E-02 |
| BP | 42776 | mitochondrial ATP synthesis coupled proton transport | 14 | 1,84 | 1,90E-02 |
| BP | 9145 | purine nucleoside triphosphate biosynthetic process | 48 | 1,84 | 3,43E-03 |
| BP | 72527 | pyrimidine-containing compound metabolic process | 37 | 1,84 | 9,68E-03 |
| BP | 43486 | histone exchange | 14 | 1,85 | 1,70E-02 |
| BP | 9206 | purine ribonucleoside triphosphate biosynthetic process | 47 | 1,86 | 8,54E-03 |
| CC | 16591 | RNA polymerase II, holoenzyme | 61 | 1,86 | 2,91E-03 |
| BP | 2181 | cytoplasmic translation | 79 | 1,86 | 1,04E-03 |
| MF | 16835 | carbon-oxygen lyase activity | 42 | 1,87 | 4,79E-03 |

| | | | | | |
|---|---|---|---|---|---|
| CC | 5753 | mitochondrial proton-transporting ATP synthase complex | 17 | 1,87 | 2,43E-02 |
| CC | 45259 | proton-transporting ATP synthase complex | 17 | 1,87 | 2,43E-02 |
| CC | 777 | condensed chromosome kinetochore | 23 | 1,89 | 6,54E-03 |
| CC | 71011 | precatalytic spliceosome | 49 | 1,89 | 3,56E-03 |
| CC | 31966 | mitochondrial membrane | 479 | 1,90 | 5,63E-09 |
| BP | 30150 | protein import into mitochondrial matrix | 18 | 1,94 | 7,22E-03 |
| BP | 15985 | energy coupled proton transport, down electrochemical grac | 19 | 1,94 | 1,11E-02 |
| BP | 15986 | ATP synthesis coupled proton transport | 19 | 1,94 | 1,11E-02 |
| MF | 51537 | 2 iron, 2 sulfur cluster binding | 19 | 1,96 | 9,92E-03 |
| CC | 307 | cyclin-dependent protein kinase holoenzyme complex | 36 | 1,96 | 3,17E-03 |
| BP | 16226 | iron-sulfur cluster assembly | 21 | 1,97 | 8,97E-03 |
| BP | 31163 | metallo-sulfur cluster assembly | 21 | 1,97 | 8,97E-03 |
| CC | 19866 | organelle inner membrane | 381 | 1,99 | 5,69E-09 |
| BP | 22904 | respiratory electron transport chain | 62 | 1,99 | 1,08E-03 |
| BP | 22900 | electron transport chain | 64 | 2,01 | 6,14E-04 |
| CC | 15935 | small ribosomal subunit | 73 | 2,03 | 2,75E-04 |
| CC | 314 | organellar small ribosomal subunit | 29 | 2,04 | 1,81E-03 |
| CC | 5763 | mitochondrial small ribosomal subunit | 29 | 2,04 | 1,81E-03 |
| MF | 16538 | cyclin-dependent protein serine/threonine kinase regulator a | 35 | 2,10 | 1,03E-03 |
| CC | 5743 | mitochondrial inner membrane | 352 | 2,11 | 3,14E-10 |
| MF | 51536 | iron-sulfur cluster binding | 44 | 2,11 | 4,67E-04 |
| MF | 51540 | metal cluster binding | 44 | 2,11 | 4,67E-04 |
| BP | 10257 | NADH dehydrogenase complex assembly | 46 | 2,14 | 2,05E-04 |
| BP | 32981 | mitochondrial respiratory chain complex I assembly | 46 | 2,14 | 2,05E-04 |
| BP | 6119 | oxidative phosphorylation | 86 | 2,17 | 1,42E-05 |
| CC | 5747 | mitochondrial respiratory chain complex I | 41 | 2,21 | 8,61E-05 |
| CC | 30964 | NADH dehydrogenase complex | 41 | 2,21 | 8,61E-05 |
| CC | 45271 | respiratory chain complex I | 41 | 2,21 | 8,61E-05 |
| BP | 42773 | ATP synthesis coupled electron transport | 51 | 2,22 | 3,57E-05 |
| BP | 42775 | mitochondrial ATP synthesis coupled electron transport | 50 | 2,23 | 4,02E-05 |
| CC | 315 | organellar large ribosomal subunit | 58 | 2,26 | 2,24E-05 |

| | | | | | |
|---|---|---|---|---|---|
| CC | 5762 | mitochondrial large ribosomal subunit | 58 | 2,26 | 2,24E-05 |
| CC | 22626 | cytosolic ribosome | 95 | 2,34 | 1,01E-07 |
| CC | 22625 | cytosolic large ribosomal subunit | 52 | 2,36 | 3,50E-06 |
| CC | 98803 | respiratory chain complex | 65 | 2,40 | 7,49E-07 |
| CC | 5759 | mitochondrial matrix | 224 | 2,40 | 3,47E-12 |
| BP | 33108 | mitochondrial respiratory chain complex assembly | 77 | 2,42 | 7,31E-08 |
| CC | 5746 | mitochondrial respirasome | 65 | 2,45 | 2,56E-07 |
| CC | 70469 | respirasome | 71 | 2,50 | 1,59E-08 |
| CC | 313 | organellar ribosome | 88 | 2,53 | 1,59E-09 |
| CC | 5761 | mitochondrial ribosome | 88 | 2,53 | 1,59E-09 |
| CC | 98800 | inner mitochondrial membrane protein complex | 113 | 2,54 | 1,40E-10 |
| CC | 15934 | large ribosomal subunit | 111 | 2,61 | 9,77E-12 |
| CC | 5840 | ribosome | 206 | 2,64 | 1,31E-15 |
| CC | 44391 | ribosomal subunit | 179 | 2,71 | 2,52E-16 |
| CC | 98798 | mitochondrial protein-containing complex | 239 | 2,80 | 9,72E-21 |
| MF | 3735 | structural constituent of ribosome | 147 | 2,89 | 5,19E-18 |

**Table S6. Compounds with a transcriptional profile similar to TGFβ-induced *Tks5*$^{+/-}$ lung fibroblasts.**

| NCS | Name | Main Target | Targets | Cell line | PMID | IPF Relevance |
|---|---|---|---|---|---|---|
| 1.760 | GSK-1070916 | Aurora kinase | AURKB\|AURKC\|AURKA\|CYP2D6\|CYP3A4 | BT20 | 32761869 | Inhibition of Aurora Kinase B attenuates fibroblast activation and pulmonary fibrosis |
| 1.733 | PSB-069 | NTPDase | ENTPD1\|ENTPD2\|ENTPD3 | MCF7 | - | - |
| 1.716 | SANT-2 | Smoothened receptor | SMO\|DHH\|IHH\|PTCH1\|SHH | HA1E | - | - |
| 1.697 | entinostat | HDAC | HDAC1\|HDAC2\|HDAC3\|HDAC9 | YAPC | 28315487 | Reversion of TGFb1-induced, XPLN/ SPARK-mediated extracellular matrix turnover in human foetal lung fibroblast cell line |
| 1.695 | pazopanib | VEGFR \|KIT \|PDGFR | KDR\|KIT\|FLT1\|FLT4\|PDGFRB\|PDGFRA\|BRAF\|CYP2B6\|CYP2C8\|CYP2E1\|DDR2\|CSF1R\|FGF1\|FGFR1\|FGFR3\|ITK\|SH2B3 | MCF10A | 21992121<br>27279371 | Tyrosine kinase inhibitor attenuated worsening of lung function, maintained quality of life and reduced acute exacerbations of IPF patients<br>PDGFR inhibition attenuated enhanced differentiation and proliferation of myofibroblasts |
| 1.683 | crizotinib | ALK | ALK\|MET\|CYP2B6\|CYP3A5\|MST1R\|ROS1 | TMD8 | 15563636 | PDGFb1-induced lung fibrosis is blocked by an orally active ALK5 kinase inhibitor |
| 1.671 | emetine | Protein synthesis | RPS2 | A375 | 18503048 | Increase of Sulf1 mRNA levels, a potential negative regulator of TGFb-1 induced fibrogenesis, through inhibition of protein translation |
| 1.667 | TAK-875 | Insulin | FFAR1\|INS | A375 | - | - |
| 1.664 | filgotinib | JAK | JAK1\|JAK2\|JAK3\|TYK2 | XC.L10 | 29440315 | Block of endothelial to mesenchymal transition and artery smooth muscle cell to myofibroblast transition via JAK inhibition thus reducing bleomycin-induced pulmonary fibrosis |
| 1.664 | arofylline | Phosphodiesterase | PDE4A\|PDE4B\|PDE4C\|PDE4D | PC3 | - | - |
| 1.663 | raloxifene | Estrogen receptor | ESR1\|ESR2\|ACVRL1\|ENG | HT29 | 35841004 | Reduction of systemic sclerosis iPSC-derived fibroblast proliferation, production |

| | | | | | | |
|---|---|---|---|---|---|---|
| | | | | | | of extracellular matrix and skin fibrosis in human organoids and mouse model. |
| 1.661 | WH-4023 | Src | LCK\|SRC\|ABL1 | MCF10A | 35998281 | Saracatinib, a Selective Src Kinase Inhibitor, Blocks Fibrotic Responses in Preclinical Models of Pulmonary Fibrosis |
| 1.661 | brefeldin-a | Protein synthesis \|BIG1 | ARFGEF1\|ARFGEF2\|GBF1\|ARF1\|CYTH2 | HCT116 | 8429043 | Inhibition of intracellular degradation and thus secretion of collagen in normal human fetal lung fibroblasts |
| 1.655 | TGX-221 | PI3K | PIK3CB\|PIK3CD | MCF10A | 21984893 | PI3K inhibition attenuates myofibroblast differentiation in the human lungs |
| 1.654 | motesanib | KIT \|PDGFR \|VEGFR | FLT1\|FLT4\|KDR\|KIT\|PDGFRA\|RET | MCF10A | - | - |

*NCS: Normalized connectivity score; PMID: PubMed ID; IPF: Idiopathic pulmonary fibrosis*
